# A genetically encoded local learning rule enables physical learning in engineered bacteria

**DOI:** 10.64898/2026.03.18.712691

**Authors:** Satya Prakash, Clenira Varela, Mark Walsh, Roberto Galizi, Mark Isalan, Alfonso Jaramillo

## Abstract

Training physical neural networks directly in matter remains difficult because most platforms do not store and update weights within the same physical substrate. Here we show that engineered *Escherichia coli* can implement a genetically encoded local learning rule acting on persistent biological memory. We introduce memregulons: bacterial memory elements in which coupled plasmids store an analogue weight as the relative abundance of P1 within the P1+P2 plasmid pool. Environmental inputs activate local promoters, and a shared sublethal kanamycin signal converts promoter activity into differential growth that lowers the stored P1 fraction. In single strains, flow-cytometry trajectories across distinct promoters support the predicted dependence of weight change on learning-channel activity and standing population variance. At the single-cell level, repeated negative learning reshapes the stored weight distribution as it approaches the lower boundary. In mixed populations and co-cultures, one shared negative learning signal selectively rewrites active memregulons, enabling externally routed supervised tic-tac-toe lessons. We then generalise the architecture across orthogonal chemical inputs and combinatorial promoters, and use experimentally measured updates in hybrid analyses of winner-take-all and nonlinear-classifier tasks. These results establish physical learning in living matter through local negative updates of genetically encoded analogue weights.

## Introduction

The energy required to train large artificial neural networks motivates the search for physical neural networks, or PNNs, that compute and learn directly in their substrate^1^. Physical systems can perform inference efficiently, but training remains the central bottleneck. Backpropagation is difficult to implement in matter because it requires a detailed system model and non-local backward transport of error signals. These requirements motivate local physical learning rules, in which each element updates from variables available at that element and a shared training signal^1, 2^. We use the term local learning rule to mean that the update of the stored memory is computed from the local state of that memory and locally available molecular signals. This definition does not require the supervisory or training signal itself to originate inside the same cell. Such rules have been demonstrated in mechanical, photonic and analogue electronic platforms, but not yet as genetically encoded update rules in living cells^1, 2^.

Synthetic biology offers a living substrate for PNNs with potential for self-repair, autonomous operation, and direct interfacing with biological environments. Recently, synthetic gene circuits have advanced beyond basic logic gates to include complex multi-level regulatory circuits^3^, classifiers, and sophisticated neural-network-like architectures in both bacterial^4, 5, 6^ and mammalian^7^ cells. Furthermore, engineered microbial consortia can now distribute computational tasks and share signaling molecules to process complex environmental inputs^8^. Yet, while these neuromorphic circuits demonstrate powerful inference capabilities, they remain fundamentally hard-coded: their parameters are fixed during design or trained *in silico* prior to assembly. True physical learning requires a writable internal memory. Recently, *in vitro* molecular systems, such as DNA strand-displacement networks, have achieved autonomous supervised learning by integrating training data into physical molecular concentrations^9^. In living cells, theoretical frameworks have recently proposed that dense molecular networks could learn environmental statistics without genetic mutations, for example via rate-sensitive autoregulation of hidden molecular species^10^. However, a genetically encoded physical learning rule acting on a persistent analogue memory has not yet been experimentally realized in living matter. Coupled-plasmid memory systems provide a route to tuneable analogue memory by encoding a numerical weight as a plasmid fraction^11, 12, 13, 14^, but these systems have not been connected to a genetically encoded local, activity-gated learning rule.

Physical learning systems require a clear separation between computation, memory and learning. A living-cell circuit can implement a neural-network-like input-output map without learning, because fixed genetic parts, externally imposed inducer concentrations or transient molecular stoichiometries can set effective weights without providing a writable internal memory. By contrast, a physical substrate for learning must contain a memory variable instantiated in living matter, capable of graded states and modifiable by molecular interactions within the system. The memory may reside in a single cell or at the population level. A population-level state qualifies as a learning memory if it is biologically instantiated, persistent across serial culture propagation, writable by local biological interactions and functionally equivalent to an adjustable weight. A key unresolved requirement therefore remains: a genetically encoded local update rule that acts on a persistent, rewritable biological memory variable and thereby provides a physical basis for learning in living matter.

Here, we show that bacterial populations can implement a genetically encoded local negative update rule acting on a persistent plasmid-ratio memory. We engineer this unit as a memregulon: a strain-level memory element in which coupled plasmids P1 and P2 share the same replicon and backbone, and the relative abundance of P1 within the coupled plasmid pool encodes the stored state. For analysis, we report this state as the normalised P1 fraction defined in Eq. (1), where n_P1_ and n_P2_ denote the copy numbers of plasmids P1 and P2.

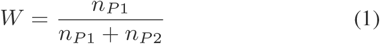

This normalisation reduces sensitivity to changes in total coupled-plasmid copy number. Environmental or Boolean inputs x activate a cognate promoter, fluorescence or promoter activity provides the output/readout y, and sublethal kanamycin K supplies a shared negative learning signal. Because P2 carries the active kanamycin-resistance module, promoter activation converts K into a local growth-rate bias that lowers W selectively in active memregulon populations. We first validate memory stability and the quantitative local update in single strains, and then test the same architecture across orthogonal chemical inputs, combinatorial promoters and co-cultures in which shared kanamycin selectively rewrites active weights in a distributed multicellular setting. We next apply the co-culture system to a winner-take-all tic-tac-toe task to show how negative physical updates can support pruning-based learning when losing branches are externally identified. In this experiment, training is supervised: offline game-tree analysis defines the lesson sequence, the operator routes each lesson to the appropriate cultures, and the cells execute only the local physical update. A complete reinforcement-learning agent would autonomously select actions and update a policy using rewards or punishments^15^. In our supervised design, the operator chooses lessons, supplies outcomes and assigns credit, while the cells execute the local physical update. Finally, we use hybrid task-level readouts to test pruning behaviour by inserting measured physical updates into externally supplied game-tree, winner-take-all and inter-layer computations. The central experimental advance is a genetically encoded local negative update rule acting on persistent biological memory.

## Results

Our main goal is to establish a genetically encoded local negative learning rule in living bacterial populations. We therefore first define the physical update rule in single-strain cultures and then ask whether the same rule remains selective when multiple memregulons share one kanamycin learning signal in mixed populations and in co-cultures. This multicellular setting allows different weights within a culture or co-culture to be updated differently without relying on chemical gradients. We use single-strain measurements to quantify the update dynamics and to parameterise the hybrid analyses used later to probe scalability. Across the experiments, learning refers to changes in stored weights rather than inference alone: in single strains, active kanamycin learning decreases a stored weight; in co-cultures, active targeted memregulons are selectively rewritten; in tic-tac-toe, an externally specified supervised lesson sequence rewrites targeted active memregulons, and measured weights then alter winner-take-all move selection in the external model; and in XOR analyses, negative-only pruning refines suitable initial architectures rather than constructing arbitrary solutions from scratch. Supplementary Notes 1–13 provide, in order, the task design, analytical rule, quantitative validation, flow-cytometry analysis, co-culture protocol, scalability constructs, bulk-weight calibration, plate-reader analysis, stability, learning, co-culture updates, XOR simulations and gating exemplar.

### Analogue memory encoded by a stable plasmid mixture

We first built a biological memory that stores a synaptic weight as a persistent plasmid ratio (Fig. 1a). Each memregulon contains two multi-copy plasmids, P1 and P2, that share the same ColE1 origin and the same ampicillin-resistance backbone. P1 and P2 also carry matched translational resistance fusions to balance plasmid size and expression burden. P1 encodes mCherry together with functional chloramphenicol resistance fused to an inactive kanamycin-resistance module, whereas P2 encodes EGFP together with inactive chloramphenicol resistance fused to the functional kanamycin-resistance module used during learning. This matched design preserves symmetry between the plasmids while conferring a kanamycin-dependent growth advantage only to P2. The local activation and kanamycin-dependent update mechanism is shown in Fig. 1b.

**Fig. 1.**
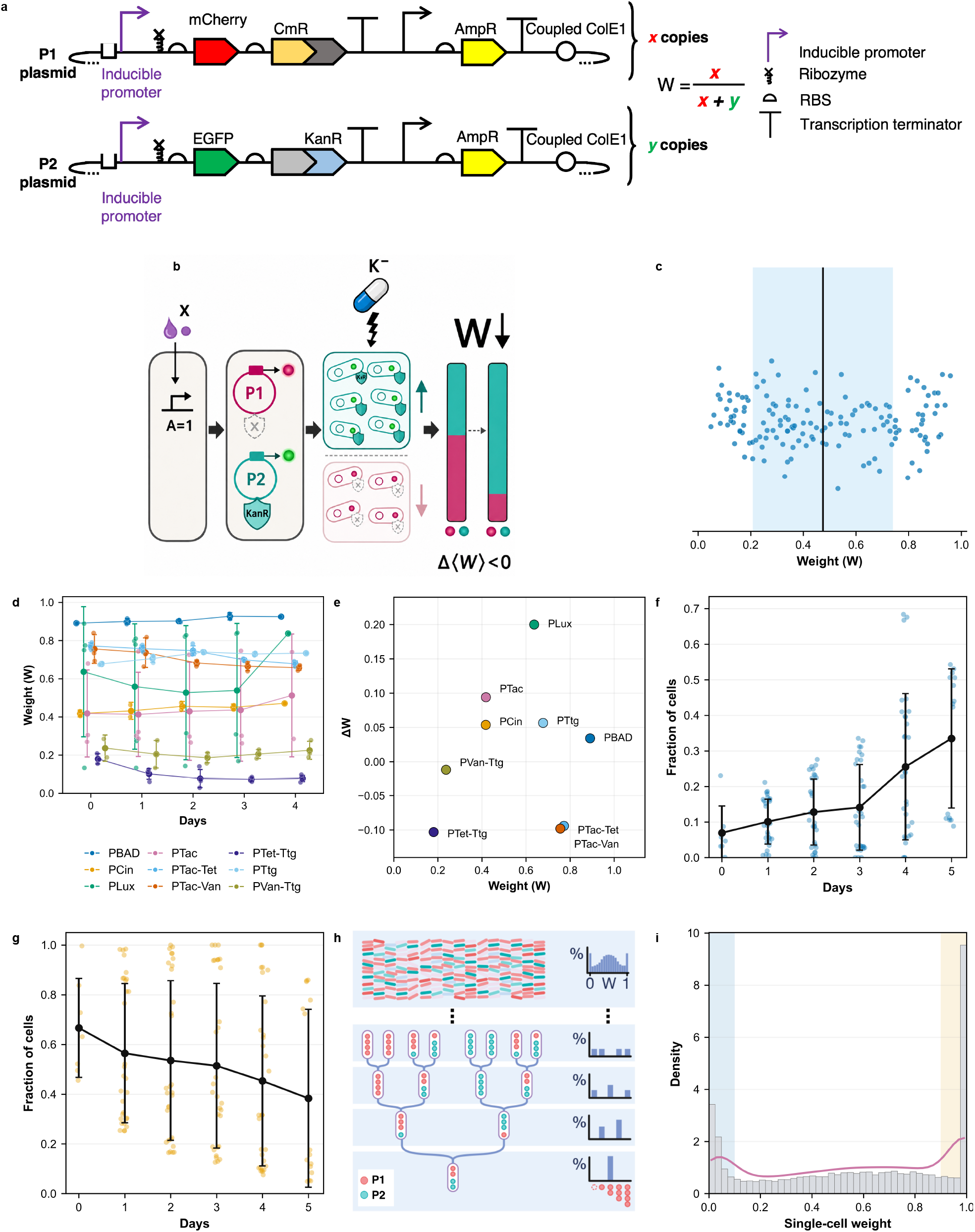
Memregulon architecture, population memory and stability without kanamycin. **a**, Two-plasmid memregulon design. P1 and P2 share a ColE1 origin and ampicillin-resistance backbone. P1 encodes mCherry and functional chloramphenicol resistance fused to an inactive kanamycin-resistance module. P2 encodes EGFP and inactive chloramphenicol resistance fused to the functional kanamycin-resistance module. The stored weight is W = x/(x + y), where x and y are the P1 and P2 copy numbers. **b**, Local activation, readout and kanamycin-dependent negative updating. **c**, Schematic of a stored-weight distribution. The vertical line marks the population mean and blue shading marks its variance. **d**, Stored weight across serial passages without kanamycin. Points are independent biological cultures, lines are means and error bars are s.d. when n > 1. Sample sizes at M0–M4 were PBAD (2/3/2/2/1), PCin (2/2/2/2/1), PLux (4/5/4/4/1), PTac (3/4/3/3/2), PTac-Tet (4/4/4/4/3), PTac-Van (3/3/3/3/3), PTet-Ttg (3/4/4/4/4), PTtg (2/3/2/2/1) and PVan-Ttg (2/3/3/3/3). **e**, Start-to-end drift for nine promoter conditions. Each point is one promoter. **f**, Fraction of cells near W = 0 for 142 independent FACS wells at M0–M5 (7/30/30/30/30/15); symbols show mean ± s.d. **g**, Fraction of cells near W = 1 for 142 independent FACS wells at M0–M5 (7/30/30/30/30/15); symbols show mean ± s.d. **h**, Origin of the interior spread and edge peaks in the inherited weight distribution. Binomial partitioning of P1 and P2 across divisions broadens the interior distribution. Stochastic loss of P1 or P2 fixes the affected lineage at W = 0 or W = 1, respectively, because its descendants inherit only the remaining plasmid. **i**, Representative PBAD-M1 distribution from three independent cultures/wells and 26,958 gated cells. The three-component fit described in Supplementary Note 4 yielded P2-like boundary, P1-like boundary and interior beta fractions of 0.058, 0.296 and 0.646, respectively. Pearson r = 0.975 and two-sided P = 3.7 × 10^-33^. Source data are provided as a Source Data file.

We define the population weight W as the fraction of P1 copies in the total coupled-plasmid pool (Fig. 1c). M-states (e.g., M0, M1, M2) denote discrete stored-memory generations of a memregulon population after defined passaging or learning cycles, and they are not continuous time units. The term day refers to calendar or daily serial-sampling axes, such as those used for stability passaging and the co-culture tournament, and is defined in the relevant figure captions. We confirmed that this weight remains stable over serial dilution in representative single-strain cultures, across the memregulon library, and in co-cultures of distinct memregulons (Fig. 1d–g). Fig. 1h explains why the inherited population contains both an interior weight distribution and peaks at the absorbing states W = 0 and W = 1. Binomial partitioning of P1 and P2 across divisions broadens the interior distribution, whereas stochastic loss of P1 or P2 fixes the affected lineage at W = 0 or W = 1, respectively, because its descendants inherit only the remaining plasmid. Flow cytometry further showed that each memregulon population maintains a broad but reproducible single-cell distribution of weights (Supplementary Note 4). For the representative PBAD-M1 population shown in Fig. 1i, a three-component mixture model captured the empirical weight density well. The model combines empirical boundary components with a beta distribution for interior weights (Supplementary Note 4), supporting the view that the stored memory is a population distribution characterised by a mean and a variance rather than a single deterministic value.

Because revival from a cryogenic stock samples only a finite subset of the stored distribution, independent biological replicates can begin with slightly different mean weights even when they derive from the same frozen population. These differences reflect sampling of pre-existing population heterogeneity rather than stochastic switching during learning. With the stability of analogue memory established, we next investigated how kanamycin-mediated growth bias reshapes this memory and whether the resulting update rule operates locally.

### Growth bias implements local weight updates

We next engineered a mechanism that converts a shared kanamycin learning signal into a persistent change in weight. The key element is the functional kanamycin-resistance cassette on P2. When an environmental or Boolean input x activates the cognate promoter, the promoter activation A(x) drives both the fluorescent/activity readout y and KanR from P2. If we then add the shared kanamycin learning signal K = kanamycin at a sublethal dose, cells with more copies of P2 express more KanR, grow faster and become enriched in the population (Fig. 1b). This adaptive differential growth, coupled to plasmid-ratio dynamics, decreases the population-average fraction of P1 and therefore lowers the stored normalised P1 fraction W in an analogue and persistent manner (Fig. 1b and Fig. 2a).

**Fig. 2.**
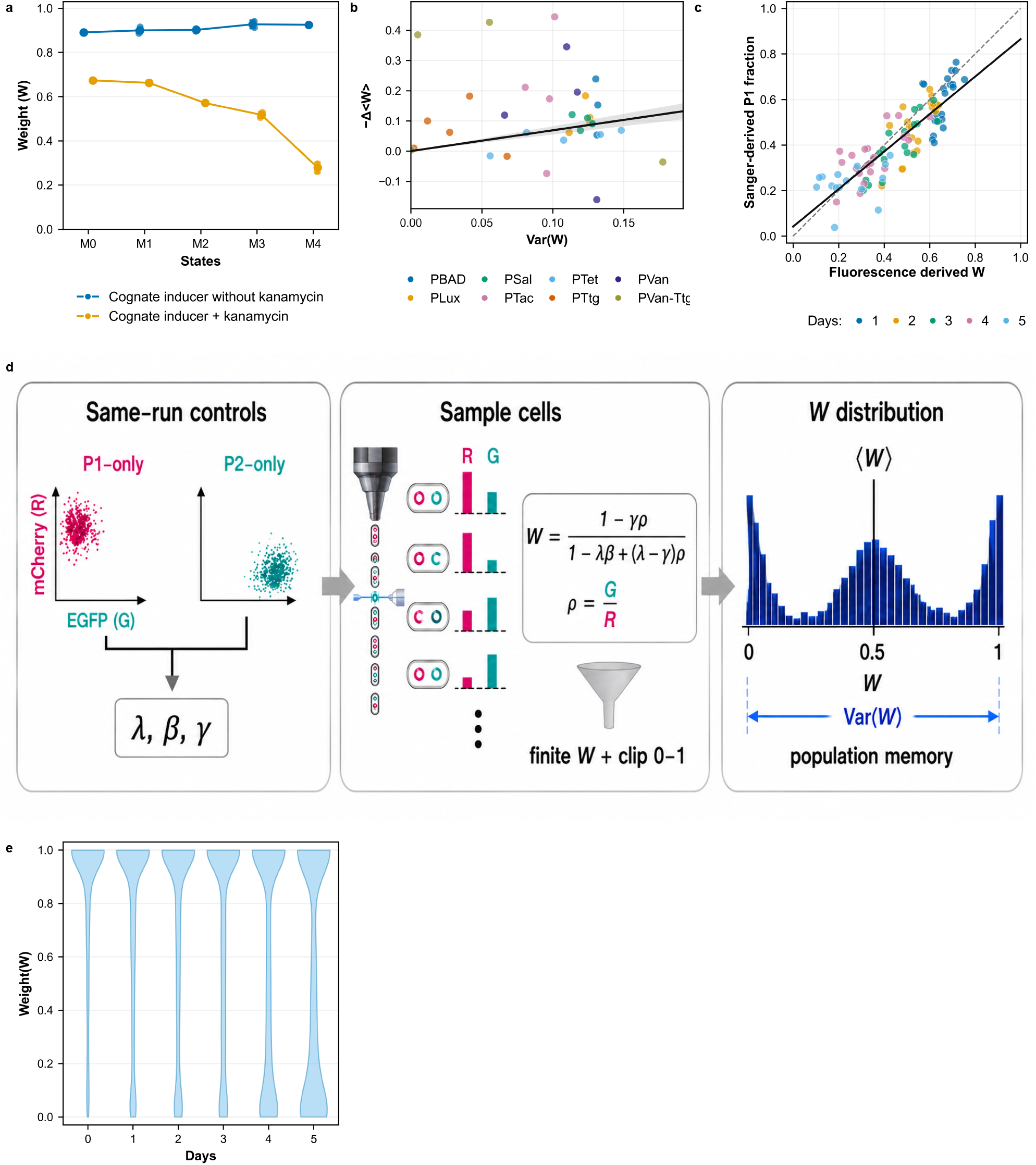
Induction control, learning-rule validation and orthogonal weight measurement. **a**, PBAD induction control across sequential stored states. Blue shows L-arabinose without kanamycin and orange shows L-arabinose plus kanamycin. Points are independent biological cultures, lines are means and error bars are s.d. when n > 1. At M0–M4, n was 1/1/1/2/2 with kanamycin and 2/3/2/2/1 without kanamycin. We use states with n < 3 as trajectory controls and base inferential claims on the full trajectories. **b**, Learning-rule validation for 34 adjacent-state transitions across eight promoters. The black line is the pooled through-origin fit and the grey band is its 95% confidence interval. The pooled inverse-variance-weighted fit was Cneg · A = 0.69 ± 0.07, estimate ± s.e., with 95% CI 0.55–0.83 and two-sided z-test P = 6.8 × 10^-23^. Promoter-level n and two-sided P values were PBAD (4, 0.075), PLux (4, 0.027), PSal (4, 0.0035), PTac (4, 0.028), PTet (5, 0.68), PTtg (5, 0.12), PVan (4, 0.037) and PVan-Ttg (4, 0.82). **c**, Fluorescence-derived W versus Sanger-derived P1 fraction for 96 matched measurements. At M1–M5, n was 21/21/18/21/15. The solid ordinary-least-squares line has slope 0.82 and intercept 0.042, with two-sided P = 3.9 × 10^-28^. The dashed line is identity and Pearson r = 0.85. **d**, Same-run P1-only and P2-only controls define λ, β and γ. The analysis converts gated-cell fluorescence to W, clips W to [0,1] and summarises the result as the population weight distribution. **e**, Single-cell W distributions across M0–M5 pooled across the measured memregulon library. The cytometer acquired up to 50,000 events per well, and the analysis used every gated event. For visual clarity, the panel displays 200 gated-cell measurements from each of 142 independent FACS wells (28,400 displayed cells in total). Source data are provided as a Source Data file.

This design makes the update local at the level of each memregulon population. Different populations can receive the same shared kanamycin learning signal, but only those activated by their cognate inputs produce enough KanR to substantially change their composition (Fig. 1b and Fig. 2a,b). The experimenter specifies the lesson context and, when required by the supervised protocol, selects the culture or co-culture to train. Within that culture, each population’s activation state and regulatory wiring determine its numerical response to the shared kanamycin learning signal. This physical negative update lets a supervised pruning workflow reduce the future influence of active weights associated with incorrect or costly outcomes. The present biological mechanism implements only weight decreases.

A simple growth-competition model predicts the update rule in Eq. (2), where A(x) denotes promoter activation by input x, Var(W) is the standing variance of the stored weight distribution, and Cneg captures the strength of the kanamycin-induced growth bias (Supplementary Note 2).

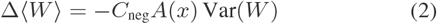

The rule has two important consequences. First, only active populations update. Second, populations with broader weight distributions update more strongly, which provides an intrinsic annealing mechanism because adaptation slows as the distribution narrows. Negative learning on the bounded interval [0,1] should also reshape the full single-cell distribution, progressively increasing its skewness as mass accumulates near the lower boundary. We next tested these predictions experimentally.

We tested this prediction across sequential adaptation trajectories for eight memregulon promoters under kanamycin (Fig. 2a,b). Average weight decreased over repeated cycles only when inducer and kanamycin were both present. To test the predicted dependence on standing variance, we quantified step-to-step changes in mean weight for each promoter and fitted weighted regressions of − Δ ⟨*W*⟩ against Var(W), constrained through the origin, as described in Supplementary Note 3. Seven of eight promoter-specific slopes were positive. The two-sided P values were PLux, 0.027, PSal, 0.0035, PTac, 0.028, PVan, 0.037, PBAD, 0.075, PTet, 0.68, PTtg, 0.12, and PVan-Ttg, 0.82. Inverse-variance weighting across promoters yielded a pooled learning coefficient Cneg · A = 0.69 ± 0.07 (estimate ± s.e., 95% CI 0.55–0.83, n = 34 total learning steps across eight promoters, two-sided z-test P = 6.8 × 10−23). While promoter-specific slope estimates carry high uncertainty because of the limited number of consecutive learning steps per promoter (n = 4–5), the pooled analysis strongly rejects the null hypothesis of no variance dependence. These data therefore quantitatively support the theoretically predicted proportionality between update magnitude and the standing variance of the stored distribution. As an additional molecular check, a limited qPCR time course on plasmid DNA from consecutive kanamycin-learning stages showed the same directional decrease in P1 fraction as the fluorescence-derived weight trajectory. We reserved this single-promoter dataset for directional orthogonal comparison and estimated the pooled Cneg · A regression exclusively from the full eight-promoter fluorescence dataset (Supplementary Fig. 2a). A matched qSanger chromatogram analysis^13^ is provided as an orthogonal comparison of fluorescence-derived W with Sanger-derived P1 fraction (Fig. 2c and Supplementary Note 7). Fig. 2d shows the same-run controlto-weight workflow. Detailed promoter-specific trajectories and the full distributional evolution are provided in Supplementary Note 3 and Supplementary Fig. 37.

We then asked whether the same mechanism remains selective in a mixed population. In a co-culture containing two memregulon strains with orthogonal inducers, only the strain activated by its cognate inducer changed weight during kanamycin exposure (Supplementary Note 4). At the single-cell level, the distribution pooled across the measured memregulon library shifted towards lower W as learning progressed (Fig. 2e and Supplementary Note 3). These results define a physical local learning rule that acts on active populations and whose magnitude depends on their stored variance.

### Co-culture use case for the local learning rule

We next used nine-strain co-cultures as a multicellular test bed for the local learning rule in a setting where different weights share one kanamycin learning signal without relying on chemical gradients. We used a simplified tic-tac-toe tournament as a compact decision-tree test bed (Fig. 3a,b). In this setting, player X always opens in the centre square. A separate memregulon co-culture represents each available move for player O. Board positions denote abstract task states, while the co-cultures remain physically separate. In Fig. 3b, the numbered key gives the experimenter-defined correspondence between X-board positions and chemical inducers. For each O decision, the experimenter supplies every candidate-O co-culture with the inducers corresponding to positions occupied by X. Within a co-culture, an inducer activates only memregulons carrying its cognate promoter. For example, OC6 activates PLux wherever a PLux memregulon is present. The experimenter therefore determines which weights respond by choosing the memregulon composition of each candidate-O co-culture. The analysis converts red and green fluorescence into calibrated weights, and an external winner-take-all (WTA) model computes a weight-derived move score for each available position. The model selects the available position with the highest score. Supplementary Table 2 lists the constituent promoter-defined memregulons and starting states for each candidate-O co-culture. Supplementary Table 38 gives the Day-0 co-culture compositions, and Supplementary Note 5 gives the lesson-by-lesson inducer schedule.

**Fig. 3.**
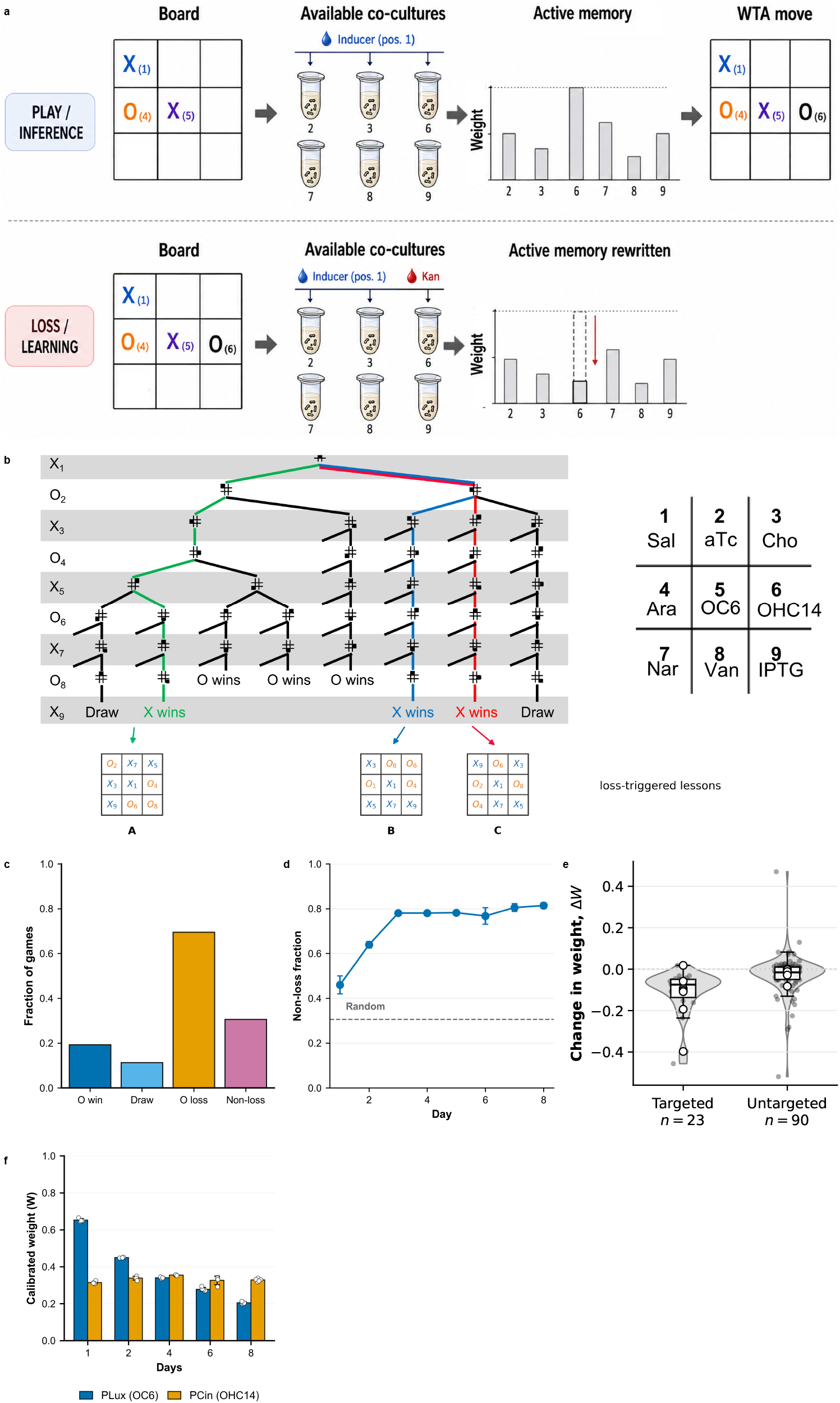
Externally routed supervised tic-tac-toe lessons in co-culture. **a**, Experimental workflow. The experimenter supplies the board context and lesson identity. Cells execute the local physical update, and an external winner-take-all rule selects moves. **b**, Hybrid lesson tree with loss-triggered supervised boards A–C and inducer key. The numbered key gives the experimenter-defined correspondence between X positions and inducers. A separate co-culture represents each candidate O move, and its composition determines which constituent memregulons respond to the supplied inducers. For example, OC6 activates each PLux memregulon present in that co-culture. Supplementary Tables 2 and 38 give the initial compositions. **c**, Fixed-first-move random baseline from 20,000 independent simulated games. There were 3,847 O wins, 2,259 draws and 13,894 O losses, giving 6,106 non-losses. Error bars are 95% Wilson binomial confidence intervals. All half-widths are ≤ 0.007 and are therefore narrower than the bar outlines. **d**, Non-loss fraction after inserting each day’s measured weights into the fixed game-tree readout. Points and error bars show the mean and s.d. across 100 Monte Carlo uncertainty-propagation evaluations per day. The dashed line is the random baseline. **e**, Targeted and untargeted weight changes. Grey violins show transition-level density and black points show 23 targeted and 90 untargeted transitions. Boxes show the median, interquartile range and 1.5×-interquartile-range whiskers. Open circles show means for each of seven paired board-position series. The seven paired series are the independent units. The seven series contain the 23 targeted and 90 untargeted transitions. Across the seven paired series, the mean targeted-minus-untargeted change was −0.11, with paired-bootstrap 95% CI −0.19 to −0.044 and two-sided exact paired sign-flip permutation P = 0.031 across 128 assignments. **f**, Position-4 PLux under OC6 and PCin under OHC14. Bars are means across independent biological cultures, overlaid points are the individual cultures and error bars are s.d. At days 1/2/4/6/8, n was 4/4/4/5/5 for PLux and 4/4/4/4/5 for PCin. Source data are provided as a Source Data file.

The experimental co-culture tic-tac-toe training is supervised learning. Before the wet-lab experiment, we used offline computational game-tree analysis to define seven training lessons from experimentally measured single-strain update rules (Supplementary Note 1). We used reinforcement-learning-style branch pruning only in that offline computational lesson-design step, where simulated games identified losing branches to prune. During the experiment, the operator followed the fixed lesson sequence, supplied the board-context inducers, trained the culture or co-culture representing the active losing branch and propagated all other positions without kanamycin (Fig. 3a,b). Thus, the co-culture experiment tests whether local activation selectively rewrites active weights within a real multi-strain population under one shared kanamycin signal and external supervision. Supplementary Note 5 gives the full daily protocol for the co-culture supervised lesson sequence. Fig. 3c,d show the fixed-first-move random baseline and measured-weight non-loss trajectory, respectively.

Across seven board-position co-culture series containing both targeted and untargeted transitions, the mean paired targeted-minus-untargeted change was −0.11 (bootstrap 95% CI −0.19 to −0.044, two-sided exact paired sign-flip permutation test P = 0.031, Fig. 3e). The plot also shows all 23 targeted and 90 untargeted transitions grouped within the seven independent series. This effect size and interval support selective rewriting of active memregulons by one shared kanamycin learning signal. At position 4, the OC6-responsive PLux weight decreased during targeted learning, whereas the PCin weight measured under cognate OHC14 remained comparatively stable (Fig. 3f).

We quantified task performance as the non-loss fraction, defined as the fraction of games in which player O either wins or draws. This metric is appropriate here because O plays second after X opens in the centre, making avoidance of loss the relevant outcome for this negative-pruning workflow. For the experimental co-culture data, we inserted the measured post-adaptation weights into the externally specified tic-tac-toe WTA readout described in Supplementary Note 1 and computed the resulting non-loss fraction. The mean ± s.d. across 100 computational uncertainty-propagation evaluations per day increased from 45.99 ± 4.03% on day 1 to 81.44 ± 1.28% on day 8 (Fig. 3d). Thus, the measured physical updates altered the fixed WTA task output across the supervised lesson sequence. Fig. 3d quantifies this task-level consequence through an externally specified WTA readout benchmarked against a simulated random baseline.

### Combinatorial promoters support XOR architecture

Negative local learning is most useful if the architecture can represent non-linear functions. We therefore expanded the library with combinatorial memregulons controlled by two-input AND promoters^16^ and with hybrid memregulons in which *P1* and *P2* carry different promoters (Supplementary Note 6). These constructs obeyed the same physical rule as the single-input memregulons: weight decreased only when the corresponding promoter logic was satisfied. This demonstrates that the local update mechanism is highly modular. It can be attached to complex regulatory elements across nine orthogonal chemical inputs, including quorum-sensing channels relevant to future cell-cell communication.

As a proof of principle, we implemented XOR by rational design using a one-stage WTA readout (Fig. 4a,b). To achieve this, the promoter layer (Fig. 4a) comprised a bias memregulon, two single-input memregulons whose pooled output acts as an OR-type gate, and one AND-type combinatorial memregulon. By physically extracting the nonlinear x1 AND x2 feature upstream of the decision step, this ensemble provides the expanded feature space required to solve XOR. Unlike a conventional single linear layer, the necessary nonlinearity is supplied directly by the engineered promoter channel. Consequently, the subsequent WTA readout can mathematically separate the classes, yielding the correct deterministic output for all four truth-table rows. Propagated weight uncertainty, correct-output advantages, and Monte Carlo correct-label fractions are reported in Fig. 4b and Supplementary Note 12.

**Fig. 4.**
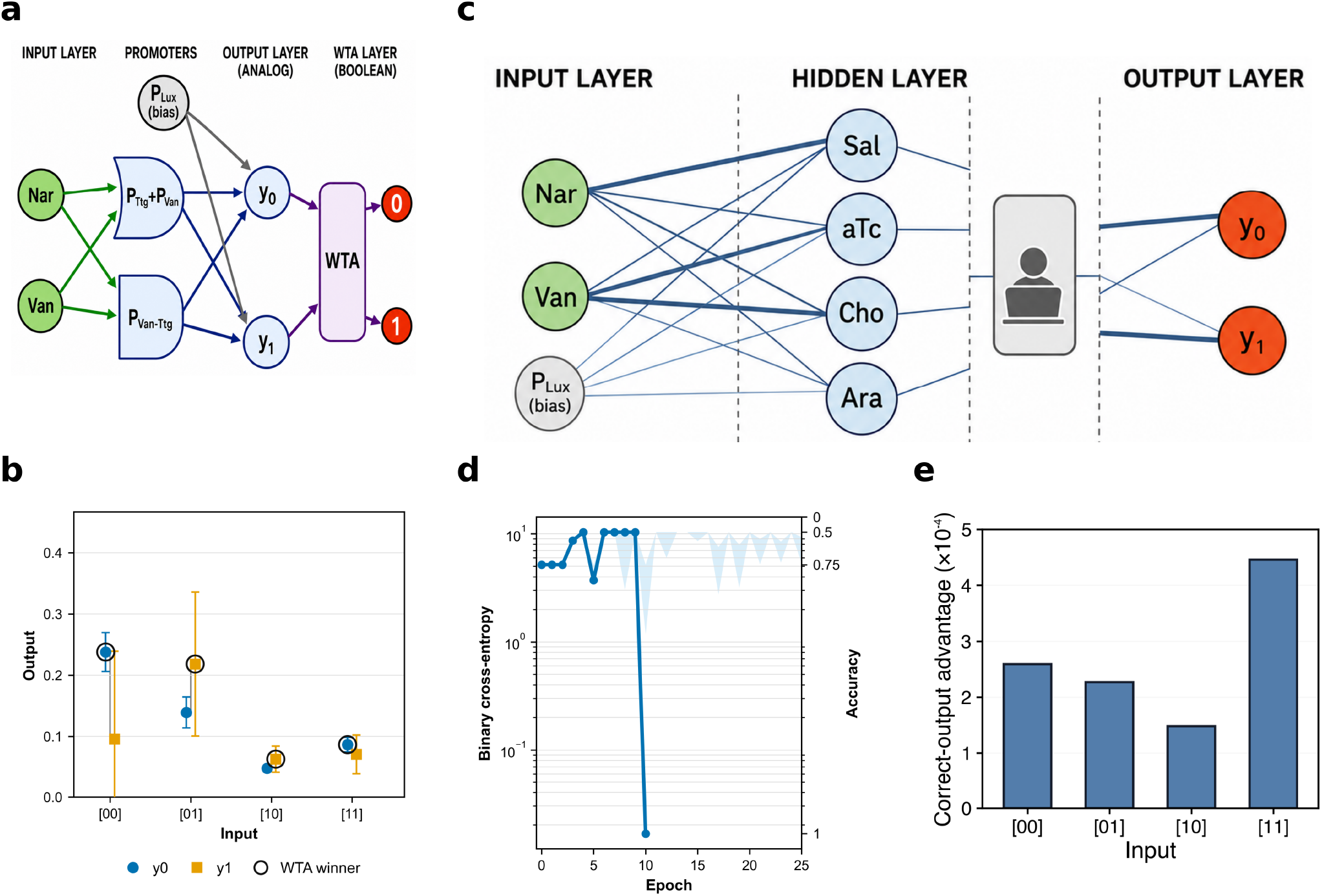
Computational XOR analyses using a physical nonlinear feature and human-in-the-loop multilayer scalability. **a**, One-stage WTA readout with an explicit physical combinatorial-promoter feature. **b**, Deterministic truth-table readout. Blue circles show y0, orange squares show y1 and open circles mark the WTA winner. Error bars propagate one s.d. of promoter-specific weight uncertainty. For input 10, OC6 = 10 µM, Nar = 1,000 µM and Van = 0 µM give x1 AND x2 = 0, y0 = 0.048 and y1 = 0.063, so WTA assigns class 1. From 5,000 independent parameter draws per input, the one-stage correct-label fractions were 0.91, 0.88, 0.83 and 0.69 for 00, 01, 10 and 11. **c**, Human/operator/computational multilayer workflow. The operator/computational workflow performs inter-layer routing and WTA decisions. **d**, Binary cross-entropy and truth-table accuracy during negative-only training. The reference run converged after 10 misclassification-triggered updates. Across 100 seeded shuffled-order runs, 45 converged. The band is the 10th–90th percentile of the 55 non-converged runs. **e**, Correct-output advantage, scaled by 10^4^. The deterministic values were 2.60, 2.27, 1.48 and 4.46 for 00, 01, 10 and 11. From 5,000 independent parameter draws per input, the multilayer correct-label fractions were 0.52, 0.49, 0.48 and 0.51. Source data are provided as a Source Data file.

### Human-in-the-loop analyses probe scalability

We next asked whether the same negative local learning rule could support human-in-the-loop multilayer scalability analyses when inter-layer communication is supplied externally. To physically isolate the learning principle from current limits in biological signalling, we used a human-in-the-loop protocol in which each co-culture was treated as one neuron. During inference, pooled fluorescence from each co-culture was converted, through the experimentally calibrated transfer functions, into the inducer concentration applied to the next layer. During learning, the incorrect output well and the active upstream wells that contributed to that decision were exposed to kanamycin under the same input context, so that each weight update remained local and physically implemented, whereas communication between layers was supplied externally (Supplementary Note 12). This protocol tests multilayer computation while confining the biological implementation to the local weight update.

XOR provided a stringent test because it cannot be implemented by a single linear layer without an additional nonlinear feature. In this hybrid framework, a network with two inputs, one bias, four hidden co-cultures and two output co-cultures reached a deterministic XOR solution by negative-only learning from a biologically suitable OR-like initial state (Supplementary Note 12). Fig. 4c shows the human/operator/computational workflow used for the multilayer scalability analysis, and Fig. 4d shows the ordinary binary crossentropy and truth-table accuracy during training. After learning, the deterministic multilayer network produced the correct WTA decision for all four XOR inputs. To show this directly, we plotted the correct-output advantage, defined for each truth-table row as the target-output score minus the non-target-output score (Fig. 4e). All four margins were positive, indicating correct deterministic XOR classification, although the small margins and the Monte Carlo analysis in the Supplementary Information show that this implementation remains sensitive to propagated biological weight uncertainty. Together with the one-stage WTA readout in Fig. 4a,b, these analyses show two constrained routes to XOR: rational one-stage WTA design with a physical nonlinear feature expansion, and multilayer negative learning in a human-in-the-loop setting. In both cases, deterministic WTA outputs implement XOR. The Monte Carlo results identify signal-to-noise engineering as the next requirement for robust execution. These multilayer results define the scope of the present claim precisely: weight updates are physically implemented and constrained by measured memregulon dynamics, whereas interlayer communication, routing and WTA decisions remain externally supplied by the operator/computational workflow (Supplementary Note 12). The result is a human-in-the-loop scalability analysis.

## Discussion

Taken together, the experiments establish a biological route to local physical weight updating. The rule is implemented in the substrate itself: a memregulon population stores a weight as a plasmid-fraction distribution, local activity gates expression of kanamycin resistance in the same cells, and one shared kanamycin learning signal rewrites only the active weights through sublethal differential growth and plasmid-ratio rebalancing. This mechanism works in single-strain cultures, in mixed populations and in genuine multistrain co-cultures. The result is a physical negative local update rule embedded in living cells and available to higher-level supervised pruning workflows. The experimenter supplies action selection and outcome assignment, and the biological mechanism implements weight decreases.

The rule is local because the update of each weight depends only on variables available at that weight: the current plasmid distribution, the local activation state, and the shared negative learning signal. The experimenter supplies the supervised lesson context and shared signals, while each population’s local biological dynamics determine its numerical weight change. This places the system within the broader paradigm of gradient-free in situ training in physical neural networks^1, 2^ while also defining its limits. The implemented negative update prunes active weights after an unfavourable signal. While autonomous physical learning has recently been demonstrated in cell-free DNA networks^9^, our memregulon system extends a constrained form of local learning into living cells. Importantly, the mechanism relies on population-level differential growth rather than intracellular molecular remodelling.

A useful feature of the mechanism is that the update magnitude scales with the standing variance of the stored distribution. Broad populations adapt quickly, whereas narrow populations resist further change. The substrate therefore carries its own adaptive learning rate: early updates are exploratory, and later updates slow automatically when the variance decreases. Supplementary Note 3 supports this prediction empirically across eight promoters by showing a positive relationship between step-to-step weight change and standing variance, with a strongly significant pooled estimate of the learning coefficient. At the same time, the single-cell distributions do not simply translate leftwards during learning. They also narrow and become increasingly skewed as weights approach the lower boundary, which is the expected signature of repeated negative learning acting on a bounded analogue memory.

The experiments operate at different abstraction levels. Single-strain and mixed-population assays validate the physical update itself; co-culture tic-tac-toe tests whether externally routed lessons can target the appropriate active cultures; and the XOR analyses test how negative-only pruning behaves in constrained WTA architectures. Across these examples, the biological component is the same local plasmid-ratio update, whereas the surrounding task logic is supplied externally. Thus, collective error reduction in the present work means that specified readouts combine several stored weights and that pruned branches become less likely to win future decisions within the fixed task readouts.

Our results demonstrate that, in this system, computation arises from the collective input-output behaviour of engineered bacterial populations, memory resides in a persistent biological state encoded in the living substrate, and learning occurs when local molecular interactions modify that stored state according to a genetically encoded rule. This implementation highlights the importance of scalable local plasticity mechanisms that couple stored biological state to local signals reporting activity, context and error.

The multilayer XOR analysis illustrates how this principle could extend to future human-in-the-loop scalability analyses. In the present work, externally supplied signalling allowed us to separate the local biological update rule from the current limits of biological communication and thereby enable an architectural scalability test (Supplementary Note 12). All weight updates followed experimentally grounded biological dynamics, whereas inter-layer communication was supplied by the human/operator/computational workflow. As synthetic multicellular communication capabilities improve, this same design logic could support persistent weights linked by biological signals. The OC6- and OHC14-responsive channels used here show that AHL-responsive promoters can gate memregulon activity like the other chemical inputs, supporting a future route in which local activation A(x) is supplied by neighbouring cells rather than by externally added inducer. Direct cell-cell signalling during learning is the next step towards an autonomous communicating learning population.

The present implementation has important limits. The learning rule is strictly negative, so it can only decrease weights and therefore operates in a carve-down regime. It cannot implement a general algorithm that is guaranteed to minimise prediction error in arbitrary tasks, because many such algorithms require coordinated weight increases as well as decreases. Nevertheless, negative-only learning can be useful when the task can be framed as pruning or suppressing wrong pathways: eliminating incorrect or high-cost actions from a winner-take-all decision set, applying delayed negative updates after losing lessons, selecting against active regulatory pathways that produced an undesired outcome, or refining a near-solution architecture such as XOR by reducing inappropriate outputs. These use cases do not require the system to know which alternative action should be strengthened; they require only that active pathways associated with a poor outcome become less likely to win future competitions. The finite dynamic range of plasmid-ratio weights further limits the size of decision trees that can be explored by repeated negative updates. Because *W* is non-negative and bounded by finite plasmid copy number, each decrement consumes writable range; with copy numbers of order tens, even one-copy changes are percent-scale, and experimentally resolvable ratiometric changes are likely larger. The practical number of useful pruning steps is therefore smaller than the absolute copy-number bound. The present system is best viewed as a physical substrate for short pruning trajectories, small decision trees and near-solution refinement, not as an indefinitely deep optimiser. Prolonged negative-only training can eventually drive weights towards collapse. The boundaries W=0 (P2-only) and W=1 (P1-only) act as absorbing states for the plasmid-ratio memory; once a lineage has effectively lost one plasmid, ordinary neutral partitioning cannot recreate it. This produces boundary accumulation, including U-shaped distributions when both edges are populated, and repeated negative-only updates consume writable range. The maximum number of useful pruning steps is therefore bounded unless reset, fusion, or bidirectional update mechanisms are introduced. To mitigate this, we computationally and experimentally explored a biologically implementable memregulon fusion operation (Supplementary Fig. 2b and Supplementary Note 1): physically mixing adapted populations with a reference culture. This operation pulls extreme weights back towards intermediate values while preserving their relative ordering, thereby replenishing population variance and extending the useful lifetime of sequential decision tasks.

Looking forward, tasks that require coordinated increases of selected connections or exact credit assignment across hidden layers will need bidirectional update rules. In neuromorphic hardware, bidirectional updates are often achieved through contrastive learning or Equilibrium Propagation (EqProp)^2, 17, 18^. Executing EqProp-like algorithms in living matter would require bidirectional, energy-based communication between cells. While traditional chemical signaling is relatively slow and unidirectional, future implementations could bridge this gap by using engineered extracellular electron transfer to form conductive cellular networks^19, 20, 21^. Currently, we use antibiotics to impose growth bias, which determines the adaptation timescale, may limit host compatibility and could favour mutational escape. Orthogonal actuators, including CRISPR-based or non-antibiotic growth-control schemes^11^, may provide faster or more portable implementations.

Several engineering challenges remain. Larger architectures will require explicit communication modules, control of metabolic burden, automated handling of long training protocols and bidirectional update mechanisms for tasks that require coordinated strengthening as well as pruning. Direct closed-loop learning in richer multicellular tasks without simulation-guided lesson design is a major next step. Even so, the present study shows that one principle of neural computation local activity-dependent weight update from a shared training signal - can be encoded genetically and executed by living populations through differential growth, while keeping the present claim focused on the experimentally demonstrated local negative update.

## Methods

### Strains and growth conditions

We grew cells aerobically at 37 °C in 14 mL tubes unless stated otherwise. We used LB medium (1% tryptone, 0.5% yeast extract, 1% NaCl), LB agar plates (1.5% Select agar, Sigma Aldrich), or M9 medium (1× M9 salts (Sigma Aldrich), 100 µM CaCl2, 2 mM MgSO4, 10 µM FeSO4 and 0.8% v/v glycerol) supplemented with 0.2% w/v casamino acids, 1 µg/mL thiamine, 20 µg/mL uracil and 30 µg/mL leucine; pH was adjusted to 7.4 with NaOH. For cloning we used *Escherichia coli* TOP10 [F-mcrA Δ(mrr-hsdRMS-mcrBC) Φ80lacZΔM15 ΔlacX74 recA1 endA1 araΔ139 Δ(ara, leu)7697 galU galK λ-rpsL (StrR) nupG] grown in LB at 37 °C and 200 rpm with appropriate antibiotics. For all experiments we used Marionette DH10B (Addgene bacterial strain #108251). We cured its native chloramphenicol-resistance cassette by electroporating 60 ng of pE-FLP plasmid^22^ (Addgene #45978) and selecting on 50 µg/mL ampicillin agar at 30 °C for 1 day. We used the cured strain for all memregulon adaptations and characterisations. To recover cultures from glycerol stocks, we inoculated 2 mL LB with 100 µg/mL ampicillin and grew cultures overnight at 37 °C and 200 rpm. Default antibiotic concentrations were carbenicillin 80 µg/mL, kanamycin 50 µg/mL, ampicillin 100 µg/mL and chloramphenicol 34 µg/mL

### Plasmid construction

We generated all plasmids by Gibson assembly^23^ using synthetic DNA fragments from IDT and PCR products amplified with Q5 DNA polymerase (NEB). P1 and P2 were designed to be identical in size (4,774 bp) and to share a common backbone containing a pMB1 origin of replication, an ampicillin-resistance cassette from pSEVA191^24^ and a spy terminator^25^. We amplified the pMB1 replicon, a ColE1 derivative, from pWT018e^11^ and introduced a T-to-C mutation in RNAII to reduce copy number^26, 27^. Into this backbone we inserted an inducible promoter, a RiboJ insulator^28^ and the weak ribosome-binding site BBa_B0064. In P1, the promoter drove mCherry from PAJ310^29^, with a C-terminal Gly-Ser-Gly extension to match EGFP size. In P2, it drove EGFP from pWT018a. Downstream, we added matched translational resistance fusions. The inactive resistance modules carried active-site-inactivating mutations and served to equalise construct architecture and expression burden. P1 contained functional chloramphenicol resistance fused through a (Gly-Ser)3 linker to inactive KanR carrying the Asp208Ala substitution, whereas P2 contained inactive chloramphenicol resistance fused to functional KanR. Plasmids generated in this study are available upon request from Alfonso Jaramillo.

### Memregulon library construction

The 9YES library comprised nine matched P1/P2 plasmid pairs, each carrying one inducible promoter: PSalTTC (PSal; sodium salicylate), PTet* (PTet; anhydrotetracycline HCl), PBetI (PBet; choline chloride), PBAD (L-arabinose), PLuxB (PLux; 3OC6-AHL), PCin (3OHC14:1-AHL), PTtg (naringenin), PVanCC (PVan; vanillic acid) and PTac (IPTG). The 4AND library comprised four two-input promoters^16^: PTet-Ttg, PVan-Ttg, PTac-Tet and PTac-Van. We built these promoters by introducing a second orthogonal Marionette operator site into an existing inducible promoter so that both cognate repressors had to be relieved for activation. The 3HYB library comprised hybrid memregulons formed by co-transforming a P1 plasmid from one 9YES member with a P2 plasmid from another. The three hybrids used here were PBAD:PSal, PTet:PBAD and PTet:PVan.

### Memregulon transformation

To create a memregulon strain, we co-transformed equimolar amounts of P1 and P2 (80 ng each) into cured Marionette DH10B. We selected transformants on LB agar containing ampicillin (60 µg/mL), kanamycin (17.5 µg/mL), chloramphenicol (11 µg/mL) and the cognate inducer or inducer pair. Inducer concentrations for selection were: Sal 60 µM, aTc 0.1 µM, choline 6,000 µM, arabinose 2,000 µM, OC6 6 µM, OHC14 5 µM, naringenin 250 µM, vanillic acid 40 µM and IPTG 250 µM. For two-input constructs we used the corresponding inducer pairs. For hybrids carrying different promoters on P1 and P2, we added both cognate inducers during selection. We picked single colonies, grew them overnight in ampicillin and stored glycerol stocks at -80 °C.

### Memregulon stability and learning

For stability measurements, we inoculated cultures from glycerol stocks into LB plus ampicillin, grew them overnight and passaged them daily for 5 days. We prepared a new glycerol stock from each day’s culture. For learning experiments, we diluted overnight cultures 1:200 into 1.5 mL fresh M9 medium containing carbenicillin, the cognate inducer(s) and a promoter-specific sublethal kanamycin concentration: PSal 1.5 µg/mL, PTet 2.5 µg/mL, PBetI 4 µg/mL, PBAD 2.5 µg/mL, PLux 1 µg/mL, PCin 1.5 µg/mL, PTtg 1.5 µg/mL, PVan 3.5 µg/mL, PTac 4 µg/mL, PTet-Ttg 1.5 µg/mL, PVan-Ttg 1.5 µg/mL, PTac-Tet 1.5 µg/mL and PTac-Van 1.5 µg/mL. Inducer concentrations were: Sal 100 µM, aTc 0.2 µM, choline 10,000 µM, arabinose 4,000 µM, OC6 10 µM, OHC14 7 µM, naringenin 1,000 µM, vanillic acid 100 µM and IPTG 500 µM; two-input constructs received the corresponding inducer pairs. After dilution, we split each culture into three independent lineages, incubated them for 8 h at 37 °C and 200 rpm, and then prepared glycerol stocks. We repeated the cycle four or five times. For co-culture learning, we first grew individual memregulon cultures overnight, mixed equal volumes, stored half of the mixture as a glycerol stock and diluted the other half 1:200 into LB containing carbenicillin, the relevant inducer(s) and kanamycin.

### Memregulon fusion

Memregulon fusion merges two cultures of the same memregulon at different weights. We inoculated two glycerol stocks of the same memregulon, grew them overnight in ampicillin, mixed 200 µL of each culture, and prepared a new glycerol stock from the mixture. A 1:1 fusion with a reference culture of weight 0.5 increases weights below 0.5 and decreases weights above 0.5, thereby replenishing variance while preserving the relative ordering used in downstream winner-take-all game-play models. We characterised this operation experimentally and used it in extended game simulations as a biologically grounded strategy to mitigate weight collapse during prolonged negative-only adaptation.

### Memregulon characterisation using fluorescence plate reader

We inoculated glycerol stocks of bacteria carrying memregulons and grew overnight with ampicillin (100 µg/mL). We refreshed cultures by diluting 1:2000 into freshly prepared M9 with the cognate inducer(s) and carbenicillin (80 µg/mL). Inducer concentrations were: PSal (Sal 100 µM), PTet (aTc 0.2 µM), PBetI (Cho 10 mM), PBAD (Ara 4 mM), PLux (OC6 10 µM), PCin (OHC14 7 µM), PTtg (Nar 1 mM), PVan (Van 100 µM), PTac (IPTG 500 µM), PTet-Ttg (aTc 0.2 µM, Nar 280 µM), PVan-Ttg (Van 25 µM, Nar 280 µM), PTac-Tet (IPTG 500 µM, aTc 200 µM) and PTac-Van (IPTG 500 µM, Van 21 µM). After 3 h, we dispensed 200 µL from each culture into a 96-well plate (Custom Corning Costar) with technical and biological replicates. We acquired data in Tecan Magellan on an Infinite F500 (serial 1402001069) at 37 °C with shaking. The Infinite F500 recorded OD_600_ and fluorescence every 15 min for 18 h using a 600 nm absorbance filter, 465/35 nm excitation and 530/25 nm emission for EGFP, and 580/20 nm excitation and 635/35 nm emission for mCherry. We also acquired data in Tecan Magellan on an Infinite 200 PRO (serial 1604000172) at 37 °C with orbital shaking. In those exports, the OD600-labelled measurement used 595 nm absorbance with a 10 nm bandwidth; EGFP used 485/20 nm excitation and 535/25 nm emission, and mCherry used 590/20 nm excitation and 625/35 nm emission.

### Fluorescence plate reader data analysis

We analysed plate-reader data with custom Python scripts. We synchronised time series at OD_600_ = 0.2, linearly interpolated fluorescence and OD_600_ at intermediate times, fitted exponential growth rates and discarded poor fits (r^2^ < 0.9). We reported fluorescence as the slope of fluorescence versus OD_600_ and converted ratiometric red and green signals into population-average weights using the calibration derived in Supplementary Note 7. We calculated weight using Eq. (3).

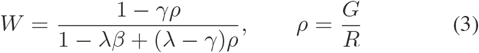

Here, ρ = G/R; RP1, GP1, RP2 and GP2 denote total red and green fluorescence in P1-only and P2-only controls (see Supplementary Information). We approximated γ = 0 and computed λ and β for each promoter (Supplementary Table 53). For distributed multicellular co-cultures, we induced one memregulon at a time and multiplied the mixed-culture weight by the number of component strains. A custom Python script then read the analysed weights, generated all figure panels and computed the associated statistical summaries. Comprehensive analyses of growth curves, fluorescence trajectories and weight variations are provided in Supplementary Notes 8, 9, 10 and 11.

### Memregulon characterisation using flow cytometry

We inoculated glycerol stocks of bacteria carrying memregulons and grew them overnight in LB with ampicillin (100 µg/mL). The next day, we refreshed cultures by adding 200 µL to 2 mL LB with carbenicillin (80 µg/mL) and the cognate inducer(s), and incubated them for 4 h at 37 °C with shaking (200 rpm). Inducer concentrations matched those used for the plate reader assay. After 4 h in log phase, we pelleted the cultures for 10 min at 4 °C in an Eppendorf Centrifuge 5810 R at 3,000 rpm (1,902 × g), resuspended them in 2 mL PBS and loaded 200 µL per well into a 96-well plate. We ran plates on a BD LSRFortessa High Throughput Sampler (HTS), acquired with BD FACSDiva v8.0.1, using a sample flow rate of 1 µL/s, sample volume of 5 µL, mixing volume of 50 µL, mixing speed of 200 µL/s, five mixes, wash volume of 200 µL and BLR enabled. We recorded up to 50,000 events per sample.

### Flow cytometry data analysis

We analysed flow-cytometry data with a custom Python script. We subtracted per-channel backgrounds using medians from Empty-cell or PBS negative controls and estimated spectral spillover between EGFP and mCherry from same-run single-colour controls. The green and red area channels were B488.530.30.A and YG561.610.20.A, respectively. The FSC/SSC cell gate was a non-parametric density polygon in log10(FSC-A, SSC-A): the script learned the 99% high-density region separately from same-run P1-only and P2-only controls and used the convex hull of the two polygons as one consensus gate for every well. The script used every event within the FSC/SSC consensus gate to calculate single-cell W, removed non-finite transformed events and then clipped W to [0,1]. For each run, the script recalculated λ, β and γ from same-run gated P1-only and P2-only controls. We calculated W = (1 − γρ)/[1 − λβ + (λ − γ)ρ], where ρ = G/R. RP1 and GP1 are background-subtracted, spillover-corrected control totals for P1-only wells, and RP2 and GP2 are the corresponding P2-only totals. Supplementary Fig. 95 and Supplementary Note 13 show the complete gating exemplar, retained-event counts and conversion.

### Memregulon characterisation using Sanger sequencing

We inoculated glycerol stocks of bacteria carrying memregulons and grew them overnight with ampicillin (100 µg/mL). We purified plasmid DNA with the GeneJET kit (Thermo Fisher Scientific). We sequenced single-strain and P1-only/P2-only controls with the common CamR reverse primer TAACTGACTGAAATGCCT-CAAAATG. We sequenced co-cultures with the promoter-specific forward primers that Supplementary Data 1 lists. The ABIF files identify an Applied Biosystems 3730 DNA Analyzer, 3730 Series Data Collection Software 4 and KB Basecaller 1.4.1.8. As previously described, we base-called mixed, P1-only and P2-only traces, globally aligned them, used matching peaks as anchors for piecewise time-dilation, normalised the traces at positions where the EGFP and mCherry references matched, and estimated P1 fraction as the mean P1/P2 peak contribution at aligned mismatch positions containing both peaks, following the qSanger framework implemented with a custom algorithm in Biopython.

Memregulon weight characterisation by qPCR for limited orthogonal comparison. We performed a limited qPCR assay on plasmid DNA extracted from bacteria carrying PLux memregulons using a StepOnePlus Real-Time PCR System with StepOne Software v2.3, Fast SYBR Green master mix and ROX passive reference. We amplified P1 with RG1083 (CCTGTCCCCTCAGTTCATGT) and RG1084 (GTCCTCGAAGTTCATCACGC) targeting mCherry, P2 with RG1087 (GACGACGGCAACTACAAGAC) and RG1088 (TCCTTGAAGTCGATGCCCTT) targeting EGFP, and the shared plasmid backbone with RG1080 (CTGACCGCGTTTCTGCATAA) and RG1082 (GGCATGGTGGTATCACGTTC). Supplementary Data 1 lists all remaining oligonucleotides. The instrument export reports Ct/Cq values for each well; in the legacy export these primer sets are labelled PR-83/84, PR-87/88 and PR-80/82, respectively. For each sample, technical replicate Ct/Cq values were first summarised by the instrument software. Target abundance was normalised to the shared backbone using ΔCt/Cq normalisation and relative abundance values were calculated assuming two-fold amplification per cycle unless primer-efficiency values are supplied with the source data. The qPCR plasmid fraction was then calculated as WqPCR = aP1/(aP1+aP2), where aP1 and aP2 denote the normalised qPCR abundances of P1 and P2. The time course contains consecutive PLux kanamycin-learning stages, with A–C sample labels treated as biological replicate identifiers where present and repeated wells within each target treated as technical replicates. We assigned WqPCR only to wells or targets with a Ct/Cq value, a target assignment and matched P1/P2 measurements. Because this dataset covers one limited trajectory rather than the full promoter library, qPCR was not used to estimate Cneg · A or to support the pooled learning-rule regression.

### Experimental strategy for memregulon tic-tac-toe

We used tic-tac-toe as an externally routed supervised-learning task to test whether active memregulons can be selectively rewritten in co-culture. Player X was a fixed trainer that opened in the centre position (position 5). Player O was the bacterial player represented by memregulon cultures assigned to the remaining board positions. Physically separate cultures represented candidate O moves associated with abstract board states. For each O decision, the experimenter supplied the board context by exposing available O-position cultures to the inducers corresponding to the positions already occupied by X. Red and green fluorescence were converted into calibrated weights W by ratiometric analysis. An external winner-take-all rule then computed a weight-derived move score for each available position and selected the available position with the highest score as O’s move. When a position contained more than one culture, we summed the corresponding weight-derived move scores. When the highest scores were within one standard deviation across biological replicates, the model chose randomly among the tied positions. After a losing lesson for O, the externally specified supervised lesson sequence identified the cultures corresponding to O’s active losing branch. We exposed only those cultures to the cognate inducer context plus kanamycin; all other positions were propagated without kanamycin. Under this external supervision, the cells executed the local negative physical update when the lesson activated the relevant promoter in the presence of kanamycin. We updated glycerol stocks after each adaptation cycle and continued the supervised lesson sequence.

### Simulation of externally routed bacterial tic-tac-toe

We simulated the externally routed tic-tac-toe task to design the supervised wet-lab lesson sequence and to evaluate non-loss fraction from measured weights. Each memregulon’s move contribution was a weight-derived score equal to the stored weight multiplied by the promoter’s effective transcription parameter fitted from experimental data. Player X was fixed and opened in the centre under the first-move convention. Player O was the bacterial player. For each simulated O turn, board-context inputs corresponding to X’s occupied positions activated the relevant promoter channels, and an external WTA rule selected the available position with the largest score. In the offline lesson-design simulation only, a reinforcementlearning-style branch-pruning routine identified losing O branches and scheduled negative updates for the active O positions on those branches. For preliminary lesson-sequence design, the in silico abstraction decreased the selected active weights by 0.05. This fixed decrement applied only to the in silico abstraction. Experimental trajectories used fluorescence- or DNA-derived measured weights directly in the same WTA readout. Supplementary Note 1 provides the offline tic-tac-toe lesson-design details. For Fig. 3d, each Monte Carlo evaluation independently drew every experimentally characterised weight from a distribution defined by its measured pooled mean and s.d., clipped the resulting weights to [0,1] and evaluated the complete fixed game tree. We ran 100 evaluations per day. For each day, Fig. 3d reports the mean and s.d. across those evaluations.

Human-in-the-loop XOR scalability analyses. We modelled each co-culture as one neuron whose state was decoded from pooled red and green fluorescence using the experimentally calibrated ratiometric transfer functions described above. We then converted the decoded neuron states into inducer concentrations for the downstream layer using the corresponding promoter response functions. Specifically, the decoded states of the hidden layer neurons dictate the concentrations of specific chemical inducers (e.g., Sal, aTc, Cho, Ara) applied to the output neurons. For the one-stage WTA readout analysis, two output neurons received a bias input, two single-input channels and one explicit physical combinatorial promoter channel. For the multilayer scalability analysis, four hidden neurons received bias, x1 and x2, and two output neurons received bias together with all hidden outputs. We constrained all synaptic weights to the biological interval [0,1], where each weight represents the fraction of the corresponding memregulon in the P1 state. One epoch was defined as one complete presentation of all four XOR truth-table inputs. After winner-take-all classification of each input, local negative updates were applied only after incorrect WTA outputs, to the incorrect output neuron and to the active upstream edges that contributed to that decision, with clipping to [0,1]. Because the rule is strictly negative, we initialised the multilayer analyses from a biologically suitable OR-like state and then tested convergence to XOR under repeated negative updates. For robustness analyses, we repeated the same initial state over 100 shuffled truth-table presentation orders; 45/100 runs converged, showing that the protocol is sample-order sensitive. Supplementary Note 12 provides the full architecture, equations, parameter assignments and training procedure.

For the multilayer XOR readout in Fig. 4e, the deterministic network produced two output scores for each input pattern, y0 and y1. Winner-take-all classification assigned the input to class 0 when y0 > y1 and to class 1 when y1 > y0. To make the deterministic classification margin visible, we plotted the correct-output advantage rather than the two raw output scores. For an input with XOR target 0, this quantity was y0 - y1; for an input with XOR target 1, it was y1 - y0. The plotted values were multiplied by 10^4^ for readability. Positive values therefore indicate that the deterministic winner-take-all readout selected the correct XOR class. For each of 5,000 Monte Carlo evaluations per truth-table input, we drew every promoter weight from a clipped normal distribution defined by its measured mean and standard deviation. The Monte Carlo correct-label fraction is the fraction of parameter draws for which the winner-take-all output matched the XOR target. The Monte Carlo correct-label fractions are reported in the Fig. 4 caption and in the Source Data/Supplementary tables, while the raw output distributions are shown in the Supplementary Information. The one-stage correct-label fractions were 0.91/0.88/0.83/0.69 and the multilayer fractions were 0.52/0.49/0.48/0.51 for inputs 00/01/10/11, respectively.

Statistics & Reproducibility. No statistical method was used to predetermine sample size. We chose sample sizes from established bacterial-culture replicate practice and the available independent cultures. No data were excluded from the analyses. Before defining analysable observations, the pipeline removed non-finite transformed flow events, assigned WqPCR only to qPCR wells with matched measurements and retained plate-reader growth fits with r^2^ ≥ 0.9. The experiments were not randomized. The Investigators were not blinded to allocation during experiments and outcome assessment. For serial-passage stability and learning-state trajectories derived from plate-reader measurements, we first averaged technical measurements within each independent biological culture and then summarised each state by the mean and s.d. across biological replicates. We used Welch’s two-sample t-test for adjacent-state comparisons and a two one-sided tests procedure with unequal variances for stability analyses. For Fig. 1i, we fitted a three-component mixture model to the representative PBAD-M1 single-cell weights and evaluated goodness-of-fit using Pearson correlation. For Fig. 2e, we compared adjacent learning-state single-cell distributions with two-sample Kolmogorov–Smirnov tests. For Fig. 3e, we treated each board-position co-culture series as one independent unit, tested seven paired series with a two-sided exact sign-flip permutation test and estimated the 95% CI with 200,000 paired bootstrap draws. The seven series contain the 23 targeted and 90 untargeted transitions. For Fig. 3f, we summarised position-4 PLux and PCin calibrated weights as the mean and s.d. across biological replicates. To validate the theoretical learning rule in Supplementary Note 3, we quantified consecutive changes in mean weight for each promoter, estimated the sampling error of each change from the corresponding single-cell variances and event counts, fitted weighted through-origin regressions of −Δ ⟨*W⟩* against Var(W), and combined promoter-level slope estimates by inverse-variance weighting.

## Data Availability

Source data are provided with this paper. The Source Data workbook contains the numerical data underlying the main figures and supplementary numerical analyses. Supplementary Data 1 provides the oligonucleotide sequences, and the accompanying Source Data archive provides annotated plasmid sequence files. Figshare contains the raw datasets, standalone analysis scripts and qSanger-derived numerical values underlying Fig. 2c^31^. The Source Data file also provides the qSanger-derived values.

## Code Availability

The custom Python codebase used for neural-network simulations, flow-cytometry and plate-reader analysis, and automated figure generation is available in Figshare^31^ and as Supplementary Software 1.

## Acknowledgements

We acknowledge M. Kushwaha, M. Fuegger and T. Nowak for discussions.

## Funding Statement

RG acknowledges the departmental allocation from the School of Life Sciences (Faculty of Natural Sciences, Keele University). MI acknowledges funding from BBSRC BB/P020615/1 (EVOENGINE), the Volkswagen Foundation (LIFE: 93 065) and the Office of Naval Research Global (ONRG N62909-23-1-2099). AJ was funded by AEI (PID2023-151174NB-I00 and PID2020-118436GB-I00), BBSRC BB/P020615/1 (EVO-ENGINE), EU grants EVOPROG 610730 and PhotoSynH2 HORIZON-EIC-2021-PATHFINDERCHALLENGES (grant agreement 101070948), EPSRC-BBSRC BB/M017982/1 (WISB centre), the Office of Naval Research Global (ONRG N62909-23-1-2008), and the departmental allocation from the School of Life Sciences (University of Warwick). The project leading to these results has received funding from the la Caixa Foundation under the project code HR22-00405 [EvoPunch].

## Author Contributions Statement

A.J. conceived the project. A.J. developed the software, performed the formal analysis and prepared the visualizations. A.J. and S.P. developed the methodology. S.P., C.V., M.W., R.G., M.I., and A.J. performed the investigation. A.J. supervised the work and wrote the original draft. S.P., C.V., M.W., R.G., M.I. and A.J. reviewed and edited the manuscript. R.G. developed and performed qPCR assays for plasmid-ratio measurement and contributed to orthogonal validation of memregulon weight changes.

## Competing Interests Statement

The authors declare no competing interests.

## Notes

### Competing Interest Statement

The authors have declared no competing interest.

### Summary of Updates

The text and figures have been revised for clarity, and the manuscript has been reformatted from its previous double-spaced layout.

