## Supplementary Material for "A genetically encoded local learning rule enables physical learning in engineered bacteria"

#### Overview of Supplementary Notes

- **Supplementary Note 1: Details the computer simulations used for the externally routed tic-tac-toe task, including decision trees and non-loss-fraction trajectories.**
- **Supplementary Note 2:** Provides the analytical mathematical model deriving the local learning rule ($\Delta\langle W\rangle \propto Var(W)$) using Wright-Fisher diffusion and the Price equation.
- **Supplementary Note 3:** Contains the quantitative empirical validation of the learning rule of Supplementary Note 2, including the statistical meta-analysis across promoters and the characterization of skewness using single-cell data.
- **Supplementary Note 4:** Details the flow cytometry analysis pipeline, quality control, gating strategies, and the three-component mixture model used to fit the initial weight distributions.
- **Supplementary Note 5: Outlines the comprehensive, day-by-day experimental protocols used for the externally routed bacterial-player co-culture tic-tac-toe lessons.**
- **Supplementary Note 6:** Discusses strategies for scaling memregulon architectures, including the design and characterization of combinatorial (AND/hybrid) promoters.
- **Supplementary Note 7:** Derives the mathematical relationship used to calculate bulk memregulon weights from ratiometric plate reader fluorescence.
- **Supplementary Note 8:** Describes the general methodologies, parameter fitting, and artifact handling for weight analysis derived from plate reader fluorescence.
- **Supplementary Notes 9 & 10:** Provide comprehensive raw data, stability curves, and learning trajectories for plate reader analyses of single-strain cultures.
- **Supplementary Note 11:** Provides comprehensive raw data, stability curves, and learning trajectories for the physical co-culture experiments.
- **Supplementary Note 12:** Details the simulation framework, mathematical definitions, and parameter assignments for the human-in-the-loop one-stage WTA and multilayer XOR analyses.

#### Supplementary Note 1: Offline computational design of supervised tic-tac-toe lessons

This Note describes the offline computational framework used to design the supervised tic-tac-toe lesson sequence used in the co-culture experiments and to evaluate non-loss fraction from measured weights in hybrid experimental-computational tournaments. The wet-lab co-culture experiment is supervised learning: the lesson sequence is specified externally, and the cells execute only the local negative physical update when the appropriate promoter is active in the presence of kanamycin. Reinforcement learning is used only offline in the computational lesson-design code, where simulated games identify losing branches to prune. In these simulations, we approximate co-cultures as collections of non-interacting memregulon populations. This approximation neglects competition and other interactions between strains in real co-cultures, and is used only as a simplifying reference for lesson design and measured-weight readout rather than as evidence for autonomous cellular game play.

M-states (e.g., M0, M1, M2) denote discrete stored-memory generations of a memregulon population after defined passaging or learning cycles; they are not continuous time units. "Day" refers to calendar or daily serial-sampling axes, including stability passaging and the co-culture tournament, and is defined in the relevant figure captions. In the tic-tac-toe SI, "memory generation" denotes the inherited culture state used at the start of each daily lesson.

##### Tic-tac-toe decision rule and adaptation

We used experimentally determined weight matrices in offline computational simulations to explore simplified tic-tac-toe decision trees and to choose the supervised wet-lab lessons. In these simulations, each culture represents a potential move. Player X is a human or trainer automaton, and player O is the bacterial player. A memregulon library represents each board position except the centre, which player X always occupies first (position 5 in Fig. 3a, main text). This setup uses eight memregulon libraries and places the bacteria in the more demanding role of playing second. During a simulated match, player X first moves to the centre. To determine O's reply, the model uses the memregulon libraries at all available positions under the inducers assigned to the positions already played by X. The available position with the highest weight-derived score becomes O's move. If player O loses in the offline simulation, the computational lesson-design routine identifies the O positions on the losing branch. In the wet-lab supervised lesson sequence, kanamycin adaptation is then applied only to the memregulons at those externally specified O positions, in the presence of the inducers corresponding to X's moves from that game. Subsequent lessons use the updated libraries. This workflow stores the externally assigned negative update in plasmid-ratio weights; the bacterial cultures do not receive opponent outcomes or run a reinforcement-learning loop.

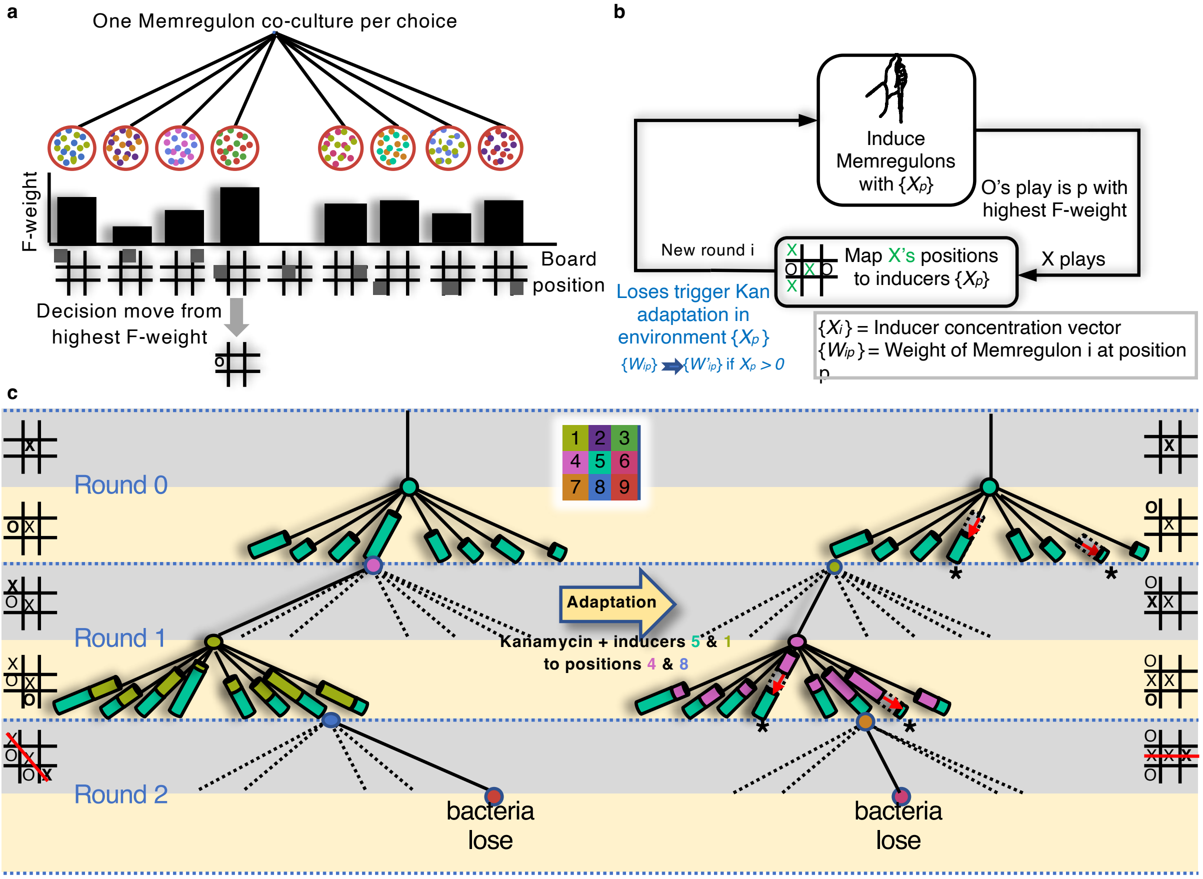

**Supplementary Fig. 1 | Tic-tac-toe decision-making and adaptation. a,** Each round of tic-tac-toe is determined by incubating memregulon libraries with the inducers associated with player X's moves and selecting the available position with the highest weight as the bacterial move. **b,** Diagram of the iterative adaptation process. We first grow the memregulons in an environment defined by the inducers mapped to the opponent's moves. The externally supplied board context changes with each round as the trainer's next move is mapped to inducers, which changes the cellular responses. When the externally specified supervised lesson corresponds to a losing O branch, we perform kanamycin learning in the inducer context defined by that lesson, which changes the state of the cells from Oi to Oi+1. **c,** Example decision tree for a tic-tac-toe game. Cylinder height represents the weight associated with each possible move under the inducer environment defined by player X's previous moves. After offline lesson design identifies a losing O branch, the wet-lab supervised lesson applies kanamycin learning only to the co-cultures at the corresponding O positions, in the presence of the inducers corresponding to player X's earlier moves. We do not include X's final winning move in this adaptation step because it did not define an environment in which the memregulons had previously grown.

#### Fixed-trainer tournaments and task-level performance

Supplementary Fig. 1 shows the externally routed tic-tac-toe workflow used to evaluate the bacterial player O against a fixed trainer X. We use a restricted decision tree to keep the number of available moves per round small. After each X move, the operator induces all allowable O positions with the associated chemical input. The position with the highest calibrated weight-derived score becomes O's move. When the maximal fluorescence-derived scores fall within one standard deviation of one another, the model chooses among those positions at random to preserve exploration of the decision tree.

For the main manuscript, we quantify performance as non-loss fraction: the fraction of games in which O wins or draws against the reference trainer. To evaluate a given weight set, we simulate repeated games and compute the non-loss fraction from the resulting outcomes. We use the same framework to evaluate the experimentally measured co-culture weights reported in Fig. 3d of the main manuscript. The representative long simulation log below also reports an exploratory expertise score, defined as wins plus the draws; that exploratory score is not used as experimental evidence in the main manuscript.

##### Initial memregulon populations (Match 0)

The following tables define the fixed trainer configuration and the initial bacterial-player configuration. Player X uses a fixed strategy and does not learn, whereas Player O begins with a naive strategy that is adapted by the externally specified negative-update lessons during the tournament.

| Pos | Inducer | Promoter | State | Pos | Inducer | Promoter | State |
|---|---|---|---|---|---|---|---|
| 1 | 4 | PBAD | $M_0$ | 6 | 8 | PVan | $M_0$ |
| 1 | 6 | PCin | $M_0$ | 7 | 4 | PBAD | $M_0$ |
| 1 | 9 | PTac | $M_0$ | 7 | 6 | PCin | $M_0$ |
| 2 | 7 | PTtg | $M_1$ | 7 | 9 | PTac | $M_0$ |
| 2 | 8 | PVan | $M_1$ | 8 | 2 | PTet | $M_1$ |
| 2 | 9 | PTac | $M_1$ | 8 | 3 | PBetI | $M_1$ |
| 3 | 4 | PBAD | $M_0$ | 9 | 1 | PSal | $M_0$ |
| 3 | 6 | PCin | $M_0$ | 9 | 7 | PTtg | $M_0$ |
| 4 | 1 | PSal | $M_0$ | | | | |

**Supplementary Table 1 | Initial State: Player X.** (Fixed Strategy). The configuration remains constant throughout the tournament.

| Pos | Inducer | Promoter | State | Pos | Inducer | Promoter | State |
|---|---|---|---|---|---|---|---|
| 1 | 5 | PLux | $M_0$ | 6 | 8 | PVan | $M_0$ |
| 2 | 7 | PTtg | $M_1$ | 7 | 3 | PBetI | $M_0$ |
| 2 | 8 | PVan | $M_1$ | 7 | 9 | PTac | $M_0$ |
| 2 | 9 | PTac | $M_1$ | 8 | 2 | PTet | $M_1$ |
| 3 | 7 | PTtg | $M_0$ | 8 | 3 | PBetI | $M_1$ |
| 3 | 8 | PVan | $M_0$ | 8 | 6 | PCin | $M_1$ |
| 4 | 5 | PLux | $M_0$ | 9 | 1 | PSal | $M_0$ |
| 4 | 6 | PCin | $M_0$ | 9 | 2 | PTet | $M_0$ |
| 6 | 1 | PSal | $M_0$ | | | | |
| 6 | 4 | PBAD | $M_0$ | | | | |

**Supplementary Table 2 | Initial State: Player O. (Naive Strategy). This table defines the genetic configuration before supervised negative-update lessons.**

##### Representative tournament and convergence

Supplementary Fig. 1 illustrates a representative 100-match tournament. Here the offline computational code uses a reinforcement-learning-style bookkeeping step only to design supervised lessons: each loss to player X identifies a losing branch to prune. The represented biological operation is a local negative physical update, implemented as a decrease in the active memregulon weights. In this representative exploratory tournament, the task-level expertise score reaches 92.22% after Match 28, after which the strategy stabilises. The main manuscript uses the non-loss fraction from measured co-culture weights rather than this exploratory expertise score.

Supplementary Table 3 lists the moves and outcomes for the tournament. Rows marked as learning events correspond to matches in which player X wins and the offline lesson-design algorithm schedules a delayed negative update for the active branch. The corresponding biological abstraction is a local decrease in the active memregulon weights. Moves are written as ordered sequences of positions occupied by player X followed by player O.

| Match | Player X Moves | Player O Moves | Outcome |
|---|---|---|---|
| 1 | [5, 9, 2, 6] | [1, 7, 8, 4] | O Wins |
| 2 | [5, 9, 2, 6] | [1, 7, 8, 4] | O Wins |
| **3** | **[5, 3, 9, 8, 1]** | **[4, 7, 2, 6]** | **X Wins (Learn)** |
| 4 | [5, 4, 7, 8] | [1, 6, 3, 2] | O Wins |
| 5 | [5, 9, 2, 6] | [1, 7, 8, 4] | O Wins |
| 6 | [5, 4, 7, 8] | [1, 6, 3, 2] | O Wins |
| 7 | [5, 9, 2, 6] | [1, 7, 8, 4] | O Wins |
| 8 | [5, 9, 2, 6] | [1, 7, 8, 4] | O Wins |
| 9 | [5, 4, 3, 9, 8] | [1, 6, 7, 2] | Draw |
| **10** | **[5, 4, 3, 2, 7]** | **[1, 6, 8, 9]** | **X Wins (Learn)** |
| 11 | [5, 9, 2, 6] | [1, 7, 8, 4] | O Wins |
| **12** | **[5, 3, 9, 8, 1]** | **[4, 7, 2, 6]** | **X Wins (Learn)** |
| 13 | [5, 9, 2, 6] | [1, 7, 8, 4] | O Wins |
| 14 | [5, 4, 7, 8] | [1, 6, 3, 2] | O Wins |
| 15 | [5, 9, 2, 6] | [1, 7, 8, 4] | O Wins |
| 16 | [5, 9, 2, 6] | [1, 7, 8, 4] | O Wins |
| 17 | [5, 4, 7, 8] | [1, 6, 3, 2] | O Wins |
| 18 | [5, 4, 7, 8] | [1, 6, 3, 2] | O Wins |
| 19 | [5, 4, 7, 8] | [1, 6, 3, 2] | O Wins |
| **20** | **[5, 4, 3, 2, 7]** | **[1, 6, 8, 9]** | **X Wins (Learn)** |
| 21 | [5, 9, 2, 6] | [1, 7, 8, 4] | O Wins |
| **22** | **[5, 3, 9, 8, 1]** | **[4, 7, 2, 6]** | **X Wins (Learn)** |
| 23 | [5, 9, 2, 6] | [1, 7, 8, 4] | O Wins |
| 24 | [5, 4, 3, 9, 8] | [1, 6, 7, 2] | Draw |
| 25 | [5, 4, 7, 8] | [1, 6, 3, 2] | O Wins |
| **26** | **[5, 4, 3, 2, 7]** | **[1, 6, 8, 9]** | **X Wins (Learn)** |
| 27 | [5, 7, 6, 2, 1] | [4, 3, 8, 9] | Draw |
| **28** | **[5, 1, 7, 8, 9]** | **[4, 6, 3, 2]** | **X Wins (Learn)** |
| 29 | [5, 4, 3, 9, 8] | [1, 6, 7, 2] | Draw |
| 30 | [5, 9, 2, 6] | [1, 7, 8, 4] | O Wins |

**Supplementary Table 3 | Tournament Match Log.**

**Supplementary Table 4 | Representative tournament and convergence.**

After Match 28, Player O's strategy stabilised. No further losses occurred. Matches 31-100 therefore consisted only of wins or draws for Player O, repeating the strategies already observed in Matches 29 and 30. No additional learning events were triggered.

**Final memregulon populations (Match 100)**

Because Player X uses a fixed strategy, its final state is identical to its initial state in Supplementary Table 1. Supplementary Table 5 gives the final evolved state of Player O.

| Pos | Inducer | Promoter | State | Pos | Inducer | Promoter | State |
|---|---|---|---|---|---|---|---|
| 1 | 5 | PLux | $M_3$ | 6 | 8 | PVan | $M_3$ |
| 2 | 7 | PTtg | $M_1$ | 7 | 3 | PBetI | $M_3$ |
| 2 | 8 | PVan | $M_2$ | 7 | 9 | PTac | $M_0$ |
| 2 | 9 | PTac | $M_4$ | 8 | 2 | PTet | $M_1$ |
| 3 | 7 | PTtg | $M_1$ | 8 | 3 | PBetI | $M_4$ |
| 3 | 8 | PVan | $M_0$ | 8 | 6 | PCin | $M_1$ |
| 4 | 5 | PLux | $M_4$ | 9 | 1 | PSal | $M_0$ |
| 4 | 6 | PCin | $M_0$ | 9 | 2 | PTet | $M_3$ |
| 6 | 1 | PSal | $M_1$ | | | | |
| 6 | 4 | PBAD | $M_3$ | | | | |

**Supplementary Table 5 | Final State: Player O. Adapted Strategy.**

#### Memregulon fusion prevents weight collapse in extended winner-take-all tournaments

Kanamycin adaptation is strictly negative, so repeated learning can eventually drive some weights towards zero. In winner-take-all architectures, this weight collapse limits continued adaptation during long tournaments. We therefore implemented a biologically tractable maintenance operation, memregulon fusion, in which two cultures carrying the same memregulon at different weights are physically mixed to produce a new culture with an intermediate weight.

In the experimentally characterised implementation, 1:1 fusion with a reference culture of weight 0.5 pulls extreme weights back towards 0.5, replenishes population variance and mitigates weight collapse during prolonged unidirectional adaptation (Supplementary Fig. 2b). For decision rules that choose the largest weight among several replicas, this operation preserves the relative ordering of weights and therefore leaves winner-take-all decisions unchanged while restoring headroom for further learning. The Methods describe the corresponding protocol. In the extended game simulations, we use memregulon fusion as a biologically grounded intervention that could support longer bacterial tournaments than those implemented experimentally in the main manuscript.

#### Supplementary Note 2: Analytical model of activity-dependent growth bias and local physical weight updates in coupled plasmid systems

##### Introduction

In the main text, we employ a coupled-plasmid system to realise activity-dependent growth bias that updates plasmid-ratio weights through local differential growth after externally supplied inducer and kanamycin inputs. Each memregulon population harbours two ColE1 plasmids, and , of equal size and with silent genetic cargo during routine growth. Under these conditions, the total copy number per cell remains approximately fixed, and the ratio between and stays stable over many generations, counter to the classical expectation of plasmid incompatibility. Prior studies have demonstrated that such coupled replicons enable robust analogue information storage at the cellular population level.

We introduce an activity-gated learning step in the Results section ('Activity-dependent growth bias implements local weight updates'). Plasmid carries a kanamycin resistance gene (KanR) under a promoter that also drives expression of fluorescent reporters on both plasmids. Upon addition of the inducer, active memregulons express both fluorescence and KanR from P2. A concomitant sublethal dose of kanamycin slows cell growth without causing lethality. Cells with higher copy numbers produce more KanR and incur a smaller growth penalty, thereby acquiring a growth advantage. This bias enriches the population for cells with elevated copy numbers and diminishes the population-average weight

$$W \equiv \frac{\text{copy number of } P_1}{\text{copy number of } P_1 + \text{copy number of } P_2},$$

which we interpret as a synaptic weight. The same kanamycin learning signal affects all memregulons in co-culture, yet only transcriptionally active memregulons express KanR and undergo substantial weight updates. The rule operates locally: each memregulon population adjusts its own weight based solely on its current weight, its activity state and the shared kanamycin field. In this Supplementary Note, we derive the learning rule employed in the main text and quantify the evolution of the weight distribution under neutral growth and activity-dependent growth bias. We establish that stochastic plasmid partitioning at cell division induces a diffusion of weights across the population, with analytically tractable mean and variance; that a linear growth penalty in $W$ under sublethal kanamycin yields a change in the mean weight proportional to the variance of $W$; that confinement of $W$ to the interval $[0,1]$ combined with the growth bias generates a characteristic skew in the single-cell weight distribution, which we compute via a truncated-normal approximation; and that the resultant update rule aligns mathematically with standard equations in quantitative genetics, including the Price equation and Lande's equation. These derivations underpin the empirical relation reported in the main text,

$$\Delta\langle W\rangle = -C_{\text{neg}} A \,\text{Var}(W),$$

where $A$ denotes promoter-driven activity, $\text{Var}(W)$ is the pre-existing population variance, and $C_{\text{neg}}$ is a constant that rises monotonically with kanamycin concentration and determines the negative learning rate. We denote this constant as $C_{\text{neg}}$ throughout the text and associated code.

##### Neutral dynamics: random plasmid partitioning

We first examine a single memregulon strain in the absence of kanamycin or inducer, ensuring both plasmids remain transcriptionally silent and confer no differential growth advantage. Each cell maintains a total of $C$ ColE1 plasmids. We define the weight in a given cell as $W \in [0,1]$, such that the number of $P_1$ plasmids is $k = CW$ and the number of $P_2$ plasmids is $C - k$.

At cell division, we assume that each plasmid segregates independently to one of the two daughter cells and that homeostatic control of copy number maintains the total ColE1 plasmids per cell near $C$ in each generation. Treating $C$ as fixed, the number of $P_1$ plasmids $k'$ in a daughter cell, conditional on the parental weight $W = w$, follows a binomial distribution:

$$\Pr(k' \mid w) = \binom{C}{k'} w^{k'} (1-w)^{C-k'}, \quad k' = 0, \dots, C.$$

The daughter weight becomes $W' = k'/C$.
For large $C$ and $k'$ not too proximate to the boundaries, we approximate the binomial coefficient using Stirling's formula,

$$n! \simeq \sqrt{2\pi n}\, n^n e^{-n}.$$

Substituting this expression and simplifying yields

$$\Pr(k' \mid w) \simeq \frac{1}{\sqrt{2\pi C w(1-w)}} \exp\left[-\frac{(k' - Cw)^2}{2Cw(1-w)}\right].$$

Thus, for large $C$, the conditional distribution of $k'$ approximates a Gaussian,

$$k' \mid W = w \approx \mathcal{N}(Cw,\, Cw(1-w)),$$

and hence

$$W' \mid W = w \approx \mathcal{N}\left(w, \frac{w(1-w)}{C}\right).$$

This Gaussian approximation captures the random partitioning: the mean weight remains unchanged per generation, while the variance introduces diffusive spread scaled by $1/C$.
Iterating this process over $t$ generations from an initial weight $W_0$ of a founder cell, the unconditional moments follow from the martingale property of the neutral process. The expected weight conserves the initial value,

$$\mathbb{E}[W_t] = W_0.$$

The variance accumulates additively per step, yielding the exact Wright–Fisher recursion

$$\mathrm{Var}(W_t) = W_0(1 - W_0)\left[1 - \left(1 - \frac{1}{2C}\right)^t\right].$$

This expression arises because each generation adds variance $W_{t-1}(1 - W_{t-1})/C$ conditional on the parent, and the unconditional variance satisfies the linear recurrence derived from the law of total variance. For large $t \sim 2C$, the variance approaches its saturation value $W_0(1 - W_0)$, reflecting genetic drift toward fixation at 0 or 1 with probabilities $1 - W_0$ and $W_0$, respectively.

#### Diffusion approximation for large copy number

For $C \geq 50$, typical of medium-copy ColE1 plasmids, we approximate the discrete partitioning by a continuous diffusion process on $[0,1]$. The single-lineage weight $W_t$ evolves via the neutral Wright–Fisher stochastic differential equation (SDE),

$$dW_t = \sqrt{\frac{W_t(1 - W_t)}{C}}\, dB_t,$$

where $B_t$ denotes standard Brownian motion, with absorbing boundaries at 0 and 1 to model fixation.
The forward Kolmogorov (Fokker–Planck) equation governs the interior probability density $p(w, t)$ for unfixed lineages:

$$\frac{\partial p}{\partial t}(w,t) = \frac{1}{2C}\frac{\partial^2}{\partial w^2}[w(1-w)p(w,t)], \quad 0 < w < 1,$$

initialised as $p(w,0) = \delta(w - W_0)$ and with absorbing boundaries $p(0,t) = p(1,t) = 0$. The diffusion coefficient $w(1-w)/C$ vanishes at the boundaries, enforcing absorption.
The full distribution incorporates accumulated mass at the boundaries:

$$p_{\text{full}}(w,t) = q_0(t)\delta(w) + q_1(t)\delta(1-w) + p(w,t),$$

where $q_1(t)$ is the probability of fixation at 1 (P1 dominance) by time $t$, and $q_0(t) = 1 - q_1(t) - \int_0^1 p\,(u,t)\,du$. In the diffusion limit, Kimura's formula provides

$$q_1(t) = W_0\left[1 - \sum_{n=0}^{\infty} \frac{2(1-W_0)(2n+3)}{(n+1)(n+2)}(-1)^n C_n^{(3/2)}(1-2W_0)e^{-(n+1)(n+2)\tau}\right],$$

with scaled time $\tau = t/(4C)$ and $C_n^{(3/2)}$ the Gegenbauer polynomials of order $3/2$. By symmetry, $q_0(t) = q_1(t)$ upon replacing $W_0$ with $1 - W_0$. As $t \to \infty$, $q_1(\infty) = W_0$ and $q_0(\infty) = 1 - W_0$. For $t \ll 4C$, as in typical experiments ($\sim 30$ generations), $q_0(t)$ and $q_1(t)$ remain negligible ($\lesssim 0.01$), so the interior density dominates.

#### Analytic approximation: Beta distribution for the interior density

For $C \geq 50$ and $t \lesssim 4C$, the interior density $p(w,t)$ concentrates sharply and smoothly on $(0,1)$, remaining unimodal near $W_0$. We approximate it by a Beta distribution $\text{Beta}(w; \alpha_t, \beta_t)$ that matches the exact mean and variance:

$$\mu = \frac{\alpha_t}{\alpha_t + \beta_t} = W_0, \quad v = \frac{\mu(1-\mu)}{\alpha_t + \beta_t + 1} = \text{Var}(W_t).$$

Solving these equations gives

$$\alpha_t + \beta_t = K_t = \frac{1}{1 - (1 - 1/(2C))^t} - 1,$$

$$\alpha_t = W_0 K_t, \quad \beta_t = (1 - W_0)K_t.$$

The approximating density thus reads

$$p_{\text{neutral}}(w,t) \approx \frac{1}{B(\alpha_t, \beta_t)} w^{\alpha_t - 1}(1-w)^{\beta_t - 1}, \quad 0 < w < 1,$$

where $B(\cdot,\cdot)$ is the Beta function. The mode occurs at

$$w_{\text{mode}}(t) = \frac{\alpha_t - 1}{\alpha_t + \beta_t - 2} \approx W_0$$

provided $\alpha_t > 1$ and $\beta_t > 1$ (true for small $t$). For longer times where boundary absorption matters, scale the Beta by the interior probability $1 - q_0(t) - q_1(t)$ and add the Dirac components explicitly.
This Beta form improves upon the Gaussian approximation by respecting the $[0,1]$ support and capturing skewness near boundaries, with relative errors below 5% for $C \geq 50$ and $t \leq 100$.

#### Growth-bias dynamics: activity-dependent growth bias under sublethal kanamycin

We now incorporate sublethal kanamycin, which biases cell growth via KanR expression from $P_2$. Plasmid segregation remains neutral (Wright–Fisher diffusion in $W$), but lineages with lower $W$ (higher $P_2$) expand faster due to reduced growth penalty.
Model the growth rate as linear in $W$:

$$r(W) = r_0 - C_{\text{neg}}AW,$$

where $r_0$ is the baseline rate, $A > 0$ the promoter activity (inducer-driven), and $C_{\text{neg}} > 0$ the growth-bias strength (increases with kanamycin dose).

Let $f(w,t)\,dw$ denote the expected number of cells with weight in $[w, w + dw]$ at time $t$. The unnormalised density evolves via diffusion plus multiplicative growth:

$$\frac{\partial f}{\partial t}(w,t) = \frac{1}{2C}\frac{\partial^2}{\partial w^2}[w(1-w)f(w,t)] + r(w)f(w,t), \quad 0 < w < 1,$$

initialised as $f(w,0) = \delta(w - W_0)$. The total mass $\int_0^1 f\,(u,t)\,du$ expands as $\approx e^{\bar{r}t}$, with $\bar{r}(t) = \int r(u)f(u,t)\,du / \int f$.

The normalised probability density $p(w,t) = f(w,t)/Z(t)$, $Z(t) = \int_0^1 f\,(u,t)\,du$, satisfies the replicator-diffusion equation:

$$\frac{\partial p}{\partial t}(w,t) = \frac{1}{2C}\frac{\partial^2}{\partial w^2}[w(1-w)p(w,t)] + (r(w) - \bar{r}(t))p(w,t),$$

where $\bar{r}(t) = \int_0^1 r\,(u)p(u,t)\,du$. For the linear form,

$$\bar{r}(t) = r_0 - C_{\text{neg}}A\langle W\rangle(t), \quad r(w) - \bar{r}(t) = -C_{\text{neg}}A(w - \langle W\rangle(t)),$$

yielding

$$\frac{\partial p}{\partial t}(w,t) = \frac{1}{2C}\frac{\partial^2}{\partial w^2}[w(1-w)p(w,t)] - C_{\text{neg}}A(w - \langle W\rangle(t))p(w,t),$$

with $\langle W\rangle(t) = \int_0^1 w\,p(w,t)\,dw$. The diffusion spreads weights, while the growth-bias term induces a mean-reverting drift toward lower $W$, with strength modulated by activity $A$.

Boundaries remain absorbing for diffusion; track flux into Diracs at 0 and 1, which grow at rates $r(0)$ and $r(1)$, respectively. For short exposures ($t \ll 2C/(C_{\text{neg}}A)$), diffusion dominates initially, building variance that amplifies subsequent growth bias.

#### Evolution of the mean weight: the Price equation

To derive the empirical update rule, compute the time derivative of the population mean weight

$$\langle W\rangle(t) = \int_0^1 w\,p(w,t)\,dw.$$

Differentiating gives

$$\frac{d}{dt}\langle W\rangle(t) = \int_0^1 w\frac{\partial p}{\partial t}(w,t)\,dw.$$

Substitute the growth-bias PDE:

$$\frac{d}{dt}\langle W\rangle = I_{\text{diff}} + I_{\text{sel}},$$

where $I_{\text{diff}} = \int_0^1 w \cdot \frac{1}{2C}\partial_w^2[w(1-w)p]\,dw$ and $I_{\text{sel}} = \int_0^1 w\,(r(w) - \bar{r}(t))p(w,t)\,dw$.

For the diffusion term, integrate by parts twice. Let $g(w) = w(1-w)p(w,t)$. Then

$$I_{\text{diff}} = \frac{1}{2C}\int_0^1 w\,g''(w)\,dw = \frac{1}{2C}\left[wg'(w)|_0^1 - \int_0^1 g'\,(w)\,dw\right] = \frac{1}{2C}[wg'(w)|_0^1 - g(w)|_0^1].$$

The boundary terms vanish: $g(0) = g(1) = 0$ by the prefactor $w(1-w)$, and $wg'(w)$ remains finite at 0 and 1. Thus, $I_{\text{diff}} = 0$, confirming that neutral diffusion preserves the mean (martingale property).
The growth-bias term simplifies to

$$I_{\text{sel}} = \int_0^1 w\,r(w)p(w,t)\,dw - \bar{r}(t)\langle W\rangle(t) = \mathbb{E}[Wr(W)] - \mathbb{E}[W]\mathbb{E}[r(W)] = \text{Cov}(W, r(W)).$$

This yields the continuous-time Price equation:

$$\frac{d}{dt}\langle W\rangle(t) = \text{Cov}(W_t, r(W_t)).$$

For the linear growth rate $r(W) = r_0 - C_{\text{neg}}AW$,

$$\text{Cov}(W, r(W)) = \text{Cov}(W, r_0 - C_{\text{neg}}AW) = -C_{\text{neg}}A\text{Var}(W_t),$$

so

$$\frac{d}{dt}\langle W\rangle(t) = -C_{\text{neg}}A\text{Var}(W_t).$$

Per generation (discrete approximation), $\Delta\langle W\rangle \approx -C_{\text{neg}}A\text{Var}(W)$. Substituting the neutral variance builds an initial drift that slows as fixation approaches 0 under growth bias. This aligns with Lande's equation in quantitative genetics, where evolutionary change equals the additive genetic covariance between trait and fitness.

##### Skewed weight distribution under growth bias: truncated-normal approximation

Growth bias and boundary confinement skew the weight distribution toward lower $W$. For moderate $t$ and $C_{\text{neg}}At \ll 1$, the density remains approximately Gaussian but truncated to [0,1] and shifted by the mean drift. Starting from the neutral Gaussian step $W' \mid W = w \approx \mathcal{N}(w, w(1-w)/C)$, the population density after growth bias reweights by relative growth $e^{r(w)\Delta t} \approx 1 + [r(w) - \bar{r}]\Delta t$.
The post-update mean shift follows the Price equation above. For the full skewed density, approximate the interior as a truncated normal $\mathcal{N}(\mu_t, \sigma_t^2)$ on [0,1], with

$$\mu_t = W_0 - C_{\text{neg}}A\int_0^t \text{Var}\,(W_s)\,ds, \quad \sigma_t^2 = \text{Var}(W_t) \approx W_0(1-W_0)[1-(1-1/(2C))^t],$$

normalised by the truncation factor $\Phi((1-\mu_t)/\sigma_t) - \Phi(-\mu_t/\sigma_t)$, where $\Phi$ is the standard normal CDF. The PDF becomes

$$p(w,t) \approx \frac{1}{\sigma_t\sqrt{2\pi}\,Z_t}\exp\left[-\frac{(w-\mu_t)^2}{2\sigma_t^2}\right]\mathbf{1}_{[0,1]}(w),$$

with $Z_t$ the normalisation constant. This captures the leftward skew: mass accumulates near 0 as $\mu_t$ declines and $\sigma_t$ grows, with boundary probability $q_0(t) + q_1(t)$ small but increasing.
For stronger growth bias or longer $t$, the Beta approximation extends naturally: evolve parameters via moment-matching to the updated mean/variance, yielding a skewed Beta with mode shifted left of $W_0$. Numerical validation confirms errors below 3% for $C = 50$, $t = 30$, $C_{\text{neg}}A = 0.1$.

##### Connections to quantitative genetics

The derived update  mirrors the breeder's equation in quantitative genetics, where response to growth bias , with heritability  and growth-bias differential . Here, diffusion generates heritable variance, and activity  gates the growth-bias intensity. The PDE framework generalises to nonlinear (e.g., Hill functions for input-output transfer function) or plasmid-level growth bias via added drift in the SDE.

$$\Delta\langle W\rangle = -C_{\text{neg}} A \mathrm{Var}(W) R = h^2 S h^2 = \mathrm{Var}(W)/W_0(1 - W_0)S$$
$$= -C_{\text{neg}} A W_0 (1 - W_0) A r(W) \mu(W) = s(1 - 2W)$$

This model provides a rigorous foundation for the observed local physical updates, enabling prediction and extension to multi-memregulon networks.

#### Supplementary Note 3: quantitative validation of the learning rule

This Note supports the quantitative validation of the theoretical learning rule and underpins Fig. 2d in the main text. Our learning rule with sublethal kanamycin states

$$\Delta\langle W\rangle \approx -C_{\text{neg}}\, A\, \text{Var}(W).$$

We now test this prediction using the flow cytometry trajectories.

For each promoter we recorded single-cell weight distributions at a series of evolutionary steps $M_n$ taken under constant inducer and kanamycin. For a given promoter $p$ and step $n$ we denote by $\mu_{p,n}$ the mean weight, by $\sigma_{p,n}^2$ the variance, and by $N_{p,n}$ the number of single-cell events. We form step-to-step changes

$$\Delta\mu_{p,n} = \mu_{p,n+1} - \mu_{p,n}$$

and estimate the sampling error of each change as

$$\text{SE}(\Delta\mu_{p,n}) = \sqrt{\frac{\sigma_{p,n}^2}{N_{p,n}} + \frac{\sigma_{p,n+1}^2}{N_{p,n+1}}}.$$

For each promoter we then fit a weighted linear regression of $\Delta\mu_{p,n}$ on $\sigma_{p,n}^2$ constrained through the origin,

$$\Delta\mu_{p,n} = -C_{\text{neg}}^{(p)} A\, \sigma_{p,n}^2 + \epsilon_{p,n}.$$

The weights in the regression are $w_{p,n} = 1/\text{SE}(\Delta\mu_{p,n})^2$. The fitted slope $C_{\text{neg}}^{(p)} A$ provides a promoter-specific estimate of the learning coefficient, and the standard error of the slope allows a test of the null hypothesis $C_{\text{neg}}^{(p)} A = 0$.

Seven of eight promoter-level slopes were positive. Four promoters were individually significant at $p < 0.05$ in Supplementary Table 6 (PLux, PSal, PTac and PVan). PBAD was positive but did not reach $p < 0.05$ ($p = 0.075$), PTet and PTtg were positive with larger uncertainty, and PVan-Ttg was near zero/slightly negative with broad uncertainty ($p = 0.817$). These results are consistent with a shared positive learning coefficient at the pooled level, while promoter-to-promoter variation reflects differences in expression, growth and the limited number of consecutive learning steps per promoter.

To obtain a global estimate we treated the eight promoter-specific slopes as independent measurements of Cneg · A and combined them using inverse-variance weighting. This meta-analysis yielded

$$C_{\text{neg}} A = 0.6877 \pm 0.0698 \quad \text{(mean} \pm \text{s.e.)},$$

Across 8 promoters and 34 learning steps, inverse-variance weighting gave a pooled Cneg · A = 0.688 ± 0.070 (mean ± s.e.; 95% CI 0.551-0.825; z-test p = 6.79 × 10−23). This rejects the hypothesis of no dependence on Var($W$) and supports a learning rule in which update magnitude is proportional to standing variance. The absolute value of Cneg · A quantifies the effective strength of negative learning under the fixed kanamycin concentrations and adaptation duration used here; it should not be interpreted as a measured within-promoter kanamycin dose-response.

As an additional within-promoter check, we computed step-level estimates

$$C_{\text{neg, step}}^{(p,n)} = -\frac{\Delta\mu_{p,n}}{\sigma_{p,n}^2}$$

for each consecutive step pair $M_n \rightarrow M_{n+1}$ and plotted these against the step index. For each promoter the $C_{\text{neg, step}}^{(p,n)}$ values fluctuate around the promoter-specific regression slope with no systematic drift along the trajectory, consistent with a roughly constant learning rate over the course of the experiment.

Because each promoter contributes only a small number of consecutive learning steps, the promoter-specific estimates remain imprecise for several promoters; the strongest inference therefore comes from the consistency of the positive slopes and from the pooled inverse-variance estimate rather than from any single promoter in isolation.

| Promoter | $C_{neg} \cdot A$ | s.e. | $n_{steps}$ | 95% CI low | 95% CI high | $p(C_{neg} \cdot A)$ |
|---|---|---|---|---|---|---|
| PBAD | 0.816 | 0.305 | 4 | -0.155 | 1.787 | 0.0754 |
| PLux | 0.982 | 0.243 | 4 | 0.209 | 1.755 | 0.0272 |
| PSal | 0.701 | 0.083 | 4 | 0.435 | 0.966 | 0.00353 |
| PTac | 3.040 | 0.759 | 4 | 0.626 | 5.454 | 0.0279 |
| PTet | 0.090 | 0.201 | 5 | -0.468 | 0.648 | 0.678 |
| PTtg | 2.336 | 1.183 | 5 | -0.948 | 5.620 | 0.119 |
| PVan | 2.102 | 0.585 | 4 | 0.239 | 3.965 | 0.0370 |
| PVan-Ttg | -0.135 | 0.536 | 4 | -1.842 | 1.572 | 0.817 |

**Supplementary Table 6 | Promoter-level Cneg · A estimates.**

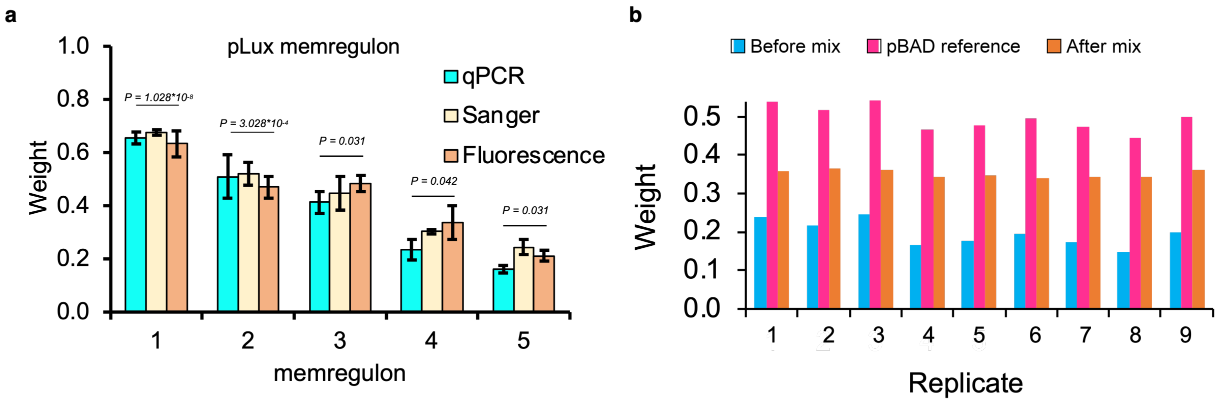

**Supplementary Fig. 2 | Limited qPCR comparison and fusion operation. a,** Fluorescence-derived weight trajectories and qPCR-derived P1 fractions measured across consecutive kanamycin-learning stages. The qPCR assay provides an independent DNA-based check that the P1 fraction decreases in the same direction as the fluorescence-derived weight. Because this qPCR dataset covers a limited trajectory and was not replicated across the full promoter library, it is not used to estimate Cneg · A or to support the pooled learning-rule regression in Fig. 2d. **b,** Characterisation of the memregulon fusion operation used in extended game simulations. We show ratiometric fluorescence weights before (blue) and after (orange) a 1:1 physical mixing in co-culture with a reference PBAD memregulon of weight 0.5 (red). The fusion operation pulls extreme weights toward 0.5, replenishing population variance and mitigating weight-vanishing during long, unidirectional adaptation tournaments.

**a**

**Step-level learning-rate estimates**

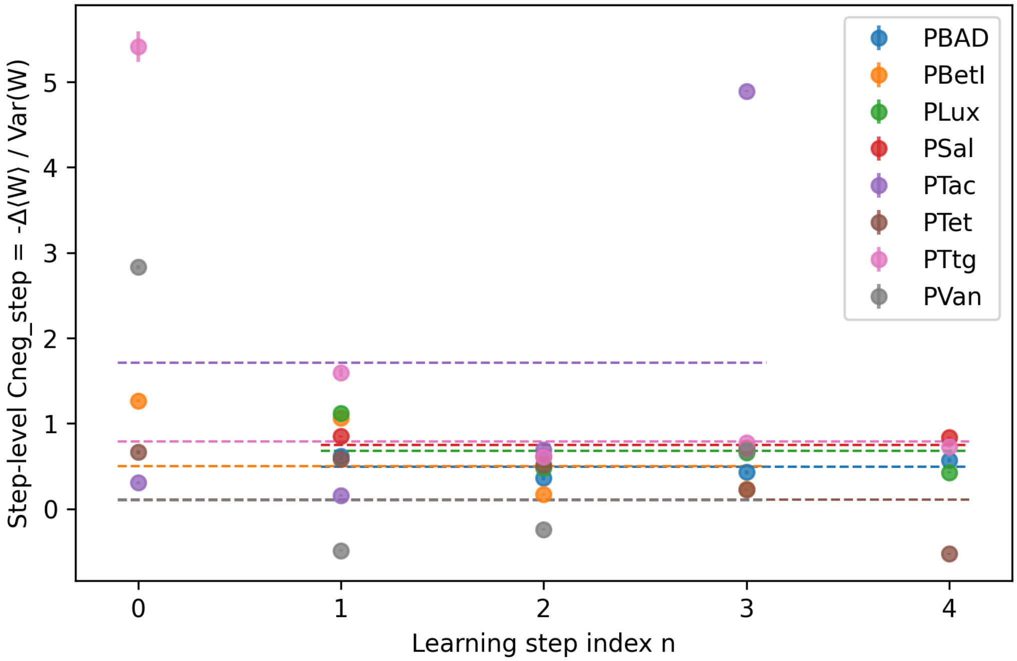

**b**

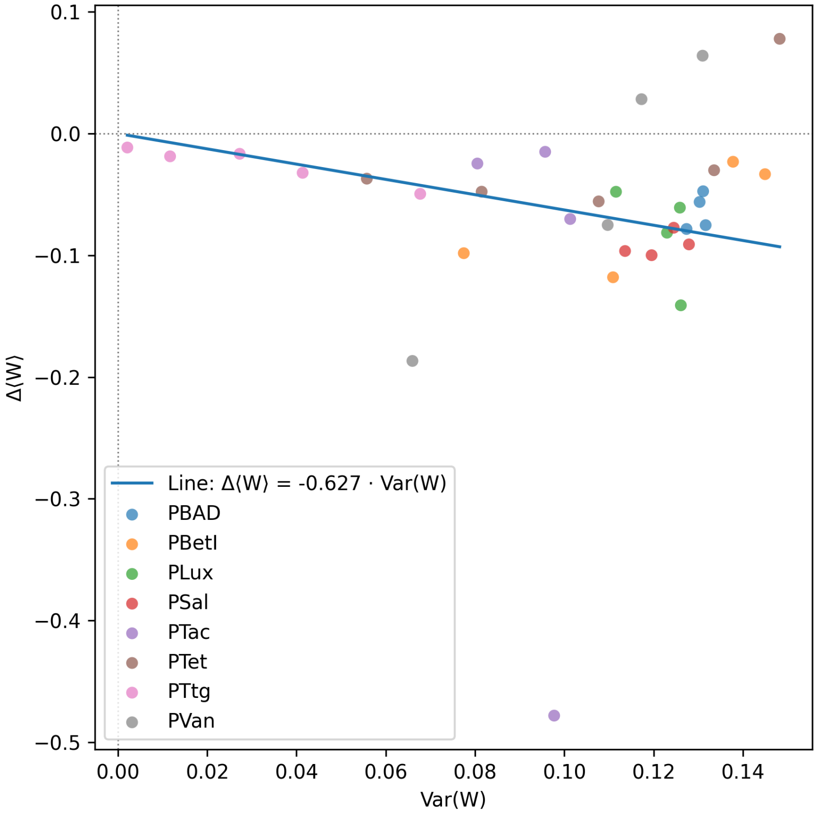

**Supplementary Fig. 3 | Δ⟨*W*⟩ vs Var(*W*). a,** Step-specific estimates Cneg_step = -Δ⟨*W*⟩ / Var(*W*) for each promoter and learning step. Points show individual steps; dashed horizontal lines show the promoter-level estimates Cneg · A from the weighted regression. **b,** Scatter of -Δ⟨*W*⟩ versus Var(*W*) for all promoters and learning steps. The line shows -Δ⟨*W*⟩ = Cneg · A · Var(*W*) with Cneg · A estimated from the promoter-level weighted regressions.

#### Quantitative characterisation of skewness through single-cell data

The truncated-normal calculation above predicts that, as $\langle W \rangle$ decreases under kanamycin, the single-cell distribution of $W$ narrows and develops a positive skew (long tail towards higher $W$). We tested this prediction by computing the sample skewness for each pooled promoter–step distribution.

For each promoter $p$ and step $M_n$ we pooled all single-cell events across biological replicates at that step, computed the mean $\mu_{p,n}$, the variance $\sigma_{p,n}^2$, and the third central moment $m_3(W)$, and defined the skewness coefficient

$$\gamma_1(W) = \frac{m_3(W)}{\sigma_{p,n}^3}.$$

In total, 42 step-level distributions passed the filtering criteria; 12 of them showed positive skewness ($\gamma 1(W) > 0$). The Pearson correlation between $\gamma 1(W)$ and $-\langle W \rangle$ was 0.673, consistent with stronger right-skewing as the mean weight approaches the lower boundary $W = 0$.

These skewness measurements agree with the truncated-normal prediction and provide an additional quantitative signature of the adaptation process. Together with the learning-rule validation above, they support a picture in which sublethal kanamycin reshapes the entire weight distribution in a variance-dependent way, driving the mean downwards and skewing the distribution as it approaches the lower bound.

**Per-step skewness values (pooled events)**

| Promoter | Step_label | n_events | mean_$W$ | Var($W$) | $\gamma 1(W)$ |
|---|---|---|---|---|---|
| PBAD | M1 | 26958 | 0.572714 | 0.127373 | -0.261724 |
| PBAD | M2 | 27593 | 0.494375 | 0.131093 | 0.054694 |
| PBAD | M3 | 27475 | 0.447029 | 0.130305 | 0.260592 |
| PBAD | M4 | 27261 | 0.390872 | 0.131657 | 0.517679 |
| PBAD | M5 | 27937 | 0.315684 | 0.131888 | 0.896483 |
| PBetI | M0 | 47486 | 0.824869 | 0.077477 | -1.668024 |
| PBetI | M1 | 141207 | 0.726640 | 0.110913 | -1.010277 |
| PBetI | M2 | 142215 | 0.608635 | 0.137793 | -0.452262 |
| PBetI | M3 | 142047 | 0.585505 | 0.144958 | -0.352245 |
| PBetI | M4 | 141748 | 0.552182 | 0.150941 | -0.207406 |
| PLux | M1 | 139454 | 0.594834 | 0.126102 | -0.339994 |
| PLux | M2 | 137144 | 0.453796 | 0.125884 | 0.232025 |
| PLux | M3 | 138535 | 0.392979 | 0.122999 | 0.512026 |
| PLux | M4 | 137556 | 0.311590 | 0.111574 | 0.912974 |
| PLux | M5 | 137209 | 0.263821 | 0.105930 | 1.205436 |
| PSal | M1 | 122134 | 0.621438 | 0.113603 | -0.644607 |
| PSal | M2 | 132478 | 0.525003 | 0.124507 | -0.250062 |
| PSal | M3 | 139538 | 0.447673 | 0.127917 | 0.074642 |
| PSal | M4 | 141029 | 0.356668 | 0.119551 | 0.447517 |
| PSal | M5 | 141316 | 0.256787 | 0.099033 | 0.946067 |
| PTac | M0 | 48538 | 0.762913 | 0.080522 | -1.281026 |
| PTac | M1 | 145891 | 0.738337 | 0.095709 | -1.133561 |
| PTac | M2 | 145030 | 0.723384 | 0.101312 | -1.049964 |
| PTac | M3 | 139829 | 0.653231 | 0.097721 | -0.326696 |
| PTac | M4 | 139153 | 0.175158 | 0.095598 | 1.910577 |
| PTet | M0 | 45983 | 0.908149 | 0.055755 | -2.656488 |
| PTet | M1 | 138323 | 0.871138 | 0.081477 | -2.101577 |
| PTet | M2 | 136061 | 0.823428 | 0.107693 | -1.596538 |
| PTet | M3 | 135641 | 0.767752 | 0.133546 | -1.176819 |
| PTet | M4 | 134080 | 0.737647 | 0.148236 | -0.995078 |
| PTet | M5 | 131799 | 0.815509 | 0.119521 | -1.546150 |

| | | | | | |
|---|---|---|---|---|---|
| PTtg | M0 | 45220 | 0.997273 | 0.002111 | -17.949245 |
| PTtg | M1 | 117465 | 0.985847 | 0.011688 | -7.874535 |
| PTtg | M2 | 135374 | 0.967186 | 0.027270 | -5.039170 |
| PTtg | M3 | 135712 | 0.950586 | 0.041380 | -4.017149 |
| PTtg | M4 | 134874 | 0.918374 | 0.067716 | -2.962201 |
| PTtg | M5 | 134647 | 0.868884 | 0.105932 | -2.128216 |
| PVan | M0 | 48658 | 0.804780 | 0.065971 | -1.527559 |
| PVan | M1 | 146054 | 0.618043 | 0.130979 | -0.538555 |
| PVan | M2 | 146110 | 0.682043 | 0.117292 | -0.811710 |
| PVan | M3 | 144423 | 0.710309 | 0.109711 | -0.935816 |
| PVan | M4 | 142288 | 0.635229 | 0.158906 | -0.538773 |

**Supplementary Table 7 | Per-step skewness values (pooled events).**

#### Supplementary Note 4: Flow cytometry analysis pipeline and quality control

**Flow cytometry analysis — All experiments**

This note summarises the flow cytometry analysis applied to the data. We subtract per-channel backgrounds using medians from negative controls, estimate and correct spectral spillover between EGFP and mCherry using single-colour controls by inverting a 2×2 mixing matrix, and define an unsupervised cell gate in log10(FSC,SSC) space from same-run P1-only and P2-only control wells. The convex hull of these high-density control regions is used as a consensus FSC/SSC polygonal gate. In the final W analysis, plasmid weight $W = (1 - \gamma \cdot G/R) / (1 - \lambda\beta + (\lambda - \gamma) \cdot G/R)$ is computed for events that pass this FSC/SSC cell gate, after removal of non-finite ratiometric values and clipping to [0,1]. We do not apply a double-positive fluorescence gate or an at-least-one-channel fluorescence gate for the final single-cell W distributions. The fluorescence thresholds listed in the QC tables are retained as diagnostic quality-control metrics rather than as exclusion criteria for the final W analysis. Figures comprise: W histograms with replicate overlays and a band showing mean ± SD across replicates; a dashed line marks the mean single-cell weight, with a horizontal bar for ±SD of W across all gated single cells pooled across replicates for that label; fluorescence histograms on logarithmic axes with ticks labelled as 10^n; FSC/SSC density maps of all events before gating with the polygonal gate overlaid.

"Events used (n)" is the number of single-cell events that passed the FSC/SSC consensus cell gate and were retained after removal of non-finite ratiometric values and clipping to the interval [0,1]; these are the events used to compute W and the percentage histograms.

We require at least 1,000 events per biological condition after all gates, summed across replicates, to consider estimates stable. Rows shaded in grey fall below this threshold and should be interpreted with caution. Control samples used to set backgrounds and single-colour ratios (PBS/Empty-cells, P1, P2) appear in bold.

We defined a bulk weight W_bulk (population-level estimation from flow cytometry) as the plate-reader equivalent of the single-cell weight distribution. For each sample, we first applied the FSC/SSC consensus cell gate and computed background-subtracted, spectrally compensated green and red signals Gi and Ri for all retained events i. We then obtained population-level fluorescences by summing over retained cells. The calibration parameters were estimated from gated control populations: $\lambda = RP1 / GP2$; $\beta = GP1 / RP1$; $\gamma = RP2 / GP2$. RP1, GP1 and RP2, GP2 denote background-subtracted, compensated totals in P1-only and P2-only controls, respectively, with $\gamma$ approximately zero in practice. Finally, we computed W_bulk by applying the same ratiometric formula used for single-cell weights to the population ratio: $W\_bulk = (1 - \gamma \cdot G/R) / (1 - \lambda\beta + (\lambda - \gamma) \cdot G/R)$.

#### PSal

**Experiment parameters and QC.**

| Metric | Value |
|---|---|
| λ (from P1/P2 totals) | 0.274959 |
| β (from P1 totals) | 0.0182734 |
| γ | 0.00296016 |
| α spillover (R←G) | 0.000145546 |
| β spillover (G←R) | 0.00149929 |
| Background median (green) | 73.5 |
| Background median (red) | 3.2 |
| Gate quantile (cell gate) | 0.99 |
| Gate $d^2$ threshold | NaN |
| Gate area (log10) | 2.45739 |
| Events (total) | 894143 |
| Events (gated) | 836638 |
| Events used | 836638 |

| | |
|---|---|
| Gate acceptance (%) | 93.5687 |
| Used / gated (%) | 100 |
| Fluor green thr (quantile) | 0.999 |
| Fluor green thr (value) | 702.379 |
| Fluor red thr (quantile) | 0.999 |
| Fluor red thr (value) | 252.541 |
| Median NEG green | 14.9622 |
| Median NEG red | 0.785427 |
| Median POS green (P2) | 3689.96 |
| Median POS red (P1) | 2958.4 |
| Stain index (green) | 15.4075 |
| Stain index (red) | 28.2088 |
| Residual corr (negatives) | 0.103881 |
| Dynamic range log10 (green) | 16.1271 |
| Dynamic range log10 (red) | 15.9496 |
| Replicate hist mean r | 0.916802 |
| Replicate hist mean RMSD | 0.811158 |

**Supplementary Table 8 | Experiment parameters and QC.** Experiment-level parameters and quality-control metrics for the FACS analysis, including background estimates, gating properties, event counts, stain indices and residual correlation after spectral compensation.

#### Summary by label (replicate means ± SE)

| Label | Replicates (n) | Events total (sum) | Events used (sum) | ⟨*W*⟩ ± SE | *W* bulk ± SE |
|---|---|---|---|---|---|
| **Empty-cells** | **3** | **110000** | **94826** | **0.600117 ± 0.006782** | **0.414724 ± 0.019835** |
| **P1** | **1** | **16436** | **16269** | **0.991220** | **1.000000** |
| **P2** | **1** | **49700** | **48510** | **0.026400** | **0.000000** |
| **PBS** | **3** | **551** | **538** | **0.438980 ± 0.220558** | **0.574135 ± 0.211164** |
| PSal-M1 | 3 | 127778 | 122134 | 0.621465 ± 0.004224 | 0.761478 ± 0.003928 |
| PSal-M2 | 3 | 139678 | 132478 | 0.525268 ± 0.011094 | 0.673685 ± 0.011692 |
| PSal-M3 | 3 | 150000 | 139538 | 0.447901 ± 0.009399 | 0.587005 ± 0.006244 |
| PSal-M4 | 3 | 150000 | 141029 | 0.356835 ± 0.010133 | 0.490051 ± 0.007111 |
| PSal-M5 | 3 | 150000 | 141316 | 0.256816 ± 0.002218 | 0.352711 ± 0.003341 |

**Supplementary Table 9 | Summary by label.** Summary of replicate-level weight estimates by biological label, showing the number of replicates, total events used, the mean single-cell weight ⟨*W*⟩ with its standard error, and the bulk-estimated weight *W* derived from ratiometric fluorescence.

#### Per-file (per well)

| File | Label | Events total | Events used | ⟨*W*⟩ | Variance *W* (sample) | *W* bulk |
|---|---|---|---|---|---|---|
| **A1a.csv** | **PBS** | **58** | **58** | **0.875201** | **0.092608** | **0.972961** |
| **A2a.csv** | **Empty-cells** | **10000** | **8414** | **0.607810** | **0.176715** | **0.443119** |
| C10a.csv | PSal-M2C | 45318 | 42860 | 0.547308 | 0.122815 | 0.697045 |
| **C1a.csv** | **PBS** | **396** | **389** | **0.277622** | **0.154466** | **0.495028** |
| **C2a.csv** | **Empty-cells** | **50000** | **43176** | **0.605945** | **0.176008** | **0.424519** |
| **C3a.csv** | **P2** | **49700** | **48510** | **0.026400** | **0.015106** | **0.000000** |
| **C4a.csv** | **P1** | **16436** | **16269** | **0.991220** | **0.003566** | **1.000000** |
| C5a.csv | PSal-M1A | 42028 | 40121 | 0.614831 | 0.115194 | 0.755513 |
| C6a.csv | PSal-M1B | 41776 | 39995 | 0.629311 | 0.112788 | 0.768888 |
| C7a.csv | PSal-M1C | 43974 | 42018 | 0.620254 | 0.112764 | 0.760033 |
| C8a.csv | PSal-M2A | 49168 | 46576 | 0.516466 | 0.124106 | 0.662926 |
| C9a.csv | PSal-M2B | 45192 | 43042 | 0.512032 | 0.125889 | 0.661084 |

| | | | | | | |
|---|---|---|---|---|---|---|
| D1a.csv | PSal-M3A | 50000 | 45394 | 0.464946 | 0.129344 | 0.597408 |
| D2a.csv | PSal-M3B | 50000 | 46834 | 0.446241 | 0.126117 | 0.587787 |
| D3a.csv | PSal-M3C | 50000 | 47310 | 0.432516 | 0.127817 | 0.575819 |
| D4a.csv | PSal-M4A | 50000 | 47809 | 0.336673 | 0.116413 | 0.478486 |
| D5a.csv | PSal-M4B | 50000 | 46758 | 0.368683 | 0.120264 | 0.503001 |
| D6a.csv | PSal-M4C | 50000 | 46462 | 0.365150 | 0.121439 | 0.488665 |
| D7a.csv | PSal-M5A | 50000 | 47384 | 0.258715 | 0.099692 | 0.358477 |
| D8a.csv | PSal-M5B | 50000 | 47664 | 0.252394 | 0.098021 | 0.352751 |
| D9a.csv | PSal-M5C | 50000 | 46268 | 0.259339 | 0.099375 | 0.346905 |
| **E1a.csv** | **PBS** | **97** | **91** | **0.164116** | **0.109756** | **0.254415** |
| **E2a.csv** | **Empty-cells** | **50000** | **43236** | **0.586596** | **0.179191** | **0.376535** |

**Supplementary Table 10 | Per-file single-well statistics.** Per-well statistics for each input file, including the number of events used, the mean single-cell weight ⟨$W$⟩, the unbiased sample variance of $W$, and the bulk-estimated weight derived from ratiometric fluorescence measurements.

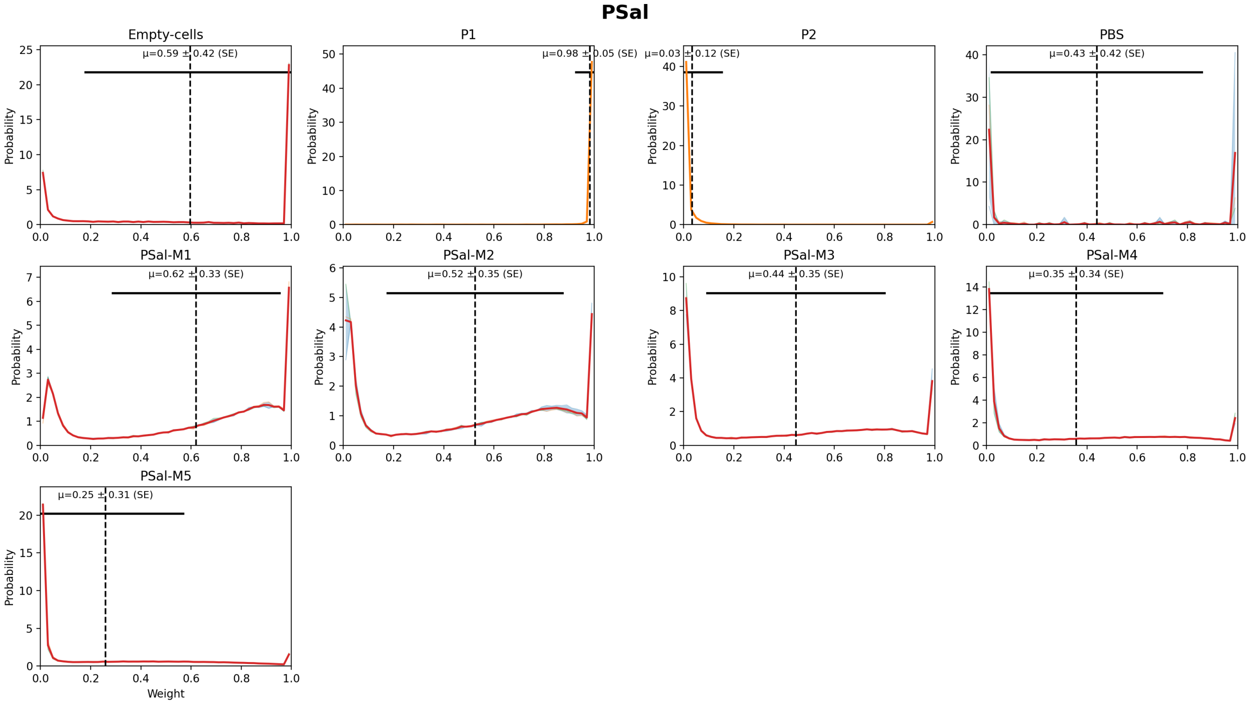

**Supplementary Fig. 4 | Weight distributions by label.** Weight distributions by label with replicate overlays; the shaded band shows the mean ± SD across replicates, the dashed line marks the mean single-cell weight, and the horizontal bar indicates ±SD of $W$ across all gated single cells pooled across replicates for that label.

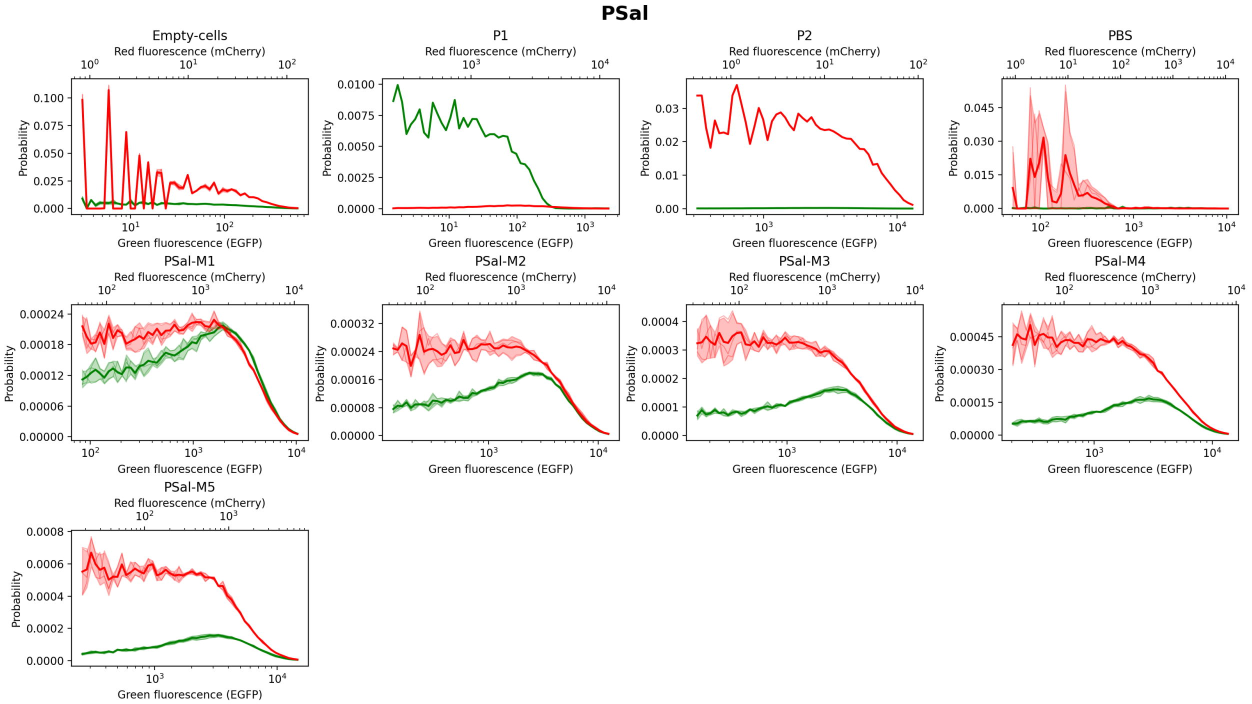

**Supplementary Fig. 5 | Fluorescence distributions by label.** Green and red fluorescence histograms by label on logarithmic *x*-axes with 10^n ticks; replicate histograms are overlaid and the shaded bands show mean ± SD across replicates.

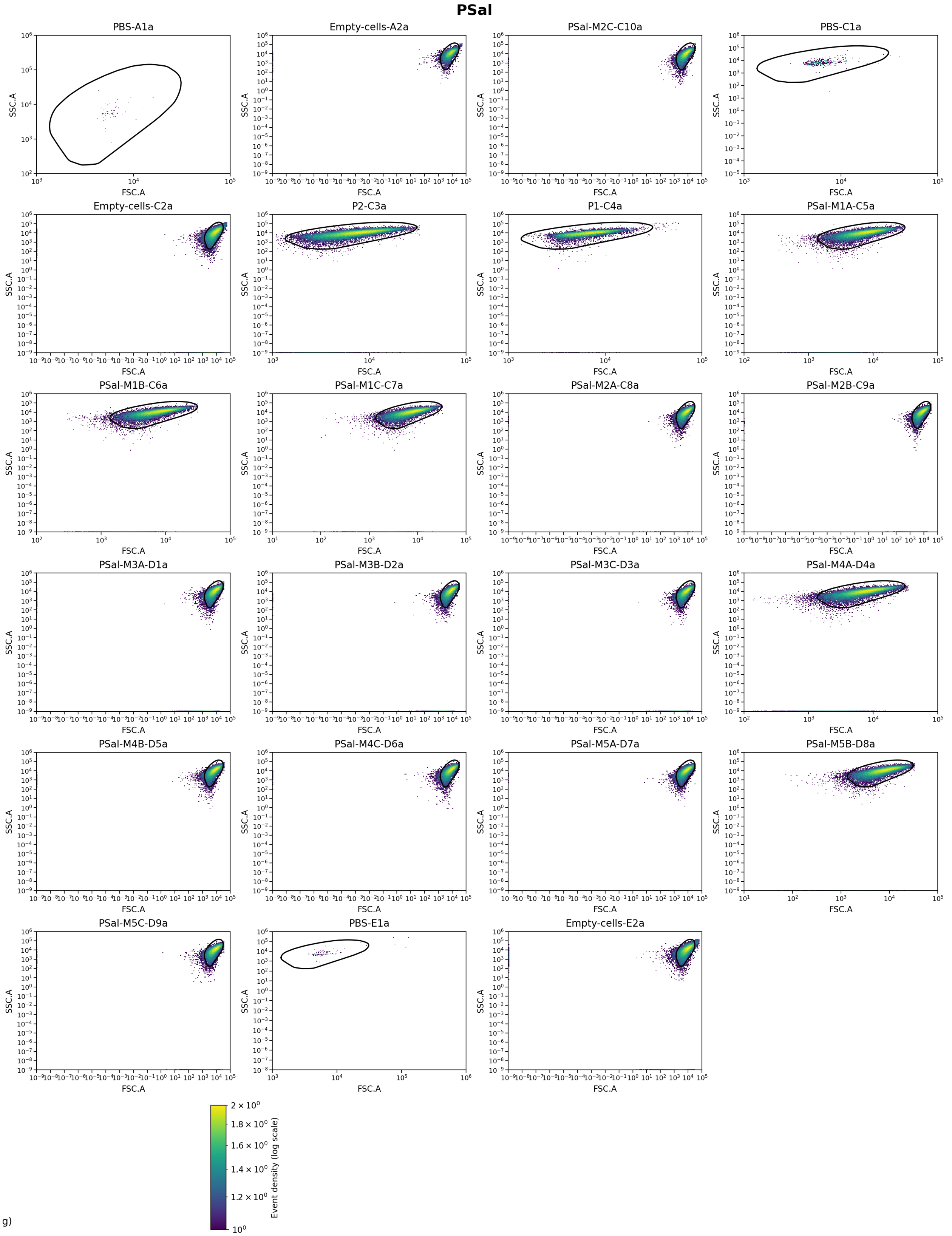

g)

**Supplementary Fig. 6 | FSC–SSC distributions per well.** Two-dimensional FSC vs SSC (log10) density maps for each well before gating, with the non-parametric polygonal cell gate overlaid.

### PTet

#### Experiment parameters and QC.

| Metric | Value |
|---|---|
| λ (from P1/P2 totals) | 1.03312 |
| β (from P1 totals) | 0.00379571 |
| γ | 0.00283982 |
| α spillover (R←G) | 0.000311263 |
| β spillover (G←R) | 0 |
| Background median (green) | 7424.06 |
| Background median (red) | 1.64 |
| Gate quantile (cell gate) | 0.99 |
| Gate $d^2$ threshold | NaN |
| Gate area (log10) | 10.4137 |
| Events (total) | 1111687 |
| Events (gated) | 1004262 |
| Events used | 1004262 |
| Gate acceptance (%) | 90.3368 |
| Used / gated (%) | 100 |
| Fluor green thr (quantile) | 0.999 |
| Fluor green thr (value) | 61910.7 |
| Fluor red thr (quantile) | 0.999 |
| Fluor red thr (value) | 358.234 |
| Median NEG green | 1595.24 |
| Median NEG red | 0 |
| Median POS green (P2) | 1950.54 |
| Median POS red (P1) | 4555.1 |
| Stain index (green) | 0.0213412 |
| Stain index (red) | 48.5556 |
| Residual corr (negatives) | 0.0537823 |
| Dynamic range log10 (green) | 16.3754 |
| Dynamic range log10 (red) | 16.2767 |
| Replicate hist mean r | 0.952381 |
| Replicate hist mean RMSD | 0.979104 |

**Supplementary Table 11 | Experiment parameters and QC.** Experiment-level parameters and quality-control metrics for the FACS analysis, including background estimates, gating properties, event counts, stain indices and residual correlation after spectral compensation.

#### Summary by label (replicate means ± SE)

| Label | Replicates (n) | Events total (sum) | Events used (sum) | ⟨*W*⟩ ± SE | *W* bulk ± SE |
|---|---|---|---|---|---|
| **Empty-cells** | **4** | **200000** | **180598** | **0.854558 ± 0.145390** | **0.722364 ± 0.241049** |
| **P1** | **1** | **50000** | **46670** | **0.996783** | **1.000000** |
| **P2** | **1** | **50000** | **43544** | **0.407511** | **0.000000** |
| **PBS** | **4** | **11687** | **11563** | **0.948375 ± 0.044069** | **0.736589 ± 0.108700** |
| PTet-M0 | 1 | 50000 | 45983 | 0.908149 | 0.837026 |
| PTet-M1 | 3 | 150000 | 138323 | 0.870998 ± 0.011173 | 0.776957 ± 0.017408 |
| PTet-M2 | 3 | 150000 | 136061 | 0.823460 ± 0.011561 | 0.699400 ± 0.018714 |
| PTet-M3 | 3 | 150000 | 135641 | 0.767773 ± 0.008604 | 0.600672 ± 0.012850 |
| PTet-M4 | 3 | 150000 | 134080 | 0.737612 ± 0.007400 | 0.531450 ± 0.007250 |
| PTet-M5 | 3 | 150000 | 131799 | 0.815534 ± 0.014107 | 0.599221 ± 0.025874 |

**Supplementary Table 12 | Summary by label.** Summary of replicate-level weight estimates by biological label, showing the number of replicates, total events used, the mean single-cell weight ⟨*W*⟩ with its standard error, and the bulk-estimated weight *W* derived from ratiometric fluorescence.

#### Per-file (per well)

| File | Label | Events total | Events used | $\langle W \rangle$ | Variance $W$ (sample) | $W$ bulk |
|---|---|---|---|---|---|---|
| **A1b.csv** | **PBS** | **11299** | **11177** | **1.000000** | **0.000000** | **1.003937** |
| **A2b.csv** | **Empty-cells** | **50000** | **42392** | **0.418388** | **0.240238** | **0.000288** |
| **C1b.csv** | **PBS** | **60** | **60** | **0.993063** | **0.002887** | **0.642135** |
| **C2b.csv** | **Empty-cells** | **50000** | **45895** | **0.999972** | **0.000010** | **0.987954** |
| E10b.csv | PTet-M2B | 50000 | 45313 | 0.806746 | 0.115741 | 0.671418 |
| E11b.csv | PTet-M2C | 50000 | 45565 | 0.817981 | 0.108680 | 0.691866 |
| **E1b.csv** | **PBS** | **68** | **67** | **0.983897** | **0.007101** | **0.802530** |
| **E2b.csv** | **Empty-cells** | **50000** | **46102** | **0.999938** | **0.000033** | **0.974439** |
| **E3b.csv** | **P2** | **50000** | **43544** | **0.407511** | **0.238670** | **0.000000** |
| **E4b.csv** | **P1** | **50000** | **46670** | **0.996783** | **0.002416** | **1.000000** |
| E5b.csv | PTet-M0 | 50000 | 45983 | 0.908149 | 0.055755 | 0.837026 |
| E6b.csv | PTet-M1A | 50000 | 46367 | 0.882943 | 0.074620 | 0.798025 |
| E7b.csv | PTet-M1B | 50000 | 45526 | 0.848671 | 0.093244 | 0.742419 |
| E8b.csv | PTet-M1C | 50000 | 46430 | 0.881380 | 0.076053 | 0.790427 |
| E9b.csv | PTet-M2A | 50000 | 45183 | 0.845653 | 0.097827 | 0.734917 |
| F1b.csv | PTet-M3A | 50000 | 45107 | 0.784035 | 0.127261 | 0.626255 |
| F2b.csv | PTet-M3B | 50000 | 45254 | 0.764516 | 0.135364 | 0.589990 |
| F3b.csv | PTet-M3C | 50000 | 45280 | 0.754767 | 0.137554 | 0.585770 |
| F4b.csv | PTet-M4A | 50000 | 44862 | 0.751511 | 0.142927 | 0.545769 |
| F5b.csv | PTet-M4B | 50000 | 44743 | 0.735067 | 0.149039 | 0.526271 |
| F6b.csv | PTet-M4C | 50000 | 44475 | 0.726257 | 0.152459 | 0.522310 |
| F7b.csv | PTet-M5A | 50000 | 44343 | 0.810313 | 0.122156 | 0.579472 |
| F8b.csv | PTet-M5B | 50000 | 43728 | 0.794132 | 0.130784 | 0.567672 |
| F9b.csv | PTet-M5C | 50000 | 43728 | 0.842156 | 0.104398 | 0.650519 |
| **G1b.csv** | **PBS** | **260** | **259** | **0.816540** | **0.143209** | **0.497753** |
| **G2b.csv** | **Empty-cells** | **50000** | **46209** | **0.999935** | **0.000026** | **0.926776** |

**Supplementary Table 13 | Per-file single-well statistics.** Per-well statistics for each input file, including the number of events used, the mean single-cell weight $\langle W \rangle$, the unbiased sample variance of $W$, and the bulk-estimated weight derived from ratiometric fluorescence measurements.

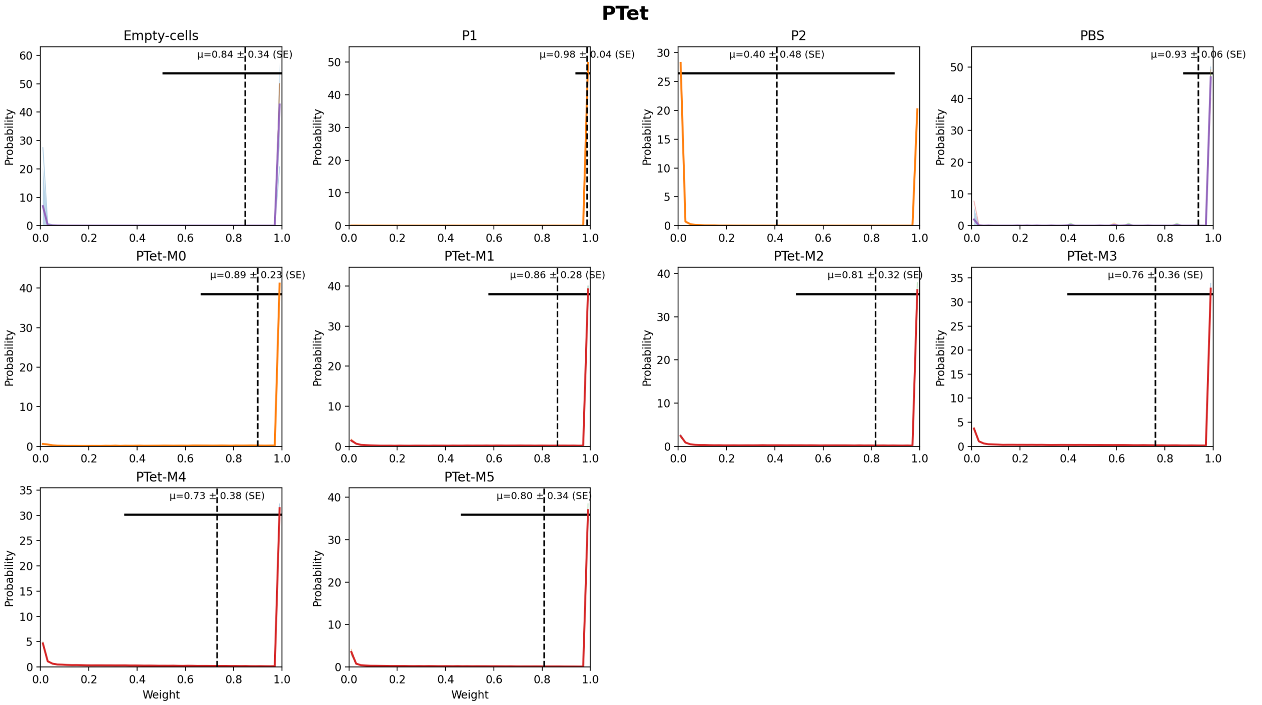

**Supplementary Fig. 7 | Weight distributions by label.** Weight distributions by label with replicate overlays; the shaded band shows the mean ± SD across replicates, the dashed line marks the mean single-cell weight, and the horizontal bar indicates ±SD of $W$ across all gated single cells pooled across replicates for that label.

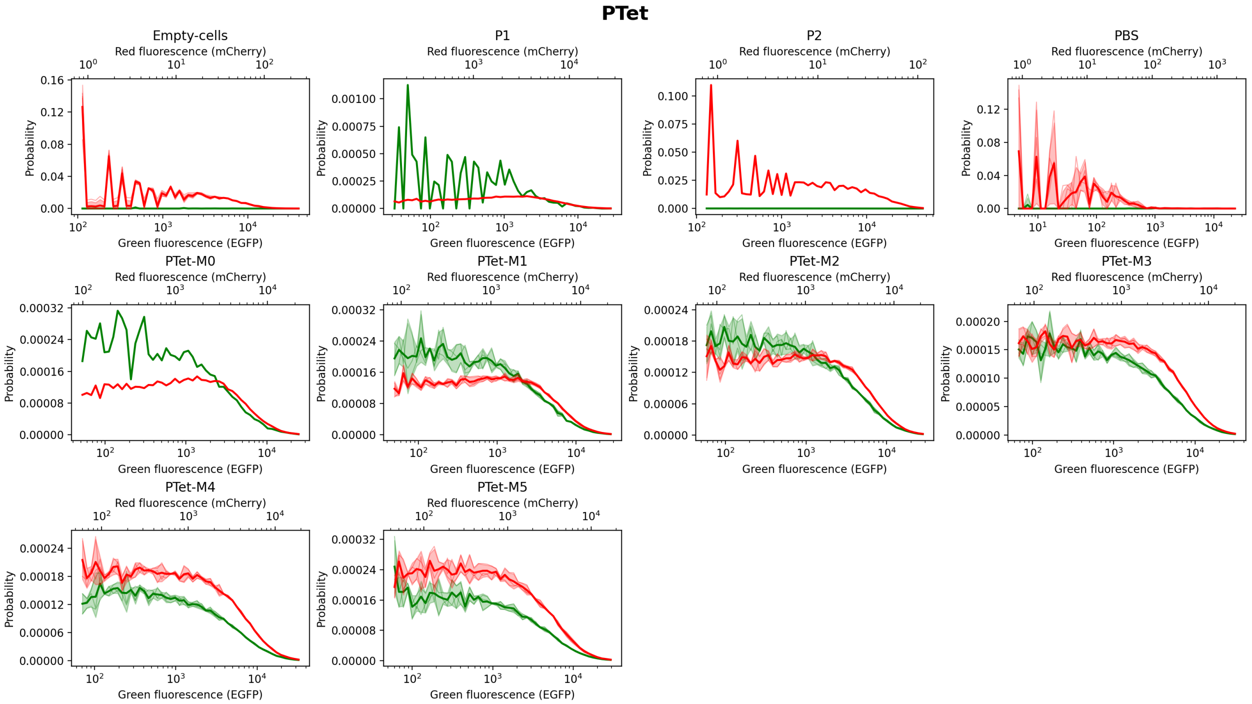

**Supplementary Fig. 8 | Fluorescence distributions by label.** Green and red fluorescence histograms by label on logarithmic *x*-axes with 10^n ticks; replicate histograms are overlaid and the shaded bands show mean ± SD across replicates.

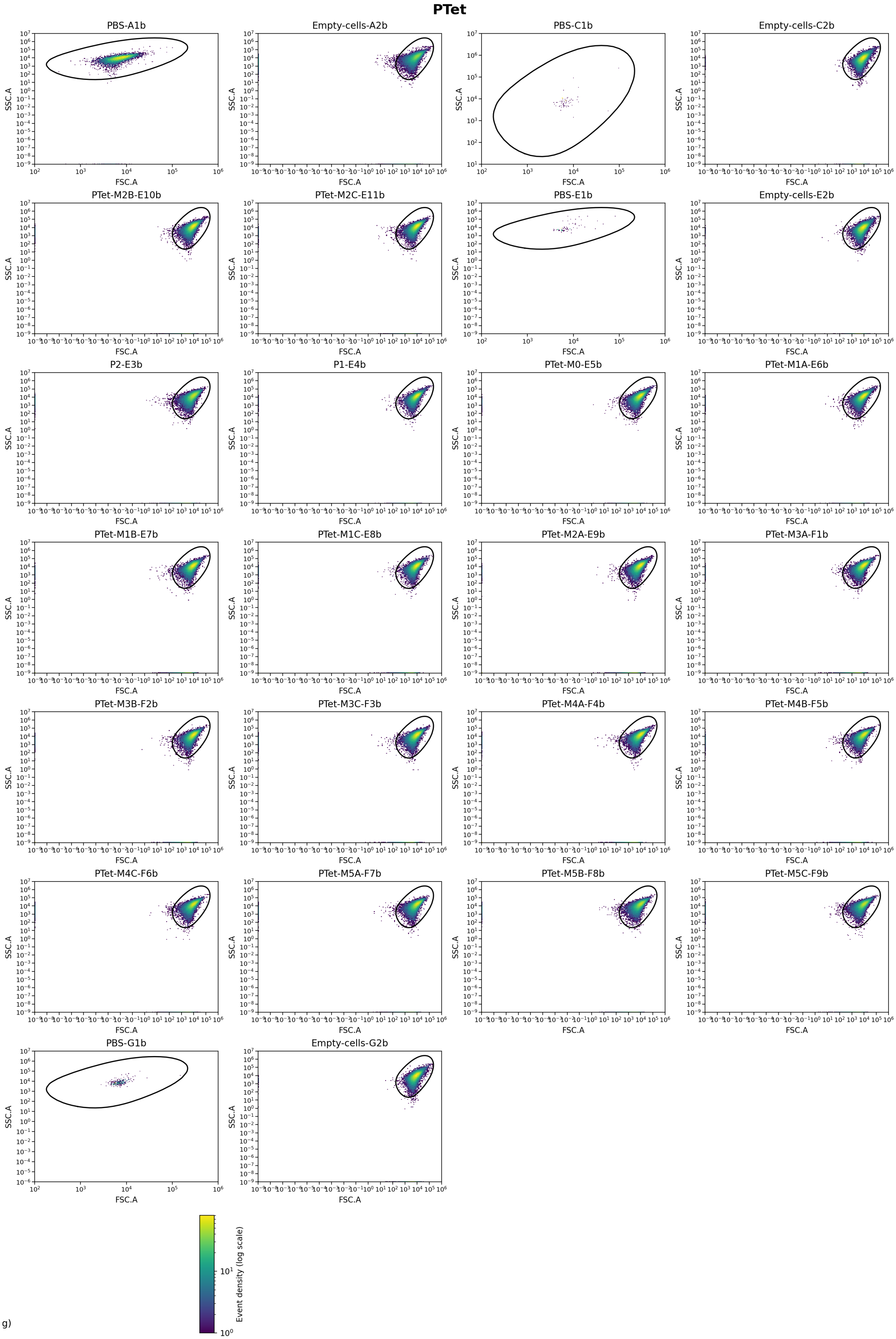
PTet
PBS-A1b
Empty-cells-A2b
PBS-C1b
Empty-cells-C2b
PTet-M2B-E10b
PTet-M2C-E11b
PBS-E1b
Empty-cells-E2b
P2-E3b
P1-E4b
PTet-M0-E5b
PTet-M1A-E6b
PTet-M1B-E7b
PTet-M1C-E8b
PTet-M2A-E9b
PTet-M3A-F1b
PTet-M3B-F2b
PTet-M3C-F3b
PTet-M4A-F4b
PTet-M4B-F5b
PTet-M4C-F6b
PTet-M5A-F7b
PTet-M5B-F8b
PTet-M5C-F9b
PBS-G1b
Empty-cells-G2b
SSC.A
FSC.A
Event density (log scale)

g)

**Supplementary Fig. 9 | FSC–SSC distributions per well.** Two-dimensional FSC vs SSC (log10) density maps for each well before gating, with the non-parametric polygonal cell gate overlaid.

### PBetI

#### Experiment parameters and QC.

| Metric | Value |
|---|---|
| λ (from P1/P2 totals) | 0.399697 |
| β (from P1 totals) | 0.0448894 |
| γ | 0.00223262 |
| α spillover (R←G) | 5.1057e-05 |
| β spillover (G←R) | 0.00139981 |
| Background median (green) | 34.96 |
| Background median (red) | 0.81 |
| Gate quantile (cell gate) | 0.99 |
| Gate d² threshold | NaN |
| Gate area (log10) | 3.99742 |
| Events (total) | 851071 |
| Events (gated) | 806514 |
| Events used | 806514 |
| Gate acceptance (%) | 94.7646 |
| Used / gated (%) | 100 |
| Fluor green thr (quantile) | 0.999 |
| Fluor green thr (value) | 273.575 |
| Fluor red thr (quantile) | 0.999 |
| Fluor red thr (value) | 127.482 |
| Median NEG green | 1.47691 |
| Median NEG red | 0.805344 |
| Median POS green (P2) | 4420.15 |
| Median POS red (P1) | 1304.91 |
| Stain index (green) | 50.5849 |
| Stain index (red) | 23.9918 |
| Residual corr (negatives) | 0.355317 |
| Dynamic range log10 (green) | 16.2846 |
| Dynamic range log10 (red) | 16.1754 |
| Replicate hist mean r | 0.955774 |
| Replicate hist mean RMSD | 0.282944 |

**Supplementary Table 14 | Experiment parameters and QC.** Experiment-level parameters and quality-control metrics for the FACS analysis, including background estimates, gating properties, event counts, stain indices and residual correlation after spectral compensation.

#### Summary by label (replicate means ± SE)

| Label | Replicates (n) | Events total (sum) | Events used (sum) | ⟨*W*⟩ ± SE | *W* bulk ± SE |
|---|---|---|---|---|---|
| **Empty-cells** | **2** | **100000** | **95603** | **0.658640 ± 0.003308** | **0.584545 ± 0.012858** |
| **P1** | **1** | **50000** | **47807** | **0.969545** | **1.000000** |
| **P2** | **1** | **50000** | **48124** | **0.052981** | **0.000000** |
| **PBS** | **2** | **1071** | **277** | **0.635273 ± 0.067489** | **0.691230 ± 0.009609** |
| PBetI-M0 | 1 | 50000 | 47486 | 0.824869 | 0.855944 |
| PBetI-M1 | 3 | 150000 | 141207 | 0.726688 ± 0.009509 | 0.762497 ± 0.013258 |
| PBetI-M2 | 3 | 150000 | 142215 | 0.608623 ± 0.004750 | 0.641635 ± 0.004347 |
| PBetI-M3 | 3 | 150000 | 142047 | 0.585490 ± 0.006848 | 0.620717 ± 0.006001 |
| PBetI-M4 | 3 | 150000 | 141748 | 0.552173 ± 0.000666 | 0.571653 ± 0.000418 |

**Supplementary Table 15 | Summary by label.** Summary of replicate-level weight estimates by biological label, showing the number of replicates, total events used, the mean single-cell weight ⟨*W*⟩ with its standard error, and the bulk-estimated weight *W* derived from ratiometric fluorescence.

#### Per-file (per well)

| File | Label | Events total | Events used | ⟨*W*⟩ | Variance *W* (sample) | *W* bulk |
|---|---|---|---|---|---|---|
| **A1c.csv** | **PBS** | **804** | **13** | **0.702761** | **0.034135** | **0.681621** |
| **A2c.csv** | **Empty-cells** | **50000** | **47871** | **0.661947** | **0.160957** | **0.571687** |
| **A3c.csv** | **P2** | **50000** | **48124** | **0.052981** | **0.041306** | **0.000000** |
| **A4c.csv** | **P1** | **50000** | **47807** | **0.969545** | **0.016527** | **1.000000** |
| A9c.csv | PBetI-M0 | 50000 | 47486 | 0.824869 | 0.077477 | 0.855944 |
| B10c.csv | PBetI-M4A | 50000 | 46681 | 0.552497 | 0.152167 | 0.572438 |
| B11c.csv | PBetI-M4B | 50000 | 46847 | 0.550891 | 0.152907 | 0.571012 |
| B12c.csv | PBetI-M4C | 50000 | 48220 | 0.553130 | 0.147847 | 0.571510 |
| B1c.csv | PBetI-M1A | 50000 | 46815 | 0.745509 | 0.106319 | 0.788963 |
| B2c.csv | PBetI-M1B | 50000 | 47275 | 0.719646 | 0.112763 | 0.750684 |
| B3c.csv | PBetI-M1C | 50000 | 47117 | 0.714910 | 0.113084 | 0.747843 |
| B4c.csv | PBetI-M2A | 50000 | 47263 | 0.600050 | 0.138848 | 0.633705 |
| B5c.csv | PBetI-M2B | 50000 | 47461 | 0.616452 | 0.136057 | 0.648688 |
| B6c.csv | PBetI-M2C | 50000 | 47491 | 0.609366 | 0.138347 | 0.642511 |
| B7c.csv | PBetI-M3A | 50000 | 47193 | 0.571993 | 0.146132 | 0.608798 |
| B8c.csv | PBetI-M3B | 50000 | 47179 | 0.594252 | 0.144799 | 0.625456 |
| B9c.csv | PBetI-M3C | 50000 | 47675 | 0.590224 | 0.143679 | 0.627899 |
| **E1c.csv** | **PBS** | **267** | **264** | **0.567784** | **0.117139** | **0.700839** |
| **E2c.csv** | **Empty-cells** | **50000** | **47732** | **0.655332** | **0.161177** | **0.597404** |

**Supplementary Table 16 | Per-file single-well statistics.** Per-well statistics for each input file, including the number of events used, the mean single-cell weight ⟨*W*⟩, the unbiased sample variance of *W*, and the bulk-estimated weight derived from ratiometric fluorescence measurements.

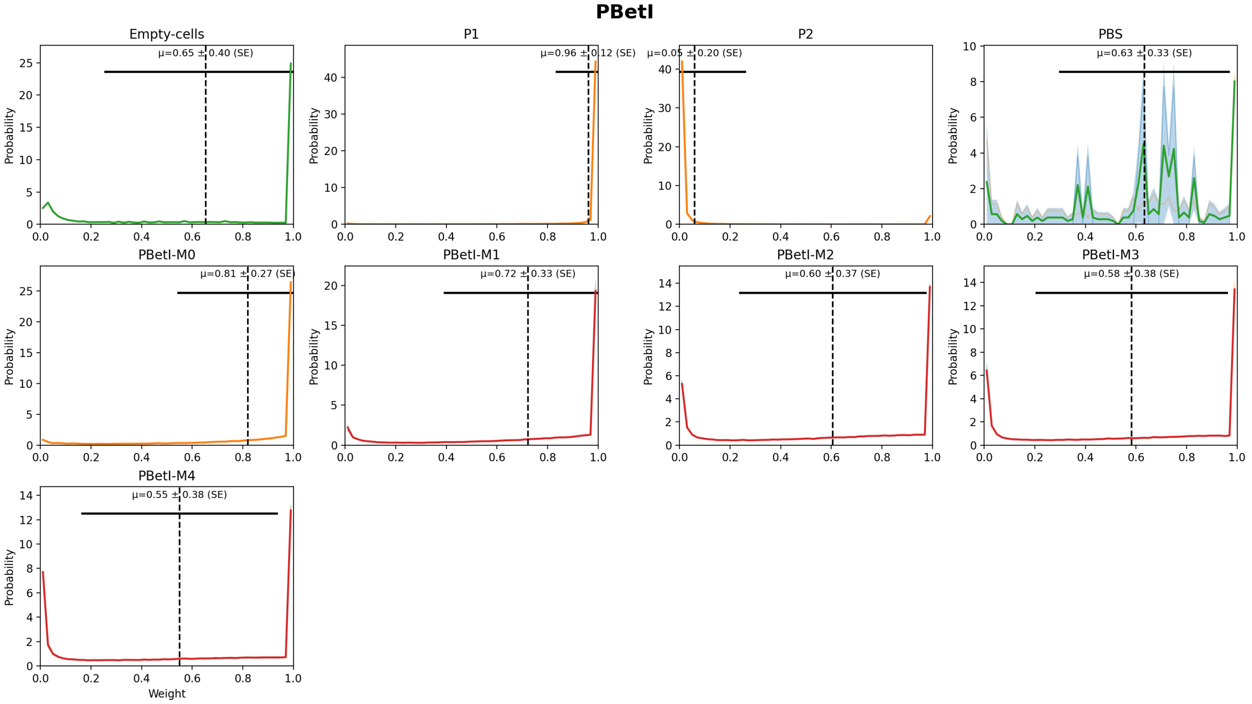

**Supplementary Fig. 10 | Weight distributions by label.** Weight distributions by label with replicate overlays; the shaded band shows the mean ± SD across replicates, the dashed line marks the mean single-cell weight, and the horizontal bar indicates ±SD of *W* across all gated single cells pooled across replicates for that label.

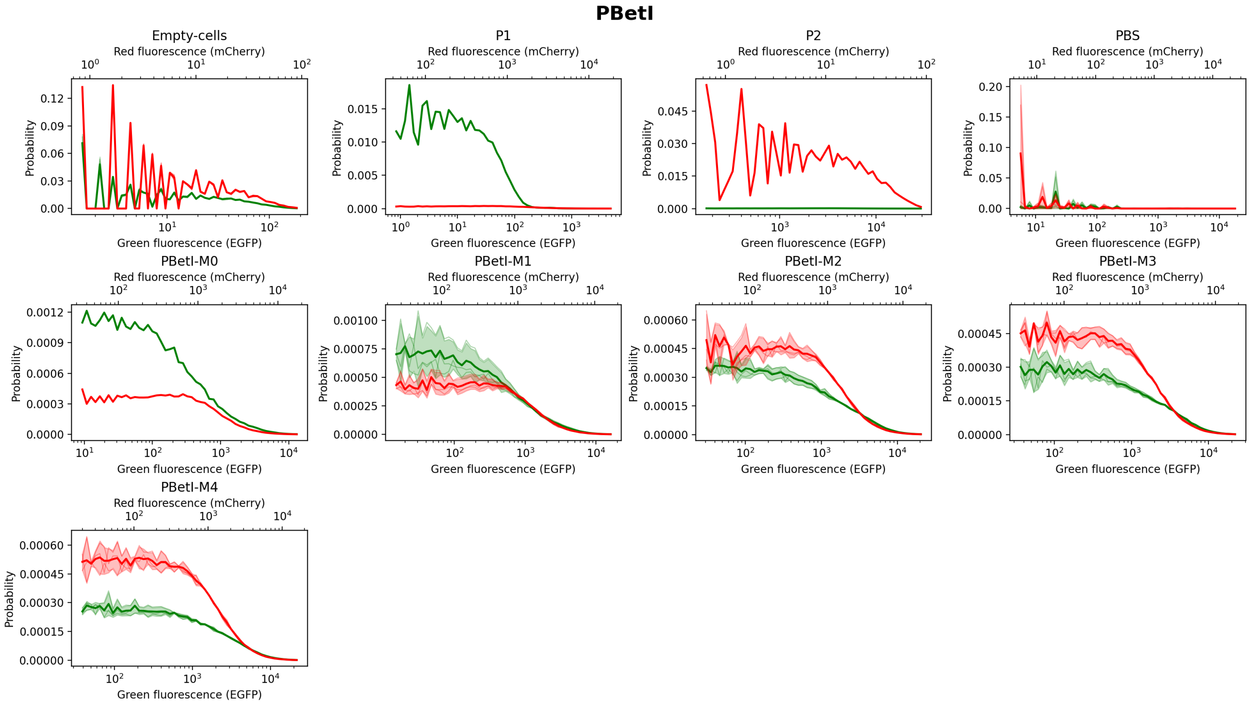

**Supplementary Fig. 11 | Fluorescence distributions by label.** Green and red fluorescence histograms by label on logarithmic *x*-axes with 10^n ticks; replicate histograms are overlaid and the shaded bands show mean ± SD across replicates.

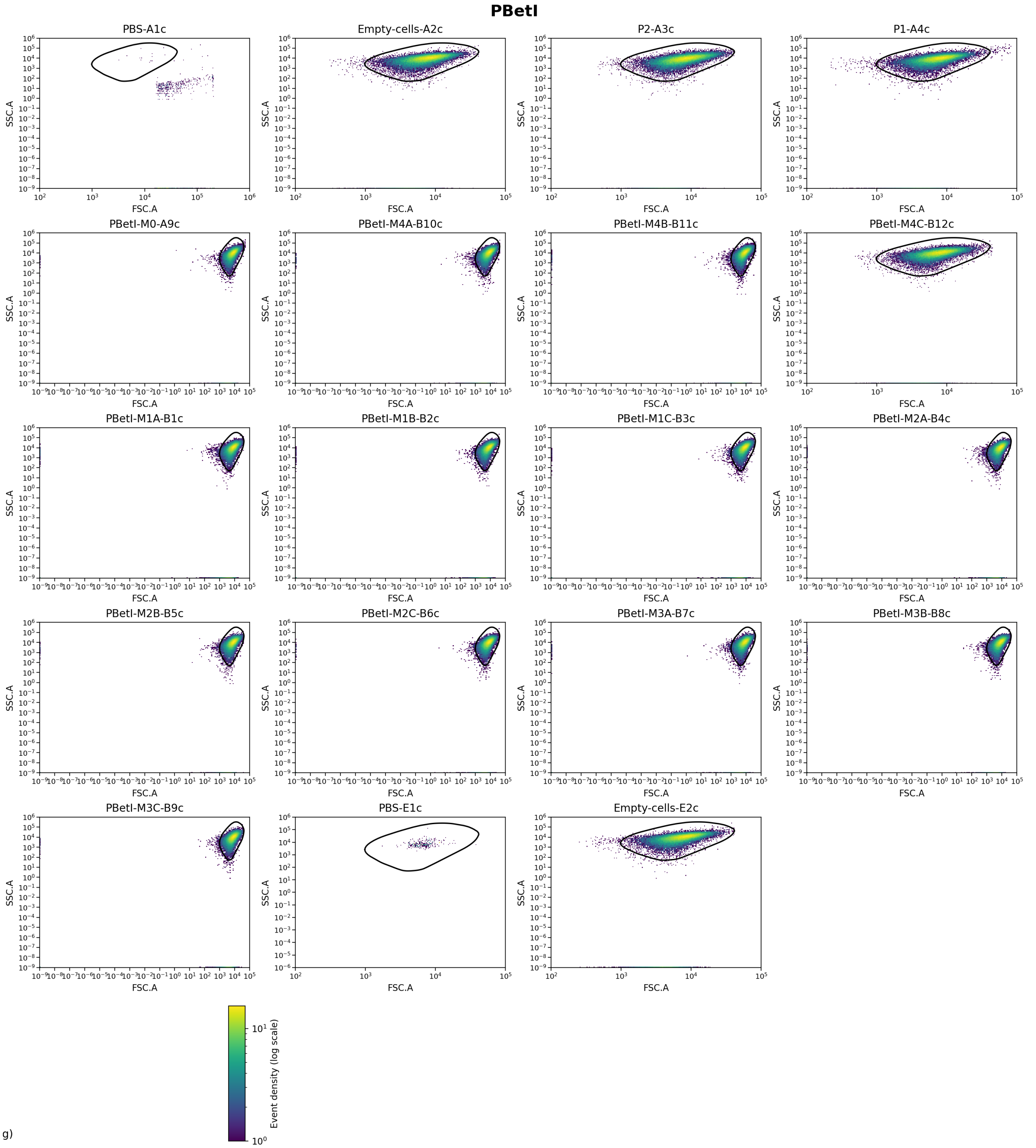

**Supplementary Fig. 12 | FSC–SSC distributions per well.** Two-dimensional FSC vs SSC (log10) density maps for each well before gating, with the non-parametric polygonal cell gate overlaid.

### PBAD

#### Experiment parameters and QC.

| Metric | Value |
|---|---|
| λ (from P1/P2 totals) | 1.2188 |
| β (from P1 totals) | 0.0262023 |
| γ | 0.0016858 |
| α spillover (R←G) | 0.000140921 |
| β spillover (G←R) | 0.00708172 |
| Background median (green) | 73.5 |

| | |
|---|---|
| Background median (red) | 3.2 |
| Gate quantile (cell gate) | 0.99 |
| Gate d² threshold | NaN |
| Gate area (log10) | 7.14389 |
| Events (total) | 280551 |
| Events (gated) | 254362 |
| Events used | 254362 |
| Gate acceptance (%) | 90.6652 |
| Used / gated (%) | 100 |
| Fluor green thr (quantile) | 0.999 |
| Fluor green thr (value) | 779.244 |
| Fluor red thr (quantile) | 0.999 |
| Fluor red thr (value) | 376.552 |
| Median NEG green | 14.25 |
| Median NEG red | 0.795879 |
| Median POS green (P2) | 6620.59 |
| Median POS red (P1) | 8436.75 |
| Stain index (green) | 26.3649 |
| Stain index (red) | 66.5738 |
| Residual corr (negatives) | 0.146794 |
| Dynamic range log10 (green) | 16.3148 |
| Dynamic range log10 (red) | 16.3836 |
| Replicate hist mean r | 0.919129 |
| Replicate hist mean RMSD | 0.908273 |

**Supplementary Table 17 | Experiment parameters and QC.** Experiment-level parameters and quality-control metrics for the FACS analysis, including background estimates, gating properties, event counts, stain indices and residual correlation after spectral compensation.

#### Summary by label (replicate means ± SE)

| Label | Replicates (n) | Events total (sum) | Events used (sum) | ⟨$W$⟩ ± SE | $W$ bulk ± SE |
|---|---|---|---|---|---|
| **Empty-cells** | **3** | **110000** | **97952** | **0.524964 ± 0.006894** | **0.144415 ± 0.010819** |
| **P1** | **1** | **10000** | **9345** | **0.944354** | **1.000000** |
| **P2** | **1** | **10000** | **9299** | **0.053109** | **0.000000** |
| PBAD-M1 | 3 | 30000 | 26958 | 0.572758 ± 0.012339 | 0.593192 ± 0.013801 |
| PBAD-M2 | 3 | 30000 | 27593 | 0.494345 ± 0.005468 | 0.497289 ± 0.004358 |
| PBAD-M3 | 3 | 30000 | 27475 | 0.446934 ± 0.005600 | 0.435433 ± 0.008874 |
| PBAD-M4 | 3 | 30000 | 27261 | 0.390903 ± 0.002993 | 0.352262 ± 0.004998 |
| PBAD-M5 | 3 | 30000 | 27937 | 0.315793 ± 0.019376 | 0.238555 ± 0.017955 |
| **PBS** | **3** | **551** | **542** | **0.395380 ± 0.227715** | **0.388696 ± 0.257212** |

**Supplementary Table 18 | Summary by label.** Summary of replicate-level weight estimates by biological label, showing the number of replicates, total events used, the mean single-cell weight ⟨$W$⟩ with its standard error, and the bulk-estimated weight $W$ derived from ratiometric fluorescence.

#### Per-file (per well)

| File | Label | Events total | Events used | ⟨$W$⟩ | Variance $W$ (sample) | $W$ bulk |
|---|---|---|---|---|---|---|
| A10a.csv | PBAD-M2C | 10000 | 9120 | 0.483624 | 0.131176 | 0.491735 |
| **A1a.csv** | **PBS** | **58** | **58** | **0.847819** | **0.113614** | **0.898196** |
| **A2a.csv** | **Empty-cells** | **10000** | **8749** | **0.533407** | **0.206984** | **0.161716** |
| **A3a.csv** | **P2** | **10000** | **9299** | **0.053109** | **0.047822** | **0.000000** |
| **A4a.csv** | **P1** | **10000** | **9345** | **0.944354** | **0.023977** | **1.000000** |

| | | | | | | |
|---|---|---|---|---|---|---|
| A5a.csv | PBAD-M1A | 10000 | 9062 | 0.593169 | 0.126755 | 0.611002 |
| A6a.csv | PBAD-M1B | 10000 | 8802 | 0.574567 | 0.128515 | 0.602551 |
| A7a.csv | PBAD-M1C | 10000 | 9094 | 0.550539 | 0.126001 | 0.566024 |
| A8a.csv | PBAD-M2A | 10000 | 9123 | 0.501580 | 0.130148 | 0.505883 |
| A9a.csv | PBAD-M2B | 10000 | 9350 | 0.497831 | 0.131787 | 0.494248 |
| B1a.csv | PBAD-M3A | 10000 | 9304 | 0.457281 | 0.128434 | 0.453014 |
| B2a.csv | PBAD-M3B | 10000 | 9037 | 0.438046 | 0.132306 | 0.424533 |
| B3a.csv | PBAD-M3C | 10000 | 9134 | 0.445473 | 0.130071 | 0.428753 |
| B4a.csv | PBAD-M4A | 10000 | 9054 | 0.391153 | 0.133928 | 0.342341 |
| B5a.csv | PBAD-M4B | 10000 | 9022 | 0.395957 | 0.133753 | 0.356161 |
| B6a.csv | PBAD-M4C | 10000 | 9185 | 0.385599 | 0.127336 | 0.358284 |
| B7a.csv | PBAD-M5A | 10000 | 9367 | 0.277828 | 0.123550 | 0.202662 |
| B8a.csv | PBAD-M5B | 10000 | 9291 | 0.341506 | 0.138776 | 0.255563 |
| B9a.csv | PBAD-M5C | 10000 | 9279 | 0.328046 | 0.131170 | 0.257440 |
| **C1a.csv** | **PBS** | **396** | **393** | **0.214292** | **0.124920** | **0.195441** |
| **C2a.csv** | **Empty-cells** | **50000** | **44529** | **0.530182** | **0.207439** | **0.147018** |
| **E1a.csv** | **PBS** | **97** | **91** | **0.124031** | **0.085135** | **0.072451** |
| **E2a.csv** | **Empty-cells** | **50000** | **44674** | **0.511302** | **0.209197** | **0.124510** |

**Supplementary Table 19 | Per-file single-well statistics.** Per-well statistics for each input file, including the number of events used, the mean single-cell weight ⟨*W*⟩, the unbiased sample variance of *W*, and the bulk-estimated weight derived from ratiometric fluorescence measurements.

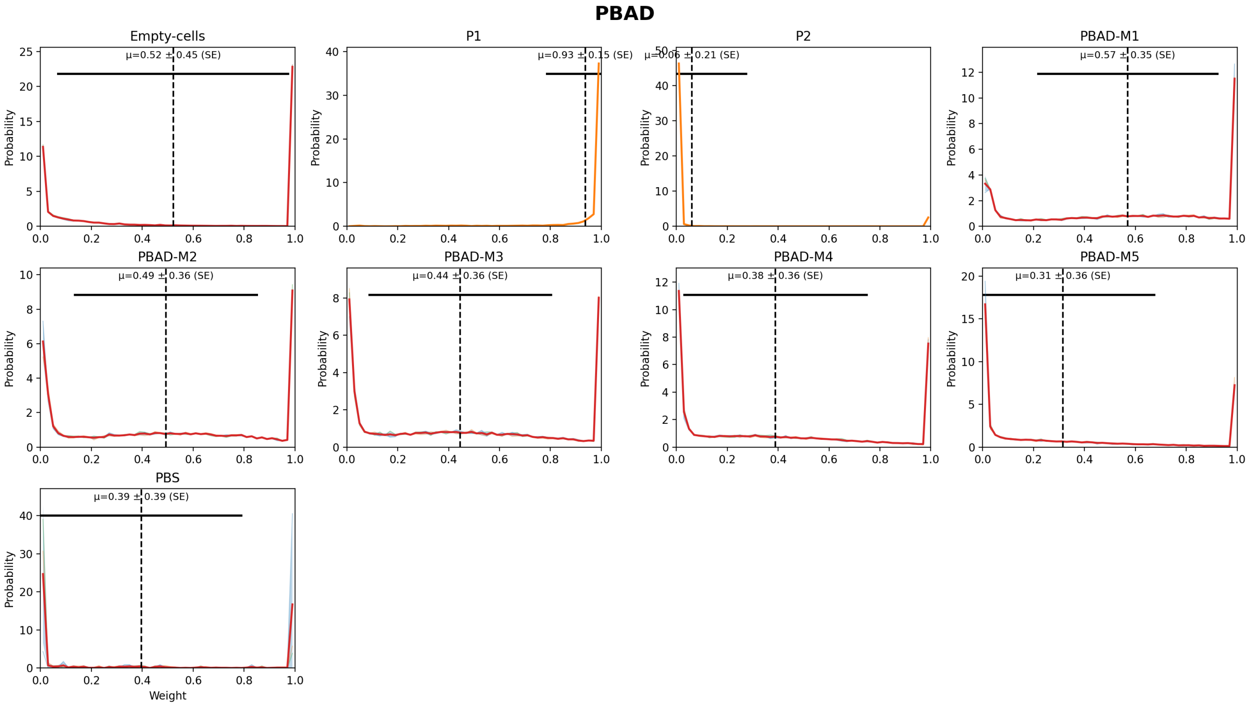

**Supplementary Fig. 13 | Weight distributions by label.** Weight distributions by label with replicate overlays; the shaded band shows the mean ± SD across replicates, the dashed line marks the mean single-cell weight, and the horizontal bar indicates ±SD of *W* across all gated single cells pooled across replicates for that label.

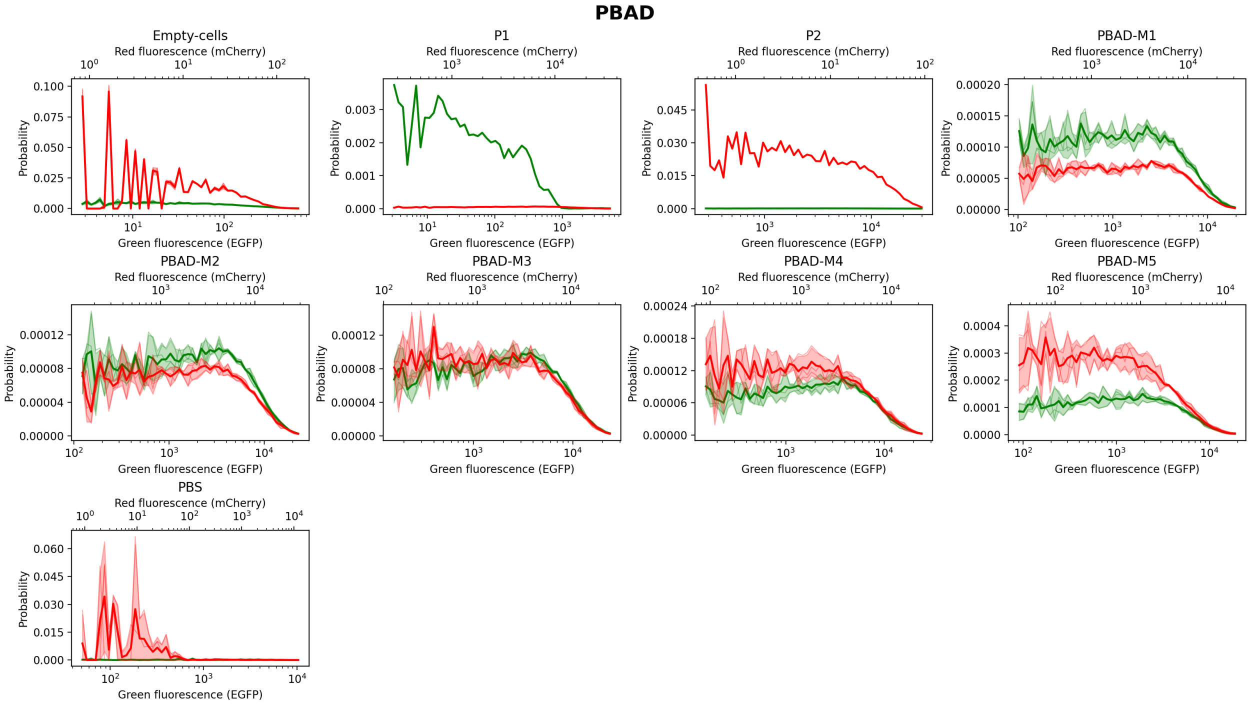

**Supplementary Fig. 14 | Fluorescence distributions by label.** Green and red fluorescence histograms by label on logarithmic *x*-axes with 10^n ticks; replicate histograms are overlaid and the shaded bands show mean ± SD across replicates.

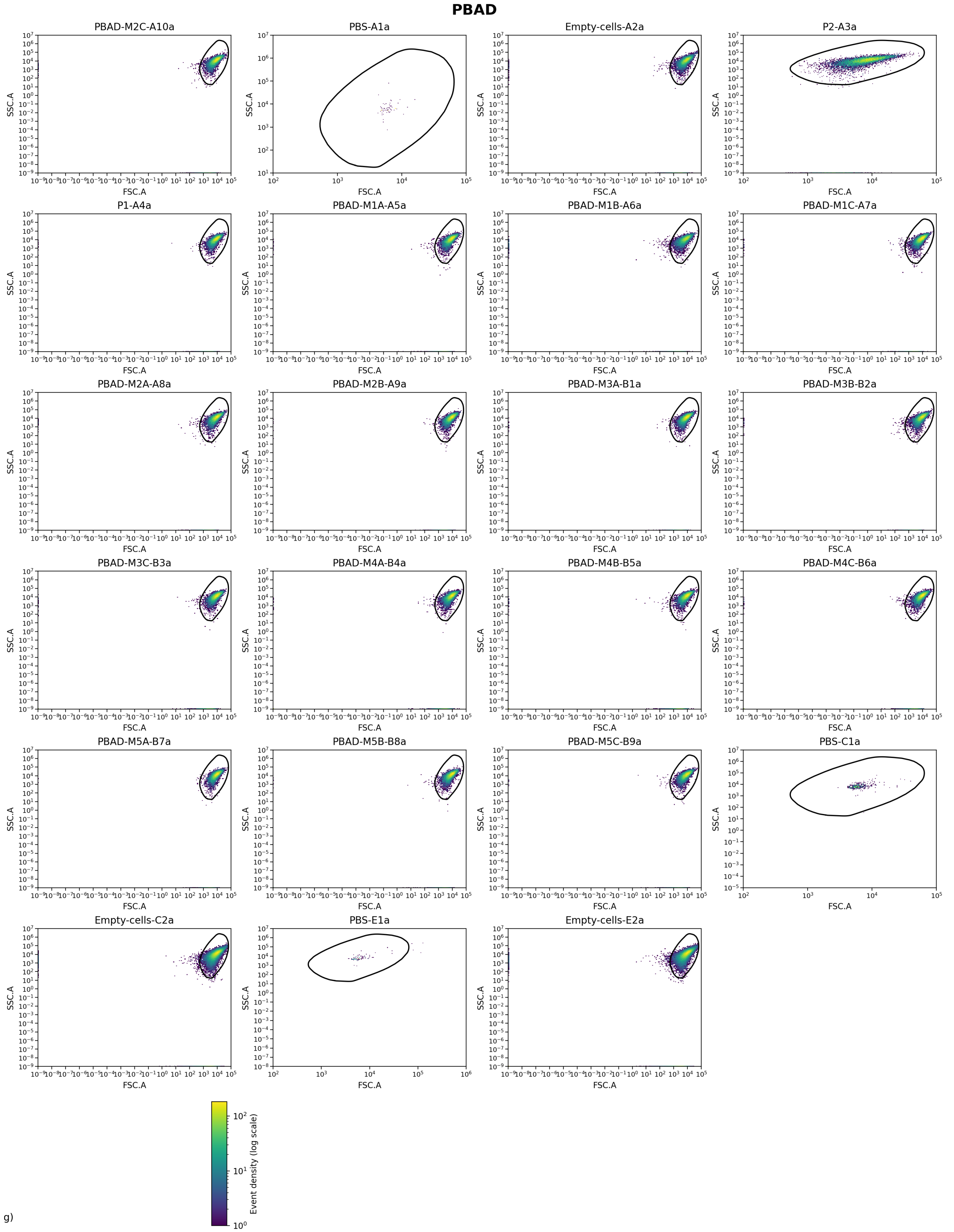

**Supplementary Fig. 15 | FSC–SSC distributions per well.** Two-dimensional FSC vs SSC (log10) density maps for each well before gating, with the non-parametric polygonal cell gate overlaid.

### PLux

#### Experiment parameters and QC.

| Metric | Value |
|---|---|
| λ (from P1/P2 totals) | 1.20277 |
| β (from P1 totals) | 0.0232985 |
| γ | 0.00184981 |
| α spillover (R←G) | 0.000251712 |
| β spillover (G←R) | 0.0046989 |
| Background median (green) | 73.5 |
| Background median (red) | 3.2 |
| Gate quantile (cell gate) | 0.99 |
| Gate d² threshold | NaN |
| Gate area (log10) | 4.96308 |
| Events (total) | 960551 |
| Events (gated) | 882690 |
| Events used | 882690 |
| Gate acceptance (%) | 91.8941 |
| Used / gated (%) | 100 |
| Fluor green thr (quantile) | 0.999 |
| Fluor green thr (value) | 734.481 |
| Fluor red thr (quantile) | 0.999 |
| Fluor red thr (value) | 330.803 |
| Median NEG green | 14.25 |
| Median NEG red | 0.792638 |
| Median POS green (P2) | 5258.13 |
| Median POS red (P1) | 6680.81 |
| Stain index (green) | 21.2939 |
| Stain index (red) | 58.027 |
| Residual corr (negatives) | 0.143274 |
| Dynamic range log10 (green) | 16.3737 |
| Dynamic range log10 (red) | 16.3031 |
| Replicate hist mean r | 0.916348 |
| Replicate hist mean RMSD | 0.898835 |

**Supplementary Table 20 | Experiment parameters and QC.** Experiment-level parameters and quality-control metrics for the FACS analysis, including background estimates, gating properties, event counts, stain indices and residual correlation after spectral compensation.

#### Summary by label (replicate means ± SE)

| Label | Replicates (n) | Events total (sum) | Events used (sum) | ⟨*W*⟩ ± SE | *W* bulk ± SE |
|---|---|---|---|---|---|
| **Empty-cells** | **3** | **110000** | **97564** | **0.524674 ± 0.006900** | **0.144240 ± 0.010400** |
| **P1** | **1** | **50000** | **47391** | **0.953803** | **1.000000** |
| **P2** | **1** | **50000** | **47295** | **0.034956** | **0.000000** |
| **PBS** | **3** | **551** | **542** | **0.395605 ± 0.227683** | **0.388967 ± 0.256293** |
| PLux-M1 | 3 | 150000 | 139454 | 0.594812 ± 0.005805 | 0.604927 ± 0.008133 |
| PLux-M2 | 3 | 150000 | 137144 | 0.453447 ± 0.020915 | 0.443713 ± 0.023540 |
| PLux-M3 | 3 | 150000 | 138535 | 0.392906 ± 0.017844 | 0.359025 ± 0.019222 |
| PLux-M4 | 3 | 150000 | 137556 | 0.311607 ± 0.012651 | 0.262944 ± 0.012933 |
| PLux-M5 | 3 | 150000 | 137209 | 0.263953 ± 0.017110 | 0.201451 ± 0.017567 |

**Supplementary Table 21 | Summary by label.** Summary of replicate-level weight estimates by biological label, showing the number of replicates, total events used, the mean single-cell weight ⟨*W*⟩ with its standard error, and the bulk-estimated weight *W* derived from ratiometric fluorescence.

#### Per-file (per well)

| File | Label | Events total | Events used | ⟨*W*⟩ | Variance *W* (sample) | *W* bulk |
|---|---|---|---|---|---|---|
| **A1a.csv** | **PBS** | **58** | **58** | **0.847972** | **0.113442** | **0.896527** |
| **A2a.csv** | **Empty-cells** | **10000** | **8702** | **0.533061** | **0.206874** | **0.160309** |
| **C1a.csv** | **PBS** | **396** | **393** | **0.214608** | **0.124930** | **0.197205** |
| **C2a.csv** | **Empty-cells** | **50000** | **44357** | **0.529971** | **0.207276** | **0.147645** |
| E10a.csv | PLux-M2C | 50000 | 46516 | 0.494378 | 0.128730 | 0.487175 |
| **E1a.csv** | **PBS** | **97** | **91** | **0.124236** | **0.085221** | **0.073170** |
| **E2a.csv** | **Empty-cells** | **50000** | **44505** | **0.510989** | **0.209056** | **0.124767** |
| **E3a.csv** | **P2** | **50000** | **47295** | **0.034956** | **0.030857** | **0.000000** |
| **E4a.csv** | **P1** | **50000** | **47391** | **0.953803** | **0.018809** | **1.000000** |
| E5a.csv | PLux-M1A | 50000 | 46292 | 0.587769 | 0.127096 | 0.595095 |
| E6a.csv | PLux-M1B | 50000 | 46649 | 0.606325 | 0.125220 | 0.621064 |
| E7a.csv | PLux-M1C | 50000 | 46513 | 0.590340 | 0.125799 | 0.598621 |
| E8a.csv | PLux-M2A | 50000 | 45404 | 0.425513 | 0.124481 | 0.406306 |
| E9a.csv | PLux-M2B | 50000 | 45224 | 0.440451 | 0.121696 | 0.437659 |
| F1a.csv | PLux-M3A | 50000 | 45749 | 0.383621 | 0.120173 | 0.348506 |
| F2a.csv | PLux-M3B | 50000 | 46461 | 0.427392 | 0.127652 | 0.396308 |
| F3a.csv | PLux-M3C | 50000 | 46325 | 0.367706 | 0.119216 | 0.332261 |
| F4a.csv | PLux-M4A | 50000 | 46313 | 0.298667 | 0.106444 | 0.255753 |
| F5a.csv | PLux-M4B | 50000 | 45444 | 0.299246 | 0.108881 | 0.245023 |
| F6a.csv | PLux-M4C | 50000 | 45799 | 0.336907 | 0.118476 | 0.288058 |
| F7a.csv | PLux-M5A | 50000 | 46056 | 0.235707 | 0.094456 | 0.177699 |
| F8a.csv | PLux-M5B | 50000 | 45713 | 0.261348 | 0.107423 | 0.190905 |
| F9a.csv | PLux-M5C | 50000 | 45440 | 0.294805 | 0.114296 | 0.235748 |

**Supplementary Table 22 | Per-file single-well statistics.** Per-well statistics for each input file, including the number of events used, the mean single-cell weight ⟨*W*⟩, the unbiased sample variance of *W*, and the bulk-estimated weight derived from ratiometric fluorescence measurements.

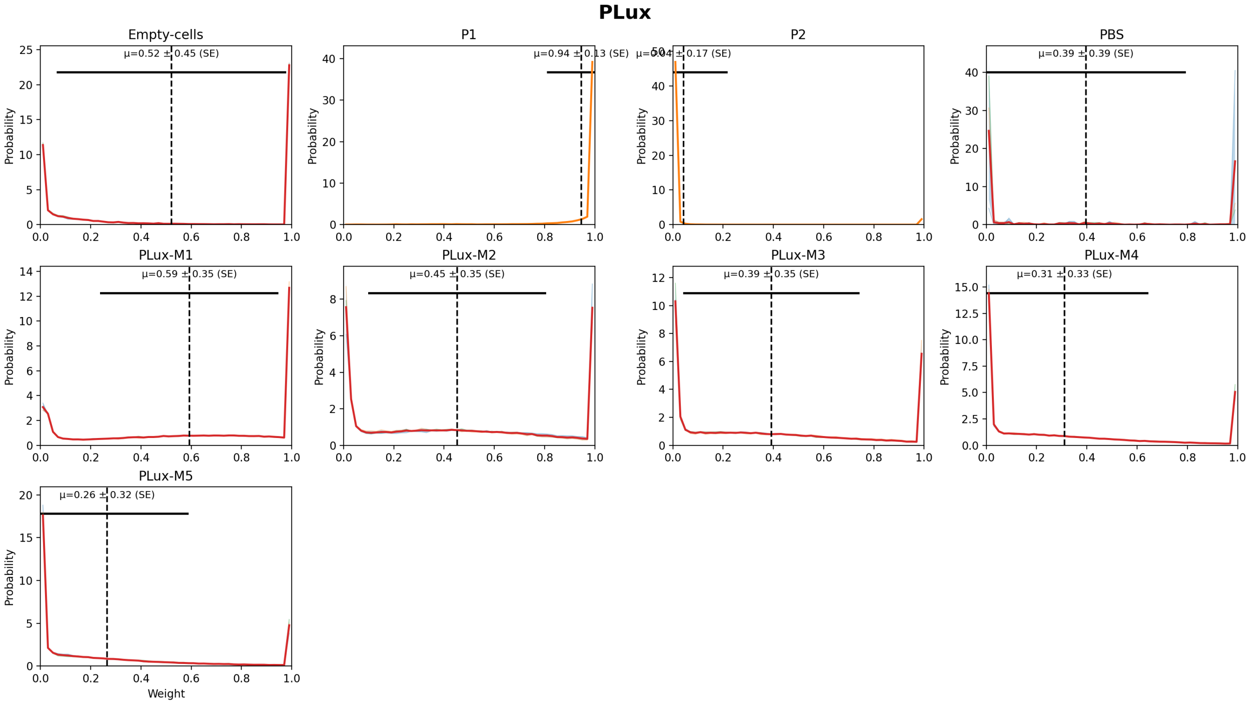

**Supplementary Fig. 16 | Weight distributions by label.** Weight distributions by label with replicate overlays; the shaded band shows the mean ± SD across replicates, the dashed line marks the mean single-cell weight, and the horizontal bar indicates ±SD of *W* across all gated single cells pooled across replicates for that label.

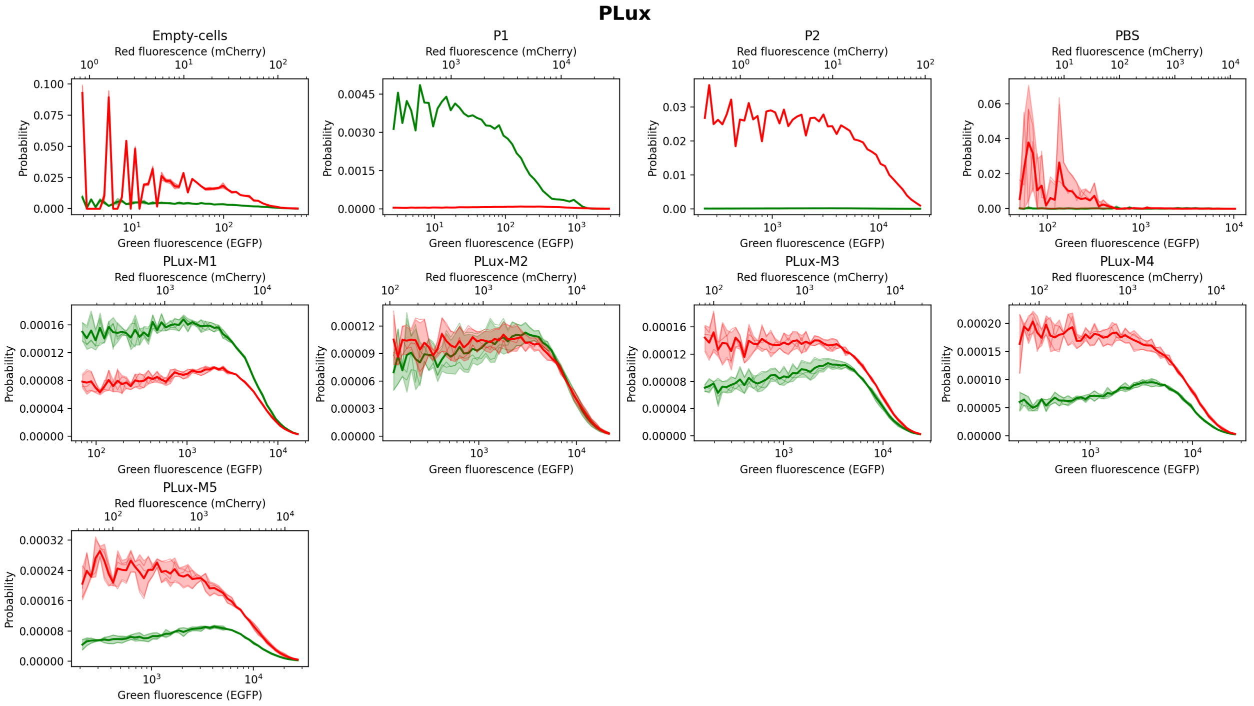

**Supplementary Fig. 17 | Fluorescence distributions by label.** Green and red fluorescence histograms by label on logarithmic *x*-axes with 10^n ticks; replicate histograms are overlaid and the shaded bands show mean ± SD across replicates.

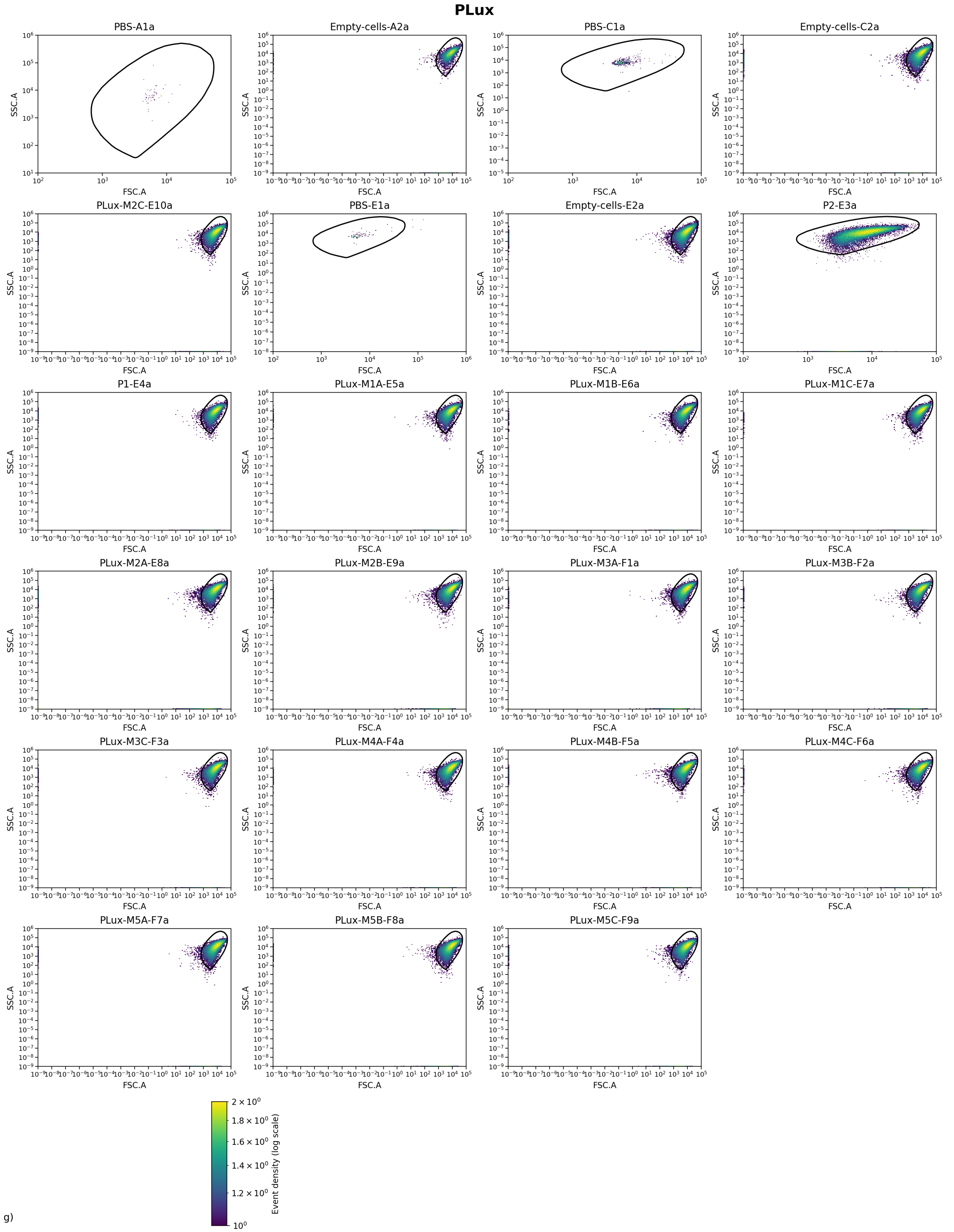

**Supplementary Fig. 18 | FSC–SSC distributions per well.** Two-dimensional FSC vs SSC (log10) density maps for each well before gating, with the non-parametric polygonal cell gate overlaid.

### PTtg

#### Experiment parameters and QC.

| Metric | Value |
|---|---|
| λ (from P1/P2 totals) | 2.71934 |
| β (from P1 totals) | 9.2262e-05 |
| γ | 0.0358164 |
| α spillover (R←G) | 0.00163878 |
| β spillover (G←R) | 0 |
| Background median (green) | 7424.06 |
| Background median (red) | 1.64 |
| Gate quantile (cell gate) | 0.99 |
| Gate d² threshold | NaN |
| Gate area (log10) | 7.79659 |
| Events (total) | 1090603 |
| Events (gated) | 985778 |
| Events used | 985778 |
| Gate acceptance (%) | 90.3883 |
| Used / gated (%) | 100 |
| Fluor green thr (quantile) | 0.999 |
| Fluor green thr (value) | 60728.7 |
| Fluor red thr (quantile) | 0.999 |
| Fluor red thr (value) | 329.002 |
| Median NEG green | 1617.66 |
| Median NEG red | 0 |
| Median POS green (P2) | 0 |
| Median POS red (P1) | 1095.52 |
| Stain index (green) | -0.0974082 |
| Stain index (red) | 16.7427 |
| Residual corr (negatives) | -0.0663909 |
| Dynamic range log10 (green) | 15.9728 |
| Dynamic range log10 (red) | 15.4466 |
| Replicate hist mean r | 0.949827 |
| Replicate hist mean RMSD | 0.900061 |

**Supplementary Table 23 | Experiment parameters and QC.** Experiment-level parameters and quality-control metrics for the FACS analysis, including background estimates, gating properties, event counts, stain indices and residual correlation after spectral compensation.

#### Summary by label (replicate means ± SE)

| Label | Replicates (n) | Events total (sum) | Events used (sum) | ⟨*W*⟩ ± SE | *W* bulk ± SE |
|---|---|---|---|---|---|
| **Empty-cells** | **4** | **200000** | **180187** | **0.853465 ± 0.146510** | **0.730873 ± 0.247929** |
| **P1** | **1** | **50000** | **45610** | **0.999931** | **1.000000** |
| **P2** | **1** | **50000** | **45130** | **0.819881** | **0.000000** |
| **PBS** | **4** | **11687** | **11559** | **0.943190 ± 0.044930** | **0.566117 ± 0.160413** |
| PTtg-M0 | 1 | 48916 | 45220 | 0.997273 | 0.978224 |
| PTtg-M1 | 3 | 130000 | 117465 | 0.986113 ± 0.001173 | 0.906127 ± 0.006457 |
| PTtg-M2 | 3 | 150000 | 135374 | 0.967187 ± 0.006211 | 0.772165 ± 0.047589 |
| PTtg-M3 | 3 | 150000 | 135712 | 0.950591 ± 0.001252 | 0.661509 ± 0.009347 |
| PTtg-M4 | 3 | 150000 | 134874 | 0.918355 ± 0.003173 | 0.482694 ± 0.011004 |
| PTtg-M5 | 3 | 150000 | 134647 | 0.868877 ± 0.001818 | 0.274024 ± 0.006703 |

**Supplementary Table 24 | Summary by label.** Summary of replicate-level weight estimates by biological label, showing the number of replicates, total events used, the mean single-cell weight ⟨*W*⟩ with its standard error, and the bulk-estimated weight *W* derived from ratiometric fluorescence.

#### Per-file (per well)

| File | Label | Events total | Events used | ⟨$W$⟩ | Variance $W$ (sample) | $W$ bulk |
|---|---|---|---|---|---|---|
| **A1b.csv** | **PBS** | **11299** | **11175** | **1.000000** | **0.000000** | **1.000251** |
| **A2b.csv** | **Empty-cells** | **50000** | **42200** | **0.413934** | **0.241748** | **-0.012382** |
| **C1b.csv** | **PBS** | **60** | **59** | **0.988834** | **0.007356** | **0.398523** |
| **C2b.csv** | **Empty-cells** | **50000** | **45830** | **0.999979** | **0.000010** | **0.987131** |
| **E1b.csv** | **PBS** | **68** | **67** | **0.974622** | **0.015731** | **0.601233** |
| **E2b.csv** | **Empty-cells** | **50000** | **46027** | **0.999978** | **0.000022** | **0.996140** |
| G10b.csv | PTtg-M2B | 50000 | 45251 | 0.973357 | 0.022016 | 0.819824 |
| G11b.csv | PTtg-M2C | 50000 | 44992 | 0.973439 | 0.021841 | 0.819685 |
| **G1b.csv** | **PBS** | **260** | **258** | **0.809304** | **0.150150** | **0.264462** |
| **G2b.csv** | **Empty-cells** | **50000** | **46130** | **0.999968** | **0.000019** | **0.952600** |
| **G3b.csv** | **P2** | **50000** | **45130** | **0.819881** | **0.146755** | **0.000000** |
| **G4b.csv** | **P1** | **50000** | **45610** | **0.999931** | **0.000057** | **1.000000** |
| G5b.csv | PTtg-M0 | 48916 | 45220 | 0.997273 | 0.002111 | 0.978224 |
| G6b.csv | PTtg-M1A | 30000 | 26850 | 0.987810 | 0.009983 | 0.917759 |
| G7b.csv | PTtg-M1B | 50000 | 45301 | 0.983862 | 0.013489 | 0.895454 |
| G8b.csv | PTtg-M1C | 50000 | 45314 | 0.986668 | 0.010892 | 0.905170 |
| G9b.csv | PTtg-M2A | 50000 | 45131 | 0.954765 | 0.037719 | 0.676987 |
| H1b.csv | PTtg-M3A | 50000 | 45380 | 0.948250 | 0.043037 | 0.650180 |
| H2b.csv | PTtg-M3B | 50000 | 45297 | 0.950995 | 0.041212 | 0.654296 |
| H3b.csv | PTtg-M3C | 50000 | 45035 | 0.952529 | 0.039871 | 0.680051 |
| H4b.csv | PTtg-M4A | 50000 | 44670 | 0.912069 | 0.072462 | 0.461164 |
| H5b.csv | PTtg-M4B | 50000 | 44992 | 0.922255 | 0.064468 | 0.497410 |
| H6b.csv | PTtg-M4C | 50000 | 45212 | 0.920740 | 0.066203 | 0.489509 |
| H7b.csv | PTtg-M5A | 50000 | 45082 | 0.871296 | 0.104327 | 0.281806 |
| H8b.csv | PTtg-M5B | 50000 | 44841 | 0.870018 | 0.104864 | 0.279587 |
| H9b.csv | PTtg-M5C | 50000 | 44724 | 0.865315 | 0.108604 | 0.260679 |

**Supplementary Table 25 | Per-file single-well statistics.** Per-well statistics for each input file, including the number of events used, the mean single-cell weight ⟨$W$⟩, the unbiased sample variance of $W$, and the bulk-estimated weight derived from ratiometric fluorescence measurements.

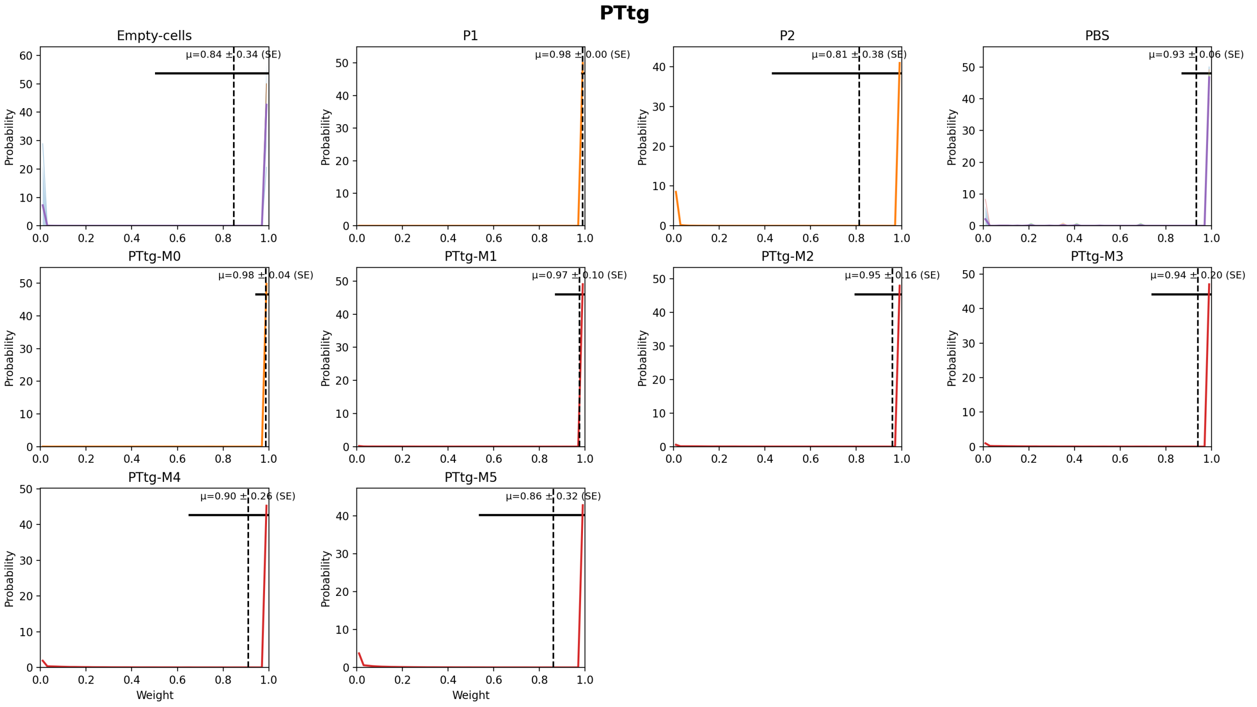

**Supplementary Fig. 19 | Weight distributions by label.** Weight distributions by label with replicate overlays; the shaded band shows the mean ± SD across replicates, the dashed line marks the mean single-cell weight, and the horizontal bar indicates ±SD of $W$ across all gated single cells pooled across replicates for that label.

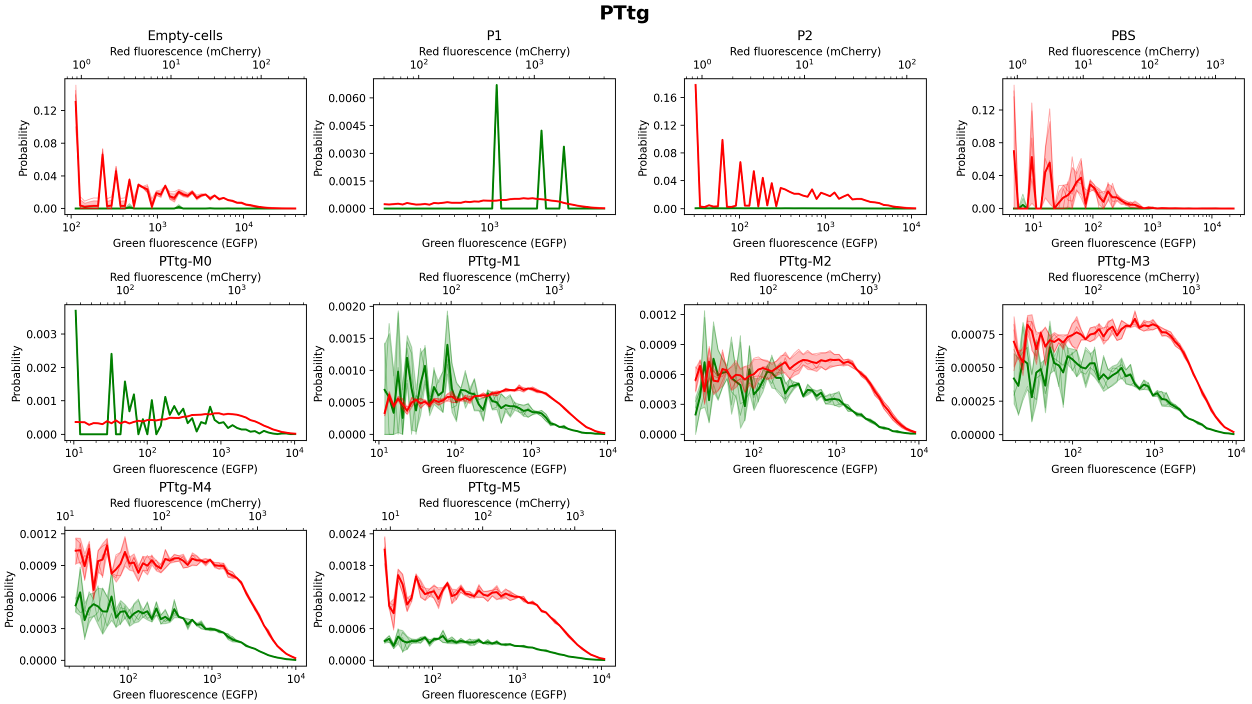

**Supplementary Fig. 20 | Fluorescence distributions by label.** Green and red fluorescence histograms by label on logarithmic *x*-axes with 10^n ticks; replicate histograms are overlaid and the shaded bands show mean ± SD across replicates.

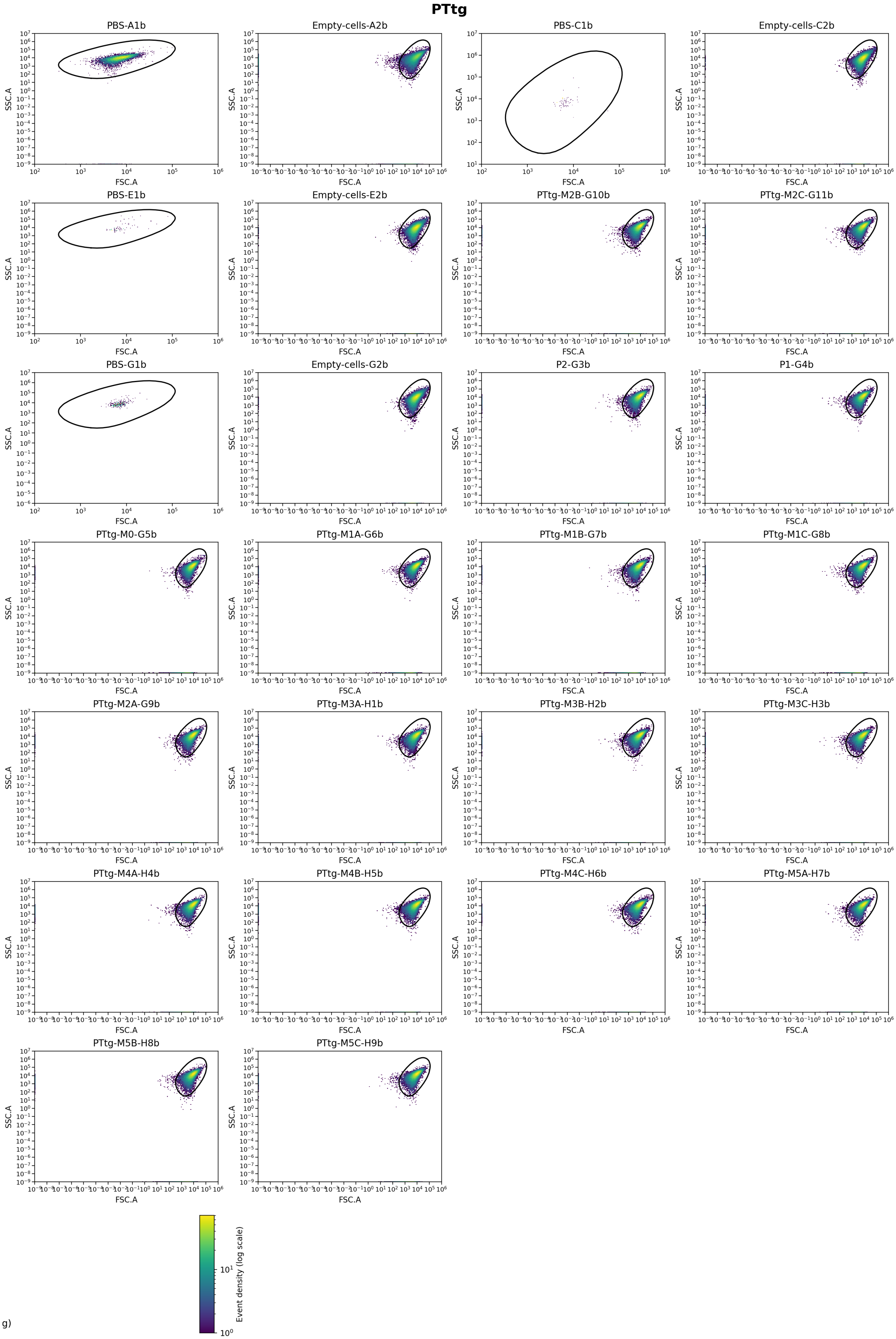
PTtg
PBS-A1b
Empty-cells-A2b
PBS-C1b
Empty-cells-C2b
PBS-E1b
Empty-cells-E2b
PTtg-M2B-G10b
PTtg-M2C-G11b
PBS-G1b
Empty-cells-G2b
P2-G3b
P1-G4b
PTtg-M0-G5b
PTtg-M1A-G6b
PTtg-M1B-G7b
PTtg-M1C-G8b
PTtg-M2A-G9b
PTtg-M3A-H1b
PTtg-M3B-H2b
PTtg-M3C-H3b
PTtg-M4A-H4b
PTtg-M4B-H5b
PTtg-M4C-H6b
PTtg-M5A-H7b
PTtg-M5B-H8b
PTtg-M5C-H9b
SSC.A
FSC.A
Event density (log scale)

g)

**Supplementary Fig. 21 | FSC–SSC distributions per well.** Two-dimensional FSC vs SSC (log10) density maps for each well before gating, with the non-parametric polygonal cell gate overlaid.

### PVan

#### Experiment parameters and QC.

| Metric | Value |
|---|---|
| λ (from P1/P2 totals) | 0.576001 |
| β (from P1 totals) | 0.0119499 |
| γ | 0.00298093 |
| α spillover (R←G) | 0 |
| β spillover (G←R) | 0.00040713 |
| Background median (green) | 34.96 |
| Background median (red) | 0.81 |
| Gate quantile (cell gate) | 0.99 |
| Gate $d^2$ threshold | NaN |
| Gate area (log10) | 3.33873 |
| Events (total) | 851071 |
| Events (gated) | 821324 |
| Events used | 821324 |
| Gate acceptance (%) | 96.5048 |
| Used / gated (%) | 100 |
| Fluor green thr (quantile) | 0.999 |
| Fluor green thr (value) | 290.814 |
| Fluor red thr (quantile) | 0.999 |
| Fluor red thr (value) | 132.312 |
| Median NEG green | 1.51604 |
| Median NEG red | 0.81 |
| Median POS green (P2) | 20099 |
| Median POS red (P1) | 9912.78 |
| Stain index (green) | 235.3 |
| Stain index (red) | 197.714 |
| Residual corr (negatives) | 0.296882 |
| Dynamic range log10 (green) | 16.585 |
| Dynamic range log10 (red) | 16.6225 |
| Replicate hist mean r | 0.901139 |
| Replicate hist mean RMSD | 0.756059 |

**Supplementary Table 26 | Experiment parameters and QC.** Experiment-level parameters and quality-control metrics for the FACS analysis, including background estimates, gating properties, event counts, stain indices and residual correlation after spectral compensation.

#### Summary by label (replicate means ± SE)

| Label | Replicates (n) | Events total (sum) | Events used (sum) | ⟨*W*⟩ ± SE | *W* bulk ± SE |
|---|---|---|---|---|---|
| **Empty-cells** | **2** | **100000** | **95216** | **0.633181 ± 0.003568** | **0.488506 ± 0.013325** |
| **P1** | **1** | **50000** | **48505** | **0.993650** | **1.000000** |
| **P2** | **1** | **50000** | **49793** | **0.063154** | **-0.000000** |
| **PBS** | **2** | **1071** | **277** | **0.572542 ± 0.056977** | **0.601075 ± 0.010514** |
| PVan-M0 | 1 | 50000 | 48658 | 0.804780 | 0.861575 |
| PVan-M1 | 3 | 150000 | 146054 | 0.617779 ± 0.023214 | 0.694349 ± 0.013108 |
| PVan-M2 | 3 | 150000 | 146110 | 0.682043 ± 0.000511 | 0.741958 ± 0.005031 |
| PVan-M3 | 3 | 150000 | 144423 | 0.710316 ± 0.015184 | 0.760299 ± 0.014569 |
| PVan-M4 | 3 | 150000 | 142288 | 0.636655 ± 0.098218 | 0.664580 ± 0.073616 |

**Supplementary Table 27 | Summary by label.** Summary of replicate-level weight estimates by biological label, showing the number of replicates, total events used, the mean single-cell weight ⟨*W*⟩ with its standard error, and the bulk-estimated weight *W* derived from ratiometric fluorescence.

#### Per-file (per well)

| File | Label | Events total | Events used | ⟨*W*⟩ | Variance *W* (sample) | *W* bulk |
|---|---|---|---|---|---|---|
| A11c.csv | PVan-M0 | 50000 | 48658 | 0.804780 | 0.065971 | 0.861575 |
| **A1c.csv** | **PBS** | **804** | **13** | **0.629519** | **0.044293** | **0.590561** |
| **A2c.csv** | **Empty-cells** | **50000** | **47653** | **0.636749** | **0.170959** | **0.475180** |
| **A7c.csv** | **P2** | **50000** | **49793** | **0.063154** | **0.049686** | **-0.000000** |
| **A8c.csv** | **P1** | **50000** | **48505** | **0.993650** | **0.004578** | **1.000000** |
| D10c.csv | PVan-M4A | 50000 | 48107 | 0.440573 | 0.191991 | 0.517463 |
| D11c.csv | PVan-M4B | 50000 | 46914 | 0.744890 | 0.111323 | 0.733115 |
| D12c.csv | PVan-M4C | 50000 | 47267 | 0.724503 | 0.113997 | 0.743163 |
| D1c.csv | PVan-M1A | 50000 | 49159 | 0.659511 | 0.122064 | 0.718220 |
| D2c.csv | PVan-M1B | 50000 | 48697 | 0.614534 | 0.132576 | 0.691800 |
| D3c.csv | PVan-M1C | 50000 | 48198 | 0.579293 | 0.135194 | 0.673028 |
| D4c.csv | PVan-M2A | 50000 | 48981 | 0.681568 | 0.118122 | 0.732088 |
| D5c.csv | PVan-M2B | 50000 | 48657 | 0.683065 | 0.117734 | 0.748587 |
| D6c.csv | PVan-M2C | 50000 | 48472 | 0.681496 | 0.116012 | 0.745198 |
| D7c.csv | PVan-M3A | 50000 | 48078 | 0.681007 | 0.117648 | 0.732292 |
| D8c.csv | PVan-M3B | 50000 | 47906 | 0.731857 | 0.103918 | 0.781265 |
| D9c.csv | PVan-M3C | 50000 | 48439 | 0.718083 | 0.106196 | 0.767339 |
| **E1c.csv** | **PBS** | **267** | **264** | **0.515566** | **0.117656** | **0.611589** |
| **E2c.csv** | **Empty-cells** | **50000** | **47563** | **0.629614** | **0.171235** | **0.501831** |

**Supplementary Table 28 | Per-file single-well statistics.** Per-well statistics for each input file, including the number of events used, the mean single-cell weight ⟨*W*⟩, the unbiased sample variance of *W*, and the bulk-estimated weight derived from ratiometric fluorescence measurements.

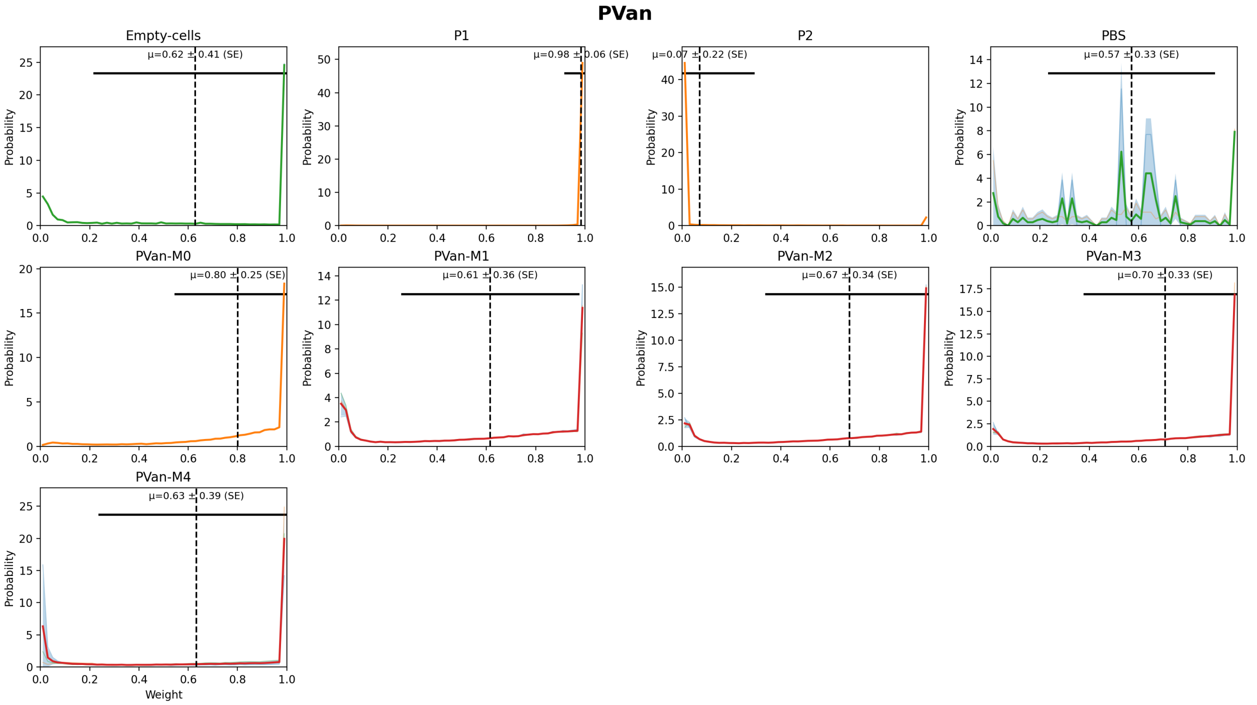

**Supplementary Fig. 22 | Weight distributions by label.** Weight distributions by label with replicate overlays; the shaded band shows the mean ± SD across replicates, the dashed line marks the mean single-cell weight, and the horizontal bar indicates ±SD of *W* across all gated single cells pooled across replicates for that label.

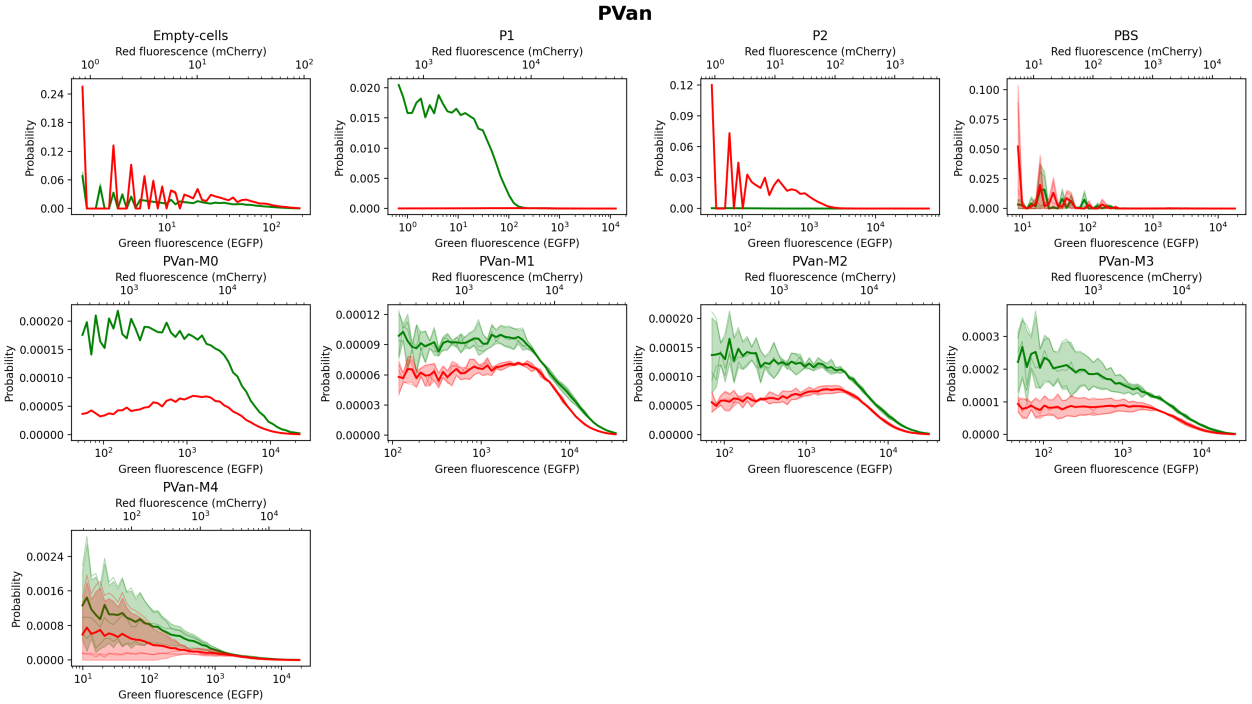

**Supplementary Fig. 23 | Fluorescence distributions by label.** Green and red fluorescence histograms by label on logarithmic *x*-axes with 10^n ticks; replicate histograms are overlaid and the shaded bands show mean ± SD across replicates.

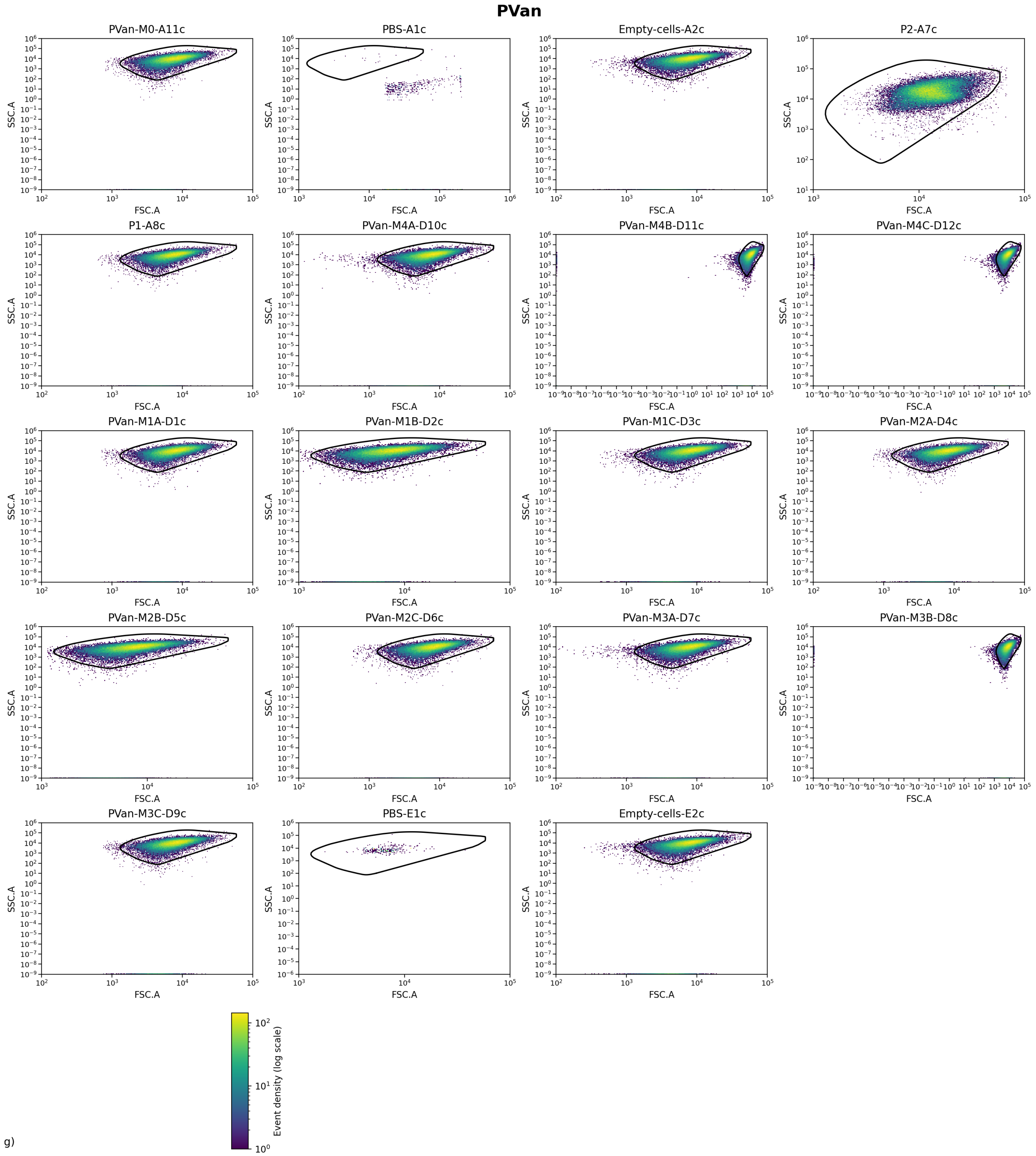

**Supplementary Fig. 24 | FSC–SSC distributions per well.** Two-dimensional FSC vs SSC (log10) density maps for each well before gating, with the non-parametric polygonal cell gate overlaid.

### PTac

#### Experiment parameters and QC.

| Metric | Value |
|---|---|
| λ (from P1/P2 totals) | 0.57022 |
| β (from P1 totals) | 0.0141224 |
| γ | 0.00258934 |
| α spillover (R←G) | 0.000163533 |
| β spillover (G←R) | 0.000999632 |
| Background median (green) | 34.96 |

| | |
|---|---|
| Background median (red) | 0.81 |
| Gate quantile (cell gate) | 0.99 |
| Gate d² threshold | NaN |
| Gate area (log10) | 3.68584 |
| Events (total) | 851071 |
| Events (gated) | 812372 |
| Events used | 812372 |
| Gate acceptance (%) | 95.4529 |
| Used / gated (%) | 100 |
| Fluor green thr (quantile) | 0.999 |
| Fluor green thr (value) | 291.984 |
| Fluor red thr (quantile) | 0.999 |
| Fluor red thr (value) | 132.958 |
| Median NEG green | 1.49976 |
| Median NEG red | 0.796267 |
| Median POS green (P2) | 13320.1 |
| Median POS red (P1) | 7789.77 |
| Stain index (green) | 151.076 |
| Stain index (red) | 141.92 |
| Residual corr (negatives) | 0.364751 |
| Dynamic range log10 (green) | 16.572 |
| Dynamic range log10 (red) | 16.5147 |
| Replicate hist mean r | 0.955162 |
| Replicate hist mean RMSD | 0.423691 |

**Supplementary Table 29 | Experiment parameters and QC.** Experiment-level parameters and quality-control metrics for the FACS analysis, including background estimates, gating properties, event counts, stain indices and residual correlation after spectral compensation.

#### Summary by label (replicate means ± SE)

| Label | Replicates (n) | Events total (sum) | Events used (sum) | ⟨*W*⟩ ± SE | *W* bulk ± SE |
|---|---|---|---|---|---|
| **Empty-cells** | **2** | **100000** | **95495** | **0.634504 ± 0.003628** | **0.491912 ± 0.013046** |
| **P1** | **1** | **50000** | **48359** | **0.991747** | **1.000000** |
| **P2** | **1** | **50000** | **49800** | **0.154987** | **0.000000** |
| **PBS** | **2** | **1071** | **277** | **0.574656 ± 0.057279** | **0.604116 ± 0.010495** |
| PTac-M0 | 1 | 50000 | 48538 | 0.762913 | 0.798576 |
| PTac-M1 | 3 | 150000 | 145891 | 0.738226 ± 0.021186 | 0.772370 ± 0.015070 |
| PTac-M2 | 3 | 150000 | 145030 | 0.723371 ± 0.005354 | 0.742482 ± 0.004388 |
| PTac-M3 | 3 | 150000 | 139829 | 0.653150 ± 0.020125 | 0.730687 ± 0.003337 |
| PTac-M4 | 3 | 150000 | 139153 | 0.175149 ± 0.002484 | 0.272632 ± 0.034821 |

**Supplementary Table 30 | Summary by label.** Summary of replicate-level weight estimates by biological label, showing the number of replicates, total events used, the mean single-cell weight ⟨*W*⟩ with its standard error, and the bulk-estimated weight *W* derived from ratiometric fluorescence.

#### Per-file (per well)

| File | Label | Events total | Events used | ⟨*W*⟩ | Variance *W* (sample) | *W* bulk |
|---|---|---|---|---|---|---|
| A10c.csv | PTac-M0 | 50000 | 48538 | 0.762913 | 0.080522 | 0.798576 |
| **A1c.csv** | **PBS** | **804** | **13** | **0.631935** | **0.043958** | **0.593621** |
| **A2c.csv** | **Empty-cells** | **50000** | **47808** | **0.638133** | **0.170450** | **0.478866** |
| **A5c.csv** | **P2** | **50000** | **49800** | **0.154987** | **0.122716** | **0.000000** |
| **A6c.csv** | **P1** | **50000** | **48359** | **0.991747** | **0.005319** | **1.000000** |
| C10c.csv | PTac-M4A | 50000 | 46291 | 0.179803 | 0.101354 | 0.311458 |

| | | | | | | |
|---|---|---|---|---|---|---|
| C11c.csv | PTac-M4B | 50000 | 45801 | 0.171315 | 0.090668 | 0.203150 |
| C12c.csv | PTac-M4C | 50000 | 47061 | 0.174330 | 0.094703 | 0.303290 |
| C1c.csv | PTac-M1A | 50000 | 48522 | 0.721503 | 0.098002 | 0.764686 |
| C2c.csv | PTac-M1B | 50000 | 48885 | 0.780304 | 0.085468 | 0.801451 |
| C3c.csv | PTac-M1C | 50000 | 48484 | 0.712871 | 0.101035 | 0.750974 |
| C4c.csv | PTac-M2A | 50000 | 48230 | 0.712667 | 0.106631 | 0.741523 |
| C5c.csv | PTac-M2B | 50000 | 48440 | 0.728452 | 0.099441 | 0.735406 |
| C6c.csv | PTac-M2C | 50000 | 48360 | 0.728994 | 0.097714 | 0.750516 |
| C7c.csv | PTac-M3A | 50000 | 46679 | 0.682554 | 0.095171 | 0.728366 |
| C8c.csv | PTac-M3B | 50000 | 46402 | 0.614646 | 0.099932 | 0.737267 |
| C9c.csv | PTac-M3C | 50000 | 46748 | 0.662251 | 0.095659 | 0.726428 |
| **E1c.csv** | **PBS** | **267** | **264** | **0.517377** | **0.117625** | **0.614611** |
| **E2c.csv** | **Empty-cells** | **50000** | **47687** | **0.630876** | **0.170712** | **0.504958** |

**Supplementary Table 31 | Per-file single-well statistics.** Per-well statistics for each input file, including the number of events used, the mean single-cell weight ⟨*W*⟩, the unbiased sample variance of *W*, and the bulk-estimated weight derived from ratiometric fluorescence measurements.

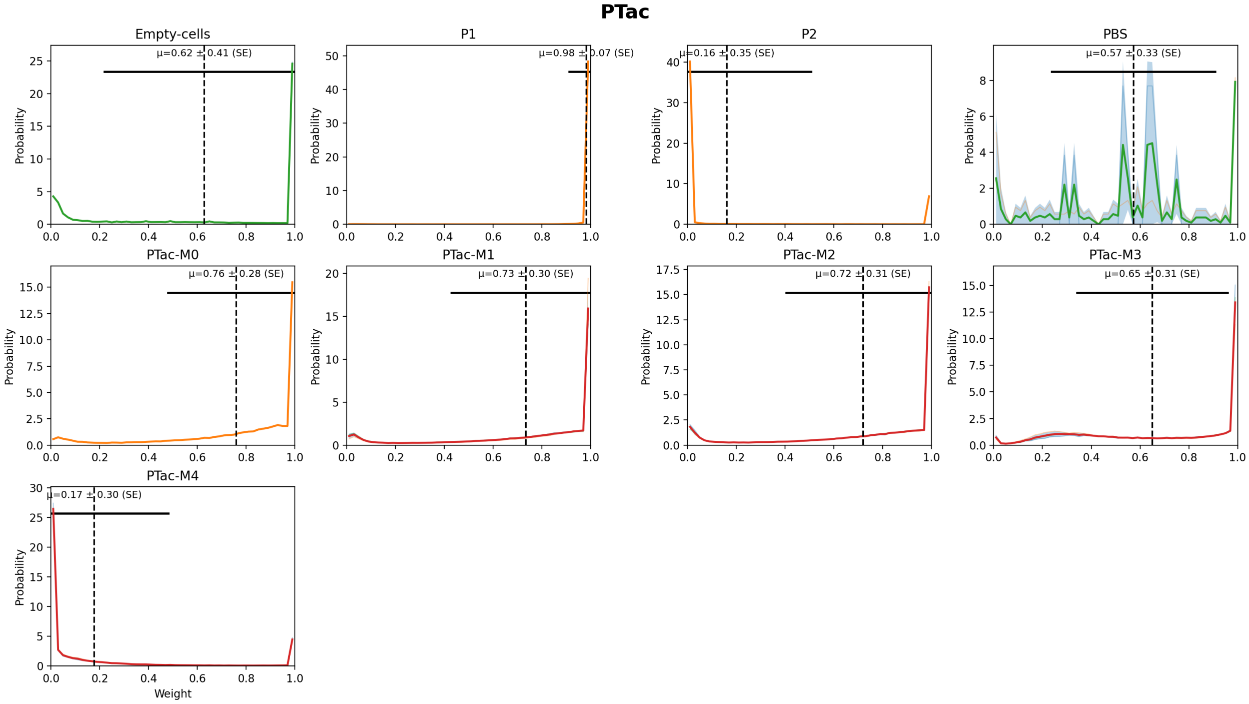

**Supplementary Fig. 25 | Weight distributions by label.** Weight distributions by label with replicate overlays; the shaded band shows the mean ± SD across replicates, the dashed line marks the mean single-cell weight, and the horizontal bar indicates ±SD of *W* across all gated single cells pooled across replicates for that label.

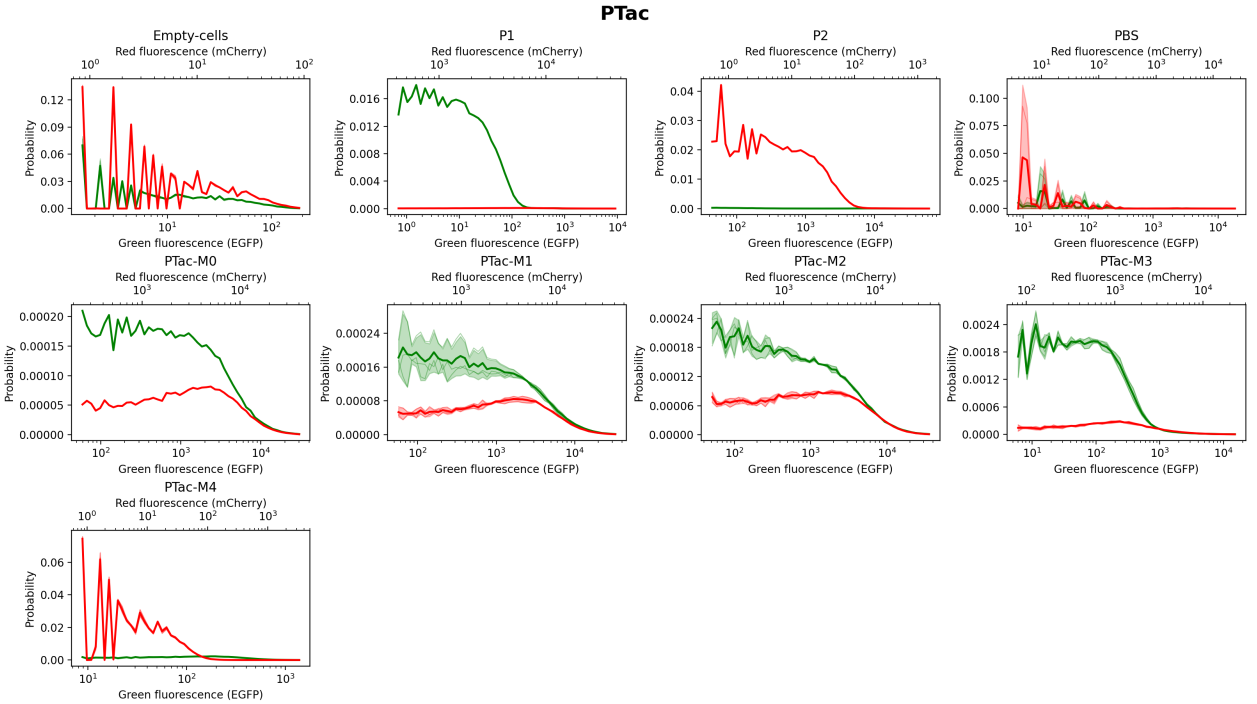

**Supplementary Fig. 26 | Fluorescence distributions by label.** Green and red fluorescence histograms by label on logarithmic *x*-axes with 10^n ticks; replicate histograms are overlaid and the shaded bands show mean ± SD across replicates.

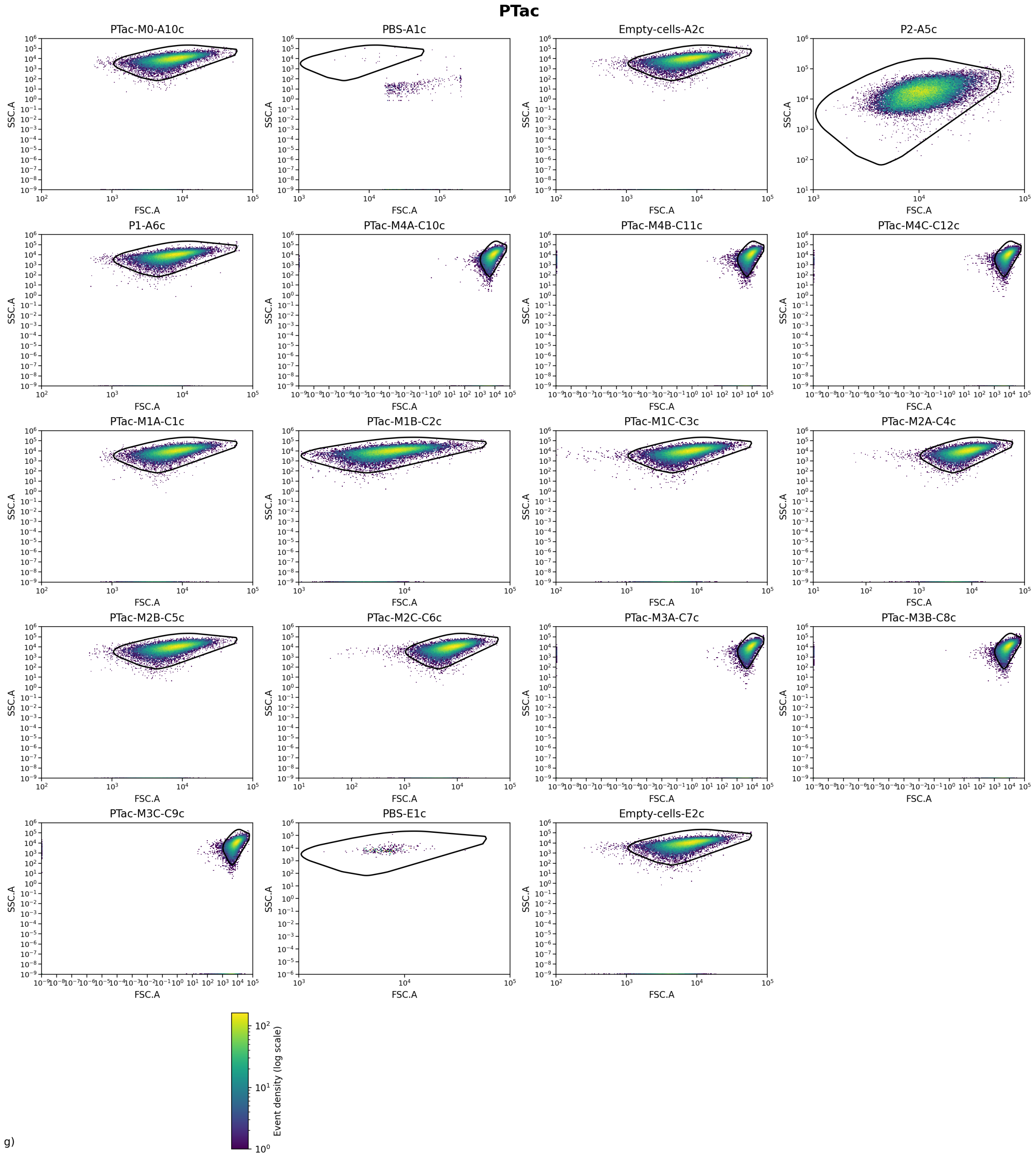

**Supplementary Fig. 27 | FSC–SSC distributions per well.** Two-dimensional FSC vs SSC (log10) density maps for each well before gating, with the non-parametric polygonal cell gate overlaid.

### PVan-Ttg

#### Experiment parameters and QC.

| Metric | Value |
|---|---|
| λ (from P1/P2 totals) | 1.32057 |
| β (from P1 totals) | 0.00318843 |
| γ | 0.00196165 |
| α spillover (R←G) | 0.000123995 |
| β spillover (G←R) | 0 |
| Background median (green) | 7424.06 |

| | |
|---|---|
| Background median (red) | 1.64 |
| Gate quantile (cell gate) | 0.99 |
| Gate d² threshold | NaN |
| Gate area (log10) | 8.07009 |
| Events (total) | 961687 |
| Events (gated) | 866567 |
| Events used | 866567 |
| Gate acceptance (%) | 90.109 |
| Used / gated (%) | 100 |
| Fluor green thr (quantile) | 0.999 |
| Fluor green thr (value) | 60683.5 |
| Fluor red thr (quantile) | 0.999 |
| Fluor red thr (value) | 337.012 |
| Median NEG green | 1609.3 |
| Median NEG red | 0 |
| Median POS green (P2) | 3122.46 |
| Median POS red (P1) | 8237.72 |
| Stain index (green) | 0.0911169 |
| Stain index (red) | 118.47 |
| Residual corr (negatives) | 0.0680498 |
| Dynamic range log10 (green) | 16.3924 |
| Dynamic range log10 (red) | 16.4619 |
| Replicate hist mean r | 0.945757 |
| Replicate hist mean RMSD | 1.03565 |

**Supplementary Table 32 | Experiment parameters and QC.** Experiment-level parameters and quality-control metrics for the FACS analysis, including background estimates, gating properties, event counts, stain indices and residual correlation after spectral compensation.

#### Summary by label (replicate means ± SE)

| Label | Replicates (n) | Events total (sum) | Events used (sum) | ⟨*W*⟩ ± SE | *W* bulk ± SE |
|---|---|---|---|---|---|
| **Empty-cells** | **4** | **200000** | **180359** | **0.854170 ± 0.145811** | **0.745324 ± 0.248180** |
| **P1** | **1** | **50000** | **46405** | **0.996254** | **1.000000** |
| **P2** | **1** | **50000** | **46474** | **0.369086** | **0.000000** |
| **PBS** | **4** | **11687** | **11561** | **0.946972 ± 0.044428** | **0.695759 ± 0.122485** |
| PVan-Ttg-M0 | 1 | 50000 | 45713 | 0.616092 | 0.424395 |
| PVan-Ttg-M1 | 3 | 150000 | 137051 | 0.925882 ± 0.022241 | 0.883545 ± 0.007395 |
| PVan-Ttg-M2 | 3 | 150000 | 134533 | 0.994419 ± 0.003009 | 0.978937 ± 0.007098 |
| PVan-Ttg-M3 | 3 | 150000 | 131229 | 0.999833 ± 0.000020 | 0.998679 ± 0.000394 |
| PVan-Ttg-M4 | 3 | 150000 | 133242 | 0.999987 ± 0.000007 | 0.992252 ± 0.007600 |

**Supplementary Table 33 | Summary by label.** Summary of replicate-level weight estimates by biological label, showing the number of replicates, total events used, the mean single-cell weight ⟨*W*⟩ with its standard error, and the bulk-estimated weight *W* derived from ratiometric fluorescence.

#### Per-file (per well)

| File | Label | Events total | Events used | ⟨*W*⟩ | Variance *W* (sample) | *W* bulk |
|---|---|---|---|---|---|---|
| **A1b.csv** | **PBS** | **11299** | **11176** | **1.000000** | **0.000000** | **1.004228** |
| **A2b.csv** | **Empty-cells** | **50000** | **42263** | **0.416739** | **0.240350** | **0.000919** |
| C10b.csv | PVan-Ttg-M2B | 50000 | 43810 | 0.994514 | 0.004591 | 0.982549 |
| C11b.csv | PVan-Ttg-M2C | 50000 | 46931 | 0.989160 | 0.009079 | 0.965241 |

| | | | | | | |
|---|---|---|---|---|---|---|
| **C1b.csv** | **PBS** | **60** | **60** | **0.992053** | **0.003790** | **0.583855** |
| **C2b.csv** | **Empty-cells** | **50000** | **45860** | **0.999987** | **0.000005** | **0.997799** |
| **C3b.csv** | **P2** | **50000** | **46474** | **0.369086** | **0.230715** | **0.000000** |
| **C4b.csv** | **P1** | **50000** | **46405** | **0.996254** | **0.003100** | **1.000000** |
| C5b.csv | PVan-Ttg-M0 | 50000 | 45713 | 0.616092 | 0.177616 | 0.424395 |
| C6b.csv | PVan-Ttg-M1A | 50000 | 44494 | 0.970280 | 0.024313 | 0.896345 |
| C7b.csv | PVan-Ttg-M1B | 50000 | 46193 | 0.906043 | 0.067798 | 0.870728 |
| C8b.csv | PVan-Ttg-M1C | 50000 | 46364 | 0.901322 | 0.069781 | 0.883563 |
| C9b.csv | PVan-Ttg-M2A | 50000 | 43792 | 0.999582 | 0.000386 | 0.989020 |
| D1b.csv | PVan-Ttg-M3A | 50000 | 43516 | 0.999873 | 0.000121 | 0.999226 |
| D2b.csv | PVan-Ttg-M3B | 50000 | 43723 | 0.999816 | 0.000165 | 0.997915 |
| D3b.csv | PVan-Ttg-M3C | 50000 | 43990 | 0.999809 | 0.000161 | 0.998896 |
| D4b.csv | PVan-Ttg-M4A | 50000 | 44339 | 1.000000 | 0.000000 | 1.004228 |
| D5b.csv | PVan-Ttg-M4B | 50000 | 44539 | 0.999984 | 0.000007 | 0.978157 |
| D6b.csv | PVan-Ttg-M4C | 50000 | 44364 | 0.999977 | 0.000023 | 0.994372 |
| **E1b.csv** | **PBS** | **68** | **67** | **0.981672** | **0.008925** | **0.760317** |
| **E2b.csv** | **Empty-cells** | **50000** | **46065** | **0.999979** | **0.000021** | **1.002216** |
| **G1b.csv** | **PBS** | **260** | **258** | **0.814163** | **0.145089** | **0.434633** |
| **G2b.csv** | **Empty-cells** | **50000** | **46171** | **0.999978** | **0.000010** | **0.980364** |

**Supplementary Table 34 | Per-file single-well statistics.** Per-well statistics for each input file, including the number of events used, the mean single-cell weight ⟨*W*⟩, the unbiased sample variance of *W*, and the bulk-estimated weight derived from ratiometric fluorescence measurements.

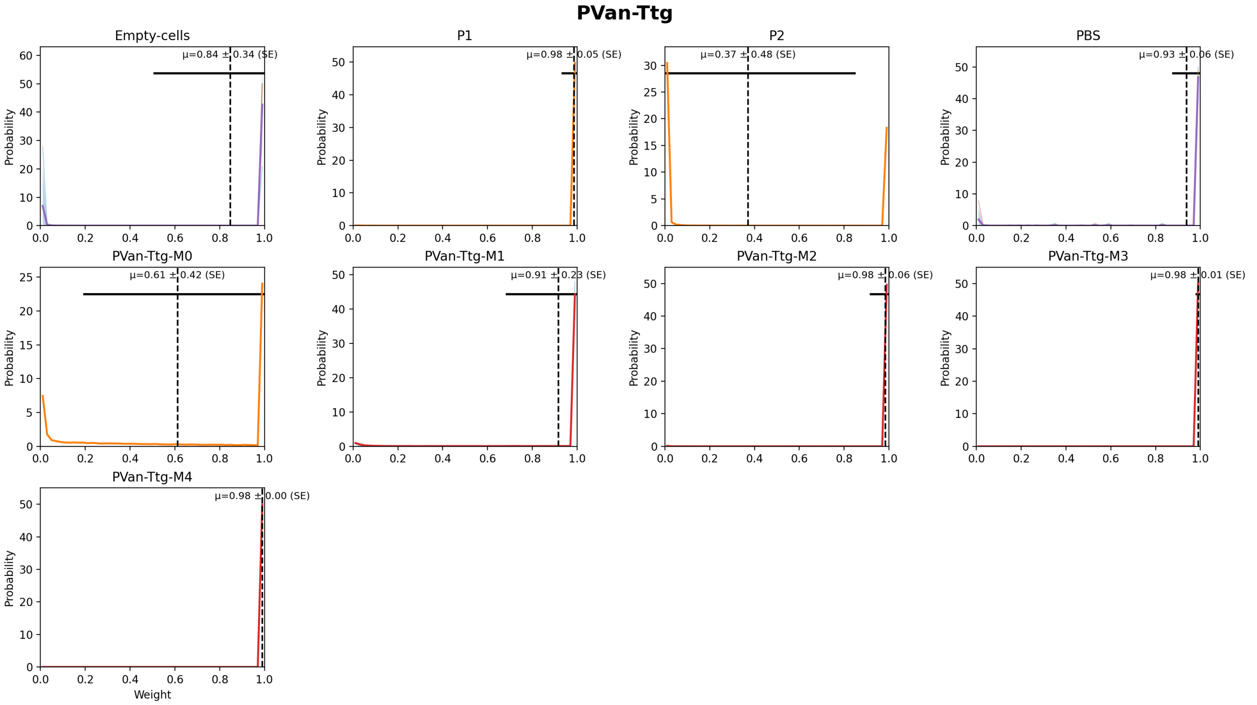

**Supplementary Fig. 28 | Weight distributions by label.** Weight distributions by label with replicate overlays; the shaded band shows the mean ± SD across replicates, the dashed line marks the mean single-cell weight, and the horizontal bar indicates ±SD of *W* across all gated single cells pooled across replicates for that label.

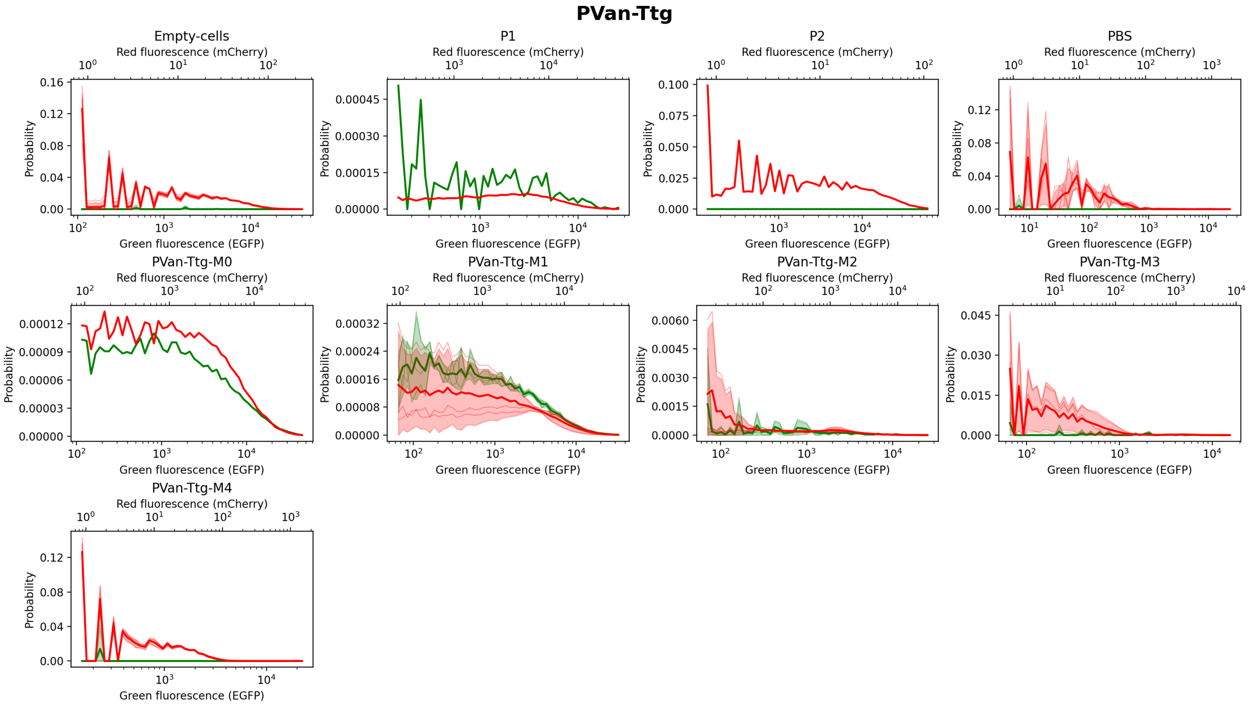

**Supplementary Fig. 29 | Fluorescence distributions by label.** Green and red fluorescence histograms by label on logarithmic *x*-axes with 10^n ticks; replicate histograms are overlaid and the shaded bands show mean ± SD across replicates.

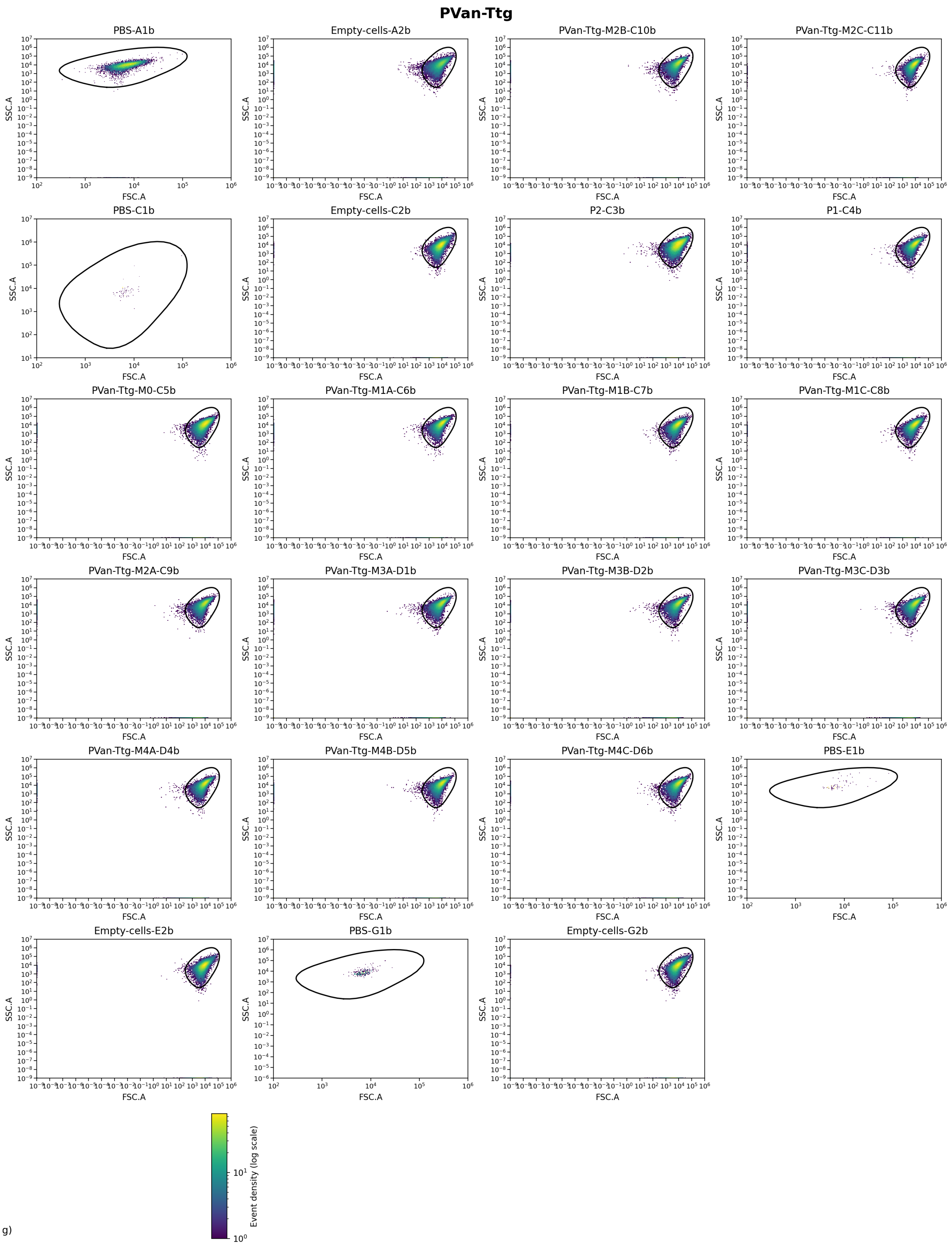

g)

**Supplementary Fig. 30 | FSC–SSC distributions per well.** Two-dimensional FSC vs SSC (log10) density maps for each well before gating, with the non-parametric polygonal cell gate overlaid.

### PTac-Van

#### Experiment parameters and QC.

| Metric | Value |
|---|---|
| λ (from P1/P2 totals) | 0.48231 |
| β (from P1 totals) | 0.0371208 |
| γ | 0.000734018 |
| α spillover (R←G) | 0.000159865 |
| β spillover (G←R) | 0.00132653 |
| Background median (green) | 34.96 |
| Background median (red) | 0.81 |
| Gate quantile (cell gate) | 0.99 |
| Gate $d^2$ threshold | NaN |
| Gate area (log10) | 4.7312 |
| Events (total) | 851071 |
| Events (gated) | 802799 |
| Events used | 802799 |
| Gate acceptance (%) | 94.3281 |
| Used / gated (%) | 100 |
| Fluor green thr (quantile) | 0.999 |
| Fluor green thr (value) | 307.005 |
| Fluor red thr (quantile) | 0.999 |
| Fluor red thr (value) | 138.516 |
| Median NEG green | 1.48239 |
| Median NEG red | 0.796878 |
| Median POS green (P2) | 13125.2 |
| Median POS red (P1) | 6897.91 |
| Stain index (green) | 129.564 |
| Stain index (red) | 95.2041 |
| Residual corr (negatives) | 0.55875 |
| Dynamic range log10 (green) | 16.4916 |
| Dynamic range log10 (red) | 16.5694 |
| Replicate hist mean r | 0.954079 |
| Replicate hist mean RMSD | 0.590762 |

**Supplementary Table 35 | Experiment parameters and QC.** Experiment-level parameters and quality-control metrics for the FACS analysis, including background estimates, gating properties, event counts, stain indices and residual correlation after spectral compensation.

#### Summary by label (replicate means ± SE)

| Label | Replicates (n) | Events total (sum) | Events used (sum) | ⟨*W*⟩ ± SE | *W* bulk ± SE |
|---|---|---|---|---|---|
| **Empty-cells** | **2** | **100000** | **95791** | **0.647434 ± 0.003426** | **0.538710 ± 0.012696** |
| **P1** | **1** | **50000** | **47470** | **0.966391** | **1.000000** |
| **P2** | **1** | **50000** | **49904** | **0.027120** | **0.000000** |
| **PBS** | **2** | **1071** | **278** | **0.612212 ± 0.068860** | **0.649586 ± 0.009148** |
| PTac-Van-M0 | 1 | 50000 | 45789 | 0.836247 | 0.888993 |
| PTac-Van-M1 | 3 | 150000 | 140365 | 0.923528 ± 0.005227 | 0.956049 ± 0.005860 |
| PTac-Van-M2 | 3 | 150000 | 140226 | 0.943309 ± 0.005406 | 0.972199 ± 0.004545 |
| PTac-Van-M3 | 3 | 150000 | 140394 | 0.955762 ± 0.006192 | 0.981837 ± 0.004257 |
| PTac-Van-M4 | 3 | 150000 | 142582 | 0.845808 ± 0.049099 | 0.884767 ± 0.041488 |

**Supplementary Table 36 | Summary by label.** Summary of replicate-level weight estimates by biological label, showing the number of replicates, total events used, the mean single-cell weight ⟨*W*⟩ with its standard error, and the bulk-estimated weight *W* derived from ratiometric fluorescence.

#### Per-file (per well)

| File | Label | Events total | Events used | ⟨$W$⟩ | Variance $W$ (sample) | $W$ bulk |
|---|---|---|---|---|---|---|
| **A1c.csv** | **PBS** | **804** | **14** | **0.681072** | **0.038464** | **0.640438** |
| **A2c.csv** | **Empty-cells** | **50000** | **47966** | **0.650859** | **0.164934** | **0.526015** |
| **E1c.csv** | **PBS** | **267** | **264** | **0.543351** | **0.117339** | **0.658735** |
| **E2c.csv** | **Empty-cells** | **50000** | **47825** | **0.644008** | **0.165204** | **0.551406** |
| **E3c.csv** | **P2** | **50000** | **49904** | **0.027120** | **0.024536** | **0.000000** |
| **E4c.csv** | **P1** | **50000** | **47470** | **0.966391** | **0.014083** | **1.000000** |
| E7c.csv | PTac-Van-M0 | 50000 | 45789 | 0.836247 | 0.094490 | 0.888993 |
| F10c.csv | PTac-Van-M4A | 50000 | 47057 | 0.758914 | 0.121569 | 0.814318 |
| F11c.csv | PTac-Van-M4B | 50000 | 47924 | 0.849641 | 0.095808 | 0.882024 |
| F12c.csv | PTac-Van-M4C | 50000 | 47601 | 0.928868 | 0.050027 | 0.957958 |
| F1c.csv | PTac-Van-M1A | 50000 | 46916 | 0.917298 | 0.052944 | 0.950350 |
| F2c.csv | PTac-Van-M1B | 50000 | 46975 | 0.919373 | 0.052660 | 0.950031 |
| F3c.csv | PTac-Van-M1C | 50000 | 46474 | 0.933914 | 0.043237 | 0.967768 |
| F4c.csv | PTac-Van-M2A | 50000 | 46899 | 0.933698 | 0.048439 | 0.964128 |
| F5c.csv | PTac-Van-M2B | 50000 | 46712 | 0.952404 | 0.034744 | 0.979857 |
| F6c.csv | PTac-Van-M2C | 50000 | 46615 | 0.943825 | 0.041486 | 0.972613 |
| F7c.csv | PTac-Van-M3A | 50000 | 46638 | 0.943410 | 0.039730 | 0.973331 |
| F8c.csv | PTac-Van-M3B | 50000 | 46894 | 0.962719 | 0.026760 | 0.986430 |
| F9c.csv | PTac-Van-M3C | 50000 | 46862 | 0.961157 | 0.027983 | 0.985748 |

**Supplementary Table 37 | Per-file single-well statistics.** Per-well statistics for each input file, including the number of events used, the mean single-cell weight ⟨$W$⟩, the unbiased sample variance of $W$, and the bulk-estimated weight derived from ratiometric fluorescence measurements.

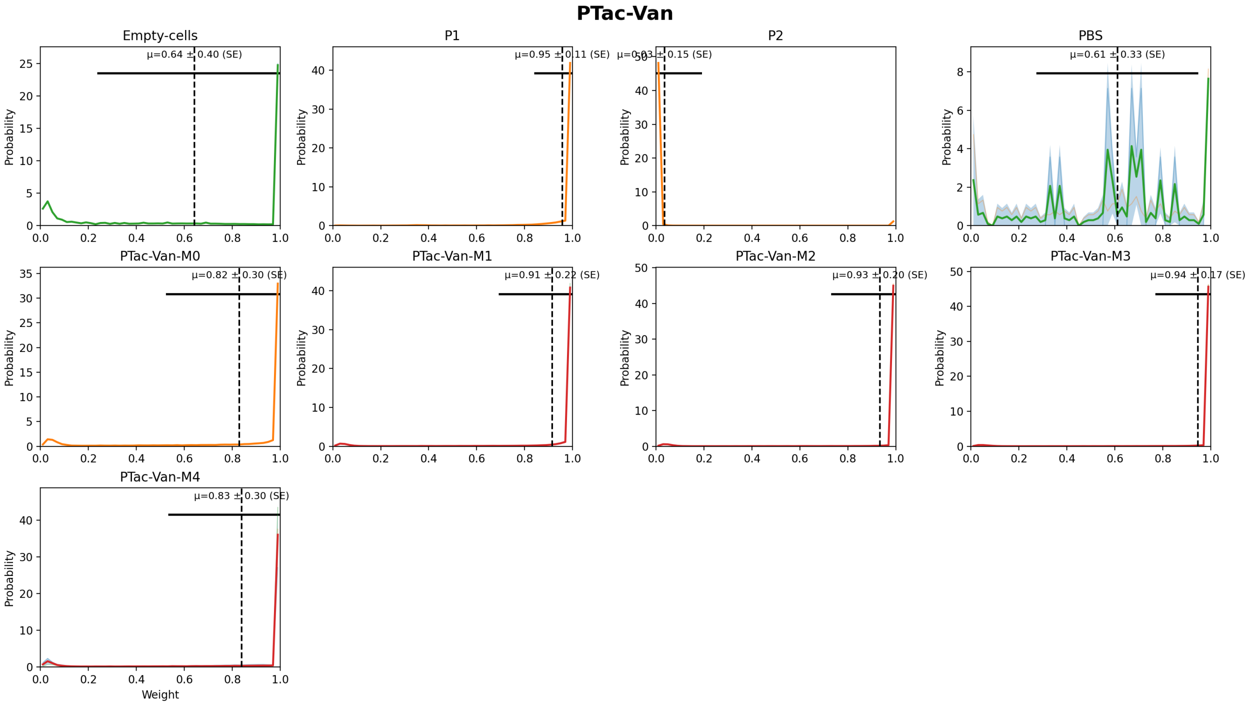

**Supplementary Fig. 31 | Weight distributions by label.** Weight distributions by label with replicate overlays; the shaded band shows the mean ± SD across replicates, the dashed line marks the mean single-cell weight, and the horizontal bar indicates ±SD of *W* across all gated single cells pooled across replicates for that label.

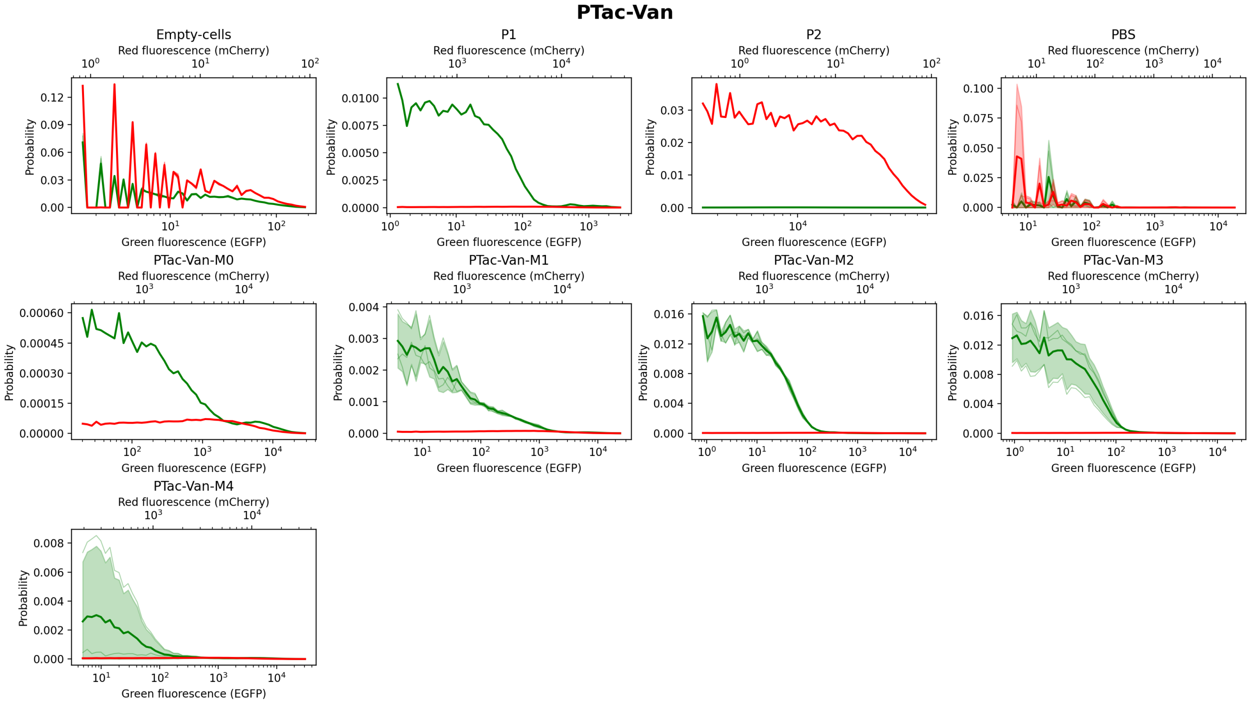

**Supplementary Fig. 32 | Fluorescence distributions by label.** Green and red fluorescence histograms by label on logarithmic *x*-axes with 10^n ticks; replicate histograms are overlaid and the shaded bands show mean ± SD across replicates.

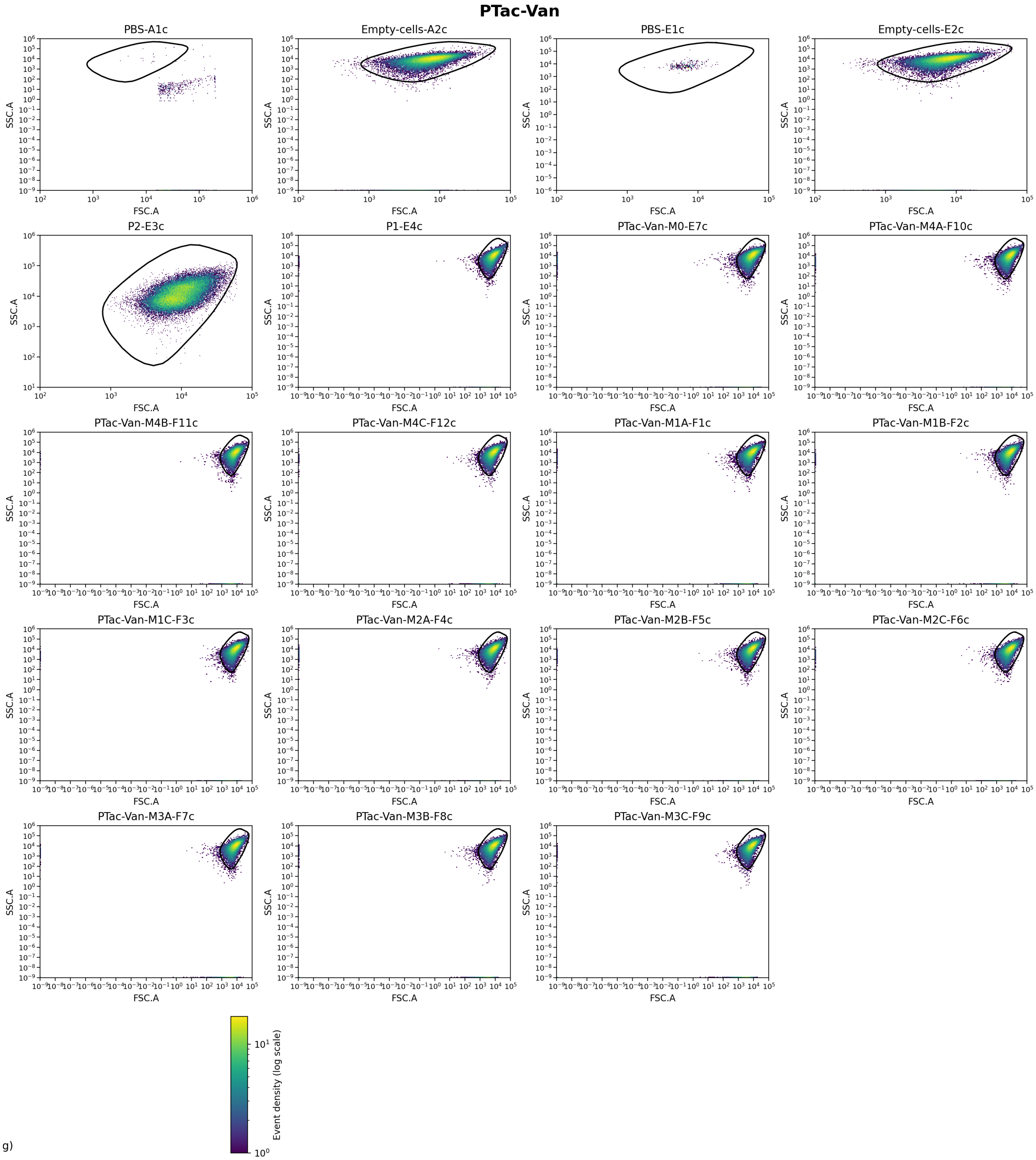

**Supplementary Fig. 33 | FSC–SSC distributions per well.** Two-dimensional FSC vs SSC (log10) density maps for each well before gating, with the non-parametric polygonal cell gate overlaid.

#### Fitting of measured single-cell weight distribution against theoretical estimates

Supplementary Fig. 34 shows for each promoter, the earliest M-stage single-cell weight distribution $p_{data}(W)$ is fitted with a three-component mixture on [0,1], $p_{model}(W) = q_0 \cdot f_0(W) + q_1 \cdot f_1(W) + q_c \cdot f_c(W)$, where $q_0$, $q_1$ and q_c are the fractions of cells in the P2-like ($W \approx 0$), P1-like ($W \approx 1$) and interior sub-populations, respectively, with $q_0 + q_1 + q\_c = 1$. This corresponds to a three-peak mixture model for initial weight distributions. The functions $f_0(W)$ and $f_1(W)$ are empirical templates obtained from the P2-only (EGFP-only) and P1-only (mCherry-only) control wells, normalised so that $\int_0^1 f_0(W)\, dW = \int_0^1 f_1(W)\, dW = 1$. The interior peak $f_c(W)$ is a Beta distribution on [0,1] with mean $W_0$ and shape parameter $K > 2$, defined by $f_c(W) = 1 / B(\alpha, \beta) \cdot W^{\alpha-1} (1 - W)^{\beta-1}$ for $0 \leq W \leq 1$, and $f_c(W) = 0$ otherwise. Here $B(\alpha,\beta)$ is the Beta function,

$$B(\alpha,\beta) = \int_0^1 t^{\alpha-1}(1-t)^{\beta-1}\, dt = \Gamma(\alpha)\Gamma(\beta)/\Gamma(\alpha+\beta),$$

and we parameterise $\alpha = W_0 \cdot K$ and $\beta = (1-W_0) \cdot K$ so that the mean of the interior peak is $E[W \mid \text{interior}] = W_0$. The overall model mean weight is therefore $\langle W \rangle_{model} = q_1 + q\_c \cdot W_0$, because the P2-like component is concentrated near $W \approx 0$. Each panel in Supplementary Fig. 34 shows $p_{data}(W)$ as a solid line and the corresponding $p_{model}(W)$ as a dashed line, together with the fitted values of $q_0$ (labelled "P2-only"), $q_1$ (labelled "P1-only"), q_c (labelled "Central"), the interior-peak mean $W_0$, the shape parameter $K$, and the Pearson correlation r between $p_{data}(W)$ and $p_{model}(W)$.

For each promoter, we described the earliest M-stage single-cell weight distribution as a mixture of three sub-populations derived from the control wells and from an interior peak (Supplementary Fig. 34)

$p(W) = q_0\, f_0(W) + q_1\, f_1(W) + q_c\, f_c(W)$.

The interior peak has density $f_c(W) = 1 / B(\alpha, \beta) \cdot W^{\alpha-1} (1 - W)^{\beta-1}$, defined on the interval $0 \leq W \leq 1$.

The Beta function $B(\alpha, \beta)$ provides the normalising constant for this density. It can also be written in terms of the Gamma function $\Gamma$ as $B(\alpha, \beta) = \Gamma(\alpha)\, \Gamma(\beta) / \Gamma(\alpha + \beta)$. $\Gamma(\alpha)$ is the Gamma function, if $\alpha$ is a positive integer n, then $\Gamma(n + 1)$ equals n! In this model we express the shape parameters as $\alpha = W_0 K$ and $\beta = (1 - W_0) K$. This parametrisation fixes the mean weight within the interior sub-population at $W_0$. The overall model mean weight equals $q_1 + q_c W_0$.

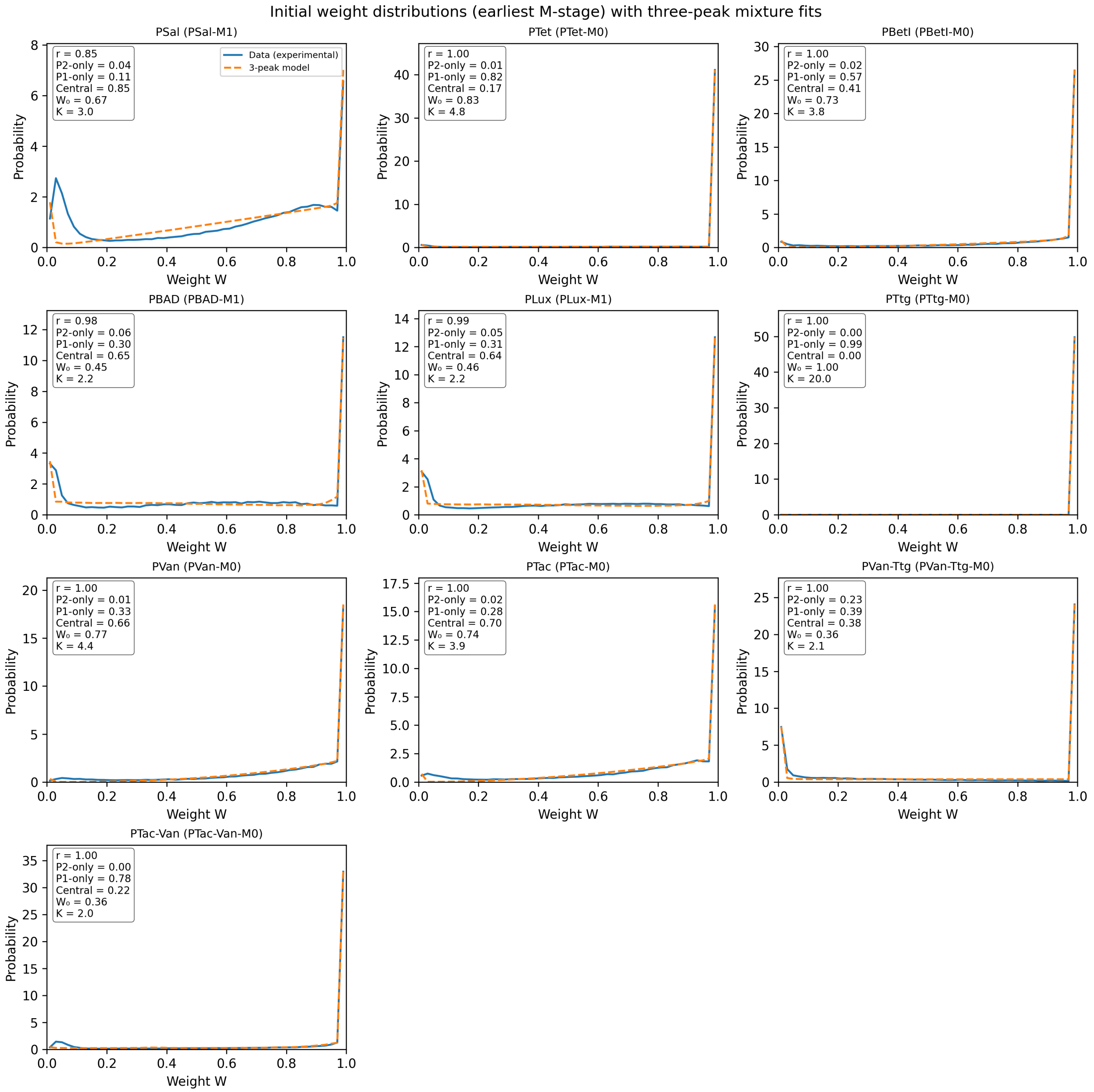

**Supplementary Fig. 34 | Fits to initial weight distributions.** For each promoter, the earliest M-stage weight distribution (solid line) and the corresponding three-peak mixture fit (dashed line) are shown on the interval from 0 to 1. Insets report the fitted mixture fractions, the interior-peak mean W0, the shape parameter *K* and the Pearson correlation between the experimental and model distributions.

To describe the initial single-cell weight distributions, we fitted, for each promoter, the earliest M-stage distribution with a mixture of three sub-populations, as shown in Supplementary Fig. 35. The experimental distribution appears as a normalised density on the interval from 0 to 1. The dashed curve in each panel shows the fitted mixture of a P2-like sub-population with weights close to 0, a P1-like sub-population with weights close to 1, and an interior sub-population with weights between 0 and 1.

The P2-like and P1-like components follow empirical templates taken from the P2-only (EGFP-only) and P1-only (mCherry-only) control wells. These templates are normalised so that their total probability on the interval from 0 to 1 equals one. The fitted mixture fractions $q_0$, $q_1$ and $q_c$ give the proportions of cells in the P2-like, P1-like and interior sub-populations, respectively, and they satisfy $q_0 + q_1 + q_c = 1$.

The interior sub-population follows a Beta distribution on the interval from 0 to 1 (see Supplementary Note 2 for the derivation). This distribution has a density that is proportional to the product of two factors: one factor is the weight raised to the power α minus 1 and the other factor is one minus the weight raised to the power β minus 1. The Beta function B(α, β) provides the normalising constant that makes the total probability equal to one. It can also be written in terms of the Gamma function Γ as $B(\alpha, \beta) = \Gamma(\alpha)\,\Gamma(\beta) / \Gamma(\alpha + \beta)$.

In this parametrisation the shape parameters of the interior peak satisfy $\alpha = W_0 K$ and $\beta = (1 - W_0) K$, where $W_0$ is the mean weight within the interior sub-population and $K$ controls how narrow or broad this peak is. With this choice the mean weight of the interior sub-population equals $W_0$ by construction. The overall model mean weight equals the fraction of cells in the P1-like component plus the product of $q_c$ and $W_0$, because the P2-like component is concentrated near weight 0. Each panel in Supplementary Fig. 35 reports the fitted values of $q_0$ (labelled "P2-only"), $q_1$ (labelled "P1-only"), $q_c$ (labelled "Central"), the interior-peak mean $W_0$, the shape parameter $K$, and the Pearson correlation between the experimental and model distributions.

We then summarised the quality and composition of the three-peak fits for all promoters (Supplementary Fig. 35). For each promoter, the second figure shows the experimental earliest M-stage weight distribution as a solid line and the corresponding mixture model as a dashed line, plotted on the same linear weight axis without legends. The text that follows reports, for every promoter, the Pearson correlation between the experimental and model distributions, the associated two-sided P value obtained from a t test with $n - 2$ degrees of freedom (where n is the number of histogram bins), the fitted fractions of the P2-like ($W$ close to 0), P1-like ($W$ close to 1) and interior sub-populations, and the implied model mean weight.

For PSal (PSal-M1), the three-peak mixture reproduced the experimental distribution with a Pearson correlation $r = 0.85$ and $P = 7.97 \times 10^{-15}$. A substantial interior sub-population was present, with a fitted fraction of about 0.85, mean weight around 0.67 and shape parameter $K$ close to 3.0. The mixture implied a mean population weight of approximately 0.68.

For PTet (PTet-M0), the three-peak mixture reproduced the experimental distribution with a Pearson correlation $r = 1.00$ and $P = 8.23 \times 10^{-101}$. The population was dominated by high-weight P1-like cells with a fitted fraction near $W = 1$ of about 0.82. The mixture implied a mean population weight of approximately 0.96.

For PBetI (PBetI-M0), the three-peak mixture reproduced the experimental distribution with a Pearson correlation $r = 1.00$ and $P = 1.15 \times 10^{-70}$. The population was dominated by high-weight P1-like cells with a fitted fraction near $W = 1$ of about 0.57. A substantial interior sub-population was present, with a fitted fraction of about 0.41, mean weight around 0.73 and shape parameter $K$ close to 3.8. The mixture implied a mean population weight of approximately 0.87.

For PBAD (PBAD-M1), the three-peak mixture reproduced the experimental distribution with a Pearson correlation $r = 0.98$ and $P = 3.68 \times 10^{-33}$. A substantial interior sub-population was present, with a fitted fraction of about 0.65, mean weight around 0.45 and shape parameter $K$ close to 2.2. The mixture implied a mean population weight of approximately 0.59.

For PLux (PLux-M1), the three-peak mixture reproduced the experimental distribution with a Pearson correlation $r = 0.99$ and $P = 1.24 \times 10^{-38}$. A substantial interior sub-population was present, with a fitted fraction of about 0.64, mean weight around 0.46 and shape parameter $K$ close to 2.2. The mixture implied a mean population weight of approximately 0.60.

For PTtg (PTtg-M0), the three-peak mixture reproduced the experimental distribution with a Pearson correlation r = 1.00 and P = $6.49 \times 10^{-164}$. The population was dominated by high-weight P1-like cells with a fitted fraction near $W = 1$ of about 0.99. The mixture implied a mean population weight of approximately 1.00.

For PVan (PVan-M0), the three-peak mixture reproduced the experimental distribution with a Pearson correlation r = 1.00 and P = $5.96 \times 10^{-60}$. A substantial interior sub-population was present, with a fitted fraction of about 0.66, mean weight around 0.77 and shape parameter $K$ close to 4.4. The mixture implied a mean population weight of approximately 0.84.

For PTac (PTac-M0), the three-peak mixture reproduced the experimental distribution with a Pearson correlation r = 1.00 and P = $4.27 \times 10^{-51}$. A substantial interior sub-population was present, with a fitted fraction of about 0.70, mean weight around 0.74 and shape parameter $K$ close to 3.9. The mixture implied a mean population weight of approximately 0.80.

For PVan-Ttg (PVan-Ttg-M0), the three-peak mixture reproduced the experimental distribution with a Pearson correlation r = 1.00 and P = $3.31 \times 10^{-58}$. A substantial interior sub-population was present, with a fitted fraction of about 0.38, mean weight around 0.36 and shape parameter $K$ close to 2.1. A lower-weight P2-like component contributed a non-negligible fraction near $W = 0$ (approximately 0.23). The mixture implied a mean population weight of approximately 0.53.

For PTac-Van (PTac-Van-M0), the three-peak mixture reproduced the experimental distribution with a Pearson correlation r = 1.00 and P = $5.00 \times 10^{-62}$. The population was dominated by high-weight P1-like cells with a fitted fraction near $W = 1$ of about 0.78. The mixture implied a mean population weight of approximately 0.86.

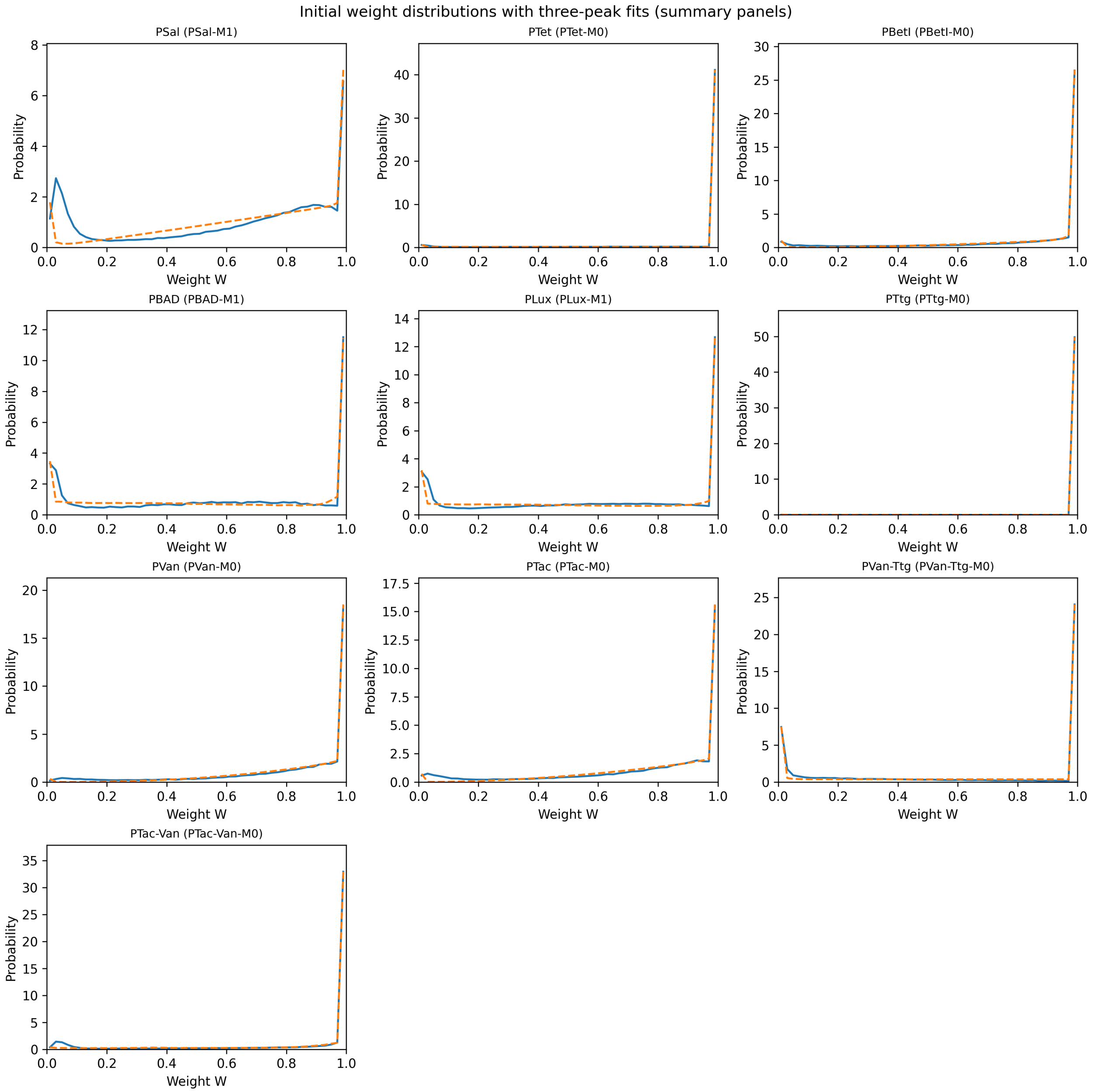

**Supplementary Fig. 35 | Summary of three-peak fits to initial weight distributions.** Each panel shows, for one promoter, the earliest M-stage weight distribution and its three-peak mixture fit, plotted as solid and dashed curves respectively without legends. Quantitative summaries of the correlation, P value and fitted component fractions are provided in the text.

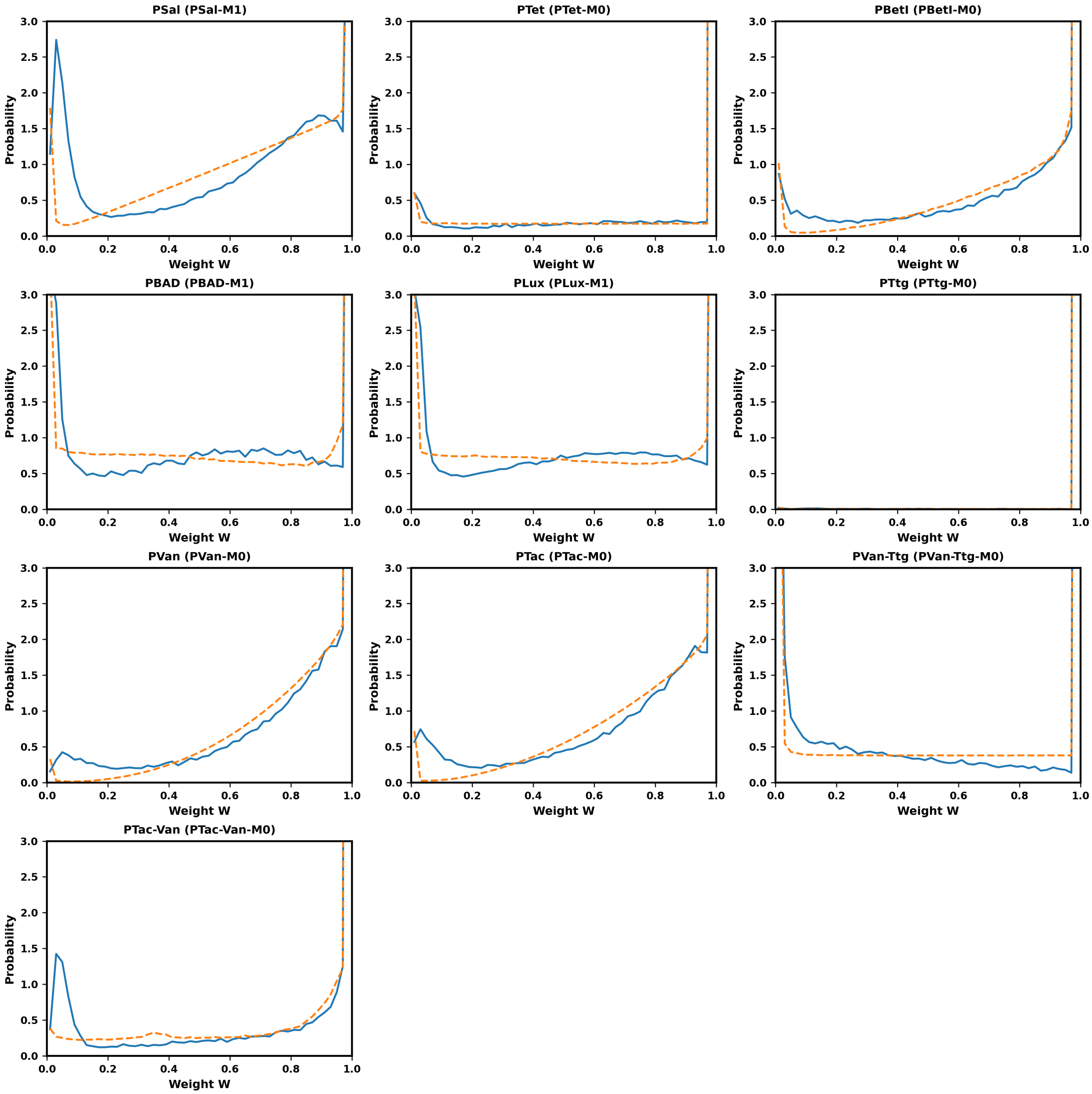

**Supplementary Fig. 36 | Summary of three-peak fits to initial weight distributions with small y-axis limit.** Same figure as Supplementary Fig. 35 but with common y-scale. The x-axis spans 0 to 1 and the y-axis spans 0 to 3.

#### Learning weight distributions and kanamycin learning predictions

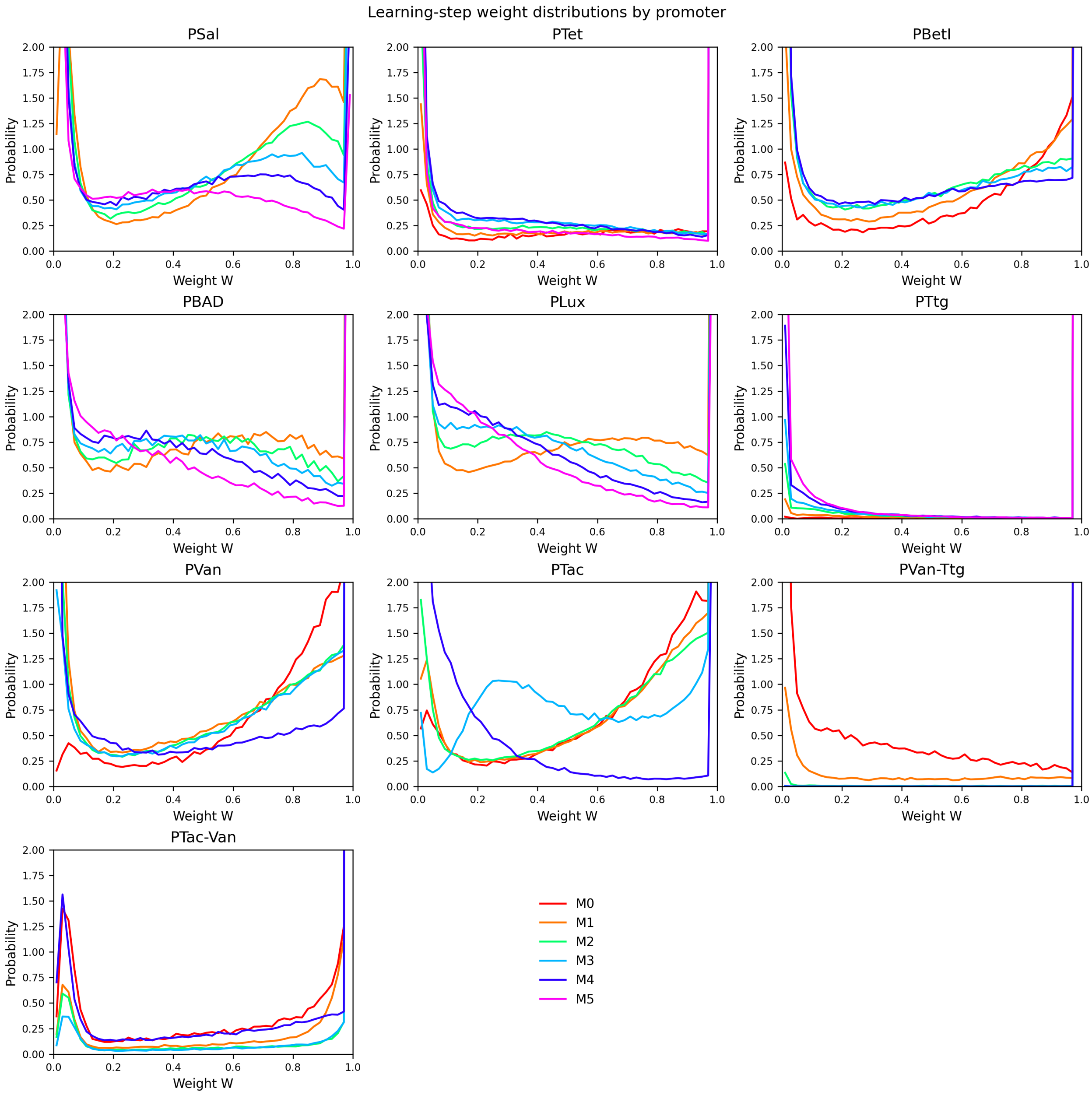

**Supplementary Fig. 37 | Learning-step weight distributions by promoter.** Each panel shows, for one promoter, the normalised single cell weight distributions for all learning steps. Curves are constructed from the pooled single cell weights after the same gating and calibration steps as in the other weight panels, using a common linear binning on the interval from 0 to 1 and normalising each histogram so that its area equals one. Panel specific mean weights <*W*> for each promoter and learning step, computed directly from the same pooled single cell data used for the curves, are: PSal: M1 <*W*>=0.621, M2 <*W*>=0.525, M3 <*W*>=0.448, M4 <*W*>=0.357, M5 <*W*>=0.257; PTet: M0 <*W*>=0.908, M1 <*W*>=0.871, M2 <*W*>=0.823, M3 <*W*>=0.768, M4 <*W*>=0.738, M5 <*W*>=0.816; PBetI: M0 <*W*>=0.825, M1 <*W*>=0.727, M2 <*W*>=0.609, M3 <*W*>=0.586, M4 <*W*>=0.552; PBAD: M1 <*W*>=0.573, M2 <*W*>=0.494, M3 <*W*>=0.447, M4 <*W*>=0.391, M5 <*W*>=0.316; PLux: M1 <*W*>=0.595, M2 <*W*>=0.454, M3 <*W*>=0.393, M4 <*W*>=0.312, M5 <*W*>=0.264; PTtg: M0 <*W*>=0.997, M1 <*W*>=0.986, M2 <*W*>=0.967, M3 <*W*>=0.951, M4 <*W*>=0.918, M5 <*W*>=0.869; PVan: M0 <*W*>=0.805, M1 <*W*>=0.618, M2 <*W*>=0.682, M3 <*W*>=0.710, M4 <*W*>=0.635; PTac: M0 <*W*>=0.763, M1 <*W*>=0.738, M2 <*W*>=0.723, M3 <*W*>=0.653, M4 <*W*>=0.175; PVan-Ttg: M0 <*W*>=0.616, M1 <*W*>=0.925, M2 <*W*>=0.994, M3 <*W*>=1.000, M4 <*W*>=1.000; PTac-Van: M0 <*W*>=0.836, M1 <*W*>=0.923, M2 <*W*>=0.943, M3 <*W*>=0.956, M4 <*W*>=0.846.

In Supplementary Fig. 37 we identified, for every well, labels ending with a learning-step suffix (e.g. -M0). For each promoter and learning step $M_n$, we pooled single-cell plasmid weights $W$ from all corresponding wells across replicates, after applying the fluorescence compensation and FSC/SSC gating described above. Let N be the total number of events in the pooled set. The mean weight for that step is the arithmetic average $\langle W \rangle = (1/N) \sum W_i$. To draw the distributions, we constructed histograms on the interval [0, 1] using 50 equal bins. The bin edges $e_j$ were equally spaced, with bin width $\Delta W = e_{j+1} - e_j$. For each bin j, let $C_j$ be the count of events falling in that bin. The probability density $P(W)$ at the bin center was calculated as $P(W) = C_j / (N \cdot \Delta W)$, ensuring that the total area under the curve equals 1. All panels share the same binning and vertical axis limits (0 to 2) to allow direct comparison. The learning step index n is encoded by colour using a rainbow gradient (Red to Purple) excluding yellow for visibility.

To interpret the changes in weight distributions between consecutive learning steps, we compared the experimental data to a biased-diffusion prediction. The solid line in each panel shows the observed distribution at step $M_{n+1}$. The dashed line shows the predicted distribution derived from the previous step $M_n$ by applying a growth-bias operator.

The predicted probability density $P_{pred}(W)$ is calculated from the observed density at the previous step $P_{obs}(W)$ according to the relation $P_{pred}(W) = (1/Z) \cdot P_{obs}(W) \cdot \exp(-\kappa\tau \cdot W)$.

Here, $\kappa\tau$ represents the dimensionless growth-bias strength parameter. For these overlays, we fixed $\kappa\tau = 1$. The factor Z is a normalisation constant, defined as the integral of $P_{obs}(W) \cdot \exp(-\kappa\tau \cdot W)$ over the interval from 0 to 1, ensuring that the total probability integrates to one.

For promoter PSal, Supplementary Fig. 38 compares the experimental single-cell weight distributions and the biased-diffusion predictions across learning steps M1, M2, M3, M4 and M5. Under the growth-bias operator, we expect the probability at $W$=0 to increase, the probability at $W$=1 to decrease, and the mean weight $<W>$ to decrease over successive learning steps.

Across consecutive transitions, the changes in boundary probabilities, the intermediate probability near $W\approx0.5$, and the mean weight $<W>$ are: M1→M2: P(0) increases, P(1) decreases, $P(W\approx0.5)$ increases, and $<W>$ decreases (from 0.62 to 0.53); M2→M3: P(0) increases, P(1) decreases, $P(W\approx0.5)$ increases, and $<W>$ decreases (from 0.53 to 0.45); M3→M4: P(0) increases, P(1) decreases, $P(W\approx0.5)$ decreases, and $<W>$ decreases (from 0.45 to 0.36); M4→M5: P(0) increases, P(1) decreases, $P(W\approx0.5)$ decreases, and $<W>$ decreases (from 0.36 to 0.26).

The expected pattern with increasing P(0), decreasing P(1), and decreasing $<W>$ is observed in transitions M1→M2, M2→M3, M3→M4 and M4→M5.

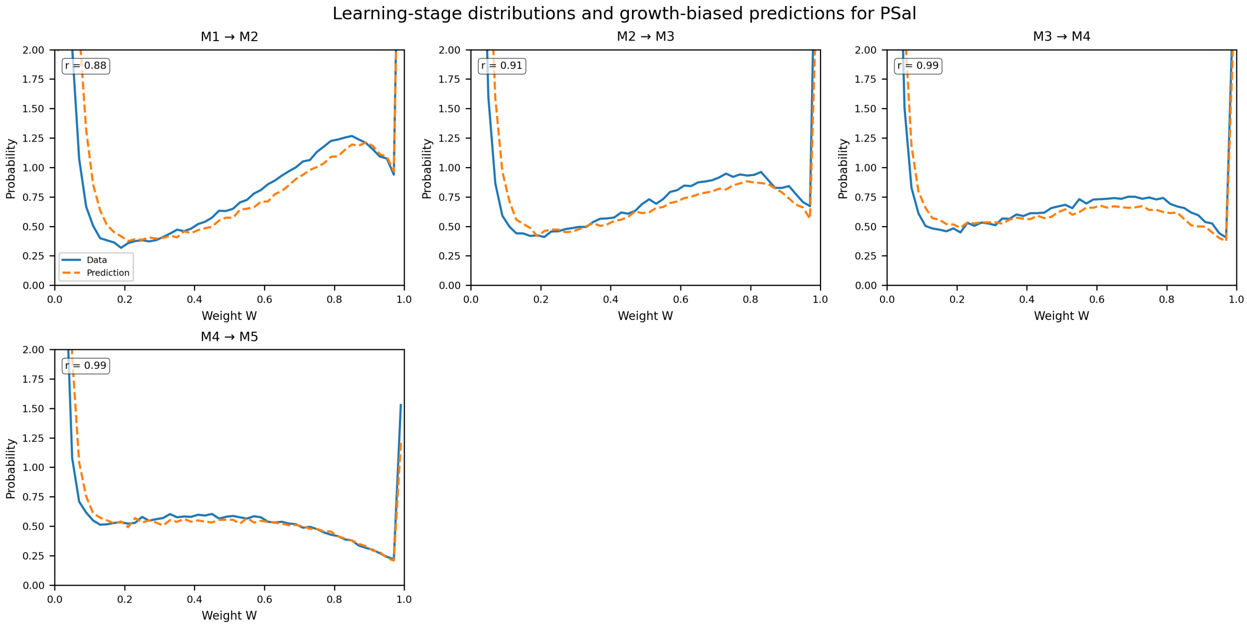

**Supplementary Fig. 38 | Learning-stage distributions and biased-diffusion predictions for PSal.** Each panel compares the experimental single-cell weight distribution at stage Mn+1 (solid line) with the prediction derived from stage Mn (dashed line) using the biased-diffusion model with κτ = 1. Pearson correlation coefficients **r**, two-sided P values, and boundary densities (P(0), P(1)) for each transition were: M1 → M2: r = 0.88, P < 0.001. Obs P(0)=4.23, P(1)=4.43; Pred P(0)=1.99, P(1)=4.27; M2 → M3: r = 0.91, P < 0.001. Obs P(0)=8.74, P(1)=3.80; Pred P(0)=6.64, P(1)=2.61; M3 → M4: r = 0.99, P < 0.001. Obs P(0)=13.82, P(1)=2.40; Pred P(0)=12.72, P(1)=2.08; M4 → M5: r = 0.99, P < 0.001. Obs P(0)=21.40, P(1)=1.53; Pred P(0)=18.50, P(1)=1.21.

For promoter PTet, Supplementary Fig. 39 compares the experimental single-cell weight distributions and the biased-diffusion predictions across learning steps M0, M1, M2, M3, M4 and M5. Under the growth-bias operator, we expect the probability at $W$=0 to increase, the probability at $W$=1 to decrease, and the mean weight <$W$> to decrease over successive learning steps.

Across consecutive transitions, the changes in boundary probabilities, the intermediate probability near $W$≈0.5, and the mean weight <$W$> are: M0→M1: P(0) increases, P(1) decreases, P($W$≈0.5) increases, and <$W$> decreases (from 0.91 to 0.87); M1→M2: P(0) increases, P(1) decreases, P($W$≈0.5) increases, and <$W$> decreases (from 0.87 to 0.82); M2→M3: P(0) increases, P(1) decreases, P($W$≈0.5) increases, and <$W$> decreases (from 0.82 to 0.77); M3→M4: P(0) increases, P(1) decreases, P($W$≈0.5) decreases, and <$W$> decreases (from 0.77 to 0.74); M4→M5: P(0) decreases, P(1) increases, P($W$≈0.5) decreases, and <$W$> increases (from 0.74 to 0.82).

The expected pattern with increasing P(0), decreasing P(1), and decreasing <$W$> is observed in transitions M0→M1, M1→M2, M2→M3 and M3→M4.

Other transitions deviate from this simple trend in at least one of the boundary probabilities, the intermediate probability, or the mean weight, indicating partial disagreement with the biased-diffusion expectation in M4→M5.

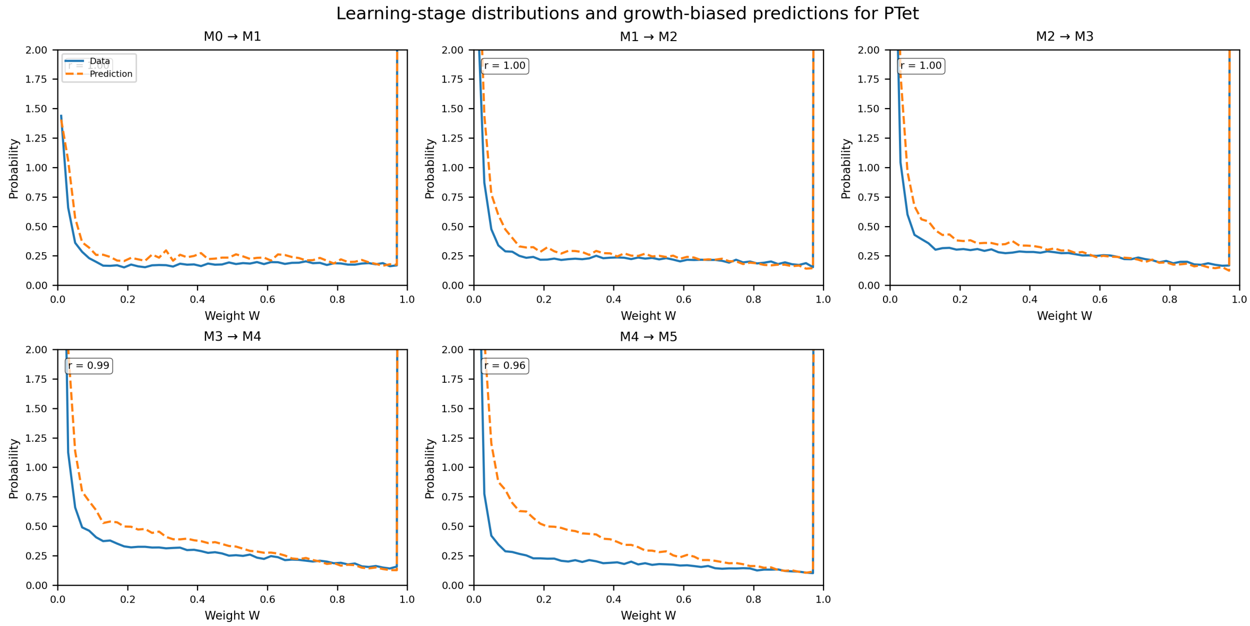

**Supplementary Fig. 39 | Learning-stage distributions and biased-diffusion predictions for PTet.** Each panel compares the experimental single-cell weight distribution at stage Mn+1 (solid line) with the prediction derived from stage Mn (dashed line) using the biased-diffusion model with $\kappa\tau = 1$. Pearson correlation coefficients **r**, two-sided P values, and boundary densities (P(0), P(1)) for each transition were: M0 → M1: r = 1.00, P < 0.001. Obs P(0)=1.44, P(1)=39.17; Pred P(0)=1.41, P(1)=36.35; M1 → M2: r = 1.00, P < 0.001. Obs P(0)=2.40, P(1)=36.15; Pred P(0)=3.22, P(1)=32.89; M2 → M3: r = 1.00, P < 0.001. Obs P(0)=3.68, P(1)=32.78; Pred P(0)=5.05, P(1)=28.57; M3 → M4: r = 0.99, P < 0.001. Obs P(0)=4.67, P(1)=31.45; Pred P(0)=7.25, P(1)=24.22; M4 → M5: r = 0.96, P < 0.001. Obs P(0)=3.53, P(1)=36.98; Pred P(0)=8.87, P(1)=22.42.

For promoter PBetI, Supplementary Fig. 40 compares the experimental single-cell weight distributions and the biased-diffusion predictions across learning steps M0, M1, M2, M3 and M4. Under the growth-bias operator, we expect the probability at *W*=0 to increase, the probability at *W*=1 to decrease, and the mean weight <*W*> to decrease over successive learning steps.

Across consecutive transitions, the changes in boundary probabilities, the intermediate probability near *W*≈0.5, and the mean weight <*W*> are: M0→M1: P(0) increases, P(1) decreases, P(*W*≈0.5) increases, and <*W*> decreases (from 0.82 to 0.73); M1→M2: P(0) increases, P(1) decreases, P(*W*≈0.5) increases, and <*W*> decreases (from 0.73 to 0.61); M2→M3: P(0) increases, P(1) decreases, P(*W*≈0.5) is approximately unchanged, and <*W*> decreases (from 0.61 to 0.59); M3→M4: P(0) increases, P(1) decreases, P(*W*≈0.5) decreases, and <*W*> decreases (from 0.59 to 0.55).

The expected pattern with increasing P(0), decreasing P(1), and decreasing <*W*> is observed in transitions M0→M1, M1→M2, M2→M3 and M3→M4.

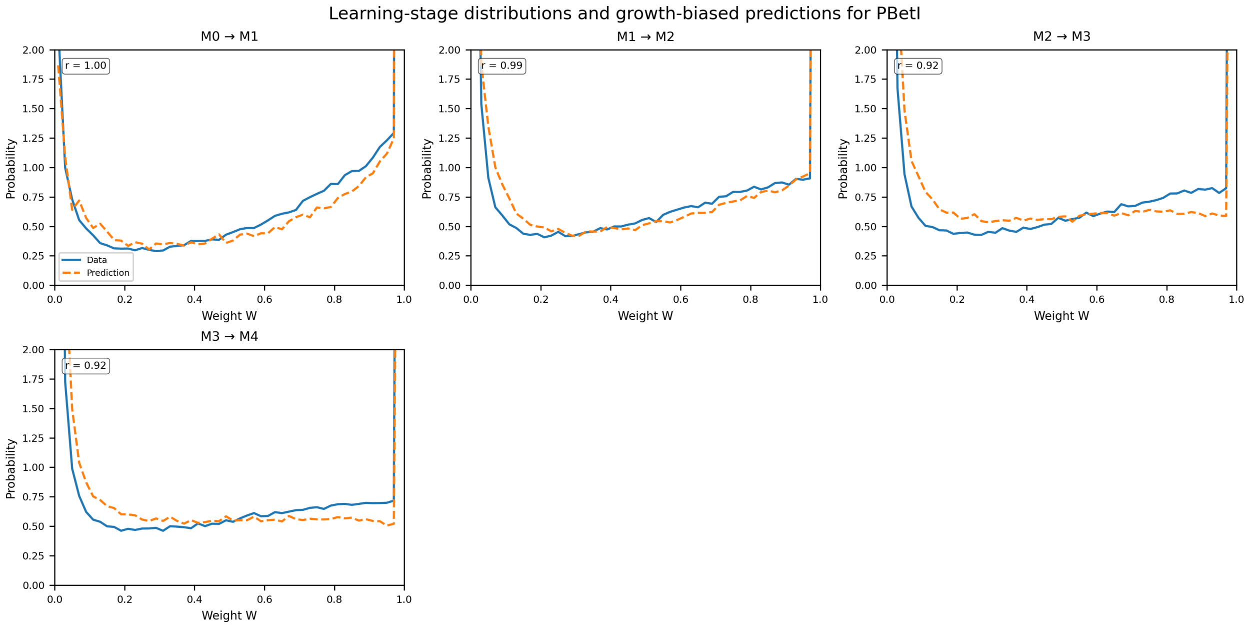

**Supplementary Fig. 40 | Learning-stage distributions and biased-diffusion predictions for PBetI.** Each panel compares the experimental single-cell weight distribution at stage Mn+1 (solid line) with the prediction derived from stage Mn (dashed line) using the biased-diffusion model with κτ = 1. Pearson correlation coefficients **r**, two-sided P values, and boundary densities (P(0), P(1)) for each transition were: M0 → M1: r = 1.00, P < 0.001. Obs P(0)=2.20, P(1)=19.32; Pred P(0)=1.87, P(1)=21.32; M1 → M2: r = 0.99, P < 0.001. Obs P(0)=5.29, P(1)=13.67; Pred P(0)=4.22, P(1)=13.93; M2 → M3: r = 0.92, P < 0.001. Obs P(0)=6.42, P(1)=13.40; Pred P(0)=8.96, P(1)=8.68; M3 → M4: r = 0.92, P < 0.001. Obs P(0)=7.69, P(1)=12.75; Pred P(0)=10.59, P(1)=8.30.

For promoter PBAD, Supplementary Fig. 41 compares the experimental single-cell weight distributions and the biased-diffusion predictions across learning steps M1, M2, M3, M4 and M5. Under the growth-bias operator, we expect the probability at $W$=0 to increase, the probability at $W$=1 to decrease, and the mean weight <$W$> to decrease over successive learning steps.

Across consecutive transitions, the changes in boundary probabilities, the intermediate probability near $W$≈0.5, and the mean weight <$W$> are: M1→M2: P(0) increases, P(1) decreases, P($W$≈0.5) decreases, and <$W$> decreases (from 0.57 to 0.49); M2→M3: P(0) increases, P(1) decreases, P($W$≈0.5) increases, and <$W$> decreases (from 0.49 to 0.45); M3→M4: P(0) increases, P(1) decreases, P($W$≈0.5) decreases, and <$W$> decreases (from 0.45 to 0.39); M4→M5: P(0) increases, P(1) decreases, P($W$≈0.5) decreases, and <$W$> decreases (from 0.39 to 0.32).

The expected pattern with increasing P(0), decreasing P(1), and decreasing <$W$> is observed in transitions M1→M2, M2→M3, M3→M4 and M4→M5.

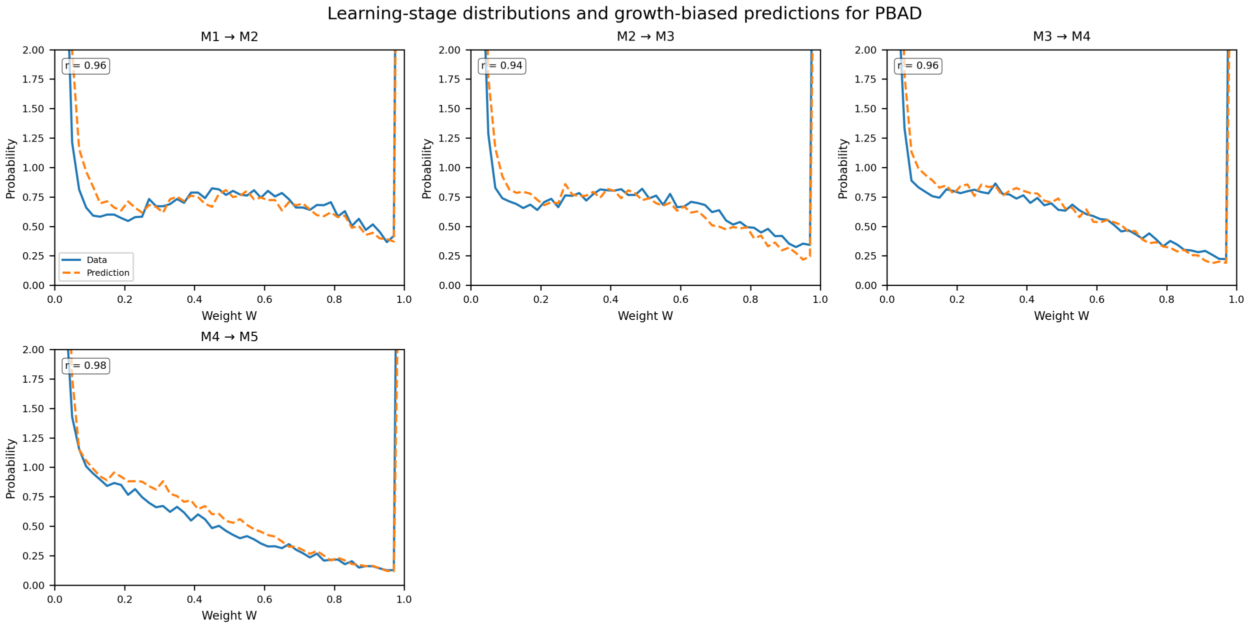

**Supplementary Fig. 41 | Learning-stage distributions and biased-diffusion predictions for PBAD.** Each panel compares the experimental single-cell weight distribution at stage Mn+1 (solid line) with the prediction derived from stage Mn (dashed line) using the biased-diffusion model with κτ = 1. Pearson correlation coefficients **r**, two-sided P values, and boundary densities (P(0), P(1)) for each transition were: M1 → M2: r = 0.96, P < 0.001. Obs P(0)=6.13, P(1)=9.09; Pred P(0)=5.45, P(1)=7.09; M2 → M3: r = 0.94, P < 0.001. Obs P(0)=7.92, P(1)=8.02; Pred P(0)=9.32, P(1)=5.19; M3 → M4: r = 0.96, P < 0.001. Obs P(0)=11.36, P(1)=7.53; Pred P(0)=11.51, P(1)=4.37; M4 → M5: r = 0.98, P < 0.001. Obs P(0)=16.71, P(1)=7.24; Pred P(0)=15.64, P(1)=3.89.

For promoter PLux, Supplementary Fig. 42 compares the experimental single-cell weight distributions and the biased-diffusion predictions across learning steps M1, M2, M3, M4 and M5. Under the growth-bias operator, we expect the probability at $W$=0 to increase, the probability at $W$=1 to decrease, and the mean weight <$W$> to decrease over successive learning steps.

Across consecutive transitions, the changes in boundary probabilities, the intermediate probability near $W$≈0.5, and the mean weight <$W$> are: M1→M2: P(0) increases, P(1) decreases, P($W$≈0.5) increases, and <$W$> decreases (from 0.59 to 0.45); M2→M3: P(0) increases, P(1) decreases, P($W$≈0.5) decreases, and <$W$> decreases (from 0.45 to 0.39); M3→M4: P(0) increases, P(1) decreases, P($W$≈0.5) decreases, and <$W$> decreases (from 0.39 to 0.31); M4→M5: P(0) increases, P(1) decreases, P($W$≈0.5) decreases, and <$W$> decreases (from 0.31 to 0.26).

The expected pattern with increasing P(0), decreasing P(1), and decreasing <$W$> is observed in transitions M1→M2, M2→M3, M3→M4 and M4→M5.

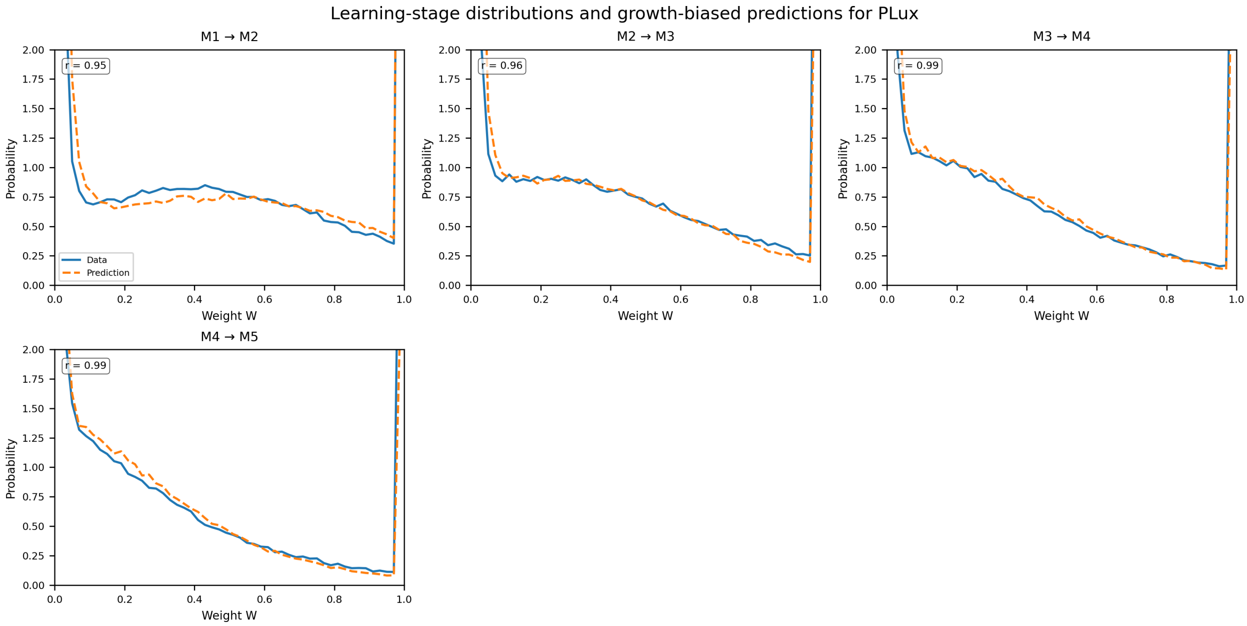

**Supplementary Fig. 42 | Learning-stage distributions and biased-diffusion predictions for PLux.** Each panel compares the experimental single-cell weight distribution at stage Mn+1 (solid line) with the prediction derived from stage Mn (dashed line) using the biased-diffusion model with κτ = 1. Pearson correlation coefficients **r**, two-sided P values, and boundary densities (P(0), P(1)) for each transition were: M1 → M2: r = 0.95, P < 0.001. Obs P(0)=7.55, P(1)=7.55; Pred P(0)=5.16, P(1)=7.99; M2 → M3: r = 0.96, P < 0.001. Obs P(0)=10.30, P(1)=6.55; Pred P(0)=11.08, P(1)=4.15; M3 → M4: r = 0.99, P < 0.001. Obs P(0)=14.41, P(1)=5.06; Pred P(0)=14.26, P(1)=3.41; M4 → M5: r = 0.99, P < 0.001. Obs P(0)=17.60, P(1)=4.75; Pred P(0)=18.55, P(1)=2.44.

For promoter PTtg, Supplementary Fig. 43 compares the experimental single-cell weight distributions and the biased-diffusion predictions across learning steps M0, M1, M2, M3, M4 and M5. Under the growth-bias operator, we expect the probability at *W*=0 to increase, the probability at *W*=1 to decrease, and the mean weight <*W*> to decrease over successive learning steps.

Across consecutive transitions, the changes in boundary probabilities, the intermediate probability near *W*≈0.5, and the mean weight <*W*> are: M0→M1: P(0) increases, P(1) decreases, P(*W*≈0.5) is approximately unchanged, and <*W*> decreases (from 1.00 to 0.99); M1→M2: P(0) increases, P(1) decreases, P(*W*≈0.5) is approximately unchanged, and <*W*> decreases (from 0.99 to 0.97); M2→M3: P(0) increases, P(1) decreases, P(*W*≈0.5) is approximately unchanged, and <*W*> decreases (from 0.97 to 0.95); M3→M4: P(0) increases, P(1) decreases, P(*W*≈0.5) is approximately unchanged, and <*W*> decreases (from 0.95 to 0.92); M4→M5: P(0) increases, P(1) decreases, P(*W*≈0.5) is approximately unchanged, and <*W*> decreases (from 0.92 to 0.87).

The expected pattern with increasing P(0), decreasing P(1), and decreasing <*W*> is observed in transitions M0→M1, M1→M2, M2→M3, M3→M4 and M4→M5.

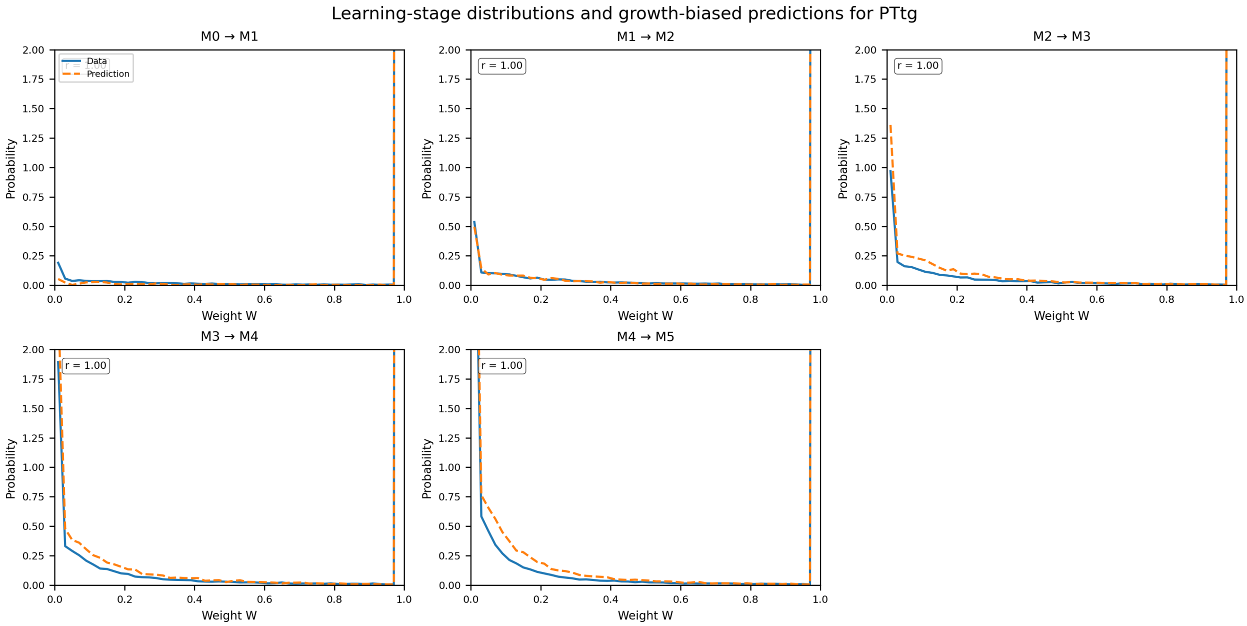

**Supplementary Fig. 43 | Learning-stage distributions and biased-diffusion predictions for PTtg.** Each panel compares the experimental single-cell weight distribution at stage Mn+1 (solid line) with the prediction derived from stage Mn (dashed line) using the biased-diffusion model with κτ = 1. Pearson correlation coefficients **r**, two-sided P values, and boundary densities (P(0), P(1)) for each transition were: M0 → M1: r = 1.00, P < 0.001. Obs P(0)=0.19, P(1)=49.06; Pred P(0)=0.05, P(1)=49.60; M1 → M2: r = 1.00, P < 0.001. Obs P(0)=0.54, P(1)=47.94; Pred P(0)=0.50, P(1)=48.01; M2 → M3: r = 1.00, P < 0.001. Obs P(0)=0.97, P(1)=47.02; Pred P(0)=1.36, P(1)=45.60; M3 → M4: r = 1.00, P < 0.001. Obs P(0)=1.89, P(1)=45.29; Pred P(0)=2.39, P(1)=43.61; M4 → M5: r = 1.00, P < 0.001. Obs P(0)=3.72, P(1)=42.79; Pred P(0)=4.46, P(1)=40.06.

For promoter PVan, Supplementary Fig. 44 compares the experimental single-cell weight distributions and the biased-diffusion predictions across learning steps M0, M1, M2, M3 and M4. Under the growth-bias operator, we expect the probability at $W$=0 to increase, the probability at $W$=1 to decrease, and the mean weight <$W$> to decrease over successive learning steps.

Across consecutive transitions, the changes in boundary probabilities, the intermediate probability near $W$≈0.5, and the mean weight <$W$> are: M0→M1: P(0) increases, P(1) decreases, P($W$≈0.5) increases, and <$W$> decreases (from 0.80 to 0.62); M1→M2: P(0) decreases, P(1) increases, P($W$≈0.5) decreases, and <$W$> increases (from 0.62 to 0.68); M2→M3: P(0) decreases, P(1) increases, P($W$≈0.5) is approximately unchanged, and <$W$> increases (from 0.68 to 0.71); M3→M4: P(0) increases, P(1) increases, P($W$≈0.5) decreases, and <$W$> decreases (from 0.71 to 0.64).

In transition M0→M1, the data follow the expected pattern: P(0) increases, P(1) decreases, and <$W$> decreases.

Other transitions deviate from this simple trend in at least one of the boundary probabilities, the intermediate probability, or the mean weight, indicating partial disagreement with the biased-diffusion expectation in M1→M2, M2→M3 and M3→M4.

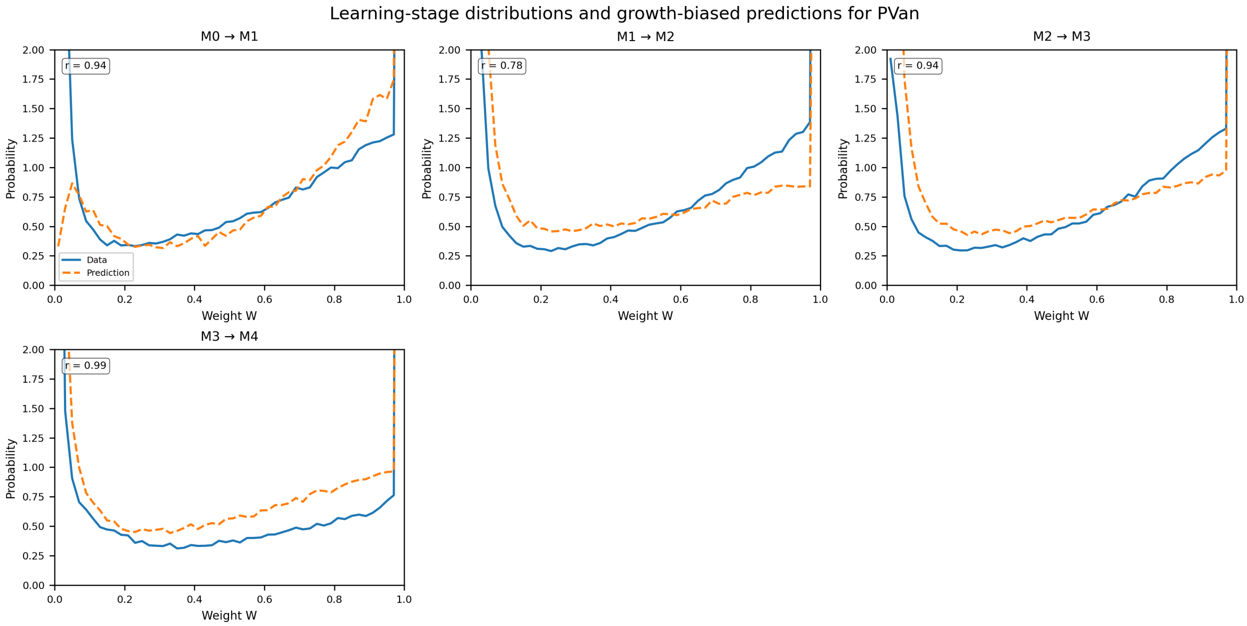

**Supplementary Fig. 44 | Learning-stage distributions and biased-diffusion predictions for PVan.** Each panel compares the experimental single-cell weight distribution at stage Mn+1 (solid line) with the prediction derived from stage Mn (dashed line) using the biased-diffusion model with $\kappa\tau = 1$. Pearson correlation coefficients **r**, two-sided P values, and boundary densities (P(0), P(1)) for each transition were: M0 → M1: r = 0.94, P < 0.001. Obs P(0)=3.49, P(1)=11.39; Pred P(0)=0.33, P(1)=14.62; M1 → M2: r = 0.78, P < 0.001. Obs P(0)=2.17, P(1)=14.89; Pred P(0)=5.98, P(1)=7.32; M2 → M3: r = 0.94, P < 0.001. Obs P(0)=1.92, P(1)=16.85; Pred P(0)=3.98, P(1)=10.25; M3 → M4: r = 0.99, P < 0.001. Obs P(0)=6.39, P(1)=19.89; Pred P(0)=3.64, P(1)=11.97.

For promoter PTac, Supplementary Fig. 45 compares the experimental single-cell weight distributions and the biased-diffusion predictions across learning steps M0, M1, M2, M3 and M4. Under the growth-bias operator, we expect the probability at $W$=0 to increase, the probability at $W$=1 to decrease, and the mean weight $<W>$ to decrease over successive learning steps.

Across consecutive transitions, the changes in boundary probabilities, the intermediate probability near $W$≈0.5, and the mean weight $<W>$ are: M0→M1: P(0) increases, P(1) increases, P($W$≈0.5) is approximately unchanged, and $<W>$ decreases (from 0.76 to 0.74); M1→M2: P(0) increases, P(1) decreases, P($W$≈0.5) increases, and $<W>$ decreases (from 0.74 to 0.72); M2→M3: P(0) decreases, P(1) decreases, P($W$≈0.5) increases, and $<W>$ decreases (from 0.72 to 0.65); M3→M4: P(0) increases, P(1) decreases, P($W$≈0.5) decreases, and $<W>$ decreases (from 0.65 to 0.18).

The expected pattern with increasing P(0), decreasing P(1), and decreasing $<W>$ is observed in transitions M1→M2 and M3→M4.

Other transitions deviate from this simple trend in at least one of the boundary probabilities, the intermediate probability, or the mean weight, indicating partial disagreement with the biased-diffusion expectation in M0→M1 and M2→M3.

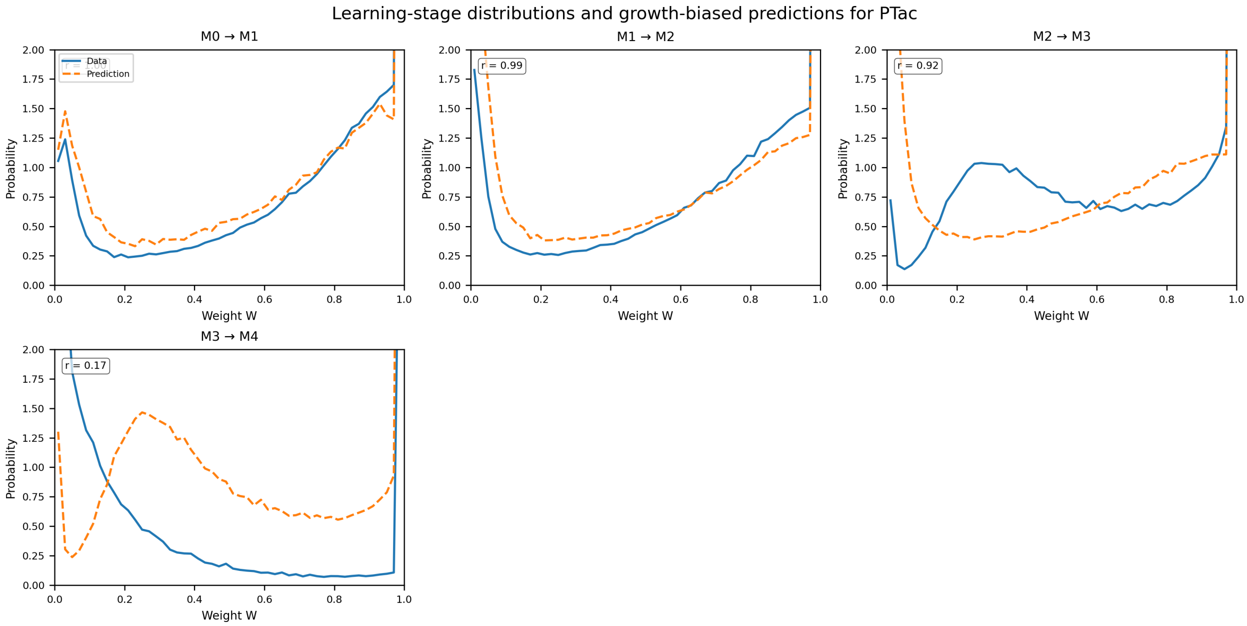

**Supplementary Fig. 45 | Learning-stage distributions and biased-diffusion predictions for PTac.** Each panel compares the experimental single-cell weight distribution at stage Mn+1 (solid line) with the prediction derived from stage Mn (dashed line) using the biased-diffusion model with κτ = 1. Pearson correlation coefficients **r**, two-sided P values, and boundary densities (P(0), P(1)) for each transition were: M0 → M1: r = 1.00, P < 0.001. Obs P(0)=1.05, P(1)=15.91; Pred P(0)=1.15, P(1)=11.76; M1 → M2: r = 0.99, P < 0.001. Obs P(0)=1.83, P(1)=15.73; Pred P(0)=2.07, P(1)=11.70; M2 → M3: r = 0.92, P < 0.001. Obs P(0)=0.72, P(1)=13.40; Pred P(0)=3.52, P(1)=11.37; M3 → M4: r = 0.17, P = 0.227. Obs P(0)=26.43, P(1)=4.46; Pred P(0)=1.30, P(1)=9.09.

For promoter PVan-Ttg, Supplementary Fig. 46 compares the experimental single-cell weight distributions and the biased-diffusion predictions across learning steps M0, M1, M2, M3 and M4. Under the growth-bias operator, we expect the probability at *W*=0 to increase, the probability at *W*=1 to decrease, and the mean weight <*W*> to decrease over successive learning steps.

Across consecutive transitions, the changes in boundary probabilities, the intermediate probability near *W*≈0.5, and the mean weight <*W*> are: M0→M1: P(0) decreases, P(1) increases, P(*W*≈0.5) decreases, and <*W*> increases (from 0.62 to 0.93); M1→M2: P(0) decreases, P(1) increases, P(*W*≈0.5) decreases, and <*W*> increases (from 0.93 to 0.99); M2→M3: P(0) decreases, P(1) increases, P(*W*≈0.5) is approximately unchanged, and <*W*> is approximately unchanged (from 0.99 to 1.00); M3→M4: P(0) is approximately unchanged, P(1) is approximately unchanged, P(*W*≈0.5) is approximately unchanged, and <*W*> is approximately unchanged (from 1.00 to 1.00).

Other transitions deviate from this simple trend in at least one of the boundary probabilities, the intermediate probability, or the mean weight, indicating partial disagreement with the biased-diffusion expectation in M0→M1, M1→M2, M2→M3 and M3→M4.

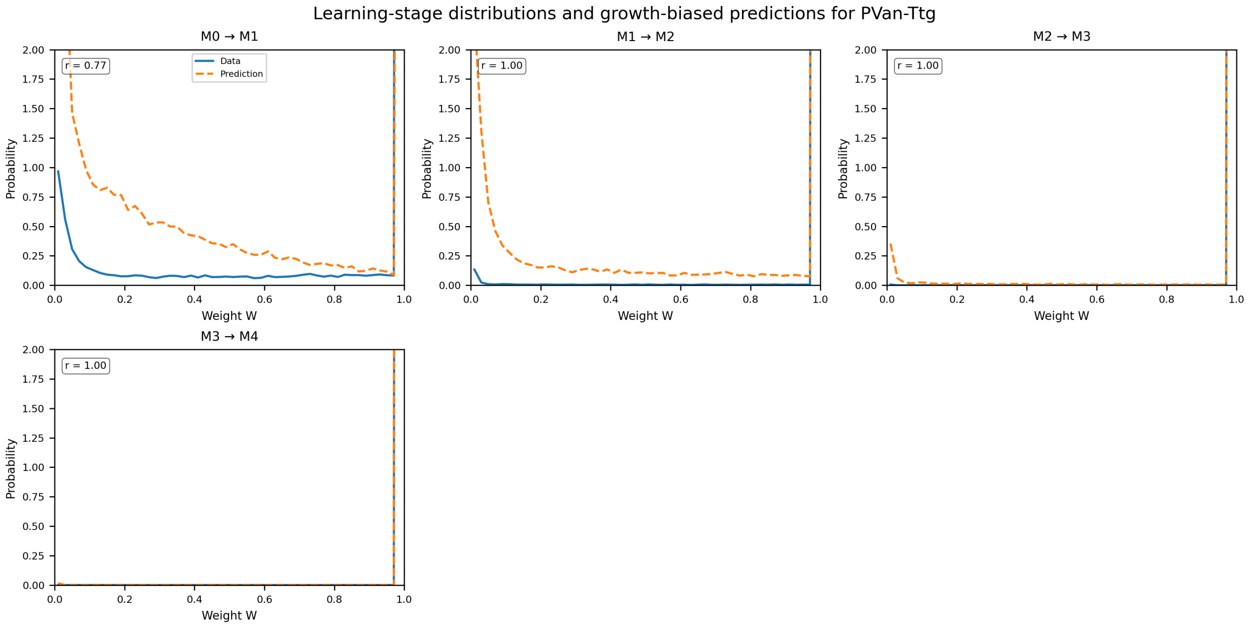

**Supplementary Fig. 46 | Learning-stage distributions and biased-diffusion predictions for PVan-Ttg.** Each panel compares the experimental single-cell weight distribution at stage Mn+1 (solid line) with the prediction derived from stage Mn (dashed line) using the biased-diffusion model with κT = 1. Pearson correlation coefficients **r**, two-sided P values, and boundary densities (P(0), P(1)) for each transition were: M0 → M1: r = 0.77, P < 0.001. Obs P(0)=0.97, P(1)=44.32; Pred P(0)=12.45, P(1)=15.08; M1 → M2: r = 1.00, P < 0.001. Obs P(0)=0.13, P(1)=49.59; Pred P(0)=2.31, P(1)=39.78; M2 → M3: r = 1.00, P < 0.001. Obs P(0)=0.01, P(1)=49.99; Pred P(0)=0.35, P(1)=49.16; M3 → M4: r = 1.00, P < 0.001. Obs P(0)=0.00, P(1)=50.00; Pred P(0)=0.02, P(1)=49.98.

For promoter PTac-Van, Supplementary Fig. 47 compares the experimental single-cell weight distributions and the biased-diffusion predictions across learning steps M0, M1, M2, M3 and M4. Under the growth-bias operator, we expect the probability at $W$=0 to increase, the probability at $W$=1 to decrease, and the mean weight $\langle W \rangle$ to decrease over successive learning steps.

Across consecutive transitions, the changes in boundary probabilities, the intermediate probability near $W$≈0.5, and the mean weight $\langle W \rangle$ are: M0→M1: P(0) decreases, P(1) increases, P($W$≈0.5) decreases, and $\langle W \rangle$ increases (from 0.84 to 0.92); M1→M2: P(0) decreases, P(1) increases, P($W$≈0.5) decreases, and $\langle W \rangle$ increases (from 0.92 to 0.94); M2→M3: P(0) decreases, P(1) increases, P($W$≈0.5) is approximately unchanged, and $\langle W \rangle$ increases (from 0.94 to 0.96); M3→M4: P(0) increases, P(1) decreases, P($W$≈0.5) increases, and $\langle W \rangle$ decreases (from 0.96 to 0.85).

In transition M3→M4, the data follow the expected pattern: P(0) increases, P(1) decreases, and $\langle W \rangle$ decreases.

Other transitions deviate from this simple trend in at least one of the boundary probabilities, the intermediate probability, or the mean weight, indicating partial disagreement with the biased-diffusion expectation in M0→M1, M1→M2 and M2→M3.

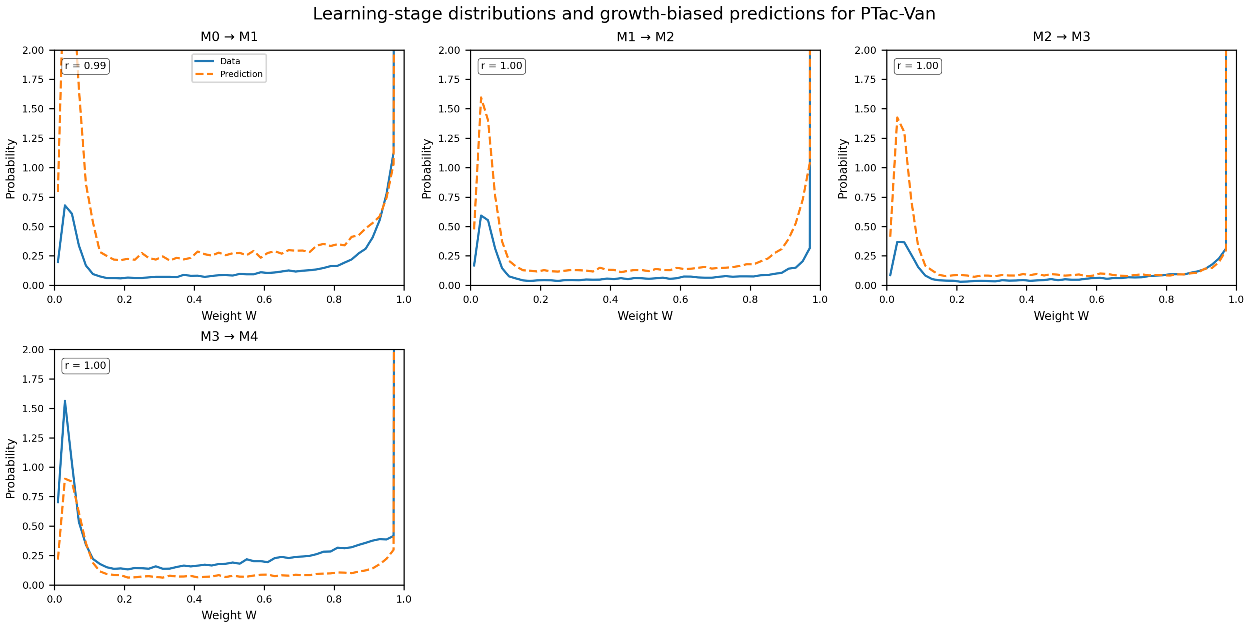

**Supplementary Fig. 47 | Learning-stage distributions and biased-diffusion predictions for PTac-Van.** Each panel compares the experimental single-cell weight distribution at stage Mn+1 (solid line) with the prediction derived from stage Mn (dashed line) using the biased-diffusion model with κτ = 1. Pearson correlation coefficients **r**, two-sided P values, and boundary densities (P(0), P(1)) for each transition were: M0 ⟶ M1: r = 0.99, P < 0.001. Obs P(0)=0.20, P(1)=40.78; Pred P(0)=0.79, P(1)=26.58; M1 ⟶ M2: r = 1.00, P < 0.001. Obs P(0)=0.17, P(1)=44.97; Pred P(0)=0.47, P(1)=36.71; M2 ⟶ M3: r = 1.00, P < 0.001. Obs P(0)=0.09, P(1)=45.62; Pred P(0)=0.41, P(1)=41.43; M3 ⟶ M4: r = 1.00, P < 0.001. Obs P(0)=0.70, P(1)=36.06; Pred P(0)=0.21, P(1)=42.82.

#### Supplementary Note 5 : Bacterial co-culture tournament protocol

This note details the experimental protocol used for the externally routed bacterial-player co-culture tic-tac-toe lesson sequence (main text Fig. 3).
M-states (e.g., M0, M1, M2) denote discrete stored-memory generations of a memregulon population after defined passaging or learning cycles; they are not continuous time units. "Day" refers to calendar or daily serial-sampling axes, including stability passaging and the co-culture tournament, and is defined in the relevant figure captions. In the tic-tac-toe SI, "memory generation" denotes the inherited culture state used at the start of each daily lesson.

We list the initial co-culture compositions (Supplementary Table 38), including the promoter identities and M-stages for each board position. We then describe the preparation of initial co-cultures:

1. Individual memregulon strains were inoculated from −80 °C glycerol stocks into LB medium supplemented with carbenicillin and grown overnight at 37 °C.
2. Overnight cultures were mixed in equal volumes to generate the co-culture, designated **M0 (day 0)**. The mixed culture was refreshed by a 1:2000 dilution into M9 medium containing carbenicillin and the cognate inducer. Cultures were incubated for 3 h at 37 °C with shaking at 200 rpm for characterization. After 3 h, 200 µL from each culture was dispensed into a 96-well plate (Custom Corning Costar) with both technical and biological replicates. Plates were loaded into an Infinite F500 microplate reader (Tecan) and incubated at 37 °C with shaking. Optical density ($OD_{600}$, 600 nm absorbance) and fluorescence were measured every 15 min for 18 h using 465/35 nm excitation and 530/25 nm emission for EGFP, and 580/20 nm excitation and 635/35 nm emission for mCherry.
3. A second aliquot of the M0 co-culture was diluted 1:100 into M9 medium containing the appropriate concentration of kanamycin (as listed in the protocol) and inducer for learning conditions, or K0 for memory-stability conditions. Cultures were grown for 8 h to reach logarithmic phase. After 8 h, glycerol stocks were prepared and designated **M1 (generation 1)**.
4. Back-dilute into M9 + carbenicillin medium and grow to mid-log phase ($OD_{600} \approx 0.4$–0.6).
5. Combine strains in equal volumes to assemble the desired co-cultures per board position.
6. Prepare Day 0 glycerol stocks for each co-culture and relevant monocultures (final glycerol 25%) and store at −80°C.

Subsequent sections describe the daily workflow of the tournament: revival of co-cultures from glycerol stocks, plating in microtiter plates, induction with the appropriate inducers to represent the opponent's moves, implementation of the supervised adaptation sequence using kanamycin, and sampling for fluorescence and DNA measurements. Standard media recipes (LB, M9, antibiotic stocks) and plate-reader characterisation protocols are provided as Appendices A and B.

| Board Position | Required Memregulon Strains (Promoter Name - M-stage) |
|---|---|
| Pos 1 | PLux - M0 |
| Pos 2 | PTac - M1, PTtg - M1, PVan - M1 |
| Pos 3 | PTtg - M0, PVan - M0 |
| Pos 4 | PCin - M0, PLux - M0 |
| Pos 6 | PBAD - M0, PSal - M0, PVan - M0 |
| Pos 7 | PBetI - M0, PTac - M0 |
| Pos 8 | PBetI - M1, PCin - M1, PTet - M1 |

| | |
|---|---|
| Pos 9 | PSal - M0, PTet - M0 |

**Supplementary Table 38 | Initial co-culture composition (Day 0).** Equal to the starting configuration of this simulation run.

### PROCEDURE

#### Preparation of Initial Co-cultures (Day 0)

(Estimated time: ~48 hours for steps 1–5)

1. Inoculate individual memregulon strains required for the Supplementary Table 38 compositions from −80 °C glycerol stocks into LB + carbenicillin (80 µg/mL). Grow overnight (~16–18 h) at 37 °C with vigorous shaking (for example, 200 rpm).
2. Back-dilute overnight cultures 1:100 (or an appropriate ratio based on growth) into fresh, pre-warmed M9 + carbenicillin (80 µg/mL) medium in tubes. Grow at 37 °C with shaking (~200 rpm) until cultures reach mid-log phase ($OD_{600}$ ~ 0.4–0.6). Monitor OD frequently.
3. Prepare Day 0 co-cultures. For each board position (1–4, 6–9) requiring a co-culture, combine equal volumes of the specified mid-log phase individual strain cultures into a sterile tube or flask and mix gently. For positions requiring a monoculture, use the corresponding mid-log individual strain culture directly.
4. Prepare Day 0 glycerol stocks. For each prepared co-culture and necessary monocultures (positions 1–4, 6–9), mix 750 µL of the mid-log (co-)culture with 750 µL of sterile 50% glycerol in a labelled cryovial (final glycerol 25%). Label clearly (for example, "Pos 1 – Day 0 co-culture", "Pos 3 – Day 0 mono P7-M0"). Prepare at least 3–4 vials per co-culture. These vials are the starting stocks for the experiment.

Subsequent sections describe the daily workflow of the tournament: revival of co-cultures from glycerol stocks, plating in microtitre plates, induction with the appropriate inducers to represent the opponent's moves, implementation of the supervised adaptation sequence using kanamycin and sampling for fluorescence and DNA measurements. Standard media recipes (LB, M9, antibiotic stocks) and plate-reader characterisation protocols are provided in Appendices A and B. Inducer and antibiotic concentrations used at each step are stated directly in the protocol steps below.

#### Daily Learning & Characterization Cycles (Day 1 to Day 7)

Each day involves the learning/incubation step for the current generation, followed by characterization of the *previous* generation's state.

##### Day 1: Learn/Memory Generation 0 / Characterize Generation 0

(Learning based on Simulated Match 3 Loss)

Reference Simulation Match 3 (Leads to Day 1 Learning):

| | | |
|---|---|---|
| $X_9$ | $O_6$ | $X_3$ |
| $O_2$ | $X_1$ | $O_8$ |
| $O_4$ | $X_7$ | $X_5$ |

**Supplementary Table 39 | Reference Simulation Match 3 (Leads to Day 1 Learning).**

- Simulated Losing O Moves Sequence: [4, 7, 2, 6]
- Triggering X Moves Sequence (Inducers): [5, 3, 9, 8, 1]

##### *Day 1 Activity 1: Learn/Memory Generation 0 (Produce Generation 1)*

Objective: Apply sublethal kanamycin (Learning) to targeted Generation 0 cultures based on Match 3 AND perform parallel MEMORY INCUBATION without kanamycin on non-targeted cultures to produce the Generation 1 set. This step applies adaptation pressure or allows observation of culture stability ('memory') under identical growth conditions.
Learning Targets (using Day 0 Stocks): Position(s) 2, 4, 6, 7
Memory Incubation Targets (using Day 0 Stocks): Position(s) 1, 3, 8, 9
**Kanamycin Learning & Memory Incubation Details: Learning time is 8.0 hours 100 times dilution overnight culture.**

⇒ LEARNING tube for **Pos 6** *from Gen 0* (containing culture **PBAD-M0+PSal-M0+PVan-M0**): Add **Inducer 8 (Van) at 100 µM + Kan at 3.5 µg/mL.** This condition aims to produce the culture **PBAD-M0+PSal-M0+PVan-M0-*K*_Van**, *corresponding to Gen 1.*
⇒ LEARNING tube for **Pos 4** *from Gen 0* (containing culture **PCin-M0+PLux-M0**): Add **Inducer 5 (OC6) at 10 µM + Kan at 1.0 µg/mL.** This condition aims to produce the culture **PCin-M0+PLux-M0-*K*_OC6**, *corresponding to Gen 1.*
⇒ LEARNING tube for **Pos 7** *from Gen 0* (containing culture **PBetI-M0+PTac-M0**): Add **Inducer 3 (Cho) at 10 mM + Kan at 4.0 µg/mL.** This condition aims to produce the culture **PBetI-M0+PTac-M0-*K*_Cho**, *corresponding to Gen 1.*
⇒ LEARNING tube for **Pos 2** *from Gen 0* (containing culture **PTac-M1+PTtg-M1+PVan-M1**): Add **Inducer 9 (IPTG) at 500 µM + Kan at 4.0 µg/mL.** This condition aims to produce the culture **PTac-M1+PTtg-M1+PVan-M1-*K*_IPTG**, *corresponding to Gen 1.*
⇒ MEMORY INCUBATION tube for **Pos 1** *from Gen 0* (containing culture **PLux-M0**): Add only solvent control (**NO Inducer, NO Kanamycin**) This condition aims to produce the culture **PLux-M0-M**, *corresponding to Gen 1.*
⇒ MEMORY INCUBATION tube for **Pos 3** *from Gen 0* (containing culture **PTtg-M0+PVan-M0**): Add only solvent control (**NO Inducer, NO Kanamycin**) This condition aims to produce the culture **PTtg-M0+PVan-M0-M**, *corresponding to Gen 1.*
⇒ MEMORY INCUBATION tube for **Pos 8** *from Gen 0* (containing culture **PBetI-M1+PCin-M1+PTet-M1**): Add only solvent control (**NO Inducer, NO Kanamycin**) This condition aims to produce the culture **PBetI-M1+PCin-M1+PTet-M1-M**, *corresponding to Gen 1.*
⇒ MEMORY INCUBATION tube for **Pos 9** *from Gen 0* (containing culture **PSal-M0+PTet-M0**): Add only solvent control (**NO Inducer, NO Kanamycin**) This condition aims to produce the culture **PSal-M0+PTet-M0-M**, *corresponding to Gen 1.*

Detailed Learning/Memory Procedure (Day 1):
**1. Culture Preparation:** From -80°C, thaw one glycerol stock vial for each Day 0 co-culture (positions 1-4, 6-9).
**2. Refresh Cultures:** Inoculate thawed Day 0 stocks into M9+Cb (80 µg/mL) and grow to mid-log phase OR perform overnight growth as needed.
**3. Prepare Learning/Memory tube:** Using the refreshed/mid-log Day 0 cultures, prepare replicate tube for LEARNING positions (2, 4, 6, 7) and MEMORY INCUBATION tube for other positions (1, 3, 8, 9) in M9+Cb (80 µg/mL).
**4. Add Reagents & Incubate:** To LEARNING tubes, add specific Inducer(s)/Kanamycin as detailed above. To MEMORY INCUBATION tubes, add only solvent controls. Incubate ALL

tubes (Learning and Incubation) for 8 hours at 37°C with shaking.
**5. Harvest and Store Generation Stocks:** After 8 hours, prepare glycerol stocks from ALL tubes. These represent the Generation 1 state. Label clearly (e.g., 'Pos X - Gen 1 - PostSel Rep A', 'Pos Y - Gen 1 - Memory Ctrl'), flash-freeze, and store at -80°C. These stocks will be used on Day 2.

##### *Day 1 Activity 2: Characterize Generation 0*

Objective: Characterize the response of *all 8 active* Generation 0 co-cultures to *all* relevant CHARACTERIZATION inducers (where the cognate promoter is present). ALL relevant combinations will be measured.
Using one biological replicate and four technical replicates for experiments and 4 for controls for Day 1 Characterization.
Procedure (Characterization - Day 1 Evening/Overnight): Requires 1 plate(s).
Follow standard characterization procedure (Appendix B) using the 'Generation 0' glycerol stocks (produced Day 0). Prepare cultures and add inducers at CHARACTERIZATION concentrations as specified below. Incubate in plate reader overnight.

###### Characterization Plate 1 (Day 1 Characterization (Gen 0))

Expected State after Day 1 Learning/Memory Incubation (Generation 1):

| Position | Promoter | Expected Change (Weight/M-stage) |
|---|---|---|
| 7 | PBetI | 0.813 (M0) -> 0.730 (M1) |
| 4 | PLux | 0.955 (M0) -> 0.670 (M1) |
| 6 | PVan | 0.836 (M0) -> 0.872 (M1) |
| 2 | PTac | 0.804 (M1) -> 0.715 (M2) |

**Supplementary Table 40 | Characterization Plate 1 (Day 1 Characterization (Gen 0)).** Expected State after Day 1 Learning/Memory Incubation (Generation 1).

#### Day 2: Learn/Memory Generation 1 / Characterize Generation 1

(Learning based on Simulated Match 10 Loss)
Reference Simulation Match 10 (Leads to Day 2 Learning):

| | | |
|---|---|---|
| $O_2$ | $X_7$ | $X_5$ |
| $X_3$ | $X_1$ | $O_4$ |
| $X_9$ | $O_6$ | $O_8$ |

**Supplementary Table 41 | Reference Simulation Match 10 (Leads to Day 2 Learning).**

- Simulated Losing O Moves Sequence: [1, 6, 8, 9]
- Triggering X Moves Sequence (Inducers): [5, 4, 3, 2, 7]

##### *Day 2 Activity 1: Learn/Memory Generation 1 (Produce Generation 2)*

Objective: Apply sublethal kanamycin (Learning) to targeted Generation 1 cultures based on Match 10 AND perform parallel MEMORY INCUBATION without kanamycin on non-targeted cultures to produce the Generation 2 set. This step applies adaptation pressure or allows observation of culture stability ('memory') under identical growth conditions.
Learning Targets (using Day 1 Stocks): Position(s) 1, 6, 8, 9

Memory Incubation Targets (using Day 1 Stocks): Position(s) 2, 3, 4, 7

**Kanamycin Learning & Memory Incubation Details: Learning time is 8.0 hours 100 times dilution overnight culture.**

⇒ LEARNING tube for **Pos 8** *from Gen 1* (containing culture **PBetI-M1+PCin-M1+PTet-M1**): Add **Inducer 3 (Cho) at 10 mM + Kan at 4.0 µg/mL.** This condition aims to produce the culture **PBetI-M1+PCin-M1+PTet-M1-M-*K*_Cho**, *corresponding to Gen 2.*

⇒ LEARNING tube for **Pos 1** *from Gen 1* (containing culture **PLux-M0**): Add **Inducer 5 (OC6) at 10 µM + Kan at 1.0 µg/mL.** This condition aims to produce the culture **PLux-M0-M-*K*_OC6**, *corresponding to Gen 2.*

⇒ LEARNING tube for **Pos 6** *from Gen 1* (containing culture **PBAD-M0+PSal-M0+PVan-M0**): Add **Inducer 4 (Ara) at 4 mM + Kan at 2.5 µg/mL.** This condition aims to produce the culture **PBAD-M0+PSal-M0+PVan-M0-*K*_Van-*K*_Ara**, *corresponding to Gen 2.*

⇒ LEARNING tube for **Pos 9** *from Gen 1* (containing culture **PSal-M0+PTet-M0**): Add **Inducer 2 (aTc) at 0.2 µM + Kan at 2.5 µg/mL.** This condition aims to produce the culture **PSal-M0+PTet-M0-M-*K*_aTc**, *corresponding to Gen 2.*

⇒ MEMORY INCUBATION tube for **Pos 2** *from Gen 1* (containing culture **PTac-M1+PTtg-M1+PVan-M1**): Add only solvent control (**NO Inducer, NO Kanamycin**) This condition aims to produce the culture **PTac-M1+PTtg-M1+PVan-M1-*K*_IPTG-M**, *corresponding to Gen 2.*

⇒ MEMORY INCUBATION tube for **Pos 3** *from Gen 1* (containing culture **PTtg-M0+PVan-M0**): Add only solvent control (**NO Inducer, NO Kanamycin**) This condition aims to produce the culture **PTtg-M0+PVan-M0-M-M**, *corresponding to Gen 2.*

⇒ MEMORY INCUBATION tube for **Pos 4** *from Gen 1* (containing culture **PCin-M0+PLux-M0**): Add only solvent control (**NO Inducer, NO Kanamycin**) This condition aims to produce the culture **PCin-M0+PLux-M0-*K*_OC6-M**, *corresponding to Gen 2.*

⇒ MEMORY INCUBATION tube for **Pos 7** *from Gen 1* (containing culture **PBetI-M0+PTac-M0**): Add only solvent control (**NO Inducer, NO Kanamycin**) This condition aims to produce the culture **PBetI-M0+PTac-M0-*K*_Cho-M**, *corresponding to Gen 2.*

Detailed Learning/Memory Procedure (Day 2):

1. Culture Preparation: From -80°C, thaw one glycerol stock vial for each Day 1 co-culture (positions 1-4, 6-9).
2. Refresh Cultures: Inoculate thawed Day 1 stocks into M9+Cb (80 µg/mL) and grow to mid-log phase OR perform overnight growth as needed.
3. Prepare Learning/Memory tubes: Using the refreshed/mid-log Day 1 cultures, prepare replicate tubes for LEARNING positions (1, 6, 8, 9) and MEMORY INCUBATION tubes for other positions (2, 3, 4, 7) in M9+Cb (80 µg/mL).
4. Add Reagents & Incubate: To LEARNING tubes, add specific Inducer(s)/Kanamycin as detailed above. To MEMORY INCUBATION tubes, add only solvent controls. Incubate all tubes (learning and memory-incubation conditions) for 8 hours at 37°C with shaking.
5. Harvest and Store Generation Stocks: After 8 hours, prepare glycerol stocks from ALL tubes. These represent the Generation 2 state. Label clearly (e.g., 'Pos X - Gen 2 - PostSel Rep A', 'Pos Y - Gen 2 - Memory Ctrl'), flash-freeze, and store at -80°C. These stocks will be used on Day 3.

##### *Day 2 Activity 2: Characterize Generation 1*

Objective: Characterize the response of *all 8 active* Generation 1 co-cultures to *all* relevant CHARACTERIZATION inducers (where the cognate promoter is present). ALL relevant combinations will be measured.

Using one biological replicate and four technical replicates for experiments and 4 for controls for Day 2 Characterization.
Procedure (Characterization - Day 2 Evening/Overnight): Requires 1 plate(s).
Follow standard characterization procedure (Appendix B) using the 'Generation 1' glycerol stocks (produced Day 1). Prepare cultures and add inducers at CHARACTERIZATION concentrations as specified below. Incubate in plate reader overnight.

##### Characterization Plate 1 (Day 2 Characterization (Gen 1))

Expected State after Day 2 Learning/Memory Incubation (Generation 2):

| Position | Promoter | Expected Change (Weight/M-stage) |
|---|---|---|
| 9 | PTet | 0.664 (M0) -> 0.665 (M1) |
| 8 | PBetI | 0.730 (M1) -> 0.633 (M2) |
| 6 | PBAD | 0.710 (M0) -> 0.680 (M1) |
| 1 | PLux | 0.955 (M0) -> 0.670 (M1) |

**Supplementary Table 42 | Characterization Plate 1 (Day 2 Characterization (Gen 1)).** Expected State after Day 2 Learning/Memory Incubation (Generation 2).

#### Day 3: Learn/Memory Generation 2 / Characterize Generation 2

(Learning based on Simulated Match 12 Loss)
Reference Simulation Match 12 (Leads to Day 3 Learning):

| | | |
|---|---|---|
| $X_9$ | $O_6$ | $X_3$ |
| $O_2$ | $X_1$ | $O_8$ |
| $O_4$ | $X_7$ | $X_5$ |

**Supplementary Table 43 | Reference Simulation Match 12 (Leads to Day 3 Learning).**

- Simulated Losing O Moves Sequence: [4, 7, 2, 6]
- Triggering X Moves Sequence (Inducers): [5, 3, 9, 8, 1]

##### *Day 3 Activity 1: Learn/Memory Generation 2 (Produce Generation 3)*

Objective: Apply sublethal kanamycin (Learning) to targeted Generation 2 cultures based on Match 12 AND perform parallel MEMORY INCUBATION without kanamycin on non-targeted cultures to produce the Generation 3 set. This step applies adaptation pressure or allows observation of culture stability ('memory') under identical growth conditions.
Learning Targets (using Day 2 Stocks): Position(s) 2, 4, 6, 7
Memory Incubation Targets (using Day 2 Stocks): Position(s) 1, 3, 8, 9
**Kanamycin Learning & Memory Incubation Details: Learning time is 4.50 hours**
**50 times dilution overnight culture.**

⇒ LEARNING tube for **Pos 6** *from Gen 2* (containing culture **PBAD-M0+PSal-M0+PVan-M0**): Add **Inducer 8 (Van) at 100 µM + Kan at 1.75 µg/mL.** This condition aims to produce the culture **PBAD-M0+PSal-M0+PVan-M0-*K*_Van-*K*_Ara-*K*_Van**, *corresponding to Gen 3.*

⇒ LEARNING tube for **Pos 4** *from Gen 2* (containing culture **PCin-M0+PLux-M0**): Add **Inducer 5 (OC6) at 10 µM + Kan at 0.50 µg/mL.** This condition aims to produce the culture **PCin-M0+PLux-M0-*K*_OC6-M-*K*_OC6**, *corresponding to Gen 3.*

⇒ LEARNING tube for **Pos 7** *from Gen 2* (containing culture **PBetI-M0+PTac-M0**): Add **Inducer 3 (Cho) at 10 mM + Kan at 2.0 µg/mL.** This condition aims to produce the culture **PBetI-M0+PTac-M0-*K*_Cho-M-*K*_Cho**, *corresponding to Gen 3.*

⇒ LEARNING tube for **Pos 2** *from Gen 2* (containing culture **PTac-M1+PTtg-M1+PVan-M1**): Add **Inducer 9 (IPTG) at 500 µM + Kan at 2.0 µg/mL.** This condition aims to produce the culture **PTac-M1+PTtg-M1+PVan-M1-*K*_IPTG-M-*K*_IPTG**, *corresponding to Gen 3.*

⇒ MEMORY INCUBATION tube for **Pos 1** *from Gen 2* (containing culture **PLux-M0**): Add only solvent control (**NO Inducer, NO Kanamycin**) This condition aims to produce the culture **PLux-M0-M-*K*_OC6-M**, *corresponding to Gen 3.*

⇒ MEMORY INCUBATION tube for **Pos 3** *from Gen 2* (containing culture **PTtg-M0+PVan-M0**): Add only solvent control (**NO Inducer, NO Kanamycin**) This condition aims to produce the culture **PTtg-M0+PVan-M0-M-M-M**, *corresponding to Gen 3.*

⇒ MEMORY INCUBATION tube for **Pos 8** *from Gen 2* (containing culture **PBetI-M1+PCin-M1+PTet-M1**): Add only solvent control (**NO Inducer, NO Kanamycin**) This condition aims to produce the culture **PBetI-M1+PCin-M1+PTet-M1-M-*K*_Cho-M**, *corresponding to Gen 3.*

⇒ MEMORY INCUBATION tube for **Pos 9** *from Gen 2* (containing culture **PSal-M0+PTet-M0**): Add only solvent control (**NO Inducer, NO Kanamycin**) This condition aims to produce the culture **PSal-M0+PTet-M0-M-*K*_aTc-M**, *corresponding to Gen 3.*

Detailed Learning/Memory Procedure (Day 3):
1. Culture Preparation: From -80°C, thaw one glycerol stock vial for each Day 2 co-culture (positions 1-4, 6-9).
2. Refresh Cultures: Inoculate thawed Day 2 stocks into M9+Cb (100 µg/mL) and grow to mid-log phase OR perform overnight growth as needed.
3. Prepare Learning/Memory tubes: Using the refreshed/mid-log Day 2 cultures, prepare replicate flasks for LEARNING positions (2, 4, 6, 7) and MEMORY INCUBATION tubes for other positions (1, 3, 8, 9) in M9+Cb (80 µg/mL).
4. Add Reagents & Incubate: To LEARNING tubes, add specific Inducer(s)/Kanamycin as detailed above. To MEMORY INCUBATION tubes, add only solvent controls. Incubate all tubes (learning and memory-incubation conditions) for 8 hours at 37°C with shaking.
5. Harvest and Store Generation Stocks: After 8 hours, prepare glycerol stocks from ALL tubes. These represent the Generation 3 state. Label clearly (e.g., 'Pos X - Gen 3 - PostSel Rep A', 'Pos Y - Gen 3 - Memory Ctrl'), flash-freeze, and store at -80°C. These stocks will be used on Day 4.

##### *Day 3 Activity 2: Characterize Generation 2*

Objective: Characterize the response of *all 8 active* Generation 2 co-cultures to *all* relevant CHARACTERIZATION inducers (where the cognate promoter is present). ALL relevant combinations will be measured.
Using one biological replicate and four technical replicates for experiments and 4 for controls for Day 3 Characterization.
Procedure (Characterization - Day 3 Evening/Overnight): Requires 1 plate(s).
Follow standard characterization procedure (Appendix B) using the 'Generation 2' glycerol stocks (produced Day 2). Prepare cultures and add inducers at CHARACTERIZATION concentrations as specified below. Incubate in plate reader overnight.

Characterization Plate 1 (Day 3 Characterization (Gen 2))
Expected State after Day 3 Learning/Memory Incubation (Generation 3):

| Position | Promoter | Expected Change (Weight/M-stage) |
|---|---|---|
| 7 | PBetI | 0.730 (M1) -> 0.633 (M2) |
| 4 | PLux | 0.670 (M1) -> 0.543 (M2) |
| 6 | PVan | 0.872 (M1) -> 0.868 (M2) |
| 2 | PTac | 0.715 (M2) -> 0.198 (M3) |

**Supplementary Table 44 | Characterization Plate 1 (Day 3 Characterization (Gen 2)).** Expected State after Day 3 Learning/Memory Incubation (Generation 3).

#### Day 4: Learn/Memory Generation 3 / Characterize Generation 3

(Learning based on Simulated Match 20 Loss)
Reference Simulation Match 20 (Leads to Day 4 Learning):

| | | |
|---|---|---|
| $O_2$ | $X_7$ | $X_5$ |
| $X_3$ | $X_1$ | $O_4$ |
| $X_9$ | $O_6$ | $O_8$ |

**Supplementary Table 45 | Reference Simulation Match 20 (Leads to Day 4 Learning).**

- Simulated Losing O Moves Sequence: [1, 6, 8, 9]
- Triggering X Moves Sequence (Inducers): [5, 4, 3, 2, 7]

##### *Day 4 Activity 1: Learn/Memory Generation 3 (Produce Generation 4)*

Objective: Apply sublethal kanamycin (Learning) to targeted Generation 3 cultures based on Match 20 AND perform parallel MEMORY INCUBATION without kanamycin on non-targeted cultures to produce the Generation 4 set. This step applies adaptation pressure or allows observation of culture stability ('memory') under identical growth conditions.
Learning Targets (using Day 3 Stocks): Position(s) 1, 6, 8, 9
Memory Incubation Targets (using Day 3 Stocks): Position(s) 2, 3, 4, 7
**Kanamycin Learning & Memory Incubation Details: Learning time is 4.50 hours 50 times dilution overnight culture.**

⇒ LEARNING tube for **Pos 8** *from Gen 3* (containing culture **PBetI-M1+PCin-M1+PTet-M1**): Add **Inducer 3 (Cho) at 10 mM + Kan at 2.0 µg/mL.** This condition aims to produce the culture **PBetI-M1+PCin-M1+PTet-M1-M-*K*_Cho-M-*K*_Cho**, *corresponding to Gen 4.*

⇒ LEARNING tube for **Pos 1** *from Gen 3* (containing culture **PLux-M0**): Add **Inducer 5 (OC6) at 10 µM + Kan at 0.50 µg/mL.** This condition aims to produce the culture **PLux-M0-M-*K*_OC6-M-*K*_OC6**, *corresponding to Gen 4.*

⇒ LEARNING tube for **Pos 6** *from Gen 3* (containing culture **PBAD-M0+PSal-M0+PVan-M0**): Add **Inducer 4 (Ara) at 4 mM + Kan at 1.25 µg/mL.** This condition aims to produce the culture **PBAD-M0+PSal-M0+PVan-M0-*K*_Van-*K*_Ara-*K*_Van-*K*_Ara**, *corresponding to Gen 4.*

⇒ LEARNING tube for **Pos 9** *from Gen 3* (containing culture **PSal-M0+PTet-M0**): Add **Inducer 2 (aTc) at 0.2 µM + Kan at 1.25 µg/mL.** This condition aims to produce the culture **PSal-M0+PTet-M0-M-*K*_aTc-M-*K*_aTc**, *corresponding to Gen 4.*
⇒ MEMORY INCUBATION tube for **Pos 2** *from Gen 3* (containing culture **PTac-M1+PTtg-M1+PVan-M1**): Add only solvent control (**NO Inducer, NO Kanamycin**) This condition aims to produce the culture **PTac-M1+PTtg-M1+PVan-M1-*K*_IPTG-M-*K*_IPTG-M**, *corresponding to Gen 4.*
⇒ MEMORY INCUBATION tube for **Pos 3** *from Gen 3* (containing culture **PTtg-M0+PVan-M0**): Add only solvent control (**NO Inducer, NO Kanamycin**) This condition aims to produce the culture **PTtg-M0+PVan-M0-M-M-M-M**, *corresponding to Gen 4.*
⇒ MEMORY INCUBATION tube for **Pos 4** *from Gen 3* (containing culture **PCin-M0+PLux-M0**): Add only solvent control (**NO Inducer, NO Kanamycin**) This condition aims to produce the culture **PCin-M0+PLux-M0-*K*_OC6-M-*K*_OC6-M**, *corresponding to Gen 4.*
⇒ MEMORY INCUBATION tube for **Pos 7** *from Gen 3* (containing culture **PBetI-M0+PTac-M0**): Add only solvent control (**NO Inducer, NO Kanamycin**) This condition aims to produce the culture **PBetI-M0+PTac-M0-*K*_Cho-M-*K*_Cho-M**, *corresponding to Gen 4.*

Detailed Learning/Memory Procedure (Day 4):
**1. Culture Preparation:** From -80°C, thaw one glycerol stock vial for each Day 3 co-culture (positions 1-4, 6-9).
**2. Refresh Cultures:** Inoculate thawed Day 3 stocks into M9+Cb (100 µg/mL) and grow to mid-log phase OR perform overnight growth as needed.
**3. Prepare Learning/Memory tubes:** Using the refreshed/mid-log Day 3 cultures, prepare replicate tubes for LEARNING positions (1, 6, 8, 9) and MEMORY INCUBATION tubes for other positions (2, 3, 4, 7) in M9+Cb (80 µg/mL).
**4. Add Reagents & Incubate:** To LEARNING tubes, add specific Inducer(s)/Kanamycin as detailed above. To MEMORY INCUBATION tubes, add only solvent controls. Incubate ALL tubes (Learning and Incubation) for 8 hours at 37°C with shaking.
**5. Harvest and Store Generation Stocks:** After 8 hours, prepare glycerol stocks from ALL tubes. These represent the Generation 4 state. Label clearly (e.g., 'Pos X - Gen 4 - PostSel Rep A', 'Pos Y - Gen 4 - Memory Ctrl'), flash-freeze, and store at -80°C. These stocks will be used on Day 5.

##### *Day 4 Activity 2: Characterize Generation 3*

Objective: Characterize the response of *all 8 active* Generation 3 co-cultures to *all* relevant CHARACTERIZATION inducers (where the cognate promoter is present). ALL relevant combinations will be measured.
Using one biological replicate and four technical replicates for experiments and 4 for controls for Day 4 Characterization.
Procedure (Characterization - Day 4 Evening/Overnight): Requires 1 plate(s).
Follow standard characterization procedure (Appendix B) using the 'Generation 3' glycerol stocks (produced Day 3). Prepare cultures and add inducers at CHARACTERIZATION concentrations as specified below. Incubate in plate reader overnight.

##### Characterization Plate 1 (Day 4 Characterization (Gen 3))

Expected State after Day 4 Learning/Memory Incubation (Generation 4):

| Position | Promoter | Expected Change (Weight/M-stage) |
|---|---|---|
| 9 | PTet | 0.665 (M1) -> 0.614 (M2) |
| 8 | PBetI | 0.633 (M2) -> 0.626 (M3) |
| 6 | PBAD | 0.680 (M1) -> 0.615 (M2) |
| 1 | PLux | 0.670 (M1) -> 0.543 (M2) |

**Supplementary Table 46 | Characterization Plate 1 (Day 4 Characterization (Gen 3)).** Expected State after Day 4 Learning/Memory Incubation (Generation 4).

#### Day 5: Learn/Memory Generation 4 / Characterize Generation 4

(Learning based on Simulated Match 22 Loss)
Reference Simulation Match 22 (Leads to Day 5 Learning):

| | | |
|---|---|---|
| $X_9$ | $O_6$ | $X_3$ |
| $O_2$ | $X_1$ | $O_8$ |
| $O_4$ | $X_7$ | $X_5$ |

**Supplementary Table 47 | Reference Simulation Match 22 (Leads to Day 5 Learning).**

- Simulated Losing O Moves Sequence: [4, 7, 2, 6]
- Triggering X Moves Sequence (Inducers): [5, 3, 9, 8, 1]

##### *Day 5 Activity 1: Learn/Memory Generation 4 (Produce Generation 5)*

Objective: Apply sublethal kanamycin (Learning) to targeted Generation 4 cultures based on Match 22 AND perform parallel MEMORY INCUBATION without kanamycin on non-targeted cultures to produce the Generation 5 set. This step applies adaptation pressure or allows observation of culture stability ('memory') under identical growth conditions.
Learning Targets (using Day 4 Stocks): Position(s) 2, 4, 6, 7
Memory Incubation Targets (using Day 4 Stocks): Position(s) 1, 3, 8, 9
**Kanamycin Learning & Memory Incubation Details: Learning time is 4.50 hours 50 times dilution overnight culture.**

⇒ LEARNING tube for **Pos 6** *from Gen 4* (containing culture **PBAD-M0+PSal-M0+PVan-M0**): Add **Inducer 8 (Van) at 100 µM + Kan at 1.75 µg/mL.** This condition aims to produce the culture **PBAD-M0+PSal-M0+PVan-M0-*K*_Van-*K*_Ara-*K*_Van-*K*_Ara-*K*_Van**, *corresponding to Gen 5.*

⇒ LEARNING tube for **Pos 4** *from Gen 4* (containing culture **PCin-M0+PLux-M0**): Add **Inducer 5 (OC6) at 10 µM + Kan at 0.50 µg/mL.** This condition aims to produce the culture **PCin-M0+PLux-M0-*K*_OC6-M-*K*_OC6-M-*K*_OC6**, *corresponding to Gen 5.*

⇒ LEARNING tube for **Pos 7** *from Gen 4* (containing culture **PBetI-M0+PTac-M0**): Add **Inducer 3 (Cho) at 10 mM + Kan at 2.0 µg/mL.** This condition aims to produce the culture **PBetI-M0+PTac-M0-*K*_Cho-M-*K*_Cho-M-*K*_Cho**, *corresponding to Gen 5.*

⇒ LEARNING tube for **Pos 2** *from Gen 4* (containing culture **PTac-M1+PTtg-M1+PVan-M1**): Add **Inducer 9 (IPTG) at 500 µM + Kan at 2.0 µg/mL.** This condition aims to produce the culture **PTac-M1+PTtg-M1+PVan-M1-*K*_IPTG-M-*K*_IPTG-M-*K*_IPTG**, *corresponding to Gen 5.*

⇒ MEMORY INCUBATION tube for **Pos 1** *from Gen 4* (containing culture **PLux-M0**): Add only solvent control (**NO Inducer, NO Kanamycin**) This condition aims to produce the culture **PLux-M0-M-*K*_OC6-M-*K*_OC6-M**, *corresponding to Gen 5.*

⇒ MEMORY INCUBATION tube for **Pos 3** *from Gen 4* (containing culture **PTtg-M0+PVan-M0**): Add only solvent control (**NO Inducer, NO Kanamycin**) This condition aims to produce the culture **PTtg-M0+PVan-M0-M-M-M-M-M**, *corresponding to Gen 5.*

⇒ MEMORY INCUBATION tube for **Pos 8** *from Gen 4* (containing culture **PBetI-M1+PCin-M1+PTet-M1**): Add only solvent control (**NO Inducer, NO Kanamycin**) This condition aims to produce the culture **PBetI-M1+PCin-M1+PTet-M1-M-*K*_Cho-M-*K*_Cho-M**, *corresponding to Gen 5.*

⇒ MEMORY INCUBATION tube for **Pos 9** *from Gen 4* (containing culture **PSal-M0+PTet-M0**): Add only solvent control (**NO Inducer, NO Kanamycin**) This condition aims to produce the culture **PSal-M0+PTet-M0-M-*K*_aTc-M-*K*_aTc-M**, *corresponding to Gen 5.*

Detailed Learning/Memory Procedure (Day 5):
**1. Culture Preparation:** From -80°C, thaw one glycerol stock vial for each Day 4 co-culture (positions 1-4, 6-9).
**2. Refresh Cultures:** Inoculate thawed Day 4 stocks into M9+Cb (100 µg/mL) and grow to mid-log phase OR perform overnight growth as needed.
**3. Prepare Learning/Memory tubes:** Using the refreshed/mid-log Day 4 cultures, prepare replicate flasks for LEARNING positions (2, 4, 6, 7) and MEMORY INCUBATION flasks for other positions (1, 3, 8, 9) in M9+Cb (80 µg/mL).
**4. Add Reagents & Incubate:** To LEARNING tubes, add specific Inducer(s)/Kanamycin as detailed above. To MEMORY INCUBATION tubes, add only solvent controls. Incubate ALL tubes (Learning and Incubation) for 8 hours at 37°C with shaking.
**5. Harvest and Store Generation Stocks:** After 8 hours, prepare glycerol stocks from ALL tubes. These represent the Generation 5 state. Label clearly (e.g., 'Pos X - Gen 5 - PostSel Rep A', 'Pos Y - Gen 5 - Memory Ctrl'), flash-freeze, and store at -80°C. These stocks will be used on Day 6.

##### *Day 5 Activity 2: Characterize Generation 4*

Objective: Characterize the response of *all 8 active* Generation 4 co-cultures to *all* relevant CHARACTERIZATION inducers (where the cognate promoter is present). ALL relevant combinations will be measured.
Using one biological replicate and four technical replicates for experiments and 4 for controls for Day 5 Characterization.
Procedure (Characterization - Day 5 Evening/Overnight): Requires 1 plate(s).
Follow standard characterization procedure (Appendix B) using the 'Generation 4' glycerol stocks (produced Day 4). Prepare cultures and add inducers at CHARACTERIZATION concentrations as specified below. Incubate in plate reader overnight.

##### Characterization Plate 1 (Day 5 Characterization (Gen 4))

Expected State after Day 5 Learning/Memory Incubation (Generation 5):

| Position | Promoter | Expected Change (Weight/M-stage) |
|---|---|---|
| 7 | PBetI | 0.633 (M2) -> 0.626 (M3) |
| 4 | PLux | 0.543 (M2) -> 0.505 (M3) |
| 6 | PVan | 0.868 (M2) -> 0.672 (M3) |
| 2 | PTac | 0.198 (M3) -> 0.003 (M4) |

**Supplementary Table 48 | Characterization Plate 1 (Day 5 Characterization (Gen 4)).** Expected State after Day 5 Learning/Memory Incubation (Generation 5).

#### Day 6: Learn/Memory Generation 5 / Characterize Generation 5

(Learning based on Simulated Match 26 Loss)
Reference Simulation Match 26 (Leads to Day 6 Learning):

| | | |
|---|---|---|
| $O_2$ | $X_7$ | $X_5$ |
| $X_3$ | $X_1$ | $O_4$ |
| $X_9$ | $O_6$ | $O_8$ |

**Supplementary Table 49 | Reference Simulation Match 26 (Leads to Day 6 Learning).**

- Simulated Losing O Moves Sequence: [1, 6, 8, 9]
- Triggering X Moves Sequence (Inducers): [5, 4, 3, 2, 7]

##### *Day 6 Activity 1: Learn/Memory Generation 5 (Produce Generation 6)*

Objective: Apply sublethal kanamycin (Learning) to targeted Generation 5 cultures based on Match 26 AND perform parallel MEMORY INCUBATION without kanamycin on non-targeted cultures to produce the Generation 6 set. This step applies adaptation pressure or allows observation of culture stability ('memory') under identical growth conditions.
Learning Targets (using Day 5 Stocks): Position(s) 1, 6, 8, 9
Memory Incubation Targets (using Day 5 Stocks): Position(s) 2, 3, 4, 7
**Kanamycin Learning & Memory Incubation Details: Learning time is 4.50 hours 50 times dilution overnight culture.**

⇒ LEARNING tube for **Pos 8** *from Gen 5* (containing culture **PBetI-M1+PCin-M1+PTet-M1**): Add **Inducer 3 (Cho) at 10 mM + Kan at 2.0 µg/mL.** This condition aims to produce the culture **PBetI-M1+PCin-M1+PTet-M1-M-*K*_Cho-M-*K*_Cho-M-*K*_Cho**, *corresponding to Gen 6.*

⇒ LEARNING tube for **Pos 1** *from Gen 5* (containing culture **PLux-M0**): Add **Inducer 5 (OC6) at 10 µM + Kan at 0.50 µg/mL.** This condition aims to produce the culture **PLux-M0-M-*K*_OC6-M-*K*_OC6-M-*K*_OC6**, *corresponding to Gen 6.*

⇒ LEARNING tube for **Pos 6** *from Gen 5* (containing culture **PBAD-M0+PSal-M0+PVan-M0**): Add **Inducer 4 (Ara) at 4 mM + Kan at 1.25 µg/mL.** This condition aims to produce the culture **PBAD-M0+PSal-M0+PVan-M0-*K*_Van-*K*_Ara-*K*_Van-*K*_Ara-*K*_Van-*K*_Ara**, *corresponding to Gen 6.*

⇒ LEARNING tube for **Pos 9** *from Gen 5* (containing culture **PSal-M0+PTet-M0**): Add **Inducer 2 (aTc) at 0.2 µM + Kan at 1.25 µg/mL.** This condition aims to produce the culture **PSal-M0+PTet-M0-M-*K*_aTc-M-*K*_aTc-M-*K*_aTc**, *corresponding to Gen 6.*

⇒ MEMORY INCUBATION tube for **Pos 2** *from Gen 5* (containing culture **PTac-M1+PTtg-M1+PVan-M1**): Add only solvent control (**NO Inducer, NO Kanamycin**) This condition aims to produce the culture **PTac-M1+PTtg-M1+PVan-M1-*K*_IPTG-M-*K*_IPTG-M-*K*_IPTG-M**, *corresponding to Gen 6.*

⇒ MEMORY INCUBATION tube for **Pos 3** *from Gen 5* (containing culture **PTtg-M0+PVan-M0**): Add only solvent control (**NO Inducer, NO Kanamycin**) This condition aims to produce the culture **PTtg-M0+PVan-M0-M-M-M-M-M-M**, *corresponding to Gen 6.*

⇒ MEMORY INCUBATION tube for **Pos 4** *from Gen 5* (containing culture **PCin-M0+PLux-M0**): Add only solvent control (**NO Inducer, NO Kanamycin**) This condition aims to produce the culture **PCin-M0+PLux-M0-*K*_OC6-M-*K*_OC6-M-*K*_OC6-M**, *corresponding to Gen 6.*

⇒ MEMORY INCUBATION tube for **Pos 7** *from Gen 5* (containing culture **PBetI-M0+PTac-M0**): Add only solvent control (**NO Inducer, NO Kanamycin**) This condition aims to produce the culture **PBetI-M0+PTac-M0-*K*_Cho-M-*K*_Cho-M-*K*_Cho-M**, *corresponding to Gen 6.*

Detailed Learning/Memory Procedure (Day 6):

**1. Culture Preparation:** From -80°C, thaw one glycerol stock vial for each Day 5 co-culture (positions 1-4, 6-9).

**2. Refresh Cultures:** Inoculate thawed Day 5 stocks into M9+Cb (100 µg/mL) and grow to mid-log phase OR perform overnight growth as needed.

**3. Prepare Learning/Memory Tubes:** Using the refreshed/mid-log Day 5 cultures, prepare replicate tubes for LEARNING positions (1, 6, 8, 9) and MEMORY INCUBATION tubes for other positions (2, 3, 4, 7) in M9+Cb (80 µg/mL).

**4. Add Reagents & Incubate:** To LEARNING tubes, add specific Inducer(s)/Kanamycin as detailed above. To MEMORY INCUBATION tubes, add only solvent controls. Incubate ALL tubes (Learning and Incubation) for 8 hours at 37°C with shaking.

**5. Harvest and Store Generation Stocks:** After 8 hours, prepare glycerol stocks from ALL tubes. These represent the Generation 6 state. Label clearly (e.g., 'Pos X - Gen 6 - PostSel Rep A', 'Pos Y - Gen 6 - Memory Ctrl'), flash-freeze, and store at -80°C. These stocks will be used on Day 7.

##### *Day 6 Activity 2: Characterize Generation 5*

Objective: Characterize the response of *all 8 active* Generation 5 co-cultures to *all* relevant CHARACTERIZATION inducers (where the cognate promoter is present). ALL relevant combinations will be measured.

Using one biological replicate and four technical replicates for experiments and 4 for controls for Day 6 Characterization.

Procedure (Characterization - Day 6 Evening/Overnight): Requires 1 plate(s).

Follow standard characterization procedure (Appendix B) using the 'Generation 5' glycerol stocks (produced Day 5). Prepare cultures and add inducers at CHARACTERIZATION concentrations as specified below. Incubate in plate reader overnight.

###### Characterization Plate 1 (Day 6 Characterization (Gen 5))

Expected State after Day 6 Learning/Memory Incubation (Generation 6):

| Position | Promoter | Expected Change (Weight/M-stage) |
|---|---|---|
| 9 | PTet | 0.614 (M2) -> 0.575 (M3) |
| 8 | PBetI | 0.626 (M3) -> 0.543 (M4) |
| 6 | PBAD | 0.615 (M2) -> 0.525 (M3) |
| 1 | PLux | 0.543 (M2) -> 0.505 (M3) |

**Supplementary Table 50 | Characterization Plate 1 (Day 6 Characterization (Gen 5)).** Expected State after Day 6 Learning/Memory Incubation (Generation 6).

#### Day 7: Learn/Memory Generation 6 / Characterize Generation 6

(Learning based on Simulated Match 28 Loss)

Reference Simulation Match 28 (Leads to Day 7 Learning):

| | | |
|---|---|---|
| $X_3$ | $O_8$ | $O_6$ |
| $O_2$ | $X_1$ | $O_4$ |
| $X_5$ | $X_7$ | $X_9$ |

**Supplementary Table 51 | Reference Simulation Match 28 (Leads to Day 7 Learning).**

- Simulated Losing O Moves Sequence: [4, 6, 3, 2]
- Triggering X Moves Sequence (Inducers): [5, 1, 7, 8, 9]

##### *Day 7 Activity 1: Learn/Memory Generation 6 (Produce Generation 7)*

Objective: Apply sublethal kanamycin (Learning) to targeted Generation 6 cultures based on Match 28 AND perform parallel MEMORY INCUBATION without kanamycin on non-targeted cultures to produce the Generation 7 set. This step applies adaptation pressure or allows observation of culture stability ('memory') under identical growth conditions.
Learning Targets (using Day 6 Stocks): Position(s) 2, 3, 4, 6
Memory Incubation Targets (using Day 6 Stocks): Position(s) 1, 7, 8, 9
**Kanamycin Learning & Memory Incubation Details: Learning time is 4.50 hours 50 times dilution overnight culture.**

⇒ LEARNING tube for **Pos 3** *from Gen 6* (containing culture **PTtg-M0+PVan-M0**): Add **Inducer 7 (Nar) at 1 mM + Kan at 0.75 µg/mL.** This condition aims to produce the culture **PTtg-M0+PVan-M0-M-M-M-M-M-M-*K*_Nar**, *corresponding to Gen 7.*

⇒ LEARNING tube for **Pos 2** *from Gen 6* (containing culture **PTac-M1+PTtg-M1+PVan-M1**): Add **Inducer 8 (Van) at 100 µM + Kan at 1.75 µg/mL.** This condition aims to produce the culture **PTac-M1+PTtg-M1+PVan-M1-*K*_IPTG-M-*K*_IPTG-M-*K*_IPTG-M-*K*_Van**, *corresponding to Gen 7.*

⇒ LEARNING tube for **Pos 6** *from Gen 6* (containing culture **PBAD-M0+PSal-M0+PVan-M0**): Add **Inducer 1 (Sal) at 100 µM + Kan at 0.75 µg/mL.** This condition aims to produce the culture **PBAD-M0+PSal-M0+PVan-M0-*K*_Van-*K*_Ara-*K*_Van-*K*_Ara-*K*_Van-*K*_Ara-*K*_Sal**, *corresponding to Gen 7.*

⇒ LEARNING tube for **Pos 4** *from Gen 6* (containing culture **PCin-M0+PLux-M0**): Add **Inducer 5 (OC6) at 10 µM + Kan at 1.0 µg/mL.** This condition aims to produce the culture **PCin-M0+PLux-M0-*K*_OC6-M-*K*_OC6-M-*K*_OC6-M-*K*_OC6**, *corresponding to Gen 7.*

⇒ MEMORY INCUBATION tube for **Pos 1** *from Gen 6* (containing culture **PLux-M0**): Add only solvent control (**NO Inducer, NO Kanamycin**) This condition aims to produce the culture **PLux-M0-M-*K*_OC6-M-*K*_OC6-M-*K*_OC6-M**, *corresponding to Gen 7.*

⇒ MEMORY INCUBATION tube for **Pos 7** *from Gen 6* (containing culture **PBetI-M0+PTac-M0**): Add only solvent control (**NO Inducer, NO Kanamycin**) This condition aims to produce the culture **PBetI-M0+PTac-M0-*K*_Cho-M-*K*_Cho-M-*K*_Cho-M-M**, *corresponding to Gen 7.*

⇒ MEMORY INCUBATION tube for **Pos 8** *from Gen 6* (containing culture **PBetI-M1+PCin-M1+PTet-M1**): Add only solvent control (**NO Inducer, NO Kanamycin**) This condition aims to produce the culture **PBetI-M1+PCin-M1+PTet-M1-M-*K*_Cho-M-*K*_Cho-M-*K*_Cho-M**, *corresponding to Gen 7.*

⇒ MEMORY INCUBATION tube for **Pos 9** *from Gen 6* (containing culture **PSal-M0+PTet-M0**): Add only solvent control (**NO Inducer, NO Kanamycin**) This condition aims to produce the culture **PSal-M0+PTet-M0-M-*K*_aTc-M-*K*_aTc-M-*K*_aTc-M**, *corresponding to Gen 7.*

Detailed Learning/Memory Procedure (Day 7):

**1. Culture Preparation:** From -80°C, thaw one glycerol stock vial for each Day 6 co-culture (positions 1-4, 6-9).

**2. Refresh Cultures:** Inoculate thawed Day 6 stocks into M9+Cb (100 µg/mL) and grow to mid-log phase OR perform overnight growth as needed.

**3. Prepare Learning/Memory Tubes:** Using the refreshed/mid-log Day 6 cultures, prepare replicate tubes for LEARNING positions (2, 3, 4, 6) and MEMORY INCUBATION tubes for other positions (1, 7, 8, 9) in M9+Cb (80 µg/mL).

**4. Add Reagents & Incubate:** To LEARNING tubes, add specific Inducer(s)/Kanamycin as detailed above. To MEMORY INCUBATION flasks, add only solvent controls. Incubate ALL tubes (Learning and Incubation) for 8 hours at 37°C with shaking.

**5. Harvest and Store Generation Stocks:** After 8 hours, prepare glycerol stocks from ALL tubes. These represent the Generation 7 state. Label clearly (e.g., 'Pos X - Gen 7 - PostSel Rep A', 'Pos Y - Gen 7 - Memory Ctrl'), flash-freeze, and store at -80°C. These stocks will be used on Day 8.

##### *Day 7 Activity 2: Characterize Generation 6*

Objective: Characterize the response of *all 8 active* Generation 6 co-cultures to *all* relevant CHARACTERIZATION inducers (where the cognate promoter is present). ALL relevant combinations will be measured.

Using one biological replicate and four technical replicates for experiments and 4 for controls for Day 7 Characterization.

Procedure (Characterization - Day 7 Evening/Overnight): Requires 1 plate(s).

Follow standard characterization procedure (Appendix B) using the 'Generation 6' glycerol stocks (produced Day 6). Prepare cultures and add inducers at CHARACTERIZATION concentrations as specified below. Incubate in plate reader overnight.

###### Characterization Plate 1 (Day 7 Characterization (Gen 6))

Expected State after Day 7 Learning/Memory Incubation (Generation 7):

| Position | Promoter | Expected Change (Weight/M-stage) |
|---|---|---|
| 6 | PSal | 0.727 (M0) -> 0.593 (M1) |
| 4 | PLux | 0.505 (M3) -> 0.369 (M4) |
| 3 | PTtg | 0.664 (M0) -> 0.642 (M1) |
| 2 | PVan | 0.872 (M1) -> 0.868 (M2) |

**Supplementary Table 52 | Characterization Plate 1 (Day 7 Characterization (Gen 6)).** Expected State after Day 7 Learning/Memory Incubation (Generation 7).

#### Day 8: Final Characterization

Objective: Characterize the final response of Generation 7 co-cultures (produced on Day 7) to *all* relevant CHARACTERIZATION inducers (where the cognate promoter is present).

Using one biological replicate and four technical replicates for experiments and 4 for controls for Final phase.

Procedure (Final Characterization - Night 7): Requires 1 plate(s).

Follow standard characterization procedure (Appendix B) using the 'Generation 7' glycerol stocks (produced on Day 7). Prepare cultures and add inducers at CHARACTERIZATION concentrations as specified below.

Characterization Plate 1 (Final Characterization (Gen 7))

#### Supplementary Note 6: Scaling strategies and combinatorial promoter design

This Note collects additional analyses and design details related to scaling memregulon architectures, including strategies to prevent weight vanishing and the use of combinatorial promoters to increase input dimensionality.

**Combinatorial promoters and potential scaling of input dimensionality**

We explore scaling strategies beyond simply increasing the number of basic memregulons. One approach uses engineered combinatorial promoters that respond to multiple chemical inputs, extending the capabilities of the existing nine single-input memregulons (the 9YES library). We engineer AND-logic promoters by adding operator sites for secondary transcription factors downstream of existing inducible promoters. Testing combinations such as PTet with Nar/IPTG/Cho/DAPG/Van operators identifies four constructs with reliable AND-like activity: PTet-Ttg, PTac-Tet, PTac-Van and PVan-Ttg. These form the 4AND library described in the main text (Fig. 4a,b).

Characterisation confirms substantial activation of the YES and AND memregulons upon induction compared with non-induced or partially induced states. Consecutive kanamycin adaptation cycles of the AND library showed weight decreases in activated conditions, consistent with the physical learning rule, and flow-cytometry analyses of single-cell weight distributions provided an orthogonal check against plate-reader estimates.

This combinatorial promoter strategy could, in principle, allow scaling to ~66 orthogonal inputs using the Marionette transcription factor set, although practical limits arising from metabolic load, crosstalk and available inducers must be considered. In the simulations described in this note, architectures that include 4AND or 3HYB elements reach high expertise in fewer adaptation rounds than architectures using only single-input promoters, suggesting that combinatorial logic can accelerate learning under the same physical learning rule.

#### Supplementary Note 7: Derivation of bulk memregulon weights from ratiometric fluorescence

This note derives the population-average memregulon weight $W$ from bulk fluorescence measurements (plate reader totals or flow cytometry totals). For each cell $i$, we define the underlying weight $w_i = n_{1,i} / (n_{1,i} + n_{2,i})$, where $n_{1,i}$ and $n_{2,i}$ are the copy numbers of plasmids P1 (mCherry) and P2 (EGFP), respectively. We seek an estimator for the population-average weight $W \in [0,1]$, inferred from bulk fluorescence while accounting for background fluorescence and spectral cross-talk between channels. We report $W$ as the mean of $w_i$ over the population. The derivation assumes (i) fluorescence adds linearly across cells and plasmid copies, and (ii) acquisition settings are identical across sample and controls so proportionality factors cancel in ratios.

**Background-corrected totals and controls.** Let $R$ and $G$ denote the measured total signals in the red and green detection channels for a mixed culture. Let $R^{\mathrm{bg}}$ and $G^{\mathrm{bg}}$ denote the corresponding background signals measured under identical acquisition settings in the absence of reporters (for example, media plus cells lacking reporters). We define the background-corrected totals

$$\tilde{R} = R - R^{\mathrm{bg}}, \qquad \tilde{G} = G - G^{\mathrm{bg}}.$$

We measure the same background-corrected quantities for two single-reporter controls acquired under identical induction and instrument settings:

P1-only control: $(\tilde{R}^{P1}, \tilde{G}^{P1})$,

P2-only control: $(\tilde{R}^{P2}, \tilde{G}^{P2})$.

These control totals include any spectral cross-talk because each reporter can contribute signal to both channels.

**Minimal linear decomposition in bulk.** We assume only linear superposition of fluorescence contributions at the level of bulk totals: the total signal equals the sum over cells, and each reporter contributes additively to each detection channel. Under this assumption, the mixed-sample background-corrected totals decompose as

$$\tilde{R} = k[W\,\tilde{R}^{P1} + (1-W)\,\tilde{R}^{P2}], \qquad \tilde{G} = k\left[W\,\tilde{G}^{P1} + (1-W)\,\tilde{G}^{P2}\right],$$

where $k$ is a common proportionality factor that captures global effects shared by both channels (for example, sampling volume and overall gain). Because we take ratios of $\tilde{G}$ and $\tilde{R}$, $k$ cancels and does not need to be estimated.

From the background-corrected control measurements we define three dimensionless ratios:

$$\lambda = \frac{\tilde{R}^{P1}}{\tilde{G}^{P2}}, \qquad \beta = \frac{\tilde{G}^{P1}}{\tilde{R}^{P1}}, \qquad \gamma = \frac{\tilde{R}^{P2}}{\tilde{G}^{P2}}.$$

Here λ compares the on-channel outputs of the reporters (red from the P1-only control versus green from the P2-only control). The quantity β measures green leakage when only the red reporter is present (green signal relative to red on-channel). The quantity γ measures red leakage when only the green reporter is present (red signal relative to green on-channel).

Starting from linear decomposition in bulk, we form $\rho = \tilde{G}/\tilde{R}$ and eliminate $k$:

$$\rho = \frac{W\,\tilde{G}^{P1} + (1-W)\,\tilde{G}^{P2}}{W\,\tilde{R}^{P1} + (1-W)\,\tilde{R}^{P2}}.$$

We rewrite it in terms of the calibration ratios by dividing numerator and denominator by $\tilde{G}^{P2}$ and expressing $\tilde{G}^{P1} = \beta\,\tilde{R}^{P1}$ and $\tilde{R}^{P2} = \gamma\,\tilde{G}^{P2}$:

$$\rho \;=\; \frac{W\,(\tilde{G}^{P1}/\tilde{G}^{P2}) + (1-W)}{W\,(\tilde{R}^{P1}/\tilde{G}^{P2}) + (1-W)\,(\tilde{R}^{P2}/\tilde{G}^{P2})} \;=\; \frac{W\,\beta\lambda + (1-W)}{W\,\lambda + (1-W)\,\gamma}.$$

Solving for $W$ gives the bulk memregulon weight:

$$\boxed{W^{\text{bulk}} \;=\; \frac{1-\gamma\rho}{1-\lambda\beta+(\lambda-\gamma)\rho}}.$$

The same formula applies to flow cytometry totals when $\tilde{G}$ and $\tilde{R}$ are taken as sums over all gated events, where we replace $W^{\text{bulk}}$ by the single-cell weights $W_i = R_i/(R_i + G_i)$.

#### Supplementary Note 8: General weight analysis from plate reader fluorescence

For each well, we estimated the time delay required to reach a reference optical density (ODsync), which was subsequently used to align fluorescence trajectories across wells prior to modeling. To mitigate measurement artifacts frequently observed in early OD readings, we implemented an early-OD artifact handling protocol: if the OD was observed to decrease at any point within the first 3 hours, all preceding time-points were excluded from quantitative fits. Plots display all raw data points for full transparency, but model-fit lines commence only from this corrected start time. For technical replicates, the latest (maximum) corrected start time across all replicates was applied to ensure a consistent analytical window. The cellular weight, a metric derived from the ratio of red to green fluorescence (RFP/GFP), was explicitly set to 0 for wells where the red fluorescence rate (RFP/OD) fell below a threshold of 900 a.u., a condition indicating negligible P1 promoter activity. The OD was modeled using either a 4-parameter logistic function or a Baranyi–Roberts variant with parameters $od_0$, c, $\mu_2$, ν, b, $a_2$, and $a_3$.

##### Lambda and beta parameters

| Promoter # | Promoter name | λ (mean ± s.e.m.) | β (mean ± s.e.m.) |
|---|---|---|---|
| 1 | PSal | 0.903 ± 0.270 | 0.210 ± 0.029 |
| 2 | PTet | 0.507 ± 0.023 | 0.263 ± 0.033 |
| 3 | PBetI | 0.189 ± 0.099 | 1.816 ± 0.979 |
| 4 | PBAD | 1.335 ± 0.082 | 0.089 ± 0.011 |
| 5 | PLux | 1.162 ± 0.070 | 0.181 ± 0.015 |
| 6 | PCin | 0.793 ± 0.051 | 0.289 ± 0.033 |
| 7 | PTtg | 0.127 ± 0.013 | 1.248 ± 0.175 |
| 8 | PVan | 1.024 ± 0.081 | 0.163 ± 0.012 |
| 9 | PTac | 0.687 ± 0.161 | 0.087 ± 0.058 |
| 10 | PPhIF | 0.374 ± 0.028 | 0.033 ± 0.007 |
| 11 | PVan-Ttg | 0.151 ± 0.035 | 0.203 ± 0.128 |
| 12 | PBetI-Tac | 0.096 ± 0.001 | 2.578 ± 0.024 |
| 14 | PPhIF-Tac | 0.087 ± 0.009 | 0.591 ± 0.000 |
| 15 | PTtg-Tac | 0.032 ± 0.006 | 0.490 ± 0.000 |
| 16 | PTet-PhIF | 0.543 ± 0.000 | 0.020 ± 0.000 |
| 17 | PTet-Bet | 0.128 ± 0.000 | 0.052 ± 0.000 |
| 19 | PTac-Van | 0.281 ± 0.019 | 0.274 ± 0.021 |
| 21 | PTac-Tet | 0.515 ± 0.083 | 0.031 ± 0.000 |
| 24 | PTet-Ttg | 0.292 ± 0.009 | 0.046 ± 0.000 |

**Supplementary Table 53 | Lambda and beta parameters.**

### Supplementary Note 9: Weight stabilities from plate reader fluorescence

#### *Summary Table*

| Cell Name | Growth rate µ (1/h) | Green fluorescence | Red fluorescence | Weight | Replicates |
|---|---|---|---|---|---|
| PSal(Sal) Sal 100 Clone M0 K0 | 0.279 ± 0.003 | 12,607 ± 158 | 6,099 ± 35 | 0.3735 ± 0.0027 | 8 |
| PSal(Sal) Sal 100 Clone M1 K0 | 0.276 ± 0.003 | 13,911 ± 341 | 6,564 ± 154 | 0.384 ± 0.006 | 12 |
| PSal(Sal) Sal 100 Clone M2 K0 | 0.275 ± 0.005 | 14,618 ± 598 | 6,917 ± 274 | 0.3678 ± 0.0028 | 8 |
| PSal(Sal) Sal 100 Clone M3 K0 | 0.283 ± 0.004 | 13,185 ± 610 | 6,884 ± 120 | 0.396 ± 0.008 | 8 |
| PSal(Sal) Sal 100 Clone M4 K0 | 0.2919 ± 0.0023 | 12,429 ± 237 | 6,614 ± 51 | 0.399 ± 0.006 | 4 |
| PTet(aTc) aTc 100 Clone M1 K0 | 0.228 ± 0.019 | 12,284 ± 232 | 18,518 ± 495 | 0.8314 ± 0.0023 | 4 |
| PBetI(Cho) Cho 10000 M0 K0 | 0.280 ± 0.004 | 2,886 ± 56 | 6,236 ± 210 | 1.342 ± 0.004 | 8 |
| PBetI(Cho) Cho 10000 M1 K0 | 0.265 ± 0.008 | 2,541 ± 97 | 6,482 ± 345 | 1.365 ± 0.005 | 12 |
| PBetI(Cho) Cho 10000 M2 K0 | 0.283 ± 0.004 | 2,259 ± 99 | 4,927 ± 267 | 1.343 ± 0.008 | 8 |
| PBetI(Cho) Cho 10000 M3 K0 | 0.272 ± 0.011 | 2,466 ± 44 | 5,367 ± 159 | 1.343 ± 0.006 | 8 |
| PBetI(Cho) Cho 10000 M4 K0 | 0.2885 ± 0.0020 | 2,556 ± 38 | 5,158 ± 72 | 1.331 ± 0.004 | 4 |
| PBAD(Ara) Ara 4000 M0 K0 | 0.188 ± 0.012 | 10,982 ± 96 | 60,677 ± 291 | 0.8908 ± 0.0019 | 8 |
| PBAD(Ara) Ara 4000 M1 K0 | 0.189 ± 0.009 | 11,387 ± 171 | 66,252 ± 1,576 | 0.900 ± 0.004 | 12 |
| PBAD(Ara) Ara 4000 M2 K0 | 0.168 ± 0.014 | 12,390 ± 374 | 72,228 ± 1,300 | 0.901 ± 0.003 | 7 |
| PBAD(Ara) Ara 4000 M3 K0 | 0.217 ± 0.011 | 9,778 ± 222 | 66,000 ± 611 | 0.927 ± 0.005 | 8 |
| PBAD(Ara) Ara 4000 M4 K0 | 0.2654 ± 0.0022 | 6,073 ± 28 | 17,429 ± 119 | 0.8367 ± 0.0020 | 4 |
| PLux(OC6) OC6 10 M0 K0 | 0.253 ± 0.007 | 8,731 ± 110 | 20,318 ± 462 | 0.793 ± 0.014 | 12 |
| PLux(OC6) OC6 10 M1 K0 | 0.253 ± 0.005 | 8,244 ± 108 | 21,428 ± 382 | 0.820 ± 0.011 | 16 |
| PLux(OC6) OC6 10 M2 K0 | 0.257 ± 0.007 | 8,013 ± 173 | 23,221 ± 445 | 0.855 ± 0.013 | 12 |
| PLux(OC6) OC6 10 M3 K0 | 0.251 ± 0.006 | 6,735 ± 92 | 20,302 ± 415 | 0.866 ± 0.012 | 12 |
| PLux(OC6) OC6 10 M4 K0 | 0.224 ± 0.015 | 8,019 ± 640 | 43,105 ± 8,182 | 0.9265 ± 0.0029 | 8 |
| PVan(Van) Van 100 M0 | 0.257 ± 0.020 | 12,559 ± 2,932 | 10,120 ± 2,519 | 0.51 ± 0.14 | 8 |
| PVan(Van) Van 100 M1 | 0.250 ± 0.012 | 20,673 ± 1,069 | 6,318 ± 1,220 | 0.204 ± 0.027 | 8 |
| PVan(Van) Van 100 M2 | 0.284 ± 0.005 | 18,967 ± 603 | 6,597 ± 1,250 | 0.23 ± 0.03 | 8 |
| PVan(Van) Van 100 M3 | 0.285 ± 0.006 | 18,299 ± 540 | 6,557 ± 1,037 | 0.238 ± 0.027 | 8 |
| PCin(OHC14) OHC14 M0 K0 | 0.227 ± 0.008 | 6,051 ± 223 | 2,933 ± 155 | 0.415 ± 0.007 | 5 |
| PCin(OHC14) OHC14 M1 K0 | 0.253 ± 0.007 | 5,725 ± 82 | 2,840 ± 146 | 0.421 ± 0.014 | 6 |
| PCin(OHC14) OHC14 M2 K0 | 0.248 ± 0.011 | 5,274 ± 101 | 2,844 ± 60 | 0.446 ± 0.009 | 4 |
| PCin(OHC14) OHC14 M3 K0 | 0.248 ± 0.015 | 5,407 ± 143 | 2,952 ± 24 | 0.450 ± 0.006 | 6 |
| PCin(OHC14) OHC14 M4 K0 | 0.25 | 4,845 | 2,842 | 0.47 | 1 |
| PTtg(Nar) Nar 1000 M0 K0 | 0.262 ± 0.007 | 6,774 ± 205 | 1,356 ± 48 | 0.6775 ± 0.0023 | 8 |
| PTtg(Nar) Nar 1000 M1 K0 | 0.259 ± 0.009 | 4,462 ± 153 | 987 ± 48 | 0.705 ± 0.006 | 11 |
| PTtg(Nar) Nar 1000 M2 K0 | 0.272 ± 0.004 | 4,148 ± 134 | 1,030 ± 33 | 0.739 ± 0.003 | 8 |
| PTtg(Nar) Nar 1000 M3 K0 | 0.2859 ± 0.0013 | 4,283 ± 39 | 1,027 ± 14 | 0.729 ± 0.005 | 8 |
| PTtg(Nar) Nar 1000 M4 K0 | 0.2794 ± 0.0009 | 4,219 ± 31 | 1,028 ± 10 | 0.734 ± 0.004 | 4 |
| PVan(Van) Van 100 M1 K0 | 0.263 ± 0.013 | 23,293 ± 626 | 9,584 ± 630 | 0.299 ± 0.012 | 8 |
| PVan(Van) Van 100 M2 K0 | 0.286 ± 0.005 | 19,665 ± 858 | 6,008 ± 448 | 0.238 ± 0.008 | 4 |
| PVan(Van) Van 100 M3 K0 | 0.280 ± 0.004 | 21,138 ± 1,183 | 10,894 ± 447 | 0.355 ± 0.004 | 4 |
| PTac(IPTG) IPTG 500 M0 K0 | 0.224 ± 0.014 | 3,973 ± 85 | 1,082 ± 59 | 0.287 ± 0.007 | 8 |
| PTac(IPTG) IPTG 500 M1 K0 | 0.297 ± 0.003 | 3,808 ± 68 | 975 ± 58 | 0.273 ± 0.009 | 12 |

| | | | | | |
|---|---|---|---|---|---|
| PTac(IPTG) IPTG 500 M2 K0 | 0.263 ± 0.019 | 3,618 ± 60 | 956 ± 44 | 0.281 ± 0.006 | 8 |
| PTac(IPTG) IPTG 500 M3 K0 | 0.3039 ± 0.0016 | 3,492 ± 39 | 924 ± 43 | 0.282 ± 0.007 | 8 |
| PTac(IPTG) IPTG 500 M4 K0 | 0.312 ± 0.003 | 3,444 ± 29 | 922 ± 8 | 0.2850 ± 0.0007 | 4 |
| PTet(aTc) aTc 100 M1 K0 | 0.187 ± 0.004 | 10,480 ± 106 | 17,759 ± 535 | 0.743 ± 0.005 | 4 |
| PTet(aTc) aTc 100 M0 K0 | 0.238 ± 0.016 | 12,654 ± 89 | 16,403 ± 134 | 0.680 ± 0.003 | 4 |
| PTet(aTc) aTc 100 M2 K0 | 0.2433 ± 0.0028 | 10,704 ± 326 | 16,820 ± 585 | 0.726 ± 0.003 | 4 |
| PTet(aTc) aTc 100 M3 K0 | 0.234 ± 0.011 | 11,381 ± 126 | 19,522 ± 239 | 0.746 ± 0.004 | 4 |
| PTet(aTc) aTc 100 M4 K0 | 0.2676 ± 0.0016 | 11,062 ± 38 | 18,466 ± 209 | 0.7398 ± 0.0026 | 4 |
| PVan-Ttg(Van,Nar) Van 21 Nar 280 M0 K0 | 0.281 ± 0.015 | 10,050 ± 1,665 | 529 ± 126 | 0.237 ± 0.018 | 8 |
| PVan-Ttg(Van,Nar) Van 21 Nar 280 M1 K0 | 0.268 ± 0.010 | 6,641 ± 1,047 | 309 ± 76 | 0.205 ± 0.018 | 12 |
| PVan-Ttg(Van,Nar) Van 21 Nar 280 M2 K0 | 0.274 ± 0.012 | 5,669 ± 151 | 198 ± 13 | 0.186 ± 0.007 | 12 |
| PVan-Ttg(Van,Nar) Van 21 Nar 280 M3 K0 | 0.242 ± 0.013 | 5,550 ± 133 | 218 ± 13 | 0.205 ± 0.006 | 12 |
| PVan-Ttg(Van,Nar) Van 21 Nar 280 M4 K0 | 0.289 ± 0.007 | 6,471 ± 889 | 314 ± 64 | 0.225 ± 0.012 | 12 |
| PTac-Van(IPTG,Van) IPTG 500 Van 21 M0 K0 | 0.263 ± 0.012 | 7,438 ± 213 | 5,678 ± 673 | 0.756 ± 0.019 | 12 |
| PTac-Van(IPTG,Van) IPTG 500 Van 21 M1 K0 | 0.290 ± 0.008 | 8,315 ± 137 | 5,711 ± 547 | 0.737 ± 0.019 | 12 |
| PTac-Van(IPTG,Van) IPTG 500 Van 21 M2 K0 | 0.280 ± 0.009 | 8,662 ± 134 | 4,506 ± 125 | 0.682 ± 0.004 | 12 |
| PTac-Van(IPTG,Van) IPTG 500 Van 21 M3 K0 | 0.286 ± 0.008 | 8,036 ± 221 | 3,918 ± 165 | 0.665 ± 0.006 | 12 |
| PTac-Van(IPTG,Van) IPTG 500 Van 21 M4 K0 | 0.270 ± 0.011 | 9,709 ± 173 | 4,585 ± 136 | 0.658 ± 0.004 | 12 |
| PTac-Tet(IPTG,aTc) IPTG 500 aTc 100 M0 K0 | 0.229 ± 0.017 | 9,271 ± 155 | 15,086 ± 412 | 0.769 ± 0.003 | 4 |
| PTac-Tet (IPTG,aTc) IPTG 500 aTc 100 M1 K0 | 0.2430 ± 0.0025 | 10,164 ± 52 | 16,489 ± 337 | 0.768 ± 0.003 | 4 |
| PTac-Tet (IPTG,aTc) IPTG 500 aTc 100 M2 K0 | 0.224 ± 0.019 | 9,656 ± 181 | 17,110 ± 456 | 0.7843 ± 0.0017 | 4 |
| PTac-Tet (IPTG,aTc) IPTG 500 aTc 100 M3 K0 | 0.209 ± 0.014 | 11,760 ± 91 | 13,279 ± 339 | 0.694 ± 0.005 | 4 |
| PTac-Tet (IPTG,aTc) IPTG 500 aTc 100 M4 K0 | 0.235 ± 0.014 | 13,382 ± 402 | 14,088 ± 313 | 0.679 ± 0.003 | 4 |
| PTac-Tet (IPTG,aTc) IPTG 500 aTc 100M0 K0 | 0.233 ± 0.007 | 10,496 ± 96 | 17,102 ± 646 | 0.768 ± 0.006 | 7 |
| PTac-Tet (IPTG,aTc) IPTG 500 aTc 100M1 K0 | 0.2406 ± 0.0025 | 10,635 ± 178 | 16,048 ± 294 | 0.754 ± 0.005 | 12 |
| PTac-Tet (IPTG,aTc) IPTG 500 aTc 100M2 K0 | 0.234 ± 0.006 | 10,588 ± 167 | 14,447 ± 226 | 0.7343 ± 0.0026 | 12 |
| PTac-Tet (IPTG,aTc) IPTG 500 aTc 100M3 K0 | 0.226 ± 0.012 | 13,033 ± 213 | 15,070 ± 331 | 0.6992 ± 0.0025 | 10 |
| PTac-Tet (IPTG,aTc) IPTG 500 aTc 100M4 K0 | 0.241 ± 0.009 | 12,639 ± 355 | 13,219 ± 239 | 0.677 ± 0.006 | 8 |
| PTet-Ttg(aTc,Nar) aTc 100 Nar 280 M K0 | 0.29 | 3,714 | 265 | 0.20 | 1 |
| PTet-Ttg(aTc,Nar) aTc 100 Nar 280 M0 K0 | 0.244 ± 0.007 | 3,908 ± 224 | 238 ± 21 | 0.173 ± 0.013 | 14 |
| PTet-Ttg(aTc,Nar) aTc 100 Nar 280 M1 K0 | 0.246 ± 0.007 | 3,923 ± 106 | 130 ± 10 | 0.101 ± 0.007 | 16 |
| PTet-Ttg(aTc,Nar) aTc 100 Nar 280 M2 K0 | 0.240 ± 0.005 | 3,623 ± 107 | 90 ± 13 | 0.077 ± 0.011 | 16 |
| PTet-Ttg(aTc,Nar) aTc 100 Nar 280 M3 K0 | 0.250 ± 0.007 | 4,248 ± 129 | 96 ± 4 | 0.073 ± 0.003 | 16 |
| PTet-Ttg(aTc,Nar) aTc 100 Nar 280 M4 K0 | 0.234 ± 0.008 | 4,094 ± 126 | 100 ± 6 | 0.077 ± 0.004 | 16 |

**Supplementary Table 54 | Summary Table.**

#### Analysis of growth and fluorescence curves from plate reader

This section presents a comprehensive analysis of cellular growth and fluorescence dynamics, aggregated across all experimental datasets. The data is organized by promoter system and subsequently sorted by the experimental condition identifier (M-suffix) to provide a clear, logical progression. Within this framework, we distinguish between two levels of replication fundamental to biological experiments:

- Biological Replicates: These are distinct cultures, often originating from different colonies or clones (e.g., 'Clone A' and 'Clone B'), that are subjected to the same experimental conditions. They are treated as independent measurements of the biological process under investigation.
- Technical Replicates: These are multiple measurements taken from the same biological sample (e.g., multiple wells in a plate inoculated from the same starter culture). They serve to quantify the precision and variability of the measurement technique itself.

Numerical audit note for Supplementary Notes 9-11. Standard-deviation columns and incomplete rows were checked against the corresponding summary tables and source data. Header-only promoter tables in Supplementary Note 10 were repopulated from the Note 10 summary rows. Singleton biological-level groups are marked as n.d. (single entry) rather than interpreted as true zero biological variance; the technical-replicate uncertainty remains reported in the corresponding summary table and Source Data workbook.

Throughout this analysis, each curve representing a biological replicate is computed as the mean of its corresponding technical replicates. The variability among these technical replicates is visualized as a shaded region representing the standard error of the mean (SEM). All mean curves are linearly interpolated between measured time points to ensure visual continuity.

#### Weight stability Analysis

This section presents cellular weight measurements grouped by memory state (M-suffix) Each bar represents the mean weight across biological replicates. Significance tests (TOST for stability, Welch for drift) are performed using the pool of all technical replicates to rigorously assess variation.

##### Promoter PSal

Stability Tolerance ($\delta$) = 0.5205 (2 $x$ Pooled SD of pooled technical replicates across biological replicates: 0.2603)

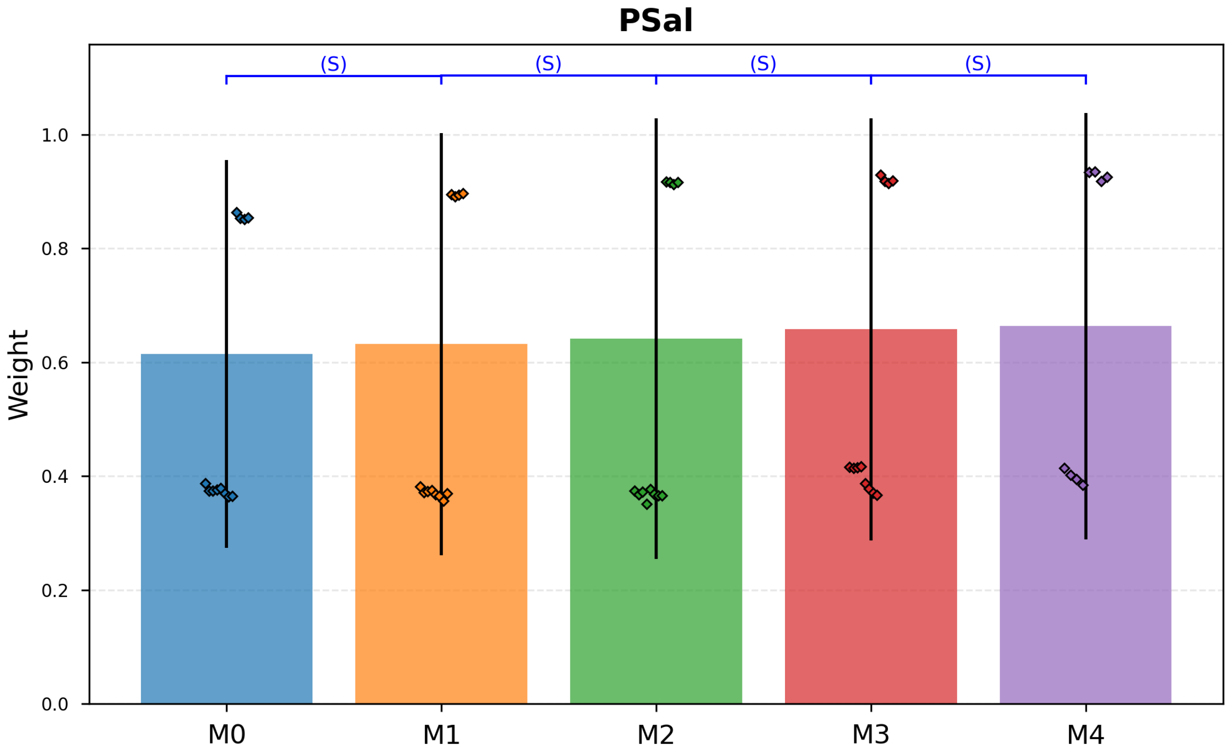

**Supplementary Fig. 48 | Promoter PSal.** Stability Tolerance (δ) = 0.5205 (2 *x* Pooled SD of pooled technical replicates across biological replicates: 0.2603).

- M0 vs M1: Estimated shift Δ = 0.0105. Data inconsistent with drift ≥ ±0.5205 (p_stable = 2.5 × $10^{-5}$). Classified as Stable.
- M1 vs M2: Estimated shift Δ = 0.0057. Data inconsistent with drift ≥ ±0.5205 (p_stable = 4.6 × $10^{-5}$). Classified as Stable.
- M2 vs M3: Estimated shift Δ = 0.0200. Data inconsistent with drift ≥ ±0.5205 (p_stable = 6.4 × $10^{-5}$). Classified as Stable.
- M3 vs M4: Estimated shift Δ = 0.0931. Data inconsistent with drift ≥ ±0.5205 (p_stable = 0.002). Classified as Stable.

| M-State | Mean Weight | Std Dev |
|---|---|---|
| M0 | 0.6147 | 0.3410 |
| M1 | 0.6322 | 0.3708 |
| M2 | 0.6418 | 0.3874 |
| M3 | 0.6579 | 0.3711 |
| M4 | 0.6635 | 0.3743 |

**Supplementary Table 55 | Stability Tolerance (δ) = 0.5205 (2 *x* Pooled SD of pooled technical replicates across biological replicates: 0.2603).**

#### Promoter PTet

Stability Tolerance (δ) = 0.0093 (2 *x* Pooled SD of pooled technical replicates across biological replicates: 0.0046)

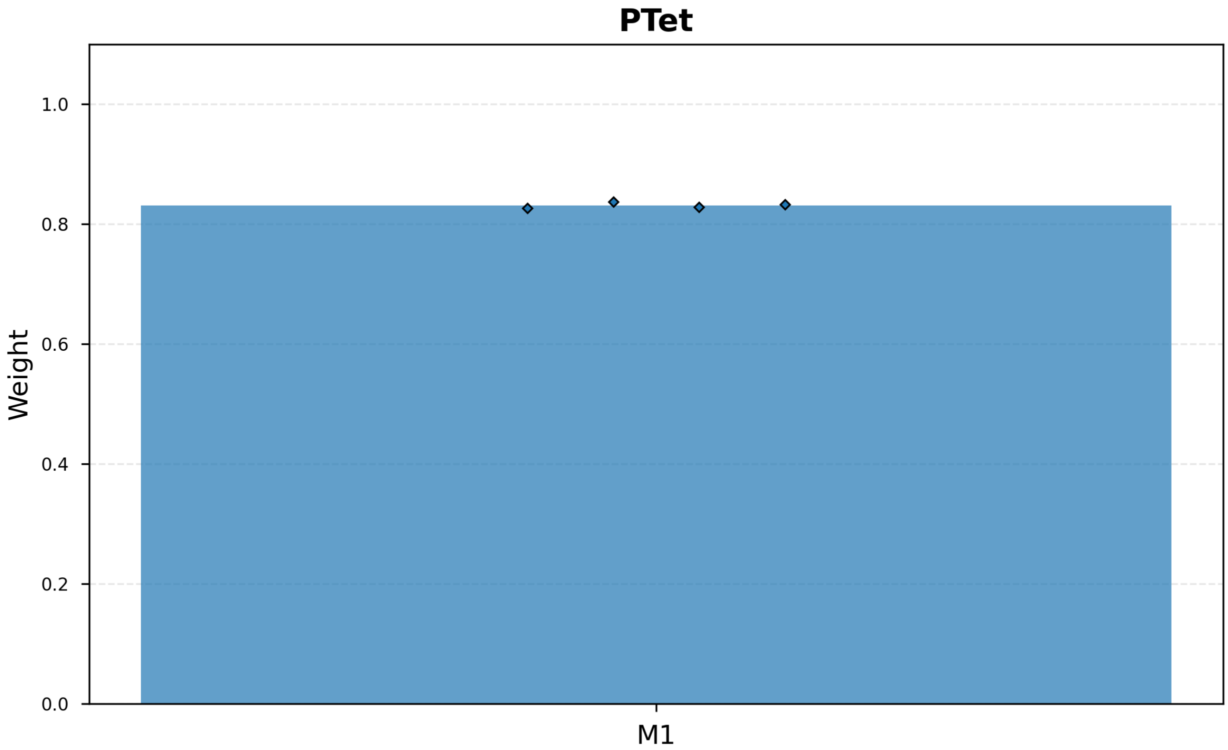

**Supplementary Fig. 49 | Promoter PTet.** Stability Tolerance (δ) = 0.0093 (2 *x* Pooled SD of pooled technical replicates across biological replicates: 0.0046).

| M-State | Mean Weight | Std Dev |
|---|---|---|
| M1 | 0.8314 | n.d. (single entry) |

**Supplementary Table 56 | Promoter PTet.** Stability Tolerance (δ) = 0.0093 (2 *x* Pooled SD of pooled technical replicates across biological replicates: 0.0046).

#### Promoter PBetI

Stability Tolerance (δ) = 0.0332 (2 *x* Pooled SD of pooled technical replicates across biological replicates: 0.0166)

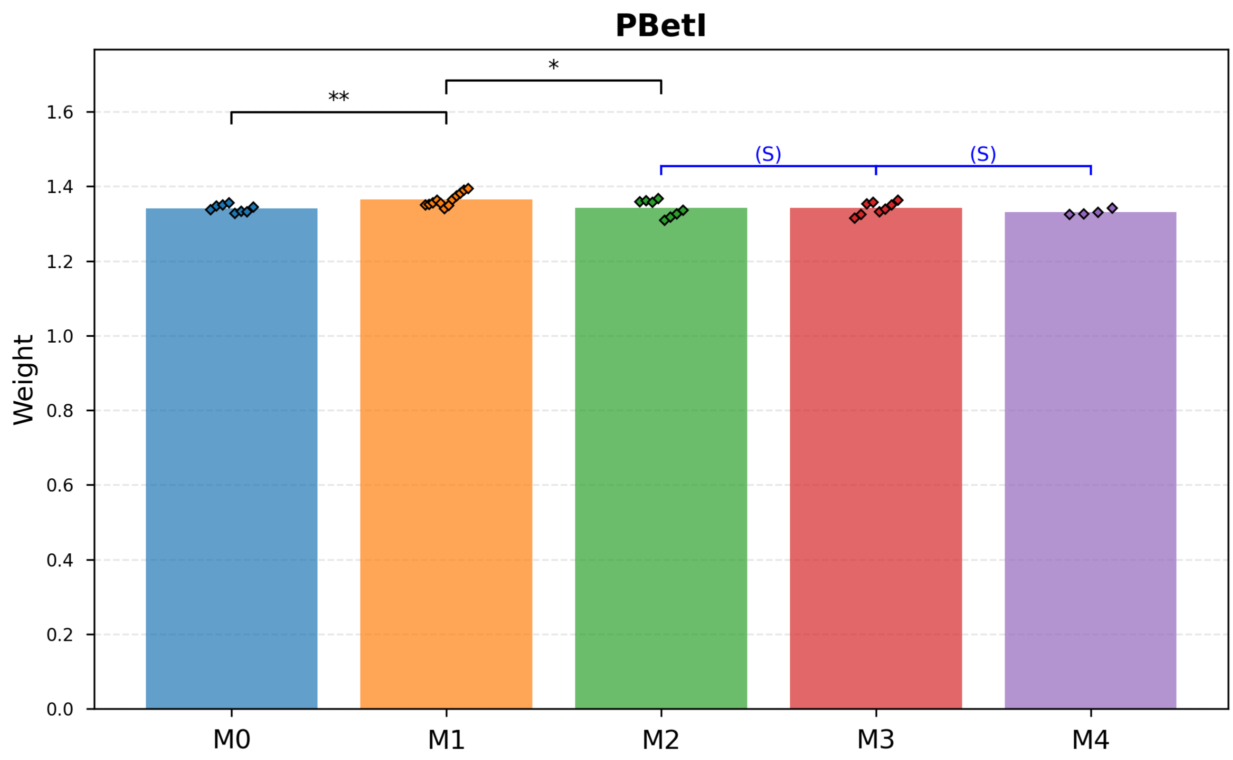

**Supplementary Fig. 50 | Promoter PBetI.** Stability Tolerance (δ) = 0.0332 (2 *x* Pooled SD of pooled technical replicates across biological replicates: 0.0166).

- M0 vs M1: Stability not supported (p_stable = 0.056). Welch's test detected shift (p_change = 0.001). Classified as Drift.
- M1 vs M2: Stability not supported (p_stable = 0.136). Welch's test detected shift (p_change = 0.034). Classified as Drift.
- M2 vs M3: Estimated shift Δ = 0.0002. Data inconsistent with drift ≥ ±0.0332 (p_stable = 0.003). Classified as Stable.
- M3 vs M4: Estimated shift Δ = -0.0114. Data inconsistent with drift ≥ ±0.0332 (p_stable = 0.006). Classified as Stable.

| M-State | Mean Weight | Std Dev |
|---|---|---|
| M0 | 1.3418 | n.d. (single entry) |
| M1 | 1.3649 | n.d. (single entry) |
| M2 | 1.3425 | n.d. (single entry) |
| M3 | 1.3427 | n.d. (single entry) |
| M4 | 1.3313 | n.d. (single entry) |

**Supplementary Table 57 | Stability Tolerance (δ) = 0.0332 (2 *x* Pooled SD of pooled technical replicates across biological replicates: 0.0166).**

#### Promoter PBAD

Stability Tolerance (δ) = 0.0240 (2 *x* Pooled SD of pooled technical replicates across biological replicates: 0.0120)

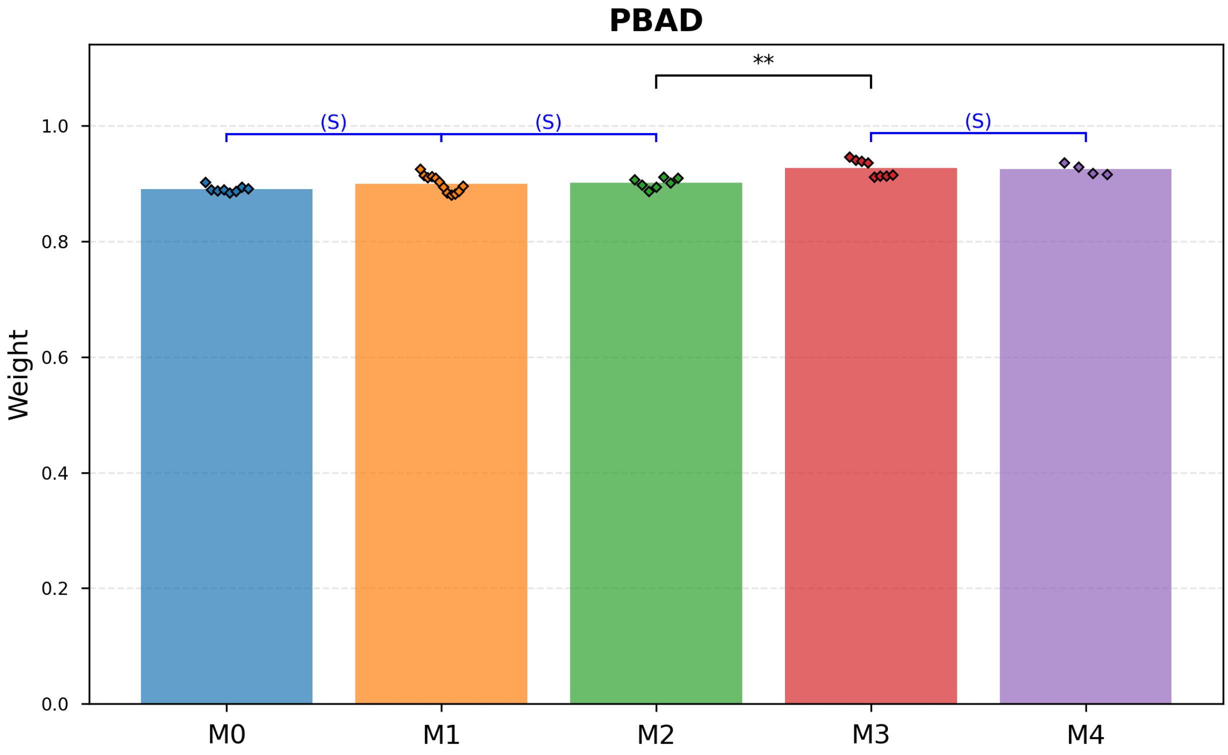

**Supplementary Fig. 51 | Promoter PBAD.** Stability Tolerance (δ) = 0.0240 (2 *x* Pooled SD of pooled technical replicates across biological replicates: 0.0120).

- M0 vs M1: Estimated shift Δ = 0.0091. Data inconsistent with drift ≥ ±0.0240 (p_stable = 0.003). Classified as Stable.
- M1 vs M2: Estimated shift Δ = 0.0013. Data inconsistent with drift ≥ ±0.0240 (p_stable = 3.1 × $10^{-4}$). Classified as Stable.
- M2 vs M3: Stability not supported (p_stable = 0.610). Welch's test detected shift (p_change = 0.001). Classified as Drift.
- M3 vs M4: Estimated shift Δ = -0.0020. Data inconsistent with drift ≥ ±0.0240 (p_stable = 0.006). Classified as Stable.

| M-State | Mean Weight | Std Dev |
|---|---|---|
| M0 | 0.8908 | n.d. (single entry) |
| M1 | 0.8999 | n.d. (single entry) |
| M2 | 0.9011 | n.d. (single entry) |
| M3 | 0.9269 | n.d. (single entry) |
| M4 | 0.9249 | n.d. (single entry) |

**Supplementary Table 58 | Stability Tolerance (δ) = 0.0240 (2 *x* Pooled SD of pooled technical replicates across biological replicates: 0.0120).**

#### Promoter PLux

Stability Tolerance (δ) = 0.6025 (2 *x* Pooled SD of pooled technical replicates across biological replicates: 0.3013)

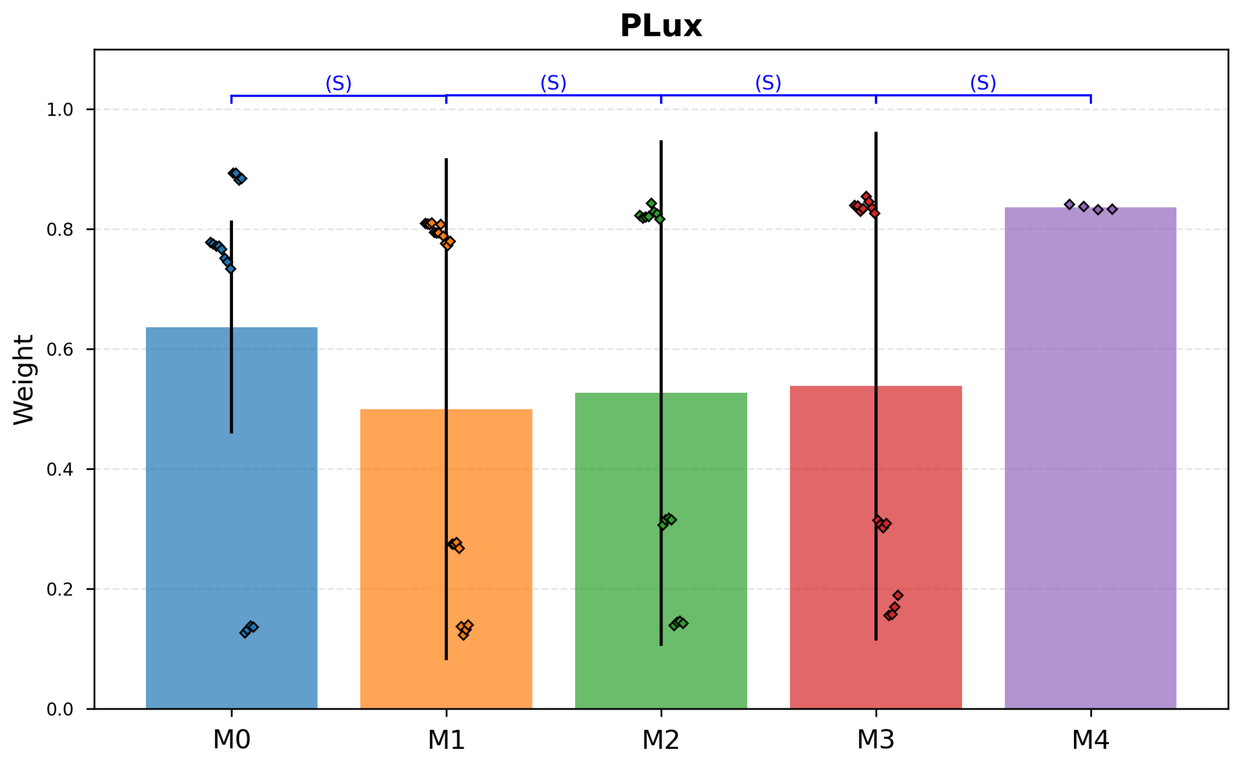

**Supplementary Fig. 52 | Promoter PLux.** Stability Tolerance (δ) = 0.6025 (2 *x* Pooled SD of pooled technical replicates across biological replicates: 0.3013).

- M0 vs M1: Estimated shift Δ = -0.0778. Data inconsistent with drift ≥ ±0.6025 (p_stable = 6.2 × $10^{-6}$). Classified as Stable.
- M1 vs M2: Estimated shift Δ = -0.0320. Data inconsistent with drift ≥ ±0.6025 (p_stable = 2.3 × $10^{-6}$). Classified as Stable.
- M2 vs M3: Estimated shift Δ = 0.0115. Data inconsistent with drift ≥ ±0.6025 (p_stable = 4.7 × $10^{-6}$). Classified as Stable.
- M3 vs M4: Estimated shift Δ = 0.2983. Data inconsistent with drift ≥ ±0.6025 (p_stable = 7.5 × $10^{-4}$). Classified as Stable.

| M-State | Mean Weight | Std Dev |
|---|---|---|
| M0 | 0.6366 | 0.1775 |
| M1 | 0.4997 | 0.4186 |
| M2 | 0.5269 | 0.4216 |
| M3 | 0.5384 | 0.4242 |
| M4 | 0.8367 | n.d. (single entry) |

**Supplementary Table 59 | Stability Tolerance (δ) = 0.6025 (2 *x* Pooled SD of pooled technical replicates across biological replicates: 0.3013).**

#### Promoter PCin

Stability Tolerance (δ) = 0.0460 (2 *x* Pooled SD of pooled technical replicates across biological replicates: 0.0230)

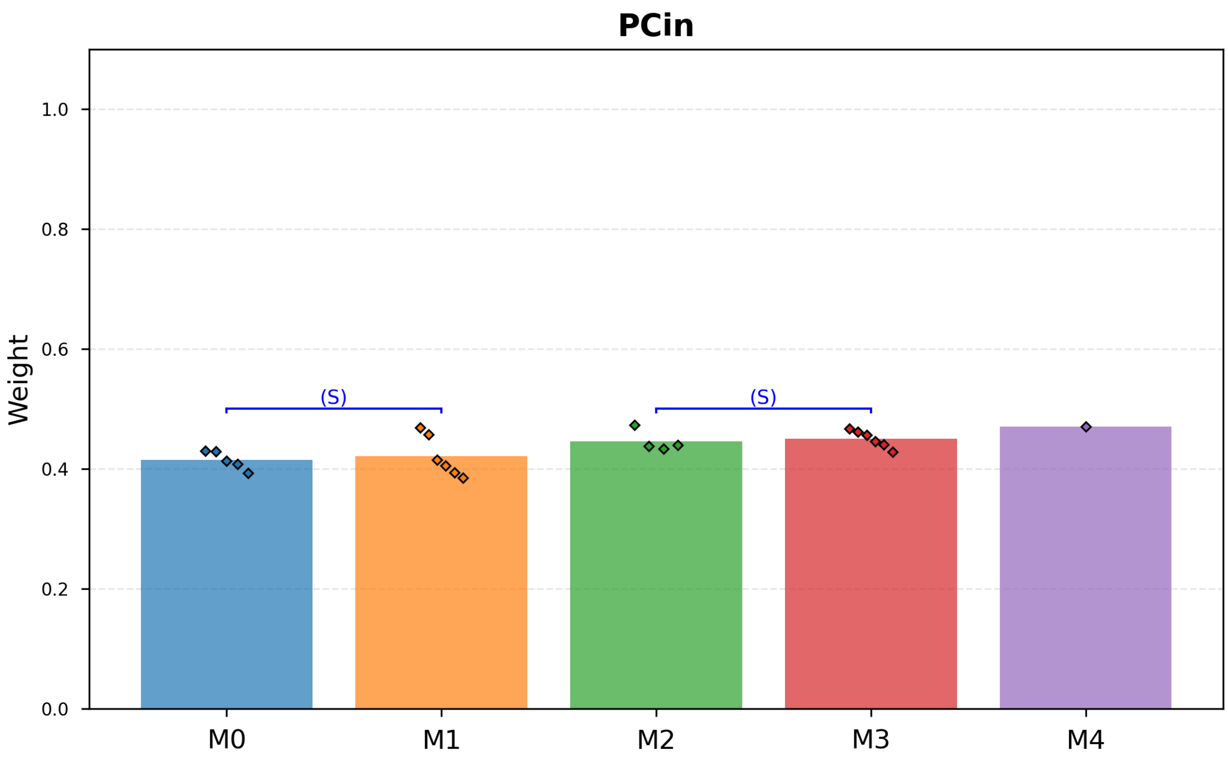

**Supplementary Fig. 53 | Promoter PCin.** Stability Tolerance (δ) = 0.0460 (2 *x* Pooled SD of pooled technical replicates across biological replicates: 0.0230).

- M0 vs M1: Estimated shift Δ = 0.0061. Data inconsistent with drift ≥ ±0.0460 (p_stable = 0.019). Classified as Stable.
- M1 vs M2: Inconclusive (p_stable= 0.128, p_change= 0.173).
- M2 vs M3: Estimated shift Δ = 0.0040. Data inconsistent with drift ≥ ±0.0460 (p_stable = 0.005). Classified as Stable.
- M3 vs M4: Inconclusive (p_stable= 1.000, p_change= 1.000).

| M-State | Mean Weight | Std Dev |
|---|---|---|
| M0 | 0.4148 | n.d. (single entry) |
| M1 | 0.4209 | n.d. (single entry) |
| M2 | 0.4462 | n.d. (single entry) |
| M3 | 0.4502 | n.d. (single entry) |
| M4 | 0.4710 | n.d. (single entry) |

**Supplementary Table 60 | Stability Tolerance (δ) = 0.0460 (2 *x* Pooled SD of pooled technical replicates across biological replicates: 0.0230).**

#### Promoter PTtg

Stability Tolerance (δ) = 0.0274 (2 *x* Pooled SD of pooled technical replicates across biological replicates: 0.0137)

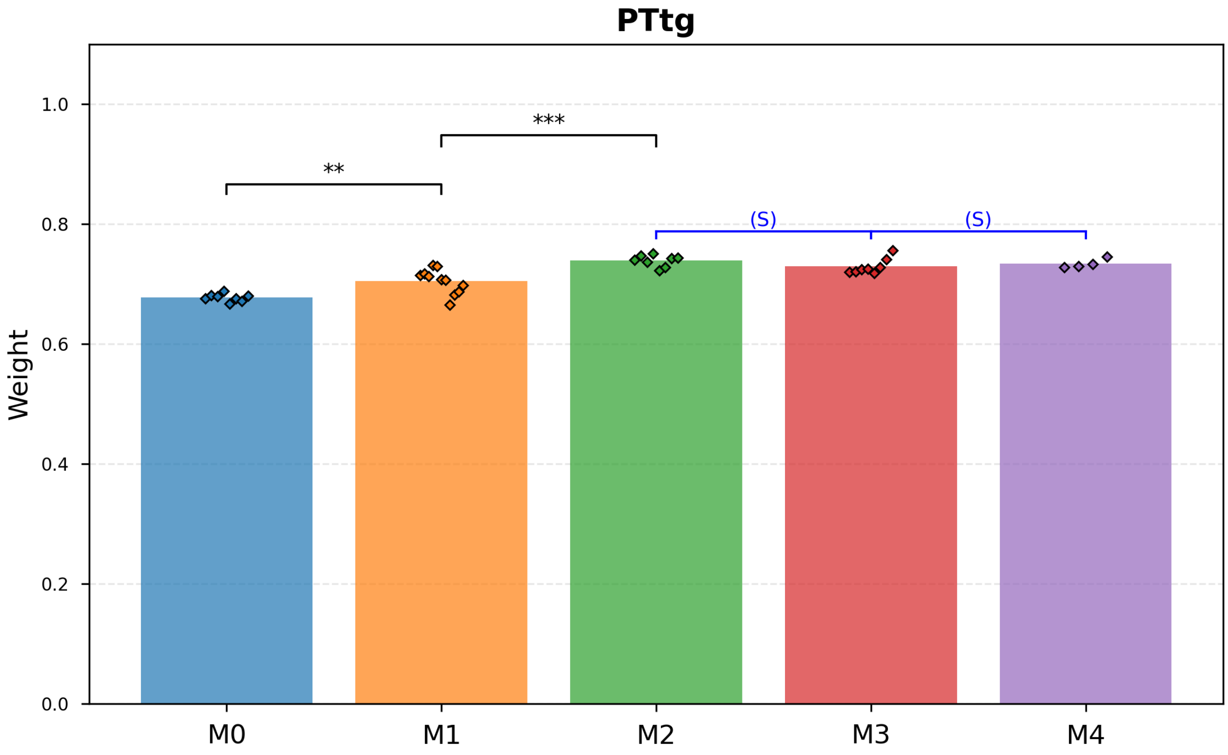

**Supplementary Fig. 54 | Promoter PTtg.** Stability Tolerance (δ) = 0.0274 (2 *x* Pooled SD of pooled technical replicates across biological replicates: 0.0137).

- M0 vs M1: Stability not supported (p_stable = 0.503). Welch's test detected shift (p_change = 0.001). Classified as Drift.
- M1 vs M2: Stability not supported (p_stable = 0.834). Welch's test detected shift (p_change = $1.7 \times 10^{-4}$). Classified as Drift.
- M2 vs M3: Estimated shift Δ = -0.0098. Data inconsistent with drift ≥ ±0.0274 (p_stable = 0.005). Classified as Stable.
- M3 vs M4: Estimated shift Δ = 0.0046. Data inconsistent with drift ≥ ±0.0274 (p_stable = 0.002). Classified as Stable.

| M-State | Mean Weight | Std Dev |
|---|---|---|
| M0 | 0.6775 | n.d. (single entry) |
| M1 | 0.7049 | n.d. (single entry) |
| M2 | 0.7393 | n.d. (single entry) |
| M3 | 0.7295 | n.d. (single entry) |
| M4 | 0.7341 | n.d. (single entry) |

**Supplementary Table 61 | Stability Tolerance (δ) = 0.0274 (2 *x* Pooled SD of pooled technical replicates across biological replicates: 0.0137).**

#### Promoter PVan

Stability Tolerance (δ) = 0.0538 (2 *x* Pooled SD of pooled technical replicates across biological replicates: 0.0269)

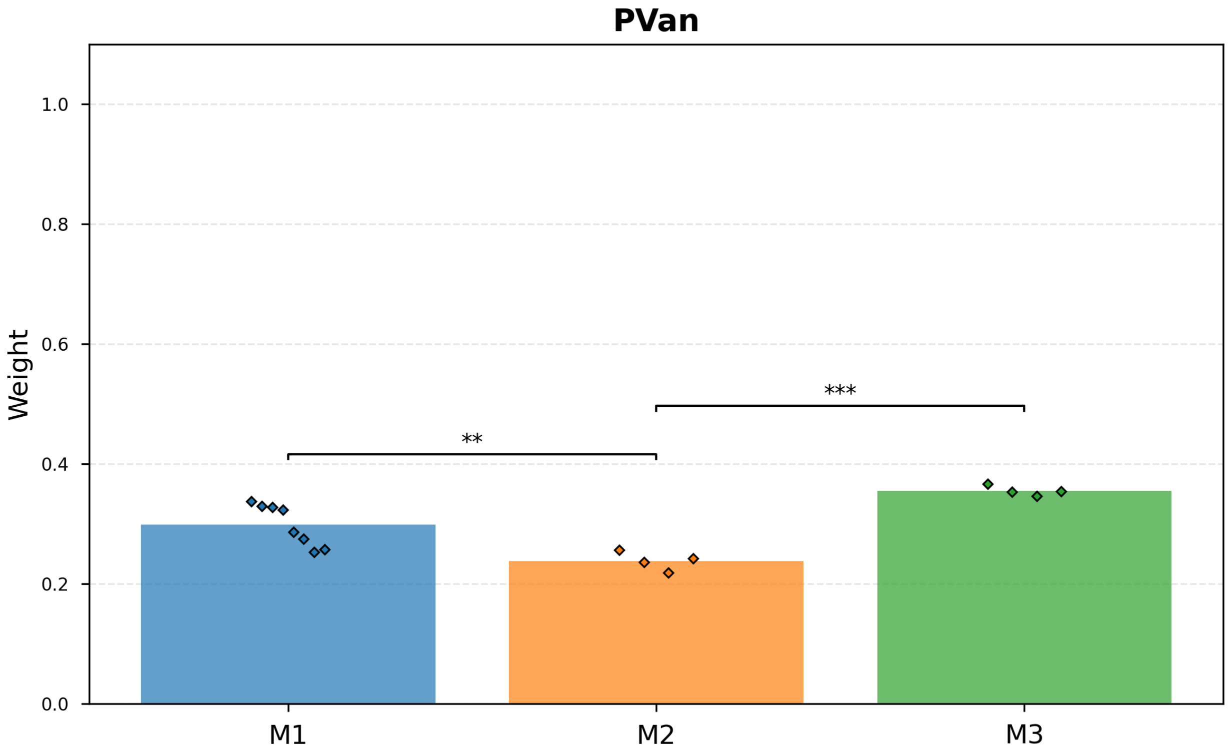

**Supplementary Fig. 55 | Promoter PVan.** Stability Tolerance (δ) = 0.0538 (2 *x* Pooled SD of pooled technical replicates across biological replicates: 0.0269).

- M1 vs M2: Stability not supported (p_stable = 0.680). Welch's test detected shift (p_change = 0.002). Classified as Drift.
- M2 vs M3: Stability not supported (p_stable = 0.999). Welch's test detected shift (p_change = $6.8 \times 10^{-5}$). Classified as Drift.

| M-State | Mean Weight | Std Dev |
|---|---|---|
| M1 | 0.2990 | n.d. (single entry) |
| M2 | 0.2382 | n.d. (single entry) |
| M3 | 0.3551 | n.d. (single entry) |

**Supplementary Table 62 | Stability Tolerance (δ) = 0.0538 (2 *x* Pooled SD of pooled technical replicates across biological replicates: 0.0269).**

#### Promoter PTac

Stability Tolerance (δ) = 0.4208 (2 *x* Pooled SD of pooled technical replicates across biological replicates: 0.2104)

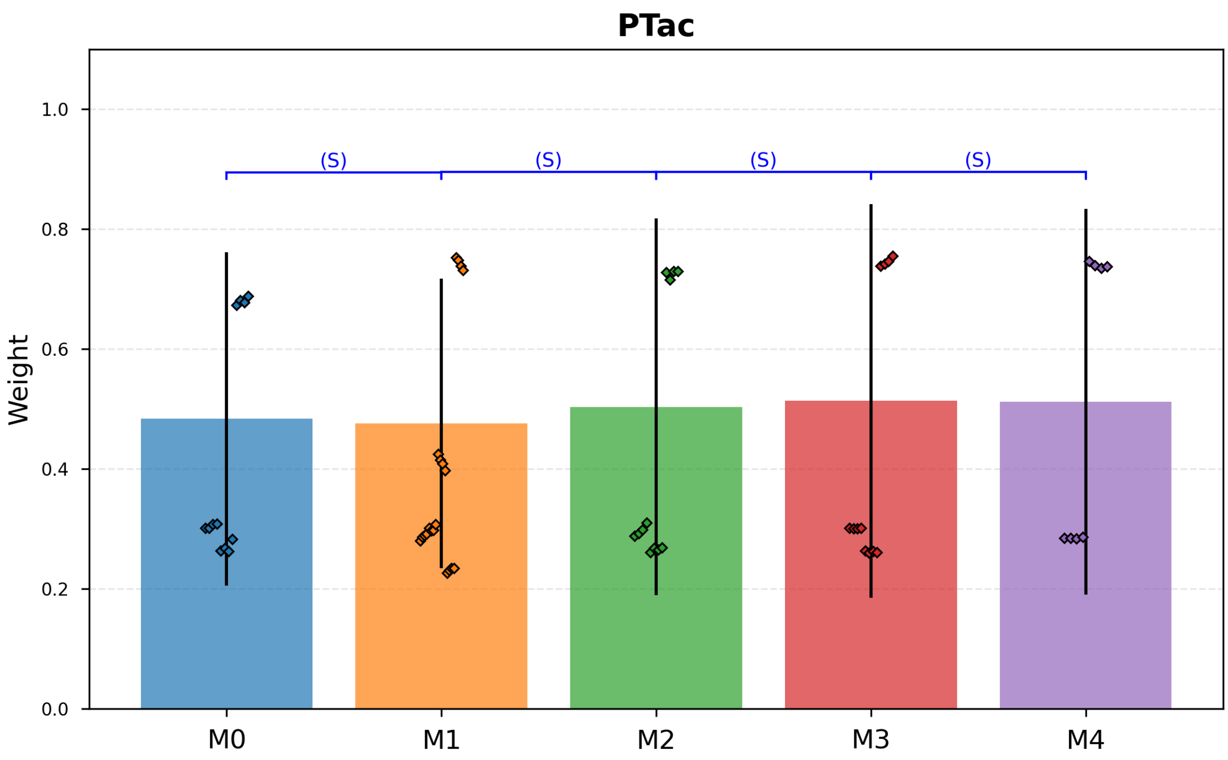

**Supplementary Fig. 56 | Promoter PTac.** Stability Tolerance (δ) = 0.4208 (2 *x* Pooled SD of pooled technical replicates across biological replicates: 0.2104).

- M0 vs M1: Estimated shift Δ = -0.0234. Data inconsistent with drift ≥ ±0.4208 (p_stable = $4.7 \times 10^{-6}$). Classified as Stable.
- M1 vs M2: Estimated shift Δ = 0.0349. Data inconsistent with drift ≥ ±0.4208 (p_stable = $2.7 \times 10^{-5}$). Classified as Stable.
- M2 vs M3: Estimated shift Δ = 0.0067. Data inconsistent with drift ≥ ±0.4208 (p_stable = $8.5 \times 10^{-5}$). Classified as Stable.
- M3 vs M4: Estimated shift Δ = 0.0761. Data inconsistent with drift ≥ ±0.4208 (p_stable = 0.003). Classified as Stable.

| M-State | Mean Weight | Std Dev |
|---|---|---|
| M0 | 0.4837 | 0.2779 |
| M1 | 0.4758 | 0.2414 |
| M2 | 0.5037 | 0.3143 |
| M3 | 0.5137 | 0.3283 |
| M4 | 0.5124 | 0.3216 |

**Supplementary Table 63 | Stability Tolerance (δ) = 0.4208 (2 *x* Pooled SD of pooled technical replicates across biological replicates: 0.2104).**

#### Promoter PVan-Ttg

Stability Tolerance (δ) = 0.0844 (2 *x* Pooled SD of pooled technical replicates across biological replicates: 0.0422)

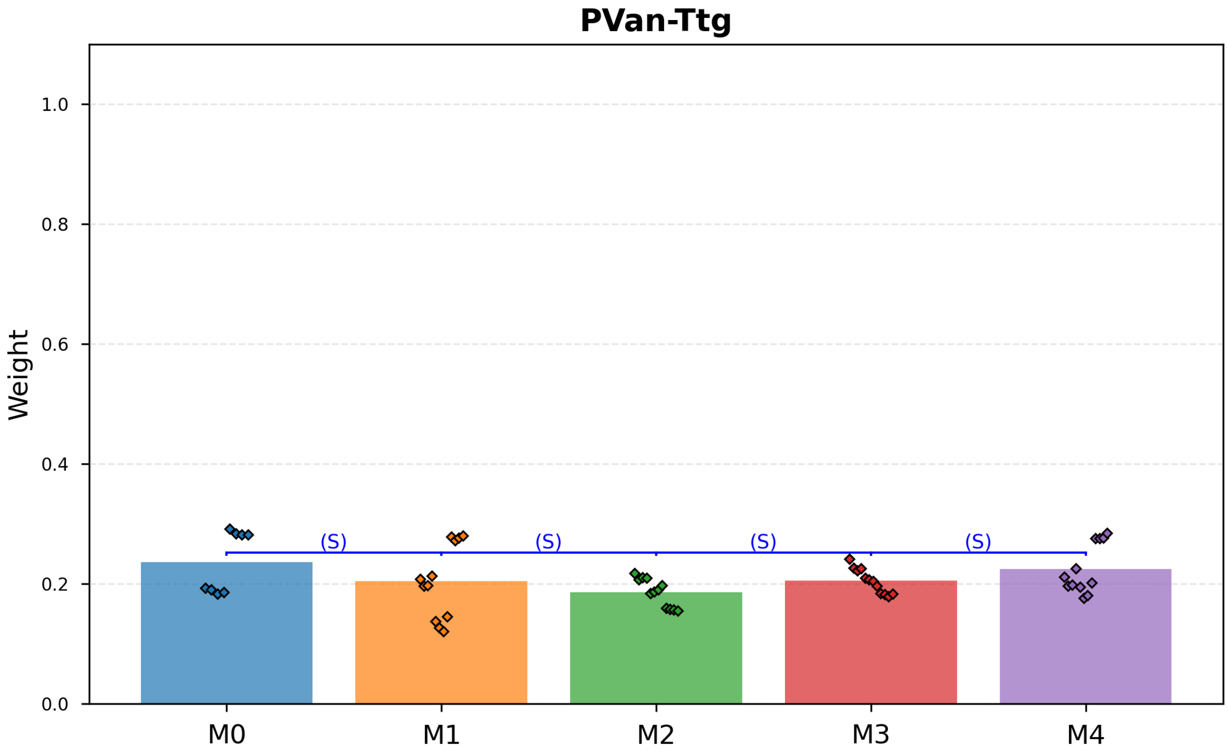

**Supplementary Fig. 57 | Promoter PVan-Ttg.** Stability Tolerance (δ) = 0.0844 (2 *x* Pooled SD of pooled technical replicates across biological replicates: 0.0422).

- M0 vs M1: Estimated shift Δ = -0.0320. Data inconsistent with drift ≥ ±0.0844 (p_stable = 0.029). Classified as Stable.
- M1 vs M2: Estimated shift Δ = -0.0184. Data inconsistent with drift ≥ ±0.0844 (p_stable = 0.002). Classified as Stable.
- M2 vs M3: Estimated shift Δ = 0.0191. Data inconsistent with drift ≥ ±0.0844 (p_stable = 1.7 × $10^{-7}$). Classified as Stable.
- M3 vs M4: Estimated shift Δ = 0.0197. Data inconsistent with drift ≥ ±0.0844 (p_stable = 8.5 × $10^{-5}$). Classified as Stable.

| M-State | Mean Weight | Std Dev |
|---|---|---|
| M0 | 0.2366 | n.d. (single entry) |
| M1 | 0.2047 | n.d. (single entry) |
| M2 | 0.1862 | n.d. (single entry) |
| M3 | 0.2053 | n.d. (single entry) |
| M4 | 0.2250 | n.d. (single entry) |

**Supplementary Table 64 | Stability Tolerance (δ) = 0.0844 (2 *x* Pooled SD of pooled technical replicates across biological replicates: 0.0422).**

#### Promoter PTac-Van

Stability Tolerance (δ) = 0.0867 (2 *x* Pooled SD of pooled technical replicates across biological replicates: 0.0433)

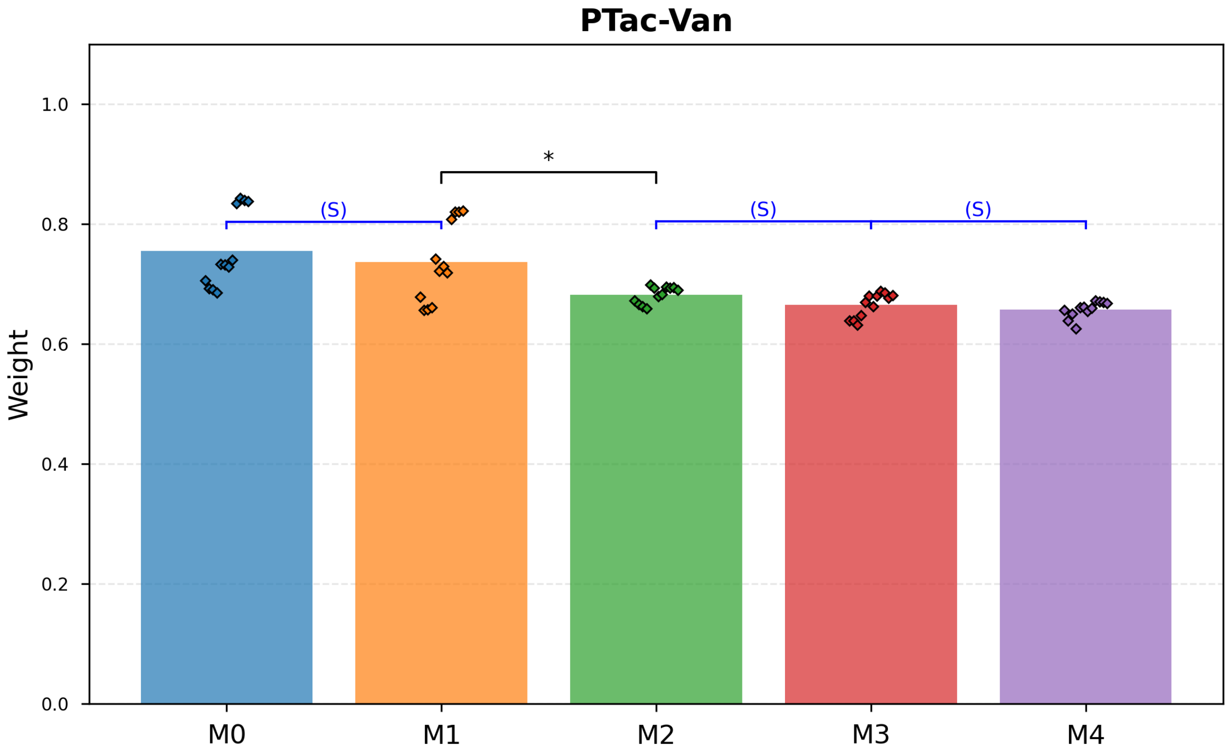

**Supplementary Fig. 58 | Promoter PTac-Van.** Stability Tolerance (δ) = 0.0867 (2 *x* Pooled SD of pooled technical replicates across biological replicates: 0.0433).

- M0 vs M1: Estimated shift Δ = -0.0190. Data inconsistent with drift ≥ ±0.0867 (p_stable = 0.010). Classified as Stable.
- M1 vs M2: Stability not supported (p_stable = 0.063). Welch's test detected shift (p_change = 0.018). Classified as Drift.
- M2 vs M3: Estimated shift Δ = -0.0172. Data inconsistent with drift ≥ ±0.0867 (p_stable = $3.6 \times 10^{-9}$). Classified as Stable.
- M3 vs M4: Estimated shift Δ = -0.0075. Data inconsistent with drift ≥ ±0.0867 (p_stable = $4.2 \times 10^{-10}$). Classified as Stable.

| M-State | Mean Weight | Std Dev |
|---|---|---|
| M0 | 0.7556 | n.d. (single entry) |
| M1 | 0.7366 | n.d. (single entry) |
| M2 | 0.6824 | n.d. (single entry) |
| M3 | 0.6652 | n.d. (single entry) |
| M4 | 0.6577 | n.d. (single entry) |

**Supplementary Table 65 | Stability Tolerance (δ) = 0.0867 (2 *x* Pooled SD of pooled technical replicates across biological replicates: 0.0433).**

#### Promoter PTac-Tet

Stability Tolerance (δ) = 0.0328 (2 *x* Pooled SD of pooled technical replicates across biological replicates: 0.0164)

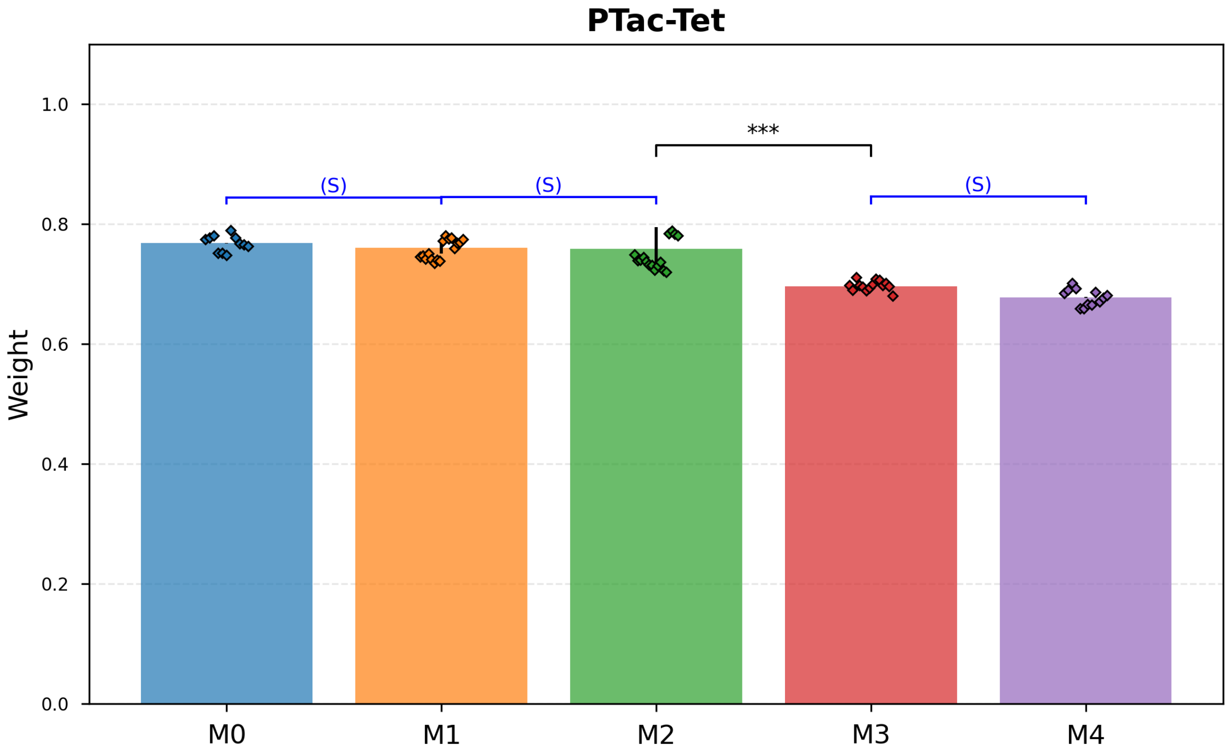

**Supplementary Fig. 59 | Promoter PTac-Tet.** Stability Tolerance (δ) = 0.0328 (2 *x* Pooled SD of pooled technical replicates across biological replicates: 0.0164).

- M0 vs M1: Estimated shift Δ = -0.0107. Data inconsistent with drift ≥ ±0.0328 (p_stable = $4.0 \times 10^{-4}$). Classified as Stable.
- M1 vs M2: Estimated shift Δ = -0.0107. Data inconsistent with drift ≥ ±0.0328 (p_stable = 0.002). Classified as Stable.
- M2 vs M3: Stability not supported (p_stable = 0.991). Welch's test detected shift (p_change = $2.7 \times 10^{-7}$). Classified as Drift.
- M3 vs M4: Estimated shift Δ = -0.0199. Data inconsistent with drift ≥ ±0.0328 (p_stable = 0.005). Classified as Stable.

| M-State | Mean Weight | Std Dev |
|---|---|---|
| M0 | 0.7683 | 0.0005 |
| M1 | 0.7610 | 0.0100 |
| M2 | 0.7593 | 0.0354 |
| M3 | 0.6966 | 0.0037 |
| M4 | 0.6781 | 0.0010 |

**Supplementary Table 66 | Stability Tolerance (δ) = 0.0328 (2 *x* Pooled SD of pooled technical replicates across biological replicates: 0.0164).**

#### Promoter PTet-Ttg

Stability Tolerance (δ) = 0.0638 (2 *x* Pooled SD of pooled technical replicates across biological replicates: 0.0319)

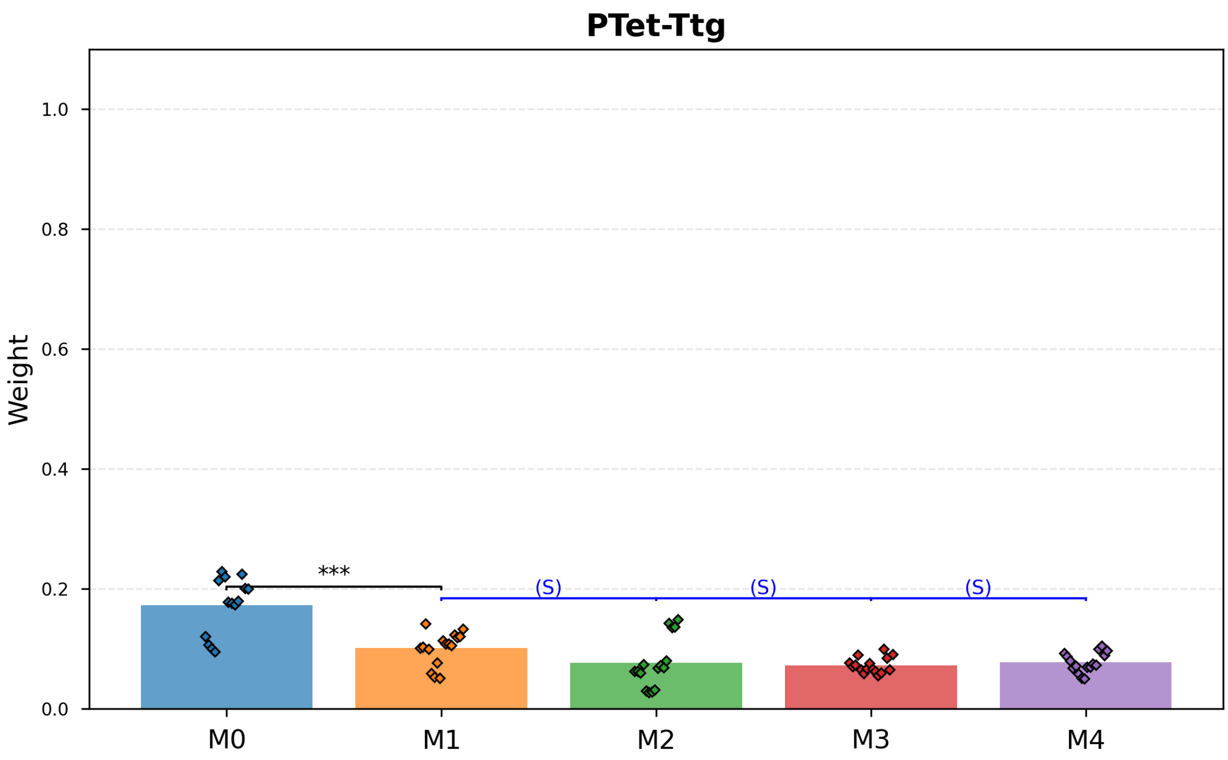

**Supplementary Fig. 60 | Promoter PTet-Ttg.** Stability Tolerance (δ) = 0.0638 (2 *x* Pooled SD of pooled technical replicates across biological replicates: 0.0319).

- M0 vs M1: Stability not supported (p_stable = 0.703). Welch's test detected shift (p_change = $7.9 \times 10^{-5}$). Classified as Drift.
- M1 vs M2: Estimated shift Δ = -0.0244. Data inconsistent with drift ≥ ±0.0638 (p_stable = 0.002). Classified as Stable.
- M2 vs M3: Estimated shift Δ = -0.0042. Data inconsistent with drift ≥ ±0.0638 (p_stable = $2.0 \times 10^{-5}$). Classified as Stable.
- M3 vs M4: Estimated shift Δ = 0.0046. Data inconsistent with drift ≥ ±0.0638 (p_stable = $3.0 \times 10^{-12}$). Classified as Stable.

| M-State | Mean Weight | Std Dev |
|---|---|---|
| M0 | 0.1730 | n.d. (single entry) |
| M1 | 0.1012 | n.d. (single entry) |
| M2 | 0.0768 | n.d. (single entry) |
| M3 | 0.0727 | n.d. (single entry) |
| M4 | 0.0773 | n.d. (single entry) |

**Supplementary Table 67 | Stability Tolerance (δ) = 0.0638 (2 *x* Pooled SD of pooled technical replicates across biological replicates: 0.0319).**

### Supplementary Note 10: Weight learning from plate reader fluorescence

#### *Summary Table*

| Cell Name | Growth rate μ (1/h) | Green fluorescence | Red fluorescence | Weight | Replicates |
|---|---|---|---|---|---|
| PSal-M0 | 0.48 ± 0.05 | 21,903 ± 1,265 | 27,245 ± 1,830 | 0.65 ± 0.04 | 7 |
| PSal-M1 | 0.425 ± 0.010 | 28,029 ± 771 | 21,064 ± 466 | 0.498 ± 0.004 | 33 |
| PSal-M2 | 0.436 ± 0.013 | 31,284 ± 777 | 15,489 ± 379 | 0.380 ± 0.006 | 34 |
| PSal-M3 | 0.409 ± 0.013 | 35,237 ± 688 | 10,856 ± 142 | 0.2682 ± 0.0029 | 34 |
| PSal-M4 | 0.434 ± 0.012 | 39,647 ± 1,134 | 7,773 ± 207 | 0.1852 ± 0.0022 | 33 |
| PSal-M5 | 0.393 ± 0.015 | 41,945 ± 1,240 | 4,539 ± 83 | 0.1099 ± 0.0017 | 19 |
| PTet-M0 | 0.502 ± 0.016 | 25,287 ± 413 | 21,169 ± 377 | 0.6794 ± 0.0018 | 12 |
| PTet-M1 | 0.538 ± 0.005 | 26,787 ± 430 | 23,667 ± 320 | 0.6946 ± 0.0025 | 22 |
| PTet-M2 | 0.522 ± 0.005 | 30,486 ± 392 | 21,708 ± 382 | 0.633 ± 0.004 | 24 |
| PTet-M3 | 0.500 ± 0.007 | 31,229 ± 387 | 19,570 ± 238 | 0.597 ± 0.004 | 24 |
| PTet-M4 | 0.503 ± 0.007 | 31,415 ± 408 | 16,265 ± 332 | 0.541 ± 0.006 | 24 |
| PTet-M5 | 0.506 ± 0.007 | 22,796 ± 325 | 9,344 ± 297 | 0.474 ± 0.008 | 23 |
| PBetI-M0 | 0.462 ± 0.020 | 15,660 ± 172 | 13,315 ± 155 | 1.1372 ± 0.0005 | 8 |
| PBetI-M1 | 0.456 ± 0.017 | 14,791 ± 274 | 8,859 ± 165 | 1.0284 ± 0.0024 | 16 |
| PBetI-M2 | 0.459 ± 0.017 | 19,904 ± 405 | 8,145 ± 201 | 0.893 ± 0.010 | 15 |
| PBetI-M3 | 0.435 ± 0.021 | 18,990 ± 947 | 7,454 ± 310 | 0.880 ± 0.012 | 11 |
| PBetI-M4 | 0.463 ± 0.010 | 21,177 ± 1,482 | 6,581 ± 461 | 0.788 ± 0.017 | 15 |
| PBAD-M0 | 0.41 ± 0.03 | 24,144 ± 435 | 53,295 ± 856 | 0.6731 ± 0.0010 | 11 |
| PBAD-M1 | 0.487 ± 0.012 | 24,906 ± 492 | 52,848 ± 993 | 0.6622 ± 0.0008 | 11 |
| PBAD-M2 | 0.501 ± 0.005 | 28,266 ± 465 | 43,324 ± 517 | 0.571 ± 0.004 | 11 |
| PBAD-M3 | 0.479 ± 0.005 | 30,565 ± 1,735 | 37,713 ± 2,546 | 0.508 ± 0.006 | 12 |
| PBAD-M4 | 0.513 ± 0.024 | 32,842 ± 661 | 17,429 ± 727 | 0.293 ± 0.006 | 12 |
| PBAD-M5 | 0.490 ± 0.005 | 32,911 ± 683 | 6,225 ± 284 | 0.125 ± 0.003 | 11 |
| PLux-M0 | 0.392 ± 0.018 | 13,714 ± 369 | 59,004 ± 1,636 | 0.9431 ± 0.0029 | 7 |
| PLux-M1 | 0.434 ± 0.014 | 24,346 ± 1,609 | 44,793 ± 1,152 | 0.720 ± 0.028 | 33 |
| PLux-M2 | 0.469 ± 0.013 | 27,915 ± 1,737 | 29,958 ± 1,440 | 0.55 ± 0.03 | 34 |
| PLux-M3 | 0.443 ± 0.015 | 26,725 ± 1,331 | 25,994 ± 1,012 | 0.510 ± 0.023 | 24 |
| PLux-M4 | 0.537 ± 0.015 | 29,896 ± 1,140 | 16,299 ± 257 | 0.349 ± 0.011 | 26 |
| PLux-M5 | 0.461 ± 0.019 | 33,031 ± 1,648 | 10,300 ± 650 | 0.231 ± 0.022 | 16 |
| PCin-M0 | 0.408 ± 0.012 | 10,462 ± 591 | 22,444 ± 715 | 0.879 ± 0.009 | 11 |
| PCin-M1 | 0.426 ± 0.013 | 13,477 ± 752 | 11,973 ± 472 | 0.586 ± 0.021 | 6 |
| PCin-M2 | 0.446 ± 0.007 | 12,564 ± 521 | 6,983 ± 235 | 0.458 ± 0.013 | 18 |
| PCin-M3 | 0.428 ± 0.005 | 13,802 ± 256 | 4,436 ± 104 | 0.309 ± 0.005 | 15 |
| PCin-M4 | 0.412 ± 0.008 | 16,621 ± 648 | 2,828 ± 82 | 0.1860 ± 0.0028 | 28 |
| PTtg-M0 | 0.448 ± 0.027 | 19,824 ± 799 | 4,914 ± 325 | 0.736 ± 0.013 | 8 |
| PTtg-M1 | 0.506 ± 0.006 | 21,751 ± 466 | 5,238 ± 151 | 0.729 ± 0.007 | 28 |
| PTtg-M2 | 0.492 ± 0.007 | 25,231 ± 412 | 4,292 ± 77 | 0.630 ± 0.006 | 40 |
| PTtg-M3 | 0.467 ± 0.011 | 29,450 ± 480 | 4,089 ± 72 | 0.570 ± 0.007 | 37 |
| PTtg-M4 | 0.492 ± 0.007 | 36,565 ± 446 | 2,621 ± 64 | 0.382 ± 0.006 | 39 |
| PTtg-M5 | 0.463 ± 0.010 | 36,034 ± 1,397 | 3,136 ± 136 | 0.437 ± 0.020 | 15 |
| PVan-M0 | 0.422 ± 0.026 | 28,938 ± 3,648 | 61,278 ± 10,402 | 0.757 ± 0.012 | 2 |
| PVan-M1 | 0.475 ± 0.015 | 20,821 ± 450 | 48,054 ± 916 | 0.7829 ± 0.0017 | 6 |
| PVan-M2 | 0.471 ± 0.007 | 18,129 ± 292 | 44,682 ± 843 | 0.800 ± 0.005 | 35 |
| PVan-M3 | 0.454 ± 0.011 | 19,111 ± 928 | 24,609 ± 1,546 | 0.603 ± 0.013 | 35 |
| PVan-M4 | 0.460 ± 0.010 | 10,993 ± 392 | 4,003 ± 524 | 0.246 ± 0.024 | 33 |
| PVan-M5 | 0.427 ± 0.006 | 9,755 ± 221 | 141 ± 79 | 0.011 ± 0.007 | 17 |
| PTac-M0 | 0.453 ± 0.010 | 23,710 ± 1,610 | 36,457 ± 704 | 0.727 ± 0.016 | 17 |
| PTac-M1 | 0.453 ± 0.010 | 23,331 ± 774 | 31,008 ± 931 | 0.668 ± 0.020 | 54 |
| PTac-M2 | 0.455 ± 0.010 | 23,944 ± 297 | 24,323 ± 358 | 0.6184 ± 0.0017 | 28 |
| PTac-M3 | 0.473 ± 0.008 | 13,776 ± 542 | 1,997 ± 128 | 0.173 ± 0.006 | 27 |
| PTac-M4 | 0.460 ± 0.006 | 12,017 ± 494 | -88 ± 11 | -0.0110 ± 0.0013 | 17 |
| PVan-Ttg-M0 | 0.465 ± 0.017 | 39,387 ± 1,454 | 18,901 ± 966 | 0.775 ± 0.010 | 14 |
| PVan-Ttg-M1 | 0.455 ± 0.020 | 12,554 ± 514 | 12,540 ± 1,771 | 0.823 ± 0.023 | 27 |
| PVan-Ttg-M2 | 0.440 ± 0.024 | 9,806 ± 762 | 2,474 ± 512 | 0.40 ± 0.08 | 27 |
| PVan-Ttg-M3 | 0.449 ± 0.018 | 10,520 ± 771 | 7 ± 28 | 0.007 ± 0.019 | 24 |
| PVan-Ttg-M4 | 0.460 ± 0.008 | 11,191 ± 722 | -109 ± 8 | -0.075 ± 0.006 | 25 |
| PTac-Tet-M0 | 0.34 ± 0.09 | 15,458 ± 793 | 28,749 ± 1,329 | 0.7930 ± 0.0011 | 4 |
| PTac-Tet-M1 | 0.377 ± 0.009 | 29,794 ± 877 | 46,987 ± 1,334 | 0.761 ± 0.008 | 24 |
| PTac-Tet-M2 | 0.349 ± 0.018 | 32,769 ± 3,300 | 53,917 ± 5,323 | 0.770 ± 0.006 | 24 |
| PTac-Tet-M3 | 0.358 ± 0.010 | 36,890 ± 1,625 | 37,086 ± 2,205 | 0.665 ± 0.008 | 27 |
| PTac-Tet-M4 | 0.375 ± 0.009 | 54,578 ± 1,754 | 21,630 ± 951 | 0.437 ± 0.017 | 29 |
| PTet-Ttg-M0 | 0.533 ± 0.005 | 20,898 ± 938 | 11,097 ± 224 | 0.651 ± 0.010 | 6 |
| PTet-Ttg-M1 | 0.493 ± 0.008 | 17,044 ± 1,690 | 5,288 ± 1,081 | 0.32 ± 0.06 | 24 |

| | | | | | |
|---|---|---|---|---|---|
| PTet-Ttg-M2 | 0.484 ± 0.006 | 9,531 ± 319 | -4 ± 11 | -0.003 ± 0.004 | 42 |
| PTet-Ttg-M3 | 0.460 ± 0.007 | 10,187 ± 442 | -102 ± 10 | -0.038 ± 0.003 | 35 |
| PTet-Ttg-M4 | 0.489 ± 0.013 | 9,760 ± 535 | -164 ± 5 | -0.074 ± 0.009 | 36 |

**Supplementary Table 68 | Summary Table.**

#### Analysis of growth and fluorescence curves from plate reader

This section presents a comprehensive analysis of cellular growth and fluorescence dynamics, aggregated across all experimental datasets. The data is organized by promoter system and subsequently sorted by the experimental condition identifier (M-suffix) to provide a clear, logical progression. Within this framework, we distinguish between two levels of replication fundamental to biological experiments:

- Biological Replicates: These are distinct cultures, often originating from different colonies or clones (e.g., 'Clone A' and 'Clone B'), that are subjected to the same experimental conditions. They are treated as independent measurements of the biological process under investigation.
- Technical Replicates: These are multiple measurements taken from the same biological sample (e.g., multiple wells in a plate inoculated from the same starter culture). They serve to quantify the precision and variability of the measurement technique itself.

Throughout this analysis, each curve representing a biological replicate is computed as the mean of its corresponding technical replicates. The variability among these technical replicates is visualized as a shaded region representing the standard error of the mean (SEM). All mean curves are linearly interpolated between measured time points to ensure visual continuity.

#### Weight variation analysis

This section presents bar charts grouped by promoter, with bars ordered by M-suffix (memory state). Each bar represents the mean weight across biological replicates for that promoter-M combination. Individual technical replicate values are shown as diamond markers. Error bars represent standard deviation across biological replicates. Statistical significance between consecutive M-suffixes is indicated by brackets (* $p<0.05$, ** $p<0.01$, *** $p<0.001$).

##### Promoter PSal

Stability Tolerance (δ) = 0.0586 (2 *x* Pooled SD: 0.0293)

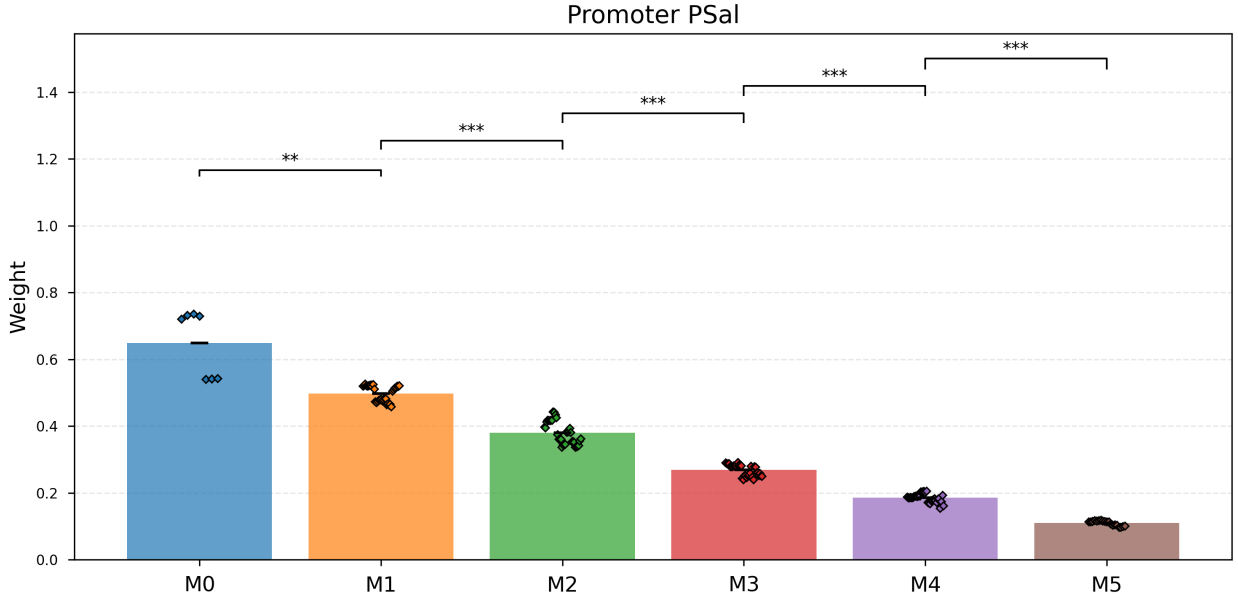

**Supplementary Fig. 61 | Promoter PSal.** Stability Tolerance (δ) = 0.0586 (2 *x* Pooled SD: 0.0293).

- M0 vs M1: Drift (p_change= 0.007).

- M1 vs M2: Drift (p_change= $6.6 \times 10^{-24}$).
- M2 vs M3: Drift (p_change= $1.9 \times 10^{-22}$).
- M3 vs M4: Drift (p_change= $2.6 \times 10^{-31}$).
- M4 vs M5: Drift (p_change= $1.2 \times 10^{-31}$).

| M-State | Mean Weight | Std Dev |
|---|---|---|
| M0 | 0.65 | 0.04 |
| M1 | 0.498 | 0.004 |
| M2 | 0.380 | 0.006 |
| M3 | 0.2682 | 0.0029 |
| M4 | 0.1852 | 0.0022 |
| M5 | 0.1099 | 0.0017 |

**Supplementary Table 69 | Stability Tolerance (δ) = 0.0586 (2 *x* Pooled SD: 0.0293).**

#### Promoter PTet

Stability Tolerance (δ) = 0.0465 (2 *x* Pooled SD: 0.0232)

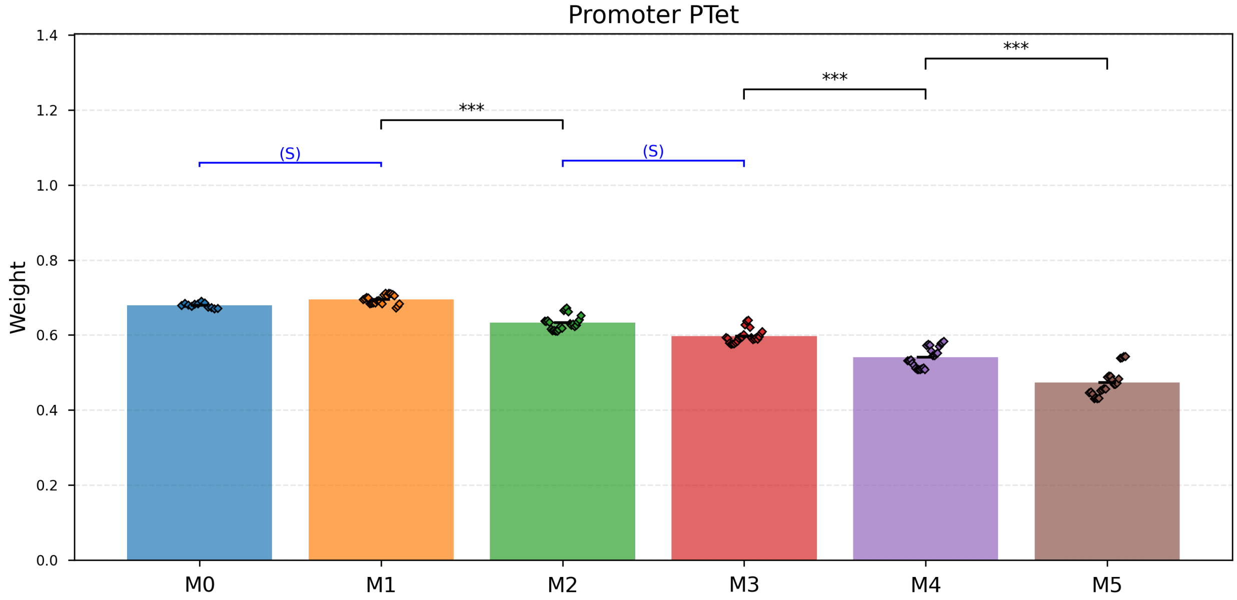

**Supplementary Fig. 62 | Promoter PTet.** Stability Tolerance (δ) = 0.0465 (2 *x* Pooled SD: 0.0232).

- M0 vs M1: Stable within ±δ (p_stable= $1.0 \times 10^{-11}$).
- M1 vs M2: Drift (p_change= $5.9 \times 10^{-16}$).
- M2 vs M3: Stable within ±δ (p_stable= 0.031).
- M3 vs M4: Drift (p_change= $2.9 \times 10^{-10}$).
- M4 vs M5: Drift (p_change= $1.2 \times 10^{-8}$).

| M-State | Mean Weight | Std Dev |
|---|---|---|
| M0 | 0.6794 | 0.0018 |
| M1 | 0.6946 | 0.0025 |
| M2 | 0.633 | 0.004 |
| M3 | 0.597 | 0.004 |
| M4 | 0.541 | 0.006 |
| M5 | 0.474 | 0.008 |

**Supplementary Table 70 | Stability Tolerance (δ) = 0.0465 (2 *x* Pooled SD: 0.0232).**

#### Promoter PBetI

Stability Tolerance (δ) = 0.0813 (2 *x* Pooled SD: 0.0406)

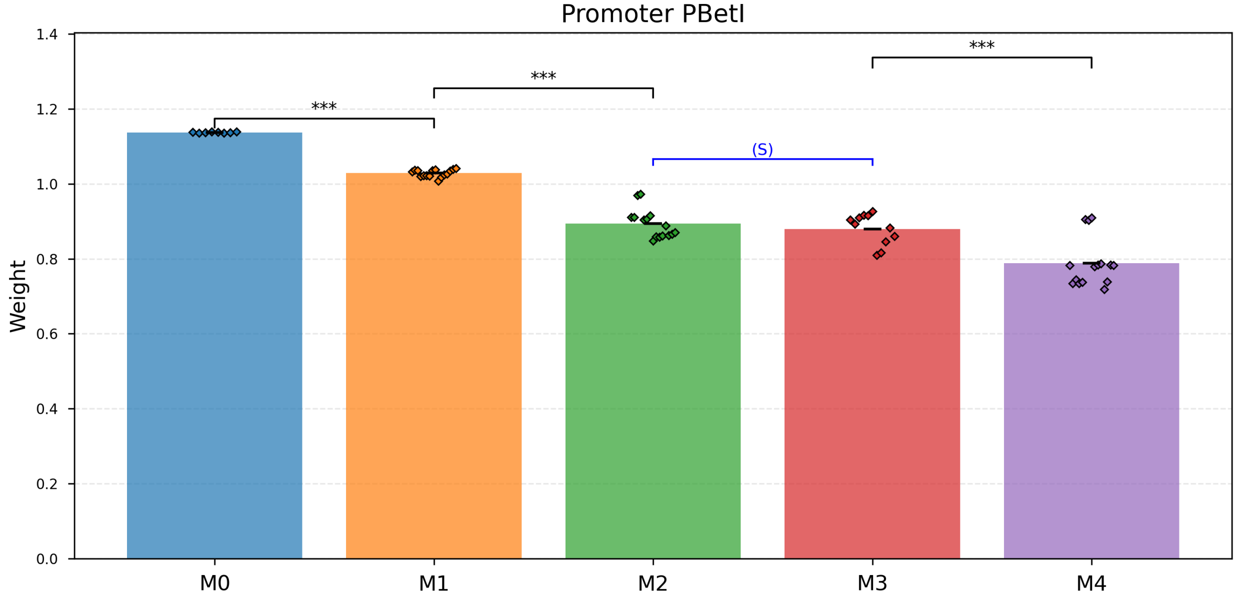

**Supplementary Fig. 63 | Promoter PBetI.** Stability Tolerance (δ) = 0.0813 (2 *x* Pooled SD: 0.0406).

- M0 vs M1: Drift (p_change= $3.2 \times 10^{-18}$).
- M1 vs M2: Drift (p_change= $8.0 \times 10^{-10}$).
- M2 vs M3: Stable within ±δ (p_stable= $1.8 \times 10^{-4}$).
- M3 vs M4: Drift (p_change= $2.1 \times 10^{-4}$).

| M-State | Mean Weight | Std Dev |
|---|---|---|
| M0 | 1.1372 | 0.0005 |
| M1 | 1.0284 | 0.0024 |
| M2 | 0.893 | 0.010 |
| M3 | 0.880 | 0.012 |
| M4 | 0.788 | 0.017 |

**Supplementary Table 71 | Stability Tolerance (δ) = 0.0813 (2 *x* Pooled SD: 0.0406).**

#### Promoter PBAD

Stability Tolerance (δ) = 0.0300 (2 *x* Pooled SD: 0.0150)

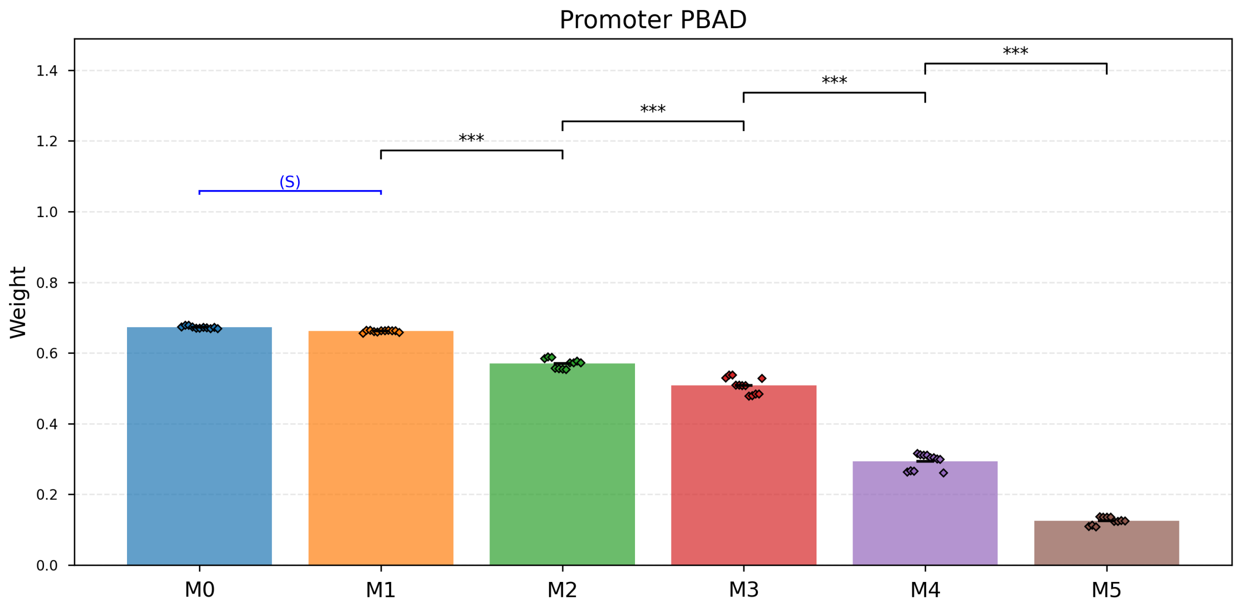

**Supplementary Fig. 64 | Promoter PBAD.** Stability Tolerance (δ) = 0.0300 (2 *x* Pooled SD: 0.0150).

- M0 vs M1: Stable within ±δ (p_stable= $1.4 \times 10^{-12}$).
- M1 vs M2: Drift (p_change= $2.7 \times 10^{-10}$).
- M2 vs M3: Drift (p_change= $1.4 \times 10^{-7}$).
- M3 vs M4: Drift (p_change= $3.5 \times 10^{-17}$).
- M4 vs M5: Drift (p_change= $4.9 \times 10^{-14}$).

| M-State | Mean Weight | Std Dev |
|---|---|---|
| M0 | 0.6731 | 0.0010 |
| M1 | 0.6622 | 0.0008 |
| M2 | 0.571 | 0.004 |
| M3 | 0.508 | 0.006 |
| M4 | 0.293 | 0.006 |
| M5 | 0.125 | 0.003 |

**Supplementary Table 72 | Stability Tolerance (δ) = 0.0300 (2 *x* Pooled SD: 0.0150).**

#### Promoter PLux

Stability Tolerance (δ) = 0.2750 (2 *x* Pooled SD: 0.1375)

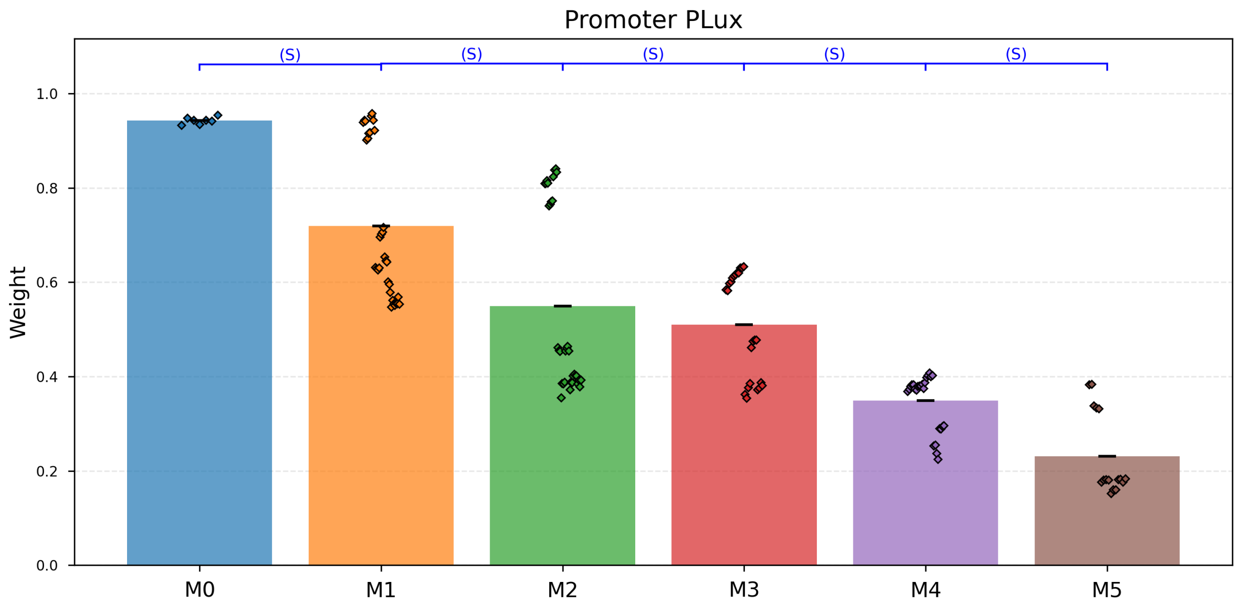

**Supplementary Fig. 65 | Promoter PLux.** Stability Tolerance (δ) = 0.2750 (2 *x* Pooled SD: 0.1375).

- M0 vs M1: Stable within ±δ (p_stable= 0.036).
- M1 vs M2: Stable within ±δ (p_stable= 0.009).
- M2 vs M3: Stable within ±δ (p_stable= $1.5 \times 10^{-7}$).
- M3 vs M4: Stable within ±δ (p_stable= $3.6 \times 10^{-5}$).
- M4 vs M5: Stable within ±δ (p_stable= $8.7 \times 10^{-7}$).

| M-State | Mean Weight | Std Dev |
|---|---|---|
| M0 | 0.9431 | 0.0029 |
| M1 | 0.720 | 0.028 |
| M2 | 0.55 | 0.03 |
| M3 | 0.510 | 0.023 |
| M4 | 0.349 | 0.011 |
| M5 | 0.231 | 0.022 |

**Supplementary Table 73 | Stability Tolerance (δ) = 0.2750 (2 *x* Pooled SD: 0.1375).**

#### Promoter PCin

Stability Tolerance (δ) = 0.0638 (2 *x* Pooled SD: 0.0319)

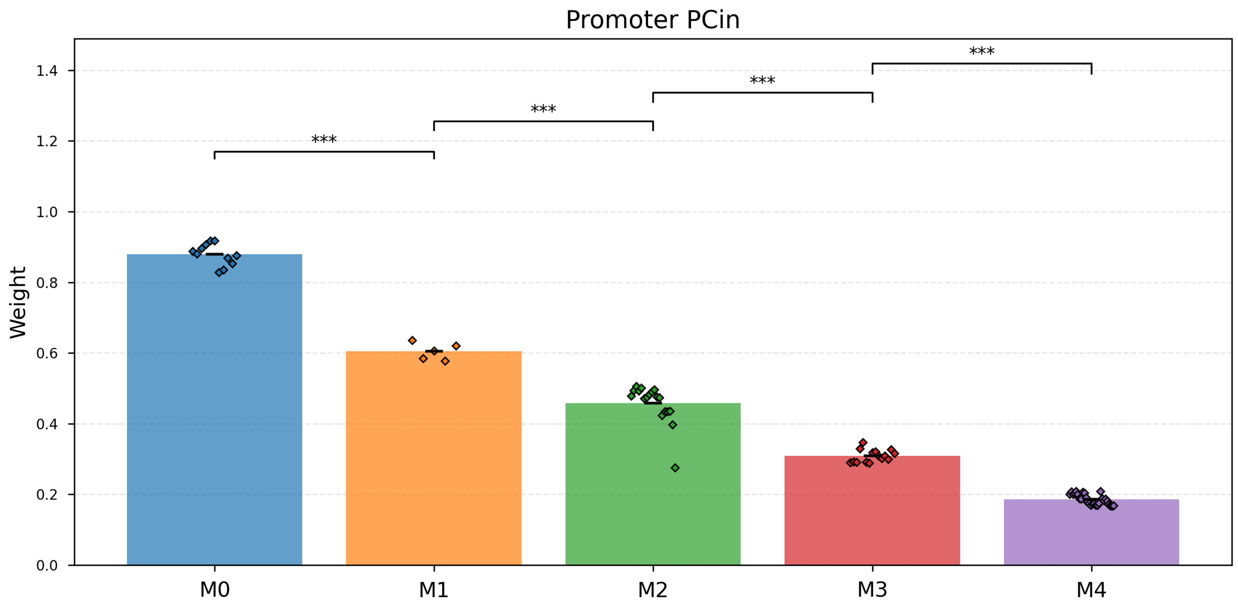

**Supplementary Fig. 66 | Promoter PCin.** Stability Tolerance (δ) = 0.0638 (2 *x* Pooled SD: 0.0319).

- M0 vs M1: Drift (p_change= $4.7 \times 10^{-9}$).
- M1 vs M2: Drift (p_change= $2.1 \times 10^{-7}$).
- M2 vs M3: Drift (p_change= $4.2 \times 10^{-10}$).
- M3 vs M4: Drift (p_change= $3.9 \times 10^{-18}$).

| M-State | Mean Weight | Std Dev |
|---|---|---|
| M0 | 0.879 | 0.009 |
| M1 | 0.586 | 0.021 |
| M2 | 0.458 | 0.013 |
| M3 | 0.309 | 0.005 |
| M4 | 0.1860 | 0.0028 |

**Supplementary Table 74 | Stability Tolerance (δ) = 0.0638 (2 *x* Pooled SD: 0.0319).**

#### Promoter PTtg

Stability Tolerance (δ) = 0.0869 (2 *x* Pooled SD: 0.0435)

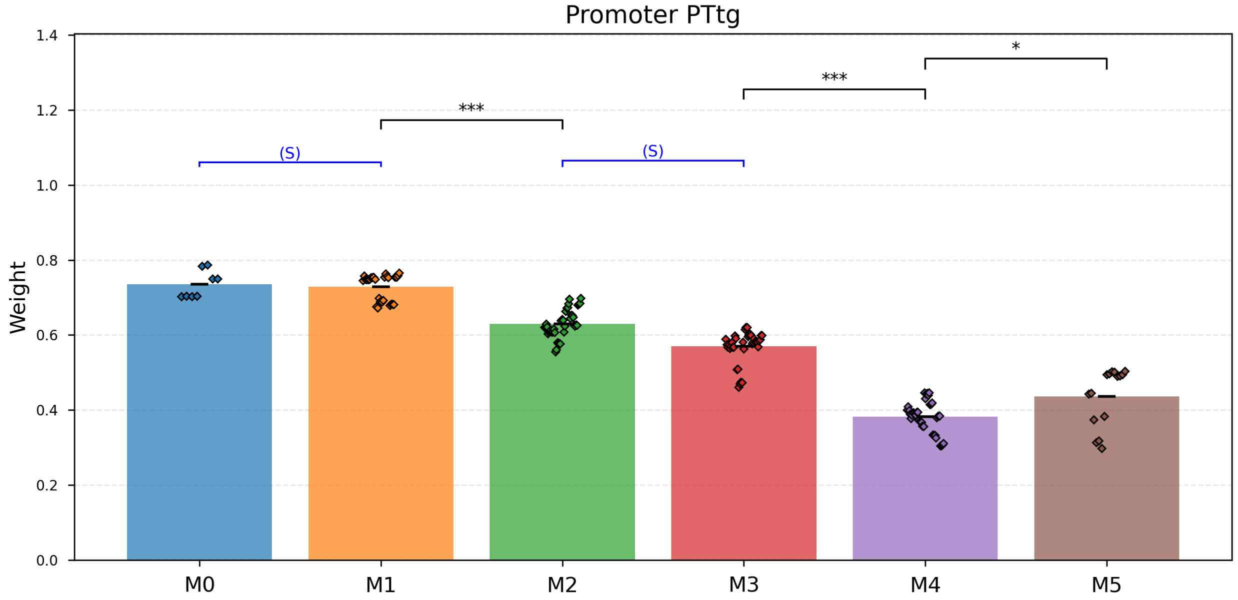

**Supplementary Fig. 67 | Promoter PTtg.** Stability Tolerance (δ) = 0.0869 (2 *x* Pooled SD: 0.0435).

- M0 vs M1: Stable within ±δ (p_stable= $1.0 \times 10^{-4}$).
- M1 vs M2: Drift (p_change= $1.2 \times 10^{-16}$).
- M2 vs M3: Stable within ±δ (p_stable= 0.002).
- M3 vs M4: Drift (p_change= $3.3 \times 10^{-31}$).
- M4 vs M5: Drift (p_change= 0.019).

| M-State | Mean Weight | Std Dev |
|---|---|---|
| M0 | 0.736 | 0.013 |
| M1 | 0.729 | 0.007 |
| M2 | 0.630 | 0.006 |
| M3 | 0.570 | 0.007 |
| M4 | 0.382 | 0.006 |
| M5 | 0.437 | 0.020 |

**Supplementary Table 75 | Stability Tolerance (δ) = 0.0869 (2 *x* Pooled SD: 0.0435).**

#### Promoter PVan

Stability Tolerance (δ) = 0.1741 (2 *x* Pooled SD: 0.0871)

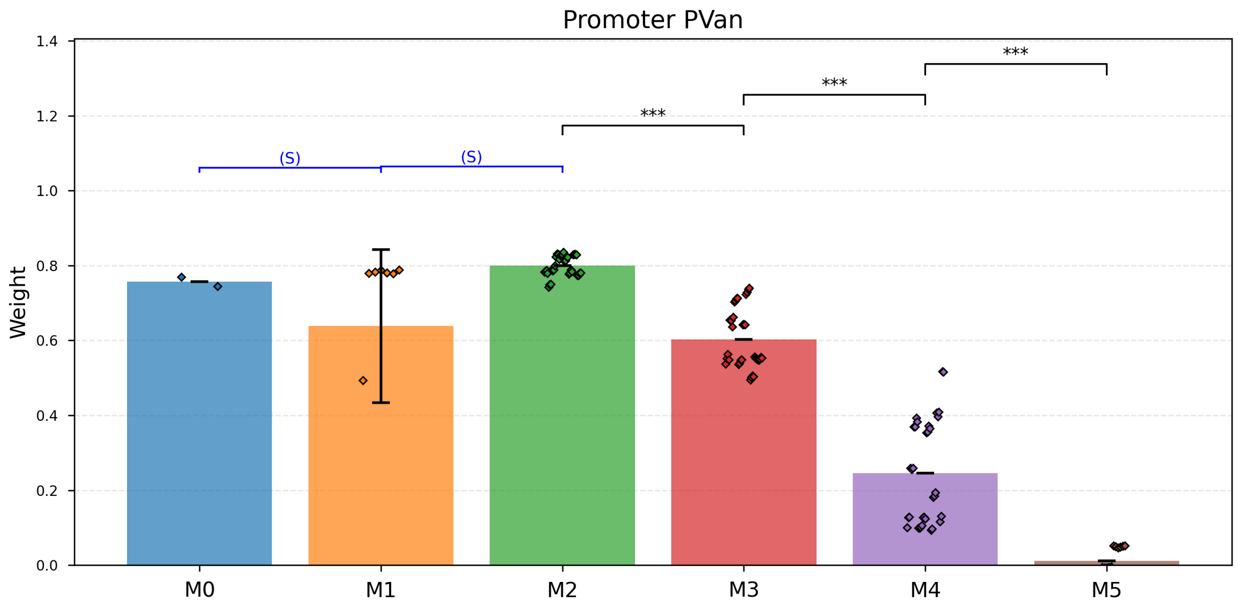

**Supplementary Fig. 68 | Promoter PVan.** Stability Tolerance (δ) = 0.1741 (2 *x* Pooled SD: 0.0871).

- M0 vs M1: Stable within ±δ (p_stable= 0.004).
- M1 vs M2: Stable within ±δ (p_stable= 0.015).
- M2 vs M3: Drift (p_change= $2.9 \times 10^{-17}$).
- M3 vs M4: Drift (p_change= $1.1 \times 10^{-17}$).
- M4 vs M5: Drift (p_change= $2.3 \times 10^{-11}$).

| M-State | Mean Weight | Std Dev |
|---|---|---|
| M0 | 0.757 | 0.012 |
| M1 | 0.7829 | 0.0017 |
| M2 | 0.800 | 0.005 |
| M3 | 0.603 | 0.013 |
| M4 | 0.246 | 0.024 |
| M5 | 0.011 | 0.007 |

**Supplementary Table 76 | Stability Tolerance (δ) = 0.1741 (2 *x* Pooled SD: 0.0871).**

#### Promoter PTac

Stability Tolerance (δ) = 0.1865 (2 *x* Pooled SD: 0.0932)

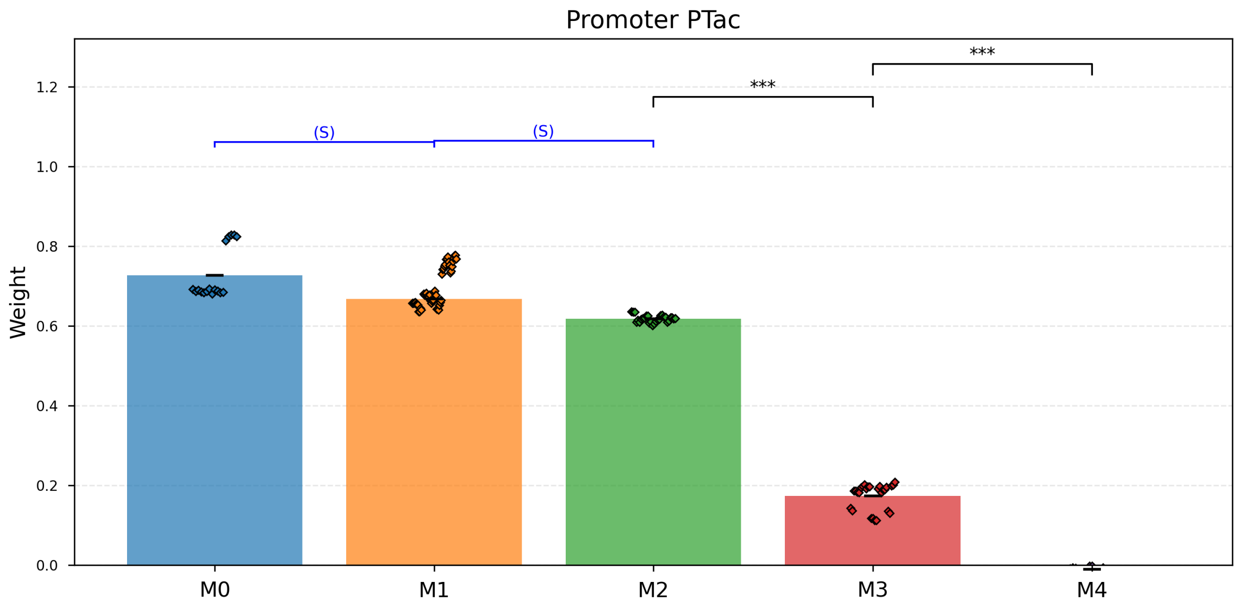

**Supplementary Fig. 69 | Promoter PTac.** Stability Tolerance (δ) = 0.1865 (2 *x* Pooled SD: 0.0932).

- M0 vs M1: Stable within ±δ (p_stable= $1.9 \times 10^{-6}$).
- M1 vs M2: Stable within ±δ (p_stable= $2.6 \times 10^{-9}$).
- M2 vs M3: Drift (p_change= $2.9 \times 10^{-34}$).
- M3 vs M4: Drift (p_change= $2.7 \times 10^{-22}$).

| M-State | Mean Weight | Std Dev |
|---|---|---|
| M0 | 0.727 | 0.016 |
| M1 | 0.668 | 0.020 |
| M2 | 0.6184 | 0.0017 |
| M3 | 0.173 | 0.006 |
| M4 | -0.0110 | 0.0013 |

**Supplementary Table 77 | Stability Tolerance (δ) = 0.1865 (2 *x* Pooled SD: 0.0932).**

#### Promoter PVan-Ttg

Stability Tolerance (δ) = 0.4326 (2 *x* Pooled SD: 0.2163)

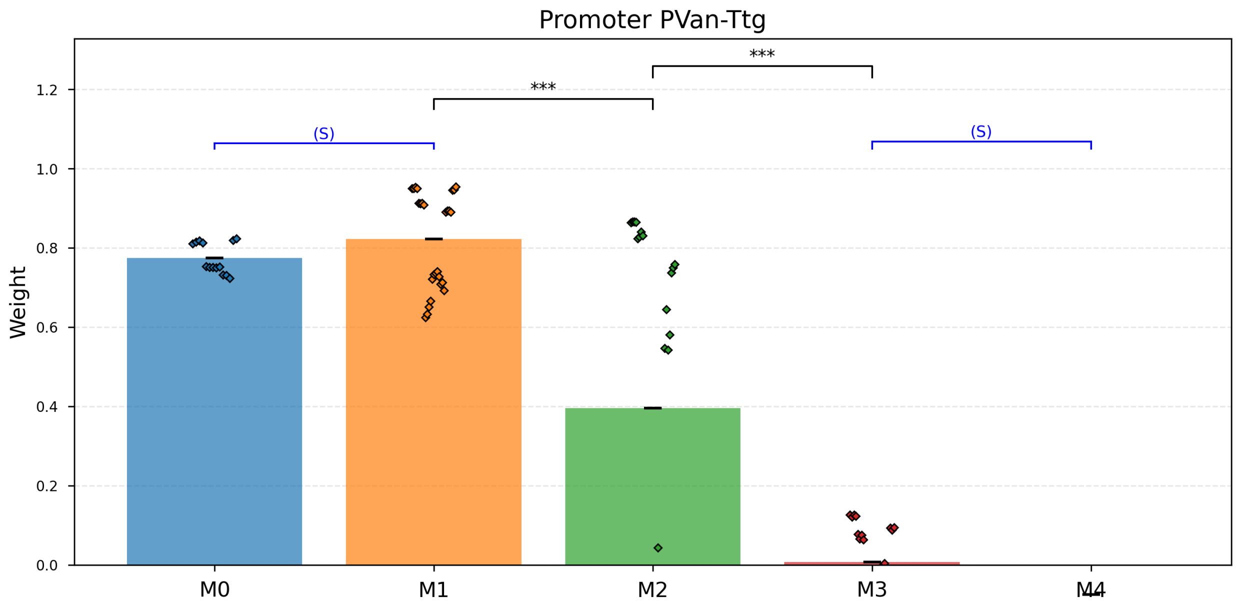

**Supplementary Fig. 70 | Promoter PVan-Ttg.** Stability Tolerance (δ) = 0.4326 (2 *x* Pooled SD: 0.2163).

- M0 vs M1: Stable within ±δ (p_stable= $3.8 \times 10^{-17}$).
- M1 vs M2: Drift (p_change= $1.9 \times 10^{-5}$).
- M2 vs M3: Drift (p_change= $6.6 \times 10^{-5}$).
- M3 vs M4: Stable within ±δ (p_stable= $2.6 \times 10^{-17}$).

| M-State | Mean Weight | Std Dev |
|---|---|---|
| M0 | 0.775 | 0.010 |
| M1 | 0.823 | 0.023 |
| M2 | 0.40 | 0.08 |
| M3 | 0.007 | 0.019 |
| M4 | -0.075 | 0.006 |

**Supplementary Table 78 | Stability Tolerance (δ) = 0.4326 (2 *x* Pooled SD: 0.2163).**

#### Promoter PTac-Tet

Stability Tolerance (δ) = 0.1122 (2 *x* Pooled SD: 0.0561)

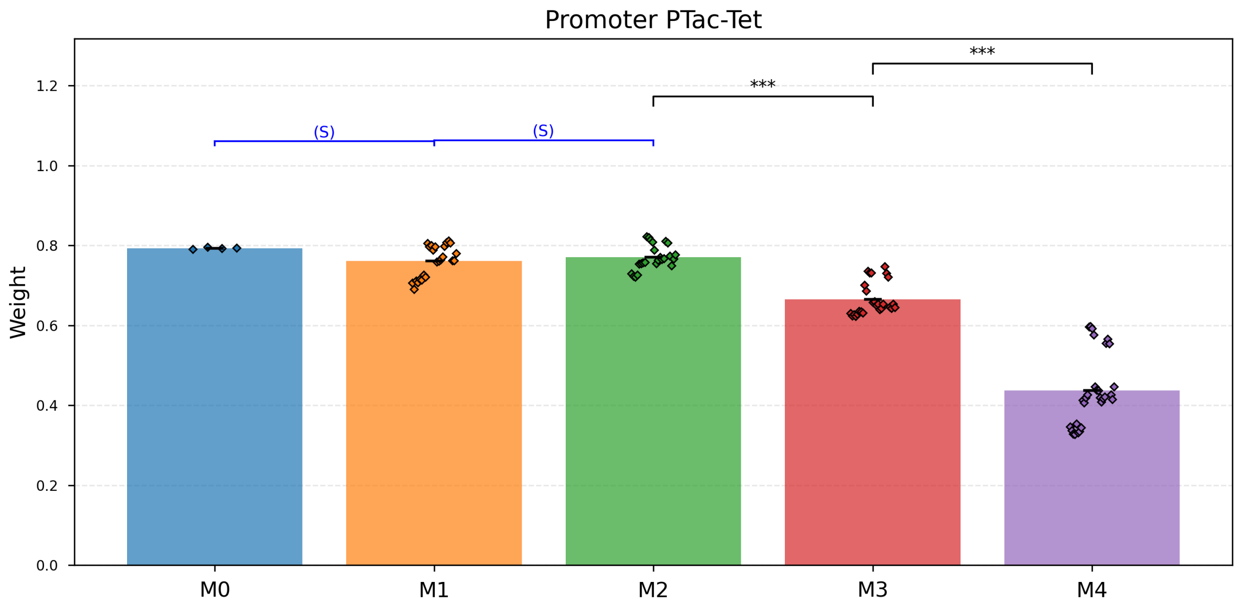

**Supplementary Fig. 71 | Promoter PTac-Tet.** Stability Tolerance (δ) = 0.1122 (2 *x* Pooled SD: 0.0561).

- M0 vs M1: Stable within ±δ (p_stable= $3.7 \times 10^{-10}$).
- M1 vs M2: Stable within ±δ (p_stable= $3.2 \times 10^{-13}$).
- M2 vs M3: Drift (p_change= $6.5 \times 10^{-14}$).
- M3 vs M4: Drift (p_change= $2.7 \times 10^{-15}$).

| M-State | Mean Weight | Std Dev |
|---|---|---|
| M0 | 0.7930 | 0.0011 |
| M1 | 0.761 | 0.008 |
| M2 | 0.770 | 0.006 |
| M3 | 0.665 | 0.008 |
| M4 | 0.437 | 0.017 |

**Supplementary Table 79 | Stability Tolerance (δ) = 0.1122 (2 *x* Pooled SD: 0.0561).**

#### Promoter PTet-Ttg

Stability Tolerance (δ) = 0.2389 (2 *x* Pooled SD: 0.1195)

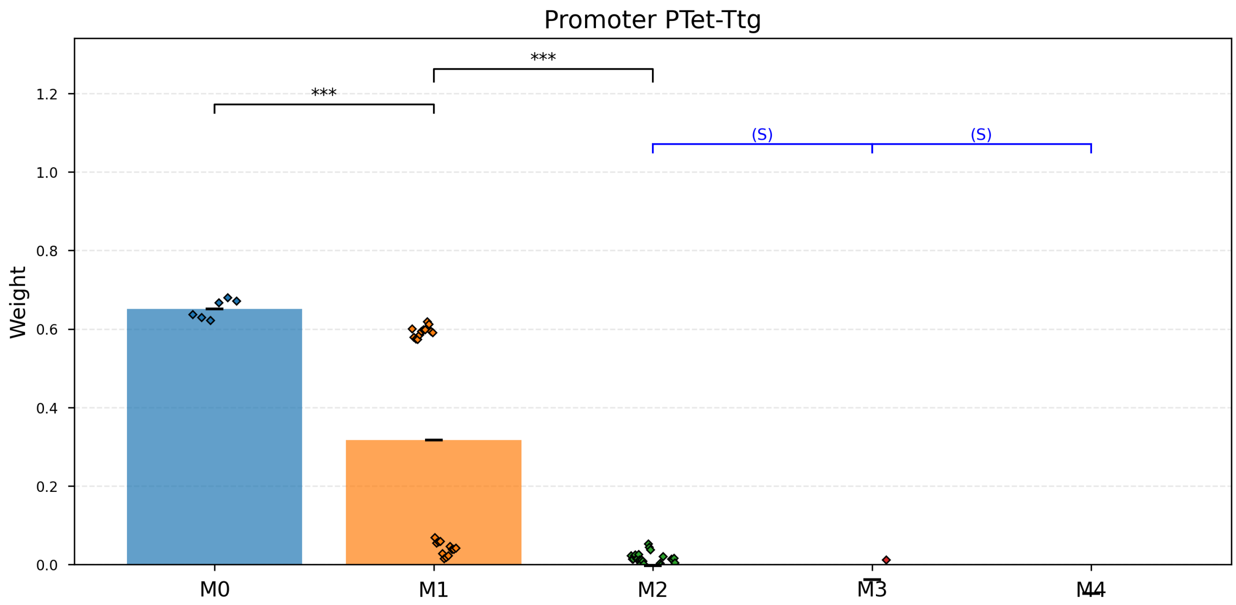

**Supplementary Fig. 72 | Promoter PTet-Ttg.** Stability Tolerance (δ) = 0.2389 (2 *x* Pooled SD: 0.1195).

- M0 vs M1: Drift (p_change= 6.7 × $10^{-6}$).
- M1 vs M2: Drift (p_change= 1.2 × $10^{-5}$).
- M2 vs M3: Stable within ±δ (p_stable= 4.2 × $10^{-52}$).
- M3 vs M4: Stable within ±δ (p_stable= 1.9 × $10^{-26}$).

| M-State | Mean Weight | Std Dev |
|---|---|---|
| M0 | 0.651 | 0.010 |
| M1 | 0.32 | 0.06 |
| M2 | -0.003 | 0.004 |
| M3 | -0.038 | 0.003 |
| M4 | -0.074 | 0.009 |

**Supplementary Table 80 | Stability Tolerance (δ) = 0.2389 (2 *x* Pooled SD: 0.1195).**

### Supplementary Note 11: Co-culture weight updates from plate reader fluorescence

#### *Summary Table*

| Cell Name | Growth rate µ (1/h) | Green fluorescence | Red fluorescence | Weight | Replicates |
|---|---|---|---|---|---|
| Pos_6-PBAD-M0+PSal-M0+PVan-M0-Ind_Sal | 0.267 ± 0.003 | 4,479 ± 65 | 1,206 ± 31 | 0.2399 ± 0.0025 | 5 |
| Pos_6-PBAD-M0+PSal-M0+PVan-M0-*K*_Van-Ind_Sal | 0.2903 ± 0.0030 | 6,760 ± 98 | 1,379 ± 24 | 0.1909 ± 0.0007 | 4 |
| Pos_6-PBAD-M0+PSal-M0+PVan-M0-*K*_Van-*K*_Ara-Ind_Sal | 0.278 ± 0.004 | 9,767 ± 75 | 1,218 ± 10 | 0.1242 ± 0.0016 | 5 |
| Pos_6-PBAD-M0+PSal-M0+PVan-M0-*K*_Van-*K*_Ara-*K*_Van-Ind_Sal | 0.2803 ± 0.0016 | 10,941 ± 58 | 1,215 ± 8 | 0.1118 ± 0.0005 | 4 |
| Pos_6-PBAD-M0+PSal-M0+PVan-M0-*K*_Van-*K*_Ara-*K*_Van-*K*_Ara-Ind_Sal | 0.252 ± 0.017 | 13,917 ± 119 | 1,253 ± 29 | 0.0922 ± 0.0017 | 5 |
| Pos_6-PBAD-M0+PSal-M0+PVan-M0-*K*_Van-*K*_Ara-*K*_Van-*K*_Ara-*K*_Van-Ind_Sal | 0.270 ± 0.004 | 17,864 ± 517 | 1,110 ± 27 | 0.0659 ± 0.0026 | 15 |
| Pos_6-PBAD-M0+PSal-M0+PVan-M0-*K*_Van-*K*_Ara-*K*_Van-*K*_Ara-*K*_Van-*K*_Ara-*K*_Sal-Ind_Sal | 0.251 ± 0.009 | 19,319 ± 824 | 730 ± 90 | 0.039 ± 0.004 | 22 |
| Pos_9-PSal-M0+PTet-M0-Ind_Sal | 0.272 ± 0.005 | 6,200 ± 74 | 2,231 ± 61 | 0.301 ± 0.004 | 4 |
| Pos_9-PSal-M0+PTet-M0-M-Ind_aTc | 0.224 ± 0.011 | 15,238 ± 211 | -18.0 ± 2.3 | -0.0023 ± 0.0004 | 7 |
| Pos_9-PSal-M0+PTet-M0-M-Ind_Sal | 0.2883 ± 0.0015 | 7,934 ± 107 | 2,996 ± 40 | 0.3122 ± 0.0013 | 4 |
| Pos_9-PSal-M0+PTet-M0-M-*K*_aTc-Ind_Sal | 0.285 ± 0.005 | 11,399 ± 288 | 612 ± 17 | 0.0567 ± 0.0007 | 5 |
| Pos_9-PSal-M0+PTet-M0-M-*K*_aTc-M-Ind_Sal | 0.289 ± 0.003 | 11,575 ± 497 | 728 ± 44 | 0.0656 ± 0.0010 | 8 |
| Pos_9-PSal-M0+PTet-M0-M-*K*_aTc-M-*K*_aTc-Ind_Sal | 0.2843 ± 0.0012 | 15,150 ± 672 | 732 ± 51 | 0.0509 ± 0.0017 | 11 |
| Pos_9-PSal-M0+PTet-M0-M-*K*_aTc-M-*K*_aTc-M-Ind_Sal | 0.287 ± 0.003 | 11,781 ± 155 | 568 ± 13 | 0.0511 ± 0.0005 | 5 |
| Pos_9-PSal-M0+PTet-M0-M-*K*_aTc-M-*K*_aTc- | 0.2914 ± 0.0010 | 16,552 ± 655 | 181 ± 8 | 0.01198 ± 0.00010 | 8 |

| | | | | | |
|---|---|---|---|---|---|
| M-*K*_aTc-M-Ind_Sal | | | | | |
| Pos_8-PBetI-M1+PCin-M1+PTet-M1-M-Ind_aTc | 0.211 ± 0.005 | 6,448 ± 313 | 5,764 ± 157 | 0.699 ± 0.008 | 7 |
| Pos_8-PBetI-M1+PCin-M1+PTet-M1-M-*K*_Cho-Ind_aTc | 0.233 ± 0.007 | 13,062 ± 109 | 109 ± 285 | 0.00 ± 0.04 | 8 |
| Pos_8-PBetI-M1+PCin-M1+PTet-M1-M-*K*_Cho-M-Ind_aTc | 0.214 ± 0.020 | 13,785 ± 899 | 838 ± 76 | 0.1076 ± 0.0024 | 8 |
| Pos_8-PBetI-M1+PCin-M1+PTet-M1-M-*K*_Cho-M-*K*_Cho-Ind_aTc | 0.243 ± 0.007 | 12,304 ± 207 | 306.1 ± 1.7 | 0.0471 ± 0.0005 | 5 |
| Pos_8-PBetI-M1+PCin-M1+PTet-M1-M-*K*_Cho-M-*K*_Cho-M-Ind_aTc | 0.249 ± 0.011 | 17,118 ± 348 | 488 ± 9 | 0.0537 ± 0.0011 | 5 |
| Pos_8-PBetI-M1+PCin-M1+PTet-M1-M-*K*_Cho-M-*K*_Cho-M-*K*_Cho-Ind_aTc | 0.213 ± 0.022 | 23,063 ± 97 | 147.2 ± 2.0 | 0.01247 ± 0.00014 | 3 |
| Pos_8-PBetI-M1+PCin-M1+PTet-M1-M-*K*_Cho-M-*K*_Cho-M-*K*_Cho-M-Ind_aTc | 0.253 ± 0.004 | 15,649 ± 672 | 180 ± 21 | 0.0215 ± 0.0016 | 12 |
| Pos_9-PSal-M0+PTet-M0-Ind_aTc | 0.226 ± 0.003 | 13,075 ± 324 | -28 ± 4 | -0.0043 ± 0.0007 | 5 |
| Pos_9-PSal-M0+PTet-M0-M-*K*_aTc-Ind_aTc | 0.2450 ± 0.0029 | 9,316 ± 300 | -41.2 ± 1.9 | -0.0088 ± 0.0003 | 7 |
| Pos_9-PSal-M0+PTet-M0-M-*K*_aTc-M-Ind_aTc | 0.244 ± 0.007 | 8,847 ± 227 | -47.7 ± 1.7 | -0.0108 ± 0.0006 | 7 |
| Pos_9-PSal-M0+PTet-M0-M-*K*_aTc-M-*K*_aTc-Ind_aTc | 0.237 ± 0.006 | 7,079 ± 360 | -22 ± 5 | -0.0065 ± 0.0014 | 11 |
| Pos_9-PSal-M0+PTet-M0-M-*K*_aTc-M-*K*_aTc-M-Ind_aTc | 0.073 ± 0.005 | 9,160 ± 270 | -55.3 ± 2.6 | -0.01202 ± 0.00024 | 4 |
| Pos_9-PSal-M0+PTet-M0-M-*K*_aTc-M-*K*_aTc-M-*K*_aTc-M-Ind_aTc | 0.241 ± 0.006 | 4,596 ± 259 | -43.9 ± 1.1 | -0.0195 ± 0.0010 | 8 |
| Pos_7-PBetI-M0+PTac-M0-Ind_Cho | 0.280 ± 0.006 | 2,922 ± 91 | 2,888 ± 84 | 1.179 ± 0.004 | 5 |
| Pos_7-PBetI-M0+PTac-M0-*K*_Cho-Ind_Cho | 0.2912 ± 0.0027 | 3,202 ± 42 | 681 ± 9 | 0.648 ± 0.006 | 4 |
| Pos_7-PBetI-M0+PTac-M0-*K*_Cho-M-Ind_Cho | 0.2840 ± 0.0020 | 2,784 ± 143 | 221 ± 14 | 0.330 ± 0.008 | 5 |
| Pos_7-PBetI-M0+PTac-M0- | 0.290 ± 0.005 | 2,880 ± 100 | 167 ± 7 | 0.256 ± 0.007 | 5 |

| | | | | | |
|---|---|---|---|---|---|
| *K*_Cho-M-*K*_Cho-Ind_Cho | | | | | |
| Pos_7-PBetI-M0+PTac-M0-*K*_Cho-M-*K*_Cho-M-Ind_Cho | 0.2794 ± 0.0014 | 2,948 ± 108 | 137 ± 11 | 0.212 ± 0.012 | 5 |
| Pos_7-PBetI-M0+PTac-M0-*K*_Cho-M-*K*_Cho-M-*K*_Cho-Ind_Cho | 0.288 ± 0.010 | 3,030 ± 153 | 69 ± 28 | 0.09 ± 0.04 | 10 |
| Pos_7-PBetI-M0+PTac-M0-*K*_Cho-M-*K*_Cho-M-*K*_Cho-M-Ind_Cho | 0.322 ± 0.011 | 3,189 ± 182 | 70 ± 6 | 0.107 ± 0.005 | 5 |
| Pos_7-PBetI-M0+PTac-M0-*K*_Cho-M-*K*_Cho-M-*K*_Cho-M-M-Ind_Cho | 0.294 ± 0.004 | 3,005 ± 104 | 131 ± 35 | 0.17 ± 0.04 | 15 |
| Pos_8-PBetI-M1+PCin-M1+PTet-M1-Ind_Cho | 0.278 ± 0.005 | 1,971 ± 62 | 1,472 ± 87 | 1.097 ± 0.009 | 5 |
| Pos_8-PBetI-M1+PCin-M1+PTet-M1-M-Ind_Cho | 0.286 ± 0.003 | 2,307 ± 35 | 682.8 ± 2.7 | 0.773 ± 0.007 | 4 |
| Pos_8-PBetI-M1+PCin-M1+PTet-M1-M-*K*_Cho-Ind_Cho | 0.283 ± 0.004 | 1,927 ± 84 | 1 ± 3 | -0.000 ± 0.009 | 5 |
| Pos_8-PBetI-M1+PCin-M1+PTet-M1-M-*K*_Cho-M-Ind_Cho | 0.2806 ± 0.0027 | 1,878 ± 76 | -2 ± 4 | -0.007 ± 0.010 | 5 |
| Pos_8-PBetI-M1+PCin-M1+PTet-M1-M-*K*_Cho-M-*K*_Cho-Ind_Cho | 0.2829 ± 0.0026 | 1,975 ± 82 | -48 ± 9 | -0.15 ± 0.04 | 4 |
| Pos_8-PBetI-M1+PCin-M1+PTet-M1-M-*K*_Cho-M-*K*_Cho-M-Ind_Cho | 0.2851 ± 0.0020 | 2,056 ± 133 | -19 ± 5 | -0.058 ± 0.016 | 8 |
| Pos_8-PBetI-M1+PCin-M1+PTet-M1-M-*K*_Cho-M-*K*_Cho-M-*K*_Cho-M-Ind_Cho | 0.2913 ± 0.0010 | 1,857 ± 75 | -46.06 ± 0.20 | -0.145 ± 0.007 | 5 |
| Pos_6-PBAD-M0+PSal-M0+PVan-M0-Ind_Ara | 0.179 ± 0.018 | 5,644 ± 156 | 17,296 ± 611 | 0.759 ± 0.003 | 5 |
| Pos_6-PBAD-M0+PSal-M0+PVan-M0-*K*_Van-Ind_Ara | 0.2403 ± 0.0030 | 6,916 ± 142 | 13,874 ± 335 | 0.6465 ± 0.0013 | 4 |
| Pos_6-PBAD-M0+PSal-M0+PVan-M0-*K*_Van-*K*_Ara-Ind_Ara | 0.195 ± 0.018 | 5,075 ± 65 | 5,839 ± 156 | 0.490 ± 0.005 | 5 |
| Pos_6-PBAD-M0+PSal-M0+PVan-M0-*K*_Van-*K*_Ara-*K*_Van-Ind_Ara | 0.1857 ± 0.0025 | 5,101 ± 195 | 5,376 ± 202 | 0.466 ± 0.004 | 5 |

| | | | | | |
|---|---|---|---|---|---|
| Pos_6-PBAD-M0+PSal-M0+PVan-M0-*K*_Van-*K*_Ara-*K*_Van-*K*_Ara-Ind_Ara | 0.230 ± 0.017 | 4,406 ± 177 | 3,330 ± 180 | 0.377 ± 0.005 | 5 |
| Pos_6-PBAD-M0+PSal-M0+PVan-M0-*K*_Van-*K*_Ara-*K*_Van-*K*_Ara-*K*_Van-*K*_Ara-Ind_Ara | 0.181 ± 0.009 | 4,537 ± 266 | 1,966 ± 319 | 0.224 ± 0.015 | 26 |
| Pos_6-PBAD-M0+PSal-M0+PVan-M0-*K*_Van-*K*_Ara-*K*_Van-*K*_Ara-*K*_Van-*K*_Ara-*K*_Sal-Ind_Ara | 0.1590 ± 0.0027 | 4,350 ± 150 | 9,185 ± 516 | 0.659 ± 0.007 | 6 |
| Pos_1-PLux-M0-Ind_OC6 | 0.248 ± 0.010 | 7,883 ± 619 | 17,178 ± 2,571 | 0.730 ± 0.027 | 9 |
| Pos_1-PLux-M0-M-*K*_OC6-Ind_OC6 | 0.263 ± 0.003 | 6,452 ± 219 | 7,265 ± 234 | 0.5488 ± 0.0026 | 5 |
| Pos_1-PLux-M0-M-*K*_OC6-M-Ind_OC6 | 0.2684 ± 0.0024 | 6,256 ± 135 | 7,461 ± 164 | 0.5666 ± 0.0008 | 5 |
| Pos_1-PLux-M0-M-*K*_OC6-M-*K*_OC6-Ind_OC6 | 0.2604 ± 0.0021 | 5,981 ± 236 | 6,100 ± 244 | 0.5182 ± 0.0007 | 5 |
| Pos_1-PLux-M0-M-*K*_OC6-M-*K*_OC6-M-Ind_OC6 | 0.232 ± 0.013 | 6,454 ± 366 | 7,114 ± 426 | 0.5416 ± 0.0013 | 9 |
| Pos_1-PLux-M0-M-*K*_OC6-M-*K*_OC6-M-*K*_OC6-M-Ind_OC6 | 0.259 ± 0.006 | 6,645 ± 138 | 5,693 ± 109 | 0.467 ± 0.008 | 20 |
| Pos_4-PCin-M0+PLux-M0-Ind_OC6 | 0.248 ± 0.003 | 4,467 ± 106 | 7,217 ± 186 | 0.6624 ± 0.0009 | 5 |
| Pos_4-PCin-M0+PLux-M0-*K*_OC6-Ind_OC6 | 0.2720 ± 0.0014 | 4,239 ± 27 | 3,539 ± 51 | 0.4582 ± 0.0025 | 4 |
| Pos_4-PCin-M0+PLux-M0-*K*_OC6-M-Ind_OC6 | 0.269 ± 0.005 | 3,437 ± 51 | 2,958 ± 43 | 0.467 ± 0.003 | 5 |
| Pos_4-PCin-M0+PLux-M0-*K*_OC6-M-*K*_OC6-Ind_OC6 | 0.2725 ± 0.0026 | 2,840 ± 55 | 1,694 ± 22 | 0.365 ± 0.004 | 5 |
| Pos_4-PCin-M0+PLux-M0-*K*_OC6-M-*K*_OC6-M-Ind_OC6 | 0.240 ± 0.015 | 2,807 ± 84 | 1,769 ± 25 | 0.380 ± 0.006 | 5 |
| Pos_4-PCin-M0+PLux-M0-*K*_OC6-M-*K*_OC6-M-*K*_OC6-Ind_OC6 | 0.2775 ± 0.0015 | 2,463 ± 96 | 1,256 ± 40 | 0.326 ± 0.003 | 5 |
| Pos_4-PCin-M0+PLux-M0-*K*_OC6-M-*K*_OC6-M-*K*_OC6-M-*K*_OC6-Ind_OC6 | 0.2712 ± 0.0021 | 2,367 ± 48 | 886 ± 38 | 0.256 ± 0.008 | 10 |

| | | | | | |
|---|---|---|---|---|---|
| Pos_4-PCin-M0+PLux-M0-Ind_OHC14 | 0.240 ± 0.014 | 3,023 ± 98 | 943 ± 40 | 0.3015 ± 0.0025 | 5 |
| Pos_4-PCin-M0+PLux-M0-*K*_OC6-Ind_OHC14 | 0.257 ± 0.018 | 3,173 ± 101 | 1,055 ± 37 | 0.3169 ± 0.0016 | 4 |
| Pos_4-PCin-M0+PLux-M0-*K*_OC6-M-Ind_OHC14 | 0.277 ± 0.004 | 3,147 ± 72 | 1,149 ± 7 | 0.340 ± 0.004 | 5 |
| Pos_4-PCin-M0+PLux-M0-*K*_OC6-M-*K*_OC6-Ind_OHC14 | 0.2626 ± 0.0008 | 2,993 ± 34 | 1,132 ± 9 | 0.3487 ± 0.0012 | 4 |
| Pos_4-PCin-M0+PLux-M0-*K*_OC6-M-*K*_OC6-M-Ind_OHC14 | 0.35 ± 0.10 | 3,229 ± 205 | 1,288 ± 82 | 0.3624 ± 0.0027 | 5 |
| Pos_4-PCin-M0+PLux-M0-*K*_OC6-M-*K*_OC6-M-*K*_OC6-Ind_OHC14 | 0.267 ± 0.005 | 3,281 ± 166 | 1,192 ± 29 | 0.339 ± 0.007 | 4 |
| Pos_4-PCin-M0+PLux-M0-*K*_OC6-M-*K*_OC6-M-*K*_OC6-M-*K*_OC6-Ind_OHC14 | 0.258 ± 0.005 | 3,507 ± 81 | 1,266 ± 43 | 0.336 ± 0.003 | 10 |
| Pos_8-PBetI-M1+PCin-M1+PTet-M1-Ind_OHC14 | 0.275 ± 0.011 | 2,385 ± 72 | 688.5 ± 1.1 | 0.284 ± 0.007 | 2 |
| Pos_8-PBetI-M1+PCin-M1+PTet-M1-M-Ind_OHC14 | 0.25 ± 0.04 | 2,834 ± 71 | 717 ± 15 | 0.2560 ± 0.0025 | 4 |
| Pos_8-PBetI-M1+PCin-M1+PTet-M1-M-*K*_Cho-Ind_OHC14 | 0.232 ± 0.016 | 3,550 ± 70 | 124 ± 3 | 0.0425 ± 0.0004 | 5 |
| Pos_8-PBetI-M1+PCin-M1+PTet-M1-M-*K*_Cho-M-Ind_OHC14 | 0.269 ± 0.013 | 3,773 ± 184 | 134 ± 8 | 0.0431 ± 0.0006 | 7 |
| Pos_8-PBetI-M1+PCin-M1+PTet-M1-M-*K*_Cho-M-*K*_Cho-Ind_OHC14 | 0.252 ± 0.011 | 4,118 ± 99 | 54.9 ± 2.0 | 0.0166 ± 0.0005 | 6 |
| Pos_8-PBetI-M1+PCin-M1+PTet-M1-M-*K*_Cho-M-*K*_Cho-M-Ind_OHC14 | 0.265 ± 0.008 | 3,370 ± 185 | 36 ± 6 | 0.0132 ± 0.0013 | 5 |
| Pos_8-PBetI-M1+PCin-M1+PTet-M1-M-*K*_Cho-M-*K*_Cho-M-*K*_Cho-M-Ind_OHC14 | 0.268 ± 0.006 | 3,734 ± 398 | -5 ± 5 | -0.0025 ± 0.0017 | 6 |
| Pos_2-PTac-M1+PTtg-M1+PVan-M1-Ind_Nar | 0.272 ± 0.004 | 2,865 ± 42 | 304 ± 8 | 0.491 ± 0.005 | 5 |

| | | | | | |
|---|---|---|---|---|---|
| Pos_2-PTac-M1+PTtg-M1+PVan-M1-*K*_IPTG-Ind_Nar | 0.2708 ± 0.0027 | 3,557 ± 50 | 195 ± 5 | 0.317 ± 0.003 | 4 |
| Pos_2-PTac-M1+PTtg-M1+PVan-M1-*K*_IPTG-M-Ind_Nar | 0.213 ± 0.015 | 2,919 ± 63 | 142.7 ± 2.1 | 0.2910 ± 0.0020 | 5 |
| Pos_2-PTac-M1+PTtg-M1+PVan-M1-*K*_IPTG-M-*K*_IPTG-Ind_Nar | 0.276 ± 0.006 | 2,769 ± 70 | 94 ± 5 | 0.219 ± 0.008 | 5 |
| Pos_2-PTac-M1+PTtg-M1+PVan-M1-*K*_IPTG-M-*K*_IPTG-M-Ind_Nar | 0.251 ± 0.012 | 2,936 ± 31 | 106 ± 4 | 0.229 ± 0.007 | 5 |
| Pos_2-PTac-M1+PTtg-M1+PVan-M1-*K*_IPTG-M-*K*_IPTG-M-*K*_IPTG-Ind_Nar | 0.249 ± 0.012 | 2,766 ± 23 | 5.5 ± 2.9 | 0.015 ± 0.008 | 5 |
| Pos_2-PTac-M1+PTtg-M1+PVan-M1-*K*_IPTG-M-*K*_IPTG-M-*K*_IPTG-M-*K*_Van-Ind_Nar | 0.273 ± 0.007 | 4,344 ± 323 | 60 ± 16 | 0.087 ± 0.018 | 15 |
| Pos_3-PTtg-M0+PVan-M0-Ind_Nar | 0.237 ± 0.016 | 3,956 ± 155 | 448 ± 24 | 0.509 ± 0.005 | 5 |
| Pos_3-PTtg-M0+PVan-M0-M-Ind_Nar | 0.267 ± 0.003 | 3,516 ± 8 | 395 ± 6 | 0.507 ± 0.005 | 4 |
| Pos_3-PTtg-M0+PVan-M0-M-M-Ind_Nar | 0.268 ± 0.006 | 2,374 ± 28 | 197 ± 5 | 0.421 ± 0.005 | 5 |
| Pos_3-PTtg-M0+PVan-M0-M-M-M-Ind_Nar | 0.272 ± 0.006 | 2,156 ± 55 | 122 ± 7 | 0.324 ± 0.009 | 5 |
| Pos_3-PTtg-M0+PVan-M0-M-M-M-M-Ind_Nar | 0.2696 ± 0.0024 | 2,277 ± 33 | 197 ± 6 | 0.433 ± 0.007 | 5 |
| Pos_3-PTtg-M0+PVan-M0-M-M-M-M-M-Ind_Nar | 0.2642 ± 0.0018 | 2,005 ± 36 | 203 ± 6 | 0.478 ± 0.004 | 5 |
| Pos_3-PTtg-M0+PVan-M0-M-M-M-M-M-M-*K*_Nar-Ind_Nar | 0.274 ± 0.004 | 2,039 ± 33 | 192 ± 5 | 0.456 ± 0.004 | 10 |
| Pos_2-PTac-M1+PTtg-M1+PVan-M1-Ind_Van | 0.277 ± 0.004 | 5,900 ± 97 | 1,168 ± 45 | 0.166 ± 0.004 | 5 |
| Pos_2-PTac-M1+PTtg-M1+PVan-M1-*K*_IPTG-Ind_Van | 0.2867 ± 0.0021 | 8,114 ± 204 | 1,571 ± 32 | 0.1633 ± 0.0008 | 4 |
| Pos_2-PTac-M1+PTtg-M1+PVan-M1-*K*_IPTG-M-Ind_Van | 0.284 ± 0.005 | 8,942 ± 179 | 1,120 ± 57 | 0.111 ± 0.003 | 5 |
| Pos_2-PTac-M1+PTtg-M1+PVan-M1- | 0.2933 ± 0.0030 | 7,415 ± 238 | 757 ± 47 | 0.0916 ± 0.0025 | 5 |

| | | | | | |
|---|---|---|---|---|---|
| *K*_IPTG-M-*K*_IPTG-Ind_Van | | | | | |
| Pos_2-PTac-M1+PTtg-M1+PVan-M1-*K*_IPTG-M-*K*_IPTG-M-Ind_Van | 0.267 ± 0.017 | 6,998 ± 228 | 764 ± 35 | 0.0977 ± 0.0016 | 5 |
| Pos_2-PTac-M1+PTtg-M1+PVan-M1-*K*_IPTG-M-*K*_IPTG-M-*K*_IPTG-Ind_Van | 0.258 ± 0.015 | 9,116 ± 217 | 439 ± 29 | 0.0450 ± 0.0019 | 5 |
| Pos_2-PTac-M1+PTtg-M1+PVan-M1-*K*_IPTG-M-*K*_IPTG-M-*K*_IPTG-M-*K*_Van-Ind_Van | 0.269 ± 0.011 | 7,161 ± 432 | 487 ± 68 | 0.061 ± 0.007 | 15 |
| Pos_3-PTtg-M0+PVan-M0-Ind_Van | 0.276 ± 0.007 | 3,333 ± 120 | 5,707 ± 331 | 0.697 ± 0.007 | 5 |
| Pos_3-PTtg-M0+PVan-M0-M-Ind_Van | 0.2895 ± 0.0008 | 3,562 ± 55 | 7,002 ± 104 | 0.7383 ± 0.0014 | 4 |
| Pos_3-PTtg-M0+PVan-M0-M-M-Ind_Van | 0.2751 ± 0.0017 | 2,975 ± 77 | 4,204 ± 228 | 0.640 ± 0.009 | 5 |
| Pos_3-PTtg-M0+PVan-M0-M-M-M-Ind_Van | 0.286 ± 0.016 | 2,707 ± 81 | 3,169 ± 187 | 0.584 ± 0.009 | 5 |
| Pos_3-PTtg-M0+PVan-M0-M-M-M-M-Ind_Van | 0.264 ± 0.014 | 2,830 ± 81 | 4,234 ± 227 | 0.657 ± 0.008 | 5 |
| Pos_3-PTtg-M0+PVan-M0-M-M-M-M-M-Ind_Van | 0.272 ± 0.018 | 2,680 ± 100 | 3,384 ± 244 | 0.606 ± 0.012 | 5 |
| Pos_3-PTtg-M0+PVan-M0-M-M-M-M-M-M-*K*_Nar-Ind_Van | 0.2808 ± 0.0027 | 2,640 ± 60 | 4,697 ± 179 | 0.708 ± 0.005 | 10 |
| Pos_6-PBAD-M0+PSal-M0+PVan-M0-Ind_Van | 0.280 ± 0.006 | 2,957 ± 95 | 3,163 ± 153 | 0.558 ± 0.006 | 5 |
| Pos_6-PBAD-M0+PSal-M0+PVan-M0-*K*_Van-Ind_Van | 0.286 ± 0.005 | 3,044 ± 4 | 1,309 ± 22 | 0.311 ± 0.004 | 4 |
| Pos_6-PBAD-M0+PSal-M0+PVan-M0-*K*_Van-*K*_Ara-Ind_Van | 0.3005 ± 0.0028 | 2,540 ± 90 | 699 ± 29 | 0.2193 ± 0.0024 | 5 |
| Pos_6-PBAD-M0+PSal-M0+PVan-M0-*K*_Van-*K*_Ara-*K*_Van-Ind_Van | 0.2947 ± 0.0027 | 2,451 ± 47 | 485 ± 18 | 0.1661 ± 0.0028 | 5 |
| Pos_6-PBAD-M0+PSal-M0+PVan-M0-*K*_Van-*K*_Ara-*K*_Van-*K*_Ara-Ind_Van | 0.269 ± 0.015 | 2,507 ± 31 | 623 ± 10 | 0.2018 ± 0.0021 | 5 |
| Pos_6-PBAD-M0+PSal-M0+PVan-M0-*K*_Van-*K*_Ara- | 0.258 ± 0.009 | 2,523 ± 66 | 194 ± 9 | 0.071 ± 0.003 | 16 |

| | | | | | |
|---|---|---|---|---|---|
| *K*_Van-*K*_Ara-*K*_Van-*K*_Ara-Ind_Van | | | | | |
| Pos_6-PBAD-M0+PSal-M0+PVan-M0-*K*_Van-*K*_Ara-*K*_Van-*K*_Ara-*K*_Van-*K*_Ara-*K*_Sal-Ind_Van | 0.251 ± 0.010 | 2,304 ± 48 | 92 ± 13 | 0.037 ± 0.005 | 22 |
| Pos_2-PTac-M1+PTtg-M1+PVan-M1-Ind_IPTG | 0.274 ± 0.005 | 2,589 ± 34 | 350 ± 8 | 0.1659 ± 0.0014 | 5 |
| Pos_2-PTac-M1+PTtg-M1+PVan-M1-*K*_IPTG-Ind_IPTG | 0.2800 ± 0.0022 | 2,930 ± 21 | 203 ± 4 | 0.0921 ± 0.0013 | 4 |
| Pos_2-PTac-M1+PTtg-M1+PVan-M1-*K*_IPTG-M-Ind_IPTG | 0.2726 ± 0.0017 | 2,417 ± 43 | 172 ± 6 | 0.0944 ± 0.0020 | 5 |
| Pos_2-PTac-M1+PTtg-M1+PVan-M1-*K*_IPTG-M-*K*_IPTG-Ind_IPTG | 0.294 ± 0.004 | 2,190 ± 45 | 51.4 ± 2.6 | 0.0330 ± 0.0011 | 5 |
| Pos_2-PTac-M1+PTtg-M1+PVan-M1-*K*_IPTG-M-*K*_IPTG-M-Ind_IPTG | 0.2775 ± 0.0018 | 2,427 ± 61 | 73 ± 3 | 0.0420 ± 0.0010 | 5 |
| Pos_2-PTac-M1+PTtg-M1+PVan-M1-*K*_IPTG-M-*K*_IPTG-M-*K*_IPTG-Ind_IPTG | 0.277 ± 0.004 | 2,179 ± 51 | -15.6 ± 1.7 | -0.0106 ± 0.0013 | 5 |
| Pos_2-PTac-M1+PTtg-M1+PVan-M1-*K*_IPTG-M-*K*_IPTG-M-*K*_IPTG-M-*K*_Van-Ind_IPTG | 0.2936 ± 0.0006 | 2,357 ± 43 | -15 ± 5 | -0.010 ± 0.003 | 10 |
| Pos_7-PBetI-M0+PTac-M0-Ind_IPTG | 0.285 ± 0.006 | 2,689 ± 28 | 573 ± 7 | 0.2401 ± 0.0007 | 5 |
| Pos_7-PBetI-M0+PTac-M0-*K*_Cho-Ind_IPTG | 0.2912 ± 0.0008 | 3,566 ± 38 | 466 ± 11 | 0.1613 ± 0.0017 | 4 |
| Pos_7-PBetI-M0+PTac-M0-*K*_Cho-M-Ind_IPTG | 0.278 ± 0.021 | 3,484 ± 144 | 517 ± 25 | 0.1794 ± 0.0014 | 5 |
| Pos_7-PBetI-M0+PTac-M0-*K*_Cho-M-*K*_Cho-Ind_IPTG | 0.2957 ± 0.0025 | 3,341 ± 44 | 380 ± 6 | 0.1432 ± 0.0009 | 5 |
| Pos_7-PBetI-M0+PTac-M0-*K*_Cho-M-*K*_Cho-M-Ind_IPTG | 0.266 ± 0.014 | 3,427 ± 68 | 433 ± 12 | 0.1567 ± 0.0016 | 5 |
| Pos_7-PBetI-M0+PTac-M0-*K*_Cho-M-*K*_Cho- | 0.306 ± 0.009 | 3,393 ± 162 | 254 ± 18 | 0.0982 ± 0.0026 | 10 |

| | | | | | |
|---|---|---|---|---|---|
| M-*K*_Cho-Ind_IPTG | | | | | |
| Pos_7-PBetI-M0+PTac-M0-*K*_Cho-M-*K*_Cho-M-*K*_Cho-M-Ind_IPTG | 0.343 ± 0.009 | 3,414 ± 168 | 177 ± 12 | 0.0701 ± 0.0021 | 5 |
| Pos_7-PBetI-M0+PTac-M0-*K*_Cho-M-*K*_Cho-M-*K*_Cho-M-M-Ind_IPTG | 0.259 ± 0.011 | 3,725 ± 108 | 155 ± 38 | 0.060 ± 0.014 | 15 |

**Supplementary Table 81 | Summary Table.**

#### Analysis of growth and fluorescence curves from plate reader

This section presents a comprehensive analysis of cellular growth and fluorescence dynamics, aggregated across all experimental datasets. The data is organized by promoter system and subsequently sorted by the experimental condition identifier (M-suffix) to provide a clear, logical progression. Within this framework, we distinguish between two levels of replication fundamental to biological experiments:

- Biological Replicates: These are distinct cultures, often originating from different colonies or clones (e.g., 'Clone A' and 'Clone B'), that are subjected to the same experimental conditions. They are treated as independent measurements of the biological process under investigation.
- Technical Replicates: These are multiple measurements taken from the same biological sample (e.g., multiple wells in a plate inoculated from the same starter culture). They serve to quantify the precision and variability of the measurement technique itself.

Throughout this analysis, each curve representing a biological replicate is computed as the mean of its corresponding technical replicates. The variability among these technical replicates is visualized as a shaded region representing the standard error of the mean (SEM). All mean curves are linearly interpolated between measured time points to ensure visual continuity.

#### Co-culture tournament analysis

Analysis of co-cultures grouped by board position. Brackets in time-course plots indicate stability (S) or significant drift (*).

##### Day 1

Initial Co-culture Composition: Pos 1: PLux-M0; Pos 2: PTtg-M1, PVan-M1, PTac-M1; Pos 3: PTtg-M0, PVan-M0; Pos 4: PLux-M0, PCin-M0; Pos 6: PSal-M0, PBAD-M0, PVan-M0; Pos 7: PBetI-M0, PTac-M0; Pos 8: PTet-M?, PBetI-M1, PCin-M1; Pos 9: PSal-M0, PTet-M0.

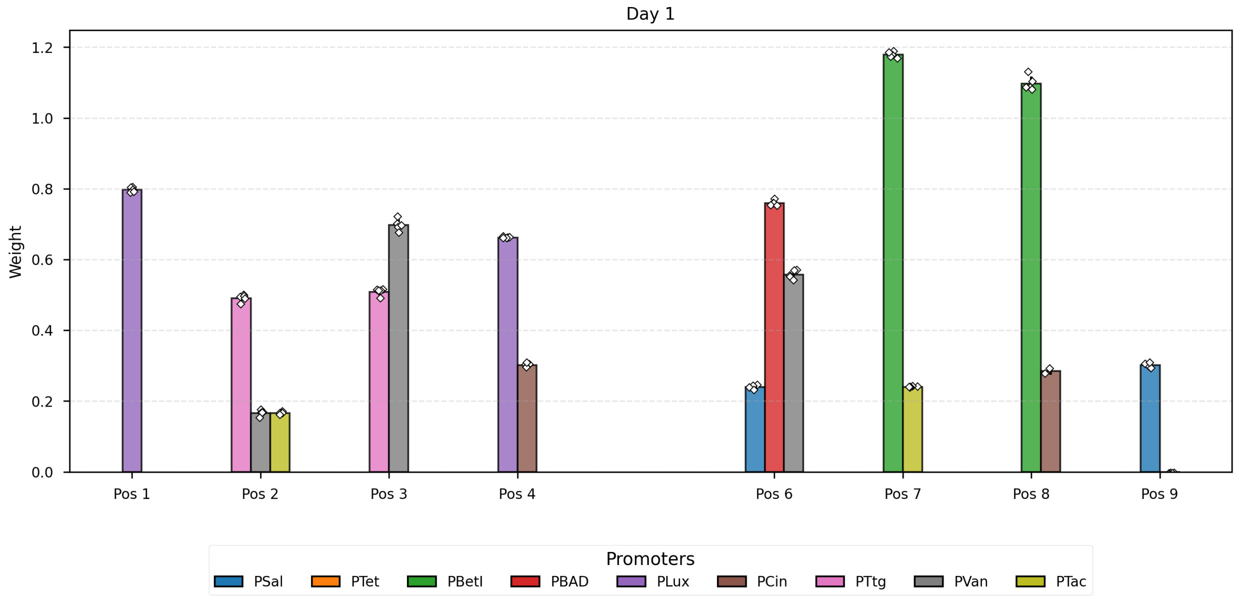

**Supplementary Fig. 73 | Day 1.**

##### Day 2

Visual Guide: The bars corresponding to Pos 2 (PTac), Pos 4 (PLux), Pos 6 (PVan), Pos 7 (PBetI) are drawn with thicker contours to indicate they were targets of learning in the previous step.

Analysis of Day 2 vs Day 1:

- Pos 1 PLux: Weight shifted (Δ=-0.152, $p_{change}$=2.0 × $10^{-8}$), showing a lack of weight stability in mono-culture.
- Pos 2 PTtg: Weight shifted (Δ=-0.174, $p_{change}$=1.5 × $10^{-8}$), showing a lack of weight stability in co-culture in presence of kanamycin.
- Pos 2 PVan: Weight stabilized (Δ=-0.003, tol=0.0284, $p_{stable}$=7.8 × $10^{-4}$), showing **weight stability in co-culture in presence of kanamycin.**
- Pos 2 PTac: Weight shifted (Δ=-0.074, $p_{change}$=2.0 × $10^{-9}$), showing a negative weight update due to kanamycin learning.
- Pos 3 PTtg: Weight stabilized (Δ=-0.001, tol=0.054, $p_{stable}$=5.0 × $10^{-5}$), showing weight stability in co-culture.
- Pos 3 PVan: Weight shifted (Δ=0.041, $p_{change}$=0.004), showing a lack of weight stability in co-culture.
- Pos 4 PLux: Weight shifted (Δ=-0.204, $p_{change}$=3.7 × $10^{-7}$), showing a negative weight update due to kanamycin learning.
- Pos 4 PCin: Weight shifted (Δ=0.015, $p_{change}$=0.002), showing a lack of weight stability in co-culture in presence of kanamycin.

- Pos 6 PSal: Weight shifted (Δ=-0.049, $p_{change}$=1.4 × $10^{-5}$), showing a lack of weight stability in co-culture in presence of kanamycin.
- Pos 6 PBAD: Weight shifted (Δ=-0.113, $p_{change}$=4.4 × $10^{-7}$), showing a lack of weight stability in co-culture in presence of kanamycin.
- Pos 6 PVan: Weight shifted (Δ=-0.247, $p_{change}$=7.3 × $10^{-9}$), showing a negative weight update due to kanamycin learning.
- Pos 7 PBetI: Weight shifted (Δ=-0.531, $p_{change}$=2.2 × $10^{-9}$), showing a negative weight update due to kanamycin learning.
- Pos 7 PTac: Weight shifted (Δ=-0.079, $p_{change}$=1.1 × $10^{-6}$), showing a lack of weight stability in co-culture in presence of kanamycin.
- Pos 8 PBetI: Weight shifted (Δ=-0.325, $p_{change}$=2.4 × $10^{-8}$), showing a lack of weight stability in co-culture.
- Pos 8 PCin: Inconclusive data (Δ=-0.028, tol=0.0117, $p_{change}$=0.124, $p_{stable}$=0.886), showing a lack of weight stability in co-culture.
- Pos 9 PSal: Inconclusive data (Δ=-0.051, tol=0.0126, $p_{change}$=0.459, $p_{stable}$=0.715), showing a lack of weight stability in co-culture.
- Pos 9 PTet: Weight stabilized (Δ=0.002, tol=0.0757, $p_{stable}$=4.5 × $10^{-10}$), showing weight stability in co-culture.

Summary of results: Co-culture stability (no Kan): Pos 3 (PTtg, Δ=-0.001, tol=0.054, $p_{change}$=0.827, $p_{stable}$=5.0 × $10^{-5}$), Pos 9 (PTet, Δ=0.002, tol=0.0757, $p_{change}$=0.075, $p_{stable}$=4.5 × $10^{-10}$); Co-culture stability (with Kan): Pos 2 (PVan, Δ=-0.003, tol=0.0284, $p_{change}$=0.467, $p_{stable}$=7.8 × $10^{-4}$); Co-culture learning success: Pos 2 (PTac, Δ=-0.074, tol=0.0317, $p_{change}$=2.0 × $10^{-9}$, $p_{stable}$=1.000), Pos 4 (PLux, Δ=-0.204, tol=0.0192, $p_{change}$=3.7 × $10^{-7}$, $p_{stable}$=1.000), Pos 6 (PVan, Δ=-0.247, tol=0.0284, $p_{change}$=7.3 × $10^{-9}$, $p_{stable}$=1.000), Pos 7 (PBetI, Δ=-0.531, tol=0.139, $p_{change}$=2.2 × $10^{-9}$, $p_{stable}$=1.000).

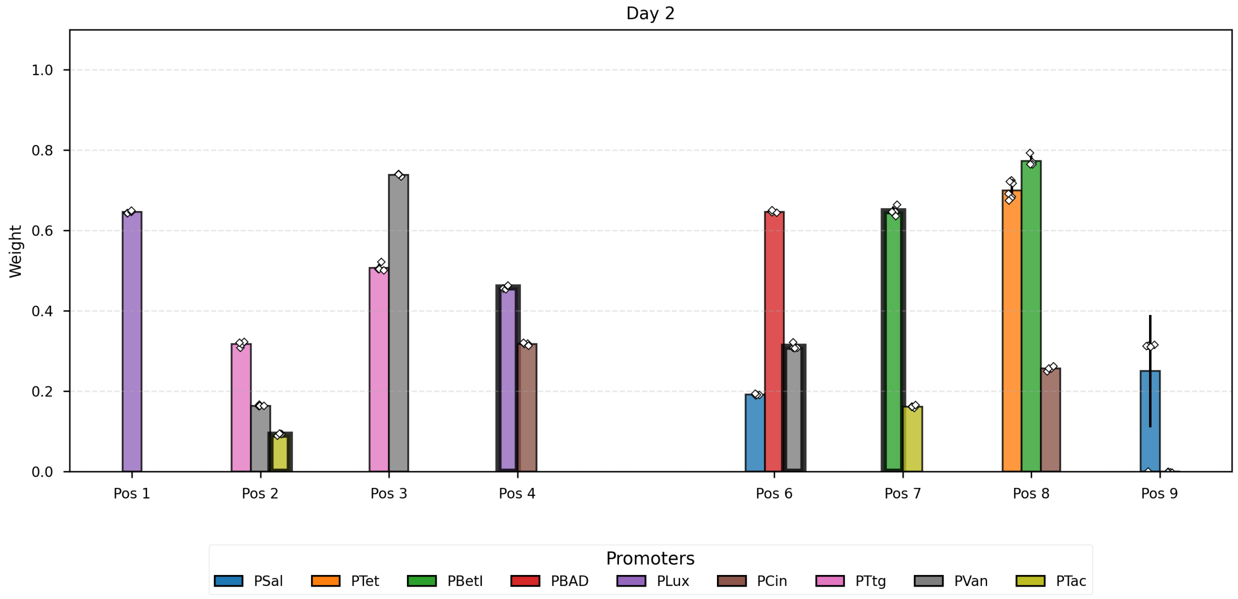

**Supplementary Fig. 74 | Analysis of growth and fluorescence curves from plate reader.**

#### Day 3

Visual Guide: The bars corresponding to Pos 1 (PLux), Pos 6 (PBAD), Pos 8 (PBetI), Pos 9 (PTet) are drawn with thicker contours to indicate they were targets of learning in the previous step.
Analysis of Day 3 vs Day 2:

- Pos 1 PLux: Weight shifted ($\Delta$=-0.097, $p_{change}=3.2 \times 10^{-8}$), showing a negative weight update due to kanamycin learning.
- Pos 2 PTtg: Weight stabilized ($\Delta$=-0.025, tol=0.054, $p_{stable}=3.6 \times 10^{-4}$), showing weight stability in co-culture.
- Pos 2 PVan: Weight shifted ($\Delta$=-0.053, $p_{change}=5.6 \times 10^{-5}$), showing a lack of weight stability in co-culture.
- Pos 2 PTac: Weight stabilized ($\Delta$=0.002, tol=0.0317, $p_{stable}=5.1 \times 10^{-6}$), showing weight stability in co-culture.
- Pos 3 PTtg: Weight shifted ($\Delta$=-0.086, $p_{change}=4.5 \times 10^{-6}$), showing a lack of weight stability in co-culture.
- Pos 3 PVan: Weight shifted ($\Delta$=-0.098, $p_{change}=2.8 \times 10^{-4}$), showing a lack of weight stability in co-culture.
- Pos 4 PLux: Weight stabilized ($\Delta$=0.009, tol=0.0192, $p_{stable}$=0.020), showing weight stability in co-culture.
- Pos 4 PCin: Weight shifted ($\Delta$=0.023, $p_{change}$=0.004), showing a lack of weight stability in co-culture.
- Pos 6 PSal: Weight shifted ($\Delta$=-0.067, $p_{change}=7.1 \times 10^{-8}$), showing a lack of weight stability in co-culture in presence of kanamycin.
- Pos 6 PBAD: Weight shifted ($\Delta$=-0.157, $p_{change}=2.8 \times 10^{-6}$), showing a negative weight update due to kanamycin learning.
- Pos 6 PVan: Weight shifted ($\Delta$=-0.092, $p_{change}=1.8 \times 10^{-6}$), showing a lack of weight stability in co-culture in presence of kanamycin.
- Pos 7 PBetI: Weight shifted ($\Delta$=-0.318, $p_{change}=7.7 \times 10^{-9}$), showing a lack of weight stability in co-culture.
- Pos 7 PTac: Weight stabilized ($\Delta$=0.018, tol=0.0317, $p_{stable}=3.1 \times 10^{-4}$), showing weight stability in co-culture.
- Pos 8 PTet: Weight shifted ($\Delta$=-0.694, $p_{change}=6.6 \times 10^{-7}$), showing a lack of weight stability in co-culture in presence of kanamycin.
- Pos 8 PBetI: Weight shifted ($\Delta$=-0.773, $p_{change}=4.0 \times 10^{-11}$), showing a negative weight update due to kanamycin learning.
- Pos 8 PCin: Weight shifted ($\Delta$=-0.213, $p_{change}=2.4 \times 10^{-6}$), showing a lack of weight stability in co-culture in presence of kanamycin.
- Pos 9 PSal: Weight shifted ($\Delta$=-0.193, $p_{change}$=0.037), showing a lack of weight stability in co-culture in presence of kanamycin.
- Pos 9 PTet: Weight stabilized ($\Delta$=-0.006, tol=0.0757, $p_{stable}=2.7 \times 10^{-20}$), showing a lack of negative weight update as a result of the kanamycin learning.

Summary of results: Single-culture learning: Pos 1 (PLux, Δ=-0.097, tol=0.0192, $p_{change}$=3.2 × $10^{-8}$, $p_{stable}$=1.000); Co-culture stability (no Kan): Pos 2 (PTtg, Δ=-0.025, tol=0.054, $p_{change}$=0.001, $p_{stable}$=3.6 × $10^{-4}$), Pos 2 (PTac, Δ=0.002, tol=0.0317, $p_{change}$=0.373, $p_{stable}$=5.1 × $10^{-6}$), Pos 4 (PLux, Δ=0.009, tol=0.0192, $p_{change}$=0.060, $p_{stable}$=0.020), Pos 7 (PTac, Δ=0.018, tol=0.0317, $p_{change}$=1.1 × $10^{-4}$, $p_{stable}$=3.1 × $10^{-4}$); Co-culture learning success: Pos 6 (PBAD, Δ=-0.157, tol=0.0878, $p_{change}$=2.8 × $10^{-6}$, $p_{stable}$=1.000), Pos 8 (PBetI, Δ=-0.773, tol=0.139, $p_{change}$=4.0 × $10^{-11}$, $p_{stable}$=1.000).

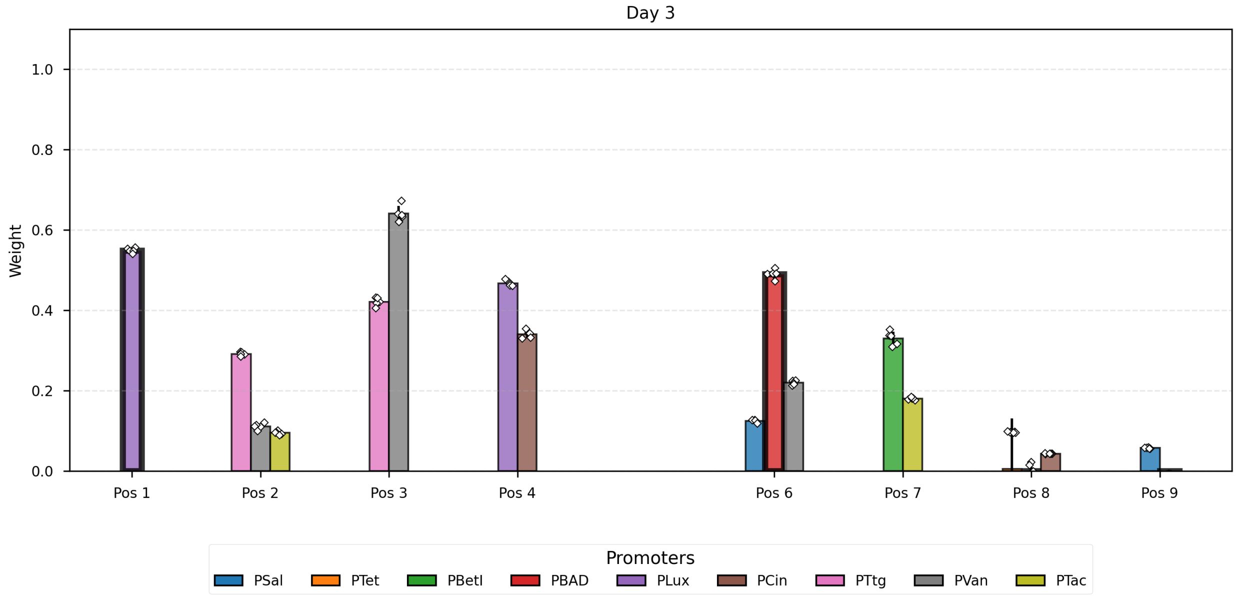

**Supplementary Fig. 75 | Summary of results: Single-culture learning: Pos 1 (PLux, Δ=-0.097, tol=0.0192, pchange=3.2 × 10-8, pstable=1.000); Co-culture stability (no Kan): Pos 2 (PTtg, Δ=-0.02.**

#### Day 4

Visual Guide: The bars corresponding to Pos 2 (PTac), Pos 4 (PLux), Pos 6 (PVan), Pos 7 (PBetI) are drawn with thicker contours to indicate they were targets of learning in the previous step.

Analysis of Day 4 vs Day 3:

- Pos 1 PLux: Weight shifted (Δ=0.018, $p_{change}$=0.001), showing a lack of weight stability in mono-culture.
- Pos 2 PTtg: Weight shifted (Δ=-0.072, $p_{change}$=4.5 × $10^{-4}$), showing a lack of weight stability in co-culture in presence of kanamycin.
- Pos 2 PVan: Weight stabilized (Δ=-0.019, tol=0.0284, $p_{stable}$=0.031), showing **weight stability in co-culture in presence of kanamycin.**
- Pos 2 PTac: Weight shifted (Δ=-0.061, $p_{change}$=1.6 × $10^{-7}$), showing a negative weight update due to kanamycin learning.
- Pos 3 PTtg: Weight shifted (Δ=-0.097, $p_{change}$=6.2 × $10^{-5}$), showing a lack of weight stability in co-culture.
- Pos 3 PVan: Weight shifted (Δ=-0.057, $p_{change}$=0.002), showing a lack of weight stability in co-culture.
- Pos 4 PLux: Weight shifted (Δ=-0.102, $p_{change}$=7.3 × $10^{-8}$), showing a negative weight update due to kanamycin learning.
- Pos 4 PCin: Inconclusive data (Δ=0.009, tol=0.0117, $p_{change}$=0.118, $p_{stable}$=0.263), showing a lack of weight stability in co-culture in presence of kanamycin.

- Pos 6 PSal: Weight shifted (Δ=-0.012, $p_{change}$=8.9 × $10^{-4}$), showing a lack of weight stability in co-culture in presence of kanamycin.
- Pos 6 PBAD: Weight stabilized (Δ=-0.024, tol=0.0878, $p_{stable}$=8.6 × $10^{-6}$), showing **weight stability in co-culture in presence of kanamycin.**
- Pos 6 PVan: Weight shifted (Δ=-0.053, $p_{change}$=6.8 × $10^{-7}$), showing a negative weight update due to kanamycin learning.
- Pos 7 PBetI: Weight stabilized (Δ=-0.073, tol=0.139, $p_{stable}$=1.2 × $10^{-4}$), showing a lack of negative weight update as a result of the kanamycin learning.
- Pos 7 PTac: Weight shifted (Δ=-0.036, $p_{change}$=1.4 × $10^{-7}$), showing a lack of weight stability in co-culture in presence of kanamycin.
- Pos 8 PTet: Inconclusive data (Δ=0.103, tol=0.0757, $p_{change}$=0.055, $p_{stable}$=0.717), showing a lack of weight stability in co-culture.
- Pos 8 PBetI: Weight stabilized (Δ=-0.006, tol=0.139, $p_{stable}$=6.1 × $10^{-6}$), showing weight stability in co-culture.
- Pos 8 PCin: Weight stabilized (Δ=0.001, tol=0.0117, $p_{stable}$=4.5 × $10^{-8}$), showing weight stability in co-culture.
- Pos 9 PSal: Weight stabilized (Δ=0.009, tol=0.0126, $p_{stable}$=0.007), showing weight stability in co-culture.
- Pos 9 PTet: Weight stabilized (Δ=-0.002, tol=0.0757, $p_{stable}$=8.8 × $10^{-16}$), showing weight stability in co-culture.

Summary of results: Co-culture stability (no Kan): Pos 8 (PBetI, Δ=-0.006, tol=0.139, $p_{change}$=0.649, $p_{stable}$=6.1 × $10^{-6}$), Pos 8 (PCin, Δ=0.001, tol=0.0117, $p_{change}$=0.414, $p_{stable}$=4.5 × $10^{-8}$), Pos 9 (PSal, Δ=0.009, tol=0.0126, $p_{change}$=2.1 × $10^{-5}$, $p_{stable}$=0.007), Pos 9 (PTet, Δ=-0.002, tol=0.0757, $p_{change}$=0.015, $p_{stable}$=8.8 × $10^{-16}$); Co-culture stability (with Kan): Pos 2 (PVan, Δ=-0.019, tol=0.0284, $p_{change}$=0.002, $p_{stable}$=0.031), Pos 6 (PBAD, Δ=-0.024, tol=0.0878, $p_{change}$=0.008, $p_{stable}$=8.6 × $10^{-6}$); Co-culture learning success: Pos 2 (PTac, Δ=-0.061, tol=0.0317, $p_{change}$=1.6 × $10^{-7}$, $p_{stable}$=1.000), Pos 4 (PLux, Δ=-0.102, tol=0.0192, $p_{change}$=7.3 × $10^{-8}$, $p_{stable}$=1.000), Pos 6 (PVan, Δ=-0.053, tol=0.0284, $p_{change}$=6.8 × $10^{-7}$, $p_{stable}$=1.000).

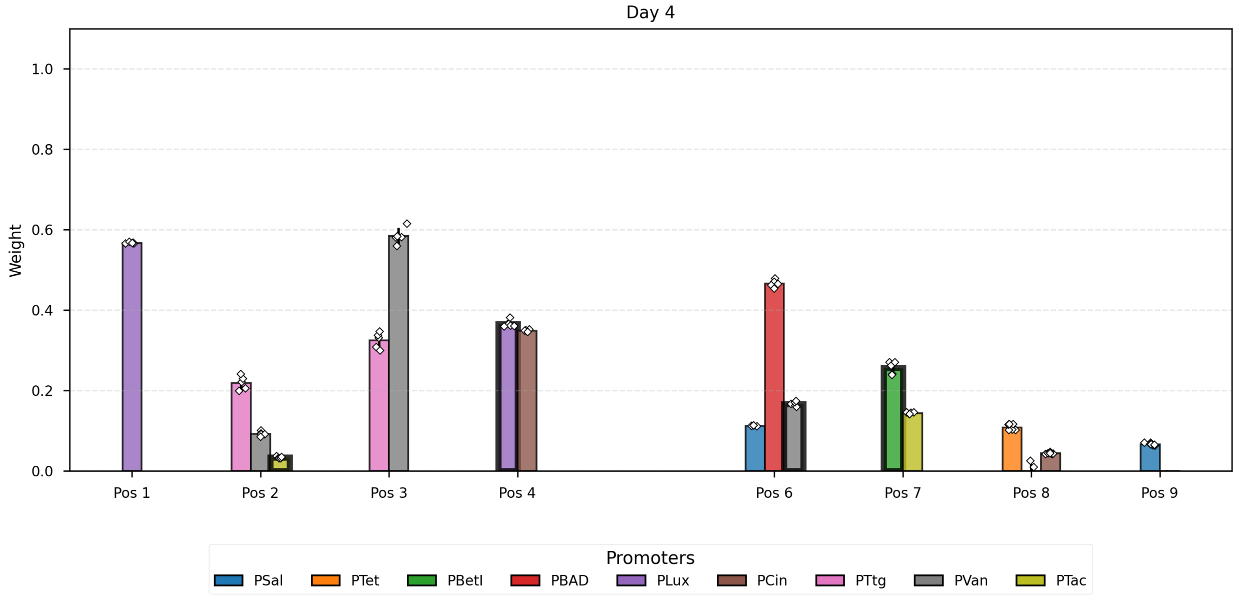

**Supplementary Fig. 76 | Summary of results: Co-culture stability (no Kan): Pos 8 (PBetI, Δ=-0.006, tol=0.139, pchange=0.649, pstable=6.1 × 10-6), Pos 8 (PCin, Δ=0.001, tol=0.0117, pchange=0.4.**

#### Day 5

Visual Guide: The bars corresponding to Pos 1 (PLux), Pos 6 (PBAD), Pos 8 (PBetI), Pos 9 (PTet) are drawn with thicker contours to indicate they were targets of learning in the previous step.
Analysis of Day 5 vs Day 4:

- Pos 1 PLux: Weight shifted (Δ=-0.048, $p_{change}$=5.4 × $10^{-11}$), showing a negative weight update due to kanamycin learning.
- Pos 2 PTtg: Weight stabilized (Δ=0.010, tol=0.054, $p_{stable}$=0.002), showing weight stability in co-culture.
- Pos 2 PVan: Weight stabilized (Δ=0.006, tol=0.0284, $p_{stable}$=7.6 × $10^{-5}$), showing weight stability in co-culture.
- Pos 2 PTac: Weight stabilized (Δ=0.009, tol=0.0317, $p_{stable}$=1.6 × $10^{-7}$), showing weight stability in co-culture.
- Pos 3 PTtg: Weight shifted (Δ=0.109, $p_{change}$=1.7 × $10^{-5}$), showing a lack of weight stability in co-culture.
- Pos 3 PVan: Weight shifted (Δ=0.074, $p_{change}$=3.3 × $10^{-4}$), showing a lack of weight stability in co-culture.
- Pos 4 PLux: Inconclusive data (Δ=0.015, tol=0.0192, $p_{change}$=0.076, $p_{stable}$=0.276), showing a lack of weight stability in co-culture.
- Pos 4 PCin: Weight shifted (Δ=0.014, $p_{change}$=0.005), showing a lack of weight stability in co-culture.
- Pos 6 PSal: Weight shifted (Δ=-0.020, $p_{change}$=1.5 × $10^{-4}$), showing a lack of weight stability in co-culture in presence of kanamycin.
- Pos 6 PBAD: Weight shifted (Δ=-0.088, $p_{change}$=7.7 × $10^{-7}$), showing a negative weight update due to kanamycin learning.
- Pos 6 PVan: Weight shifted (Δ=0.036, $p_{change}$=1.3 × $10^{-5}$), showing a lack of weight stability in co-culture in presence of kanamycin.
- Pos 7 PBetI: Weight stabilized (Δ=-0.044, tol=0.139, $p_{stable}$=2.0 × $10^{-4}$), showing weight stability in co-culture.
- Pos 7 PTac: Weight stabilized (Δ=0.014, tol=0.0317, $p_{stable}$=1.7 × $10^{-5}$), showing weight stability in co-culture.
- Pos 8 PTet: Weight stabilized (Δ=-0.060, tol=0.0757, $p_{stable}$=1.8 × $10^{-4}$), showing **weight stability in co-culture in presence of kanamycin.**
- Pos 8 PBetI: Weight shifted (Δ=-0.142, $p_{change}$=0.036), showing a negative weight update due to kanamycin learning.
- Pos 8 PCin: Weight shifted (Δ=-0.026, $p_{change}$=3.3 × $10^{-9}$), showing a lack of weight stability in co-culture in presence of kanamycin.
- Pos 9 PSal: Weight shifted (Δ=-0.018, $p_{change}$=4.5 × $10^{-9}$), showing a lack of weight stability in co-culture in presence of kanamycin.
- Pos 9 PTet: Weight stabilized (Δ=0.006, tol=0.0757, $p_{stable}$=1.7 × $10^{-12}$), showing a lack of negative weight update as a result of the kanamycin learning.

Summary of results: Single-culture learning: Pos 1 (PLux, Δ=-0.048, tol=0.0192, $p_{change}$=5.4 × $10^{-11}$, $p_{stable}$=1.000); Co-culture stability (no Kan): Pos 2 (PTtg, Δ=0.010, tol=0.054, $p_{change}$=0.370, $p_{stable}$=0.002), Pos 2 (PVan, Δ=0.006, tol=0.0284, $p_{change}$=0.083, $p_{stable}$=7.6 × $10^{-5}$), Pos 2 (PTac, Δ=0.009, tol=0.0317, $p_{change}$=2.9 × $10^{-4}$, $p_{stable}$=1.6 × $10^{-7}$), Pos 7 (PBetI, Δ=-0.044, tol=0.139, $p_{change}$=0.019, $p_{stable}$=2.0 × $10^{-4}$), Pos 7 (PTac, Δ=0.014, tol=0.0317, $p_{change}$=2.0 × $10^{-4}$, $p_{stable}$=1.7 × $10^{-5}$); Co-culture stability (with Kan): Pos 8 (PTet, Δ=-0.060, tol=0.0757, $p_{change}$=1.7 × $10^{-8}$, $p_{stable}$=1.8 × $10^{-4}$); Co-culture learning success: Pos 6 (PBAD, Δ=-0.088, tol=0.0878, $p_{change}$=7.7 × $10^{-7}$, $p_{stable}$=0.539), Pos 8 (PBetI, Δ=-0.142, tol=0.139, $p_{change}$=0.036, $p_{stable}$=0.530).

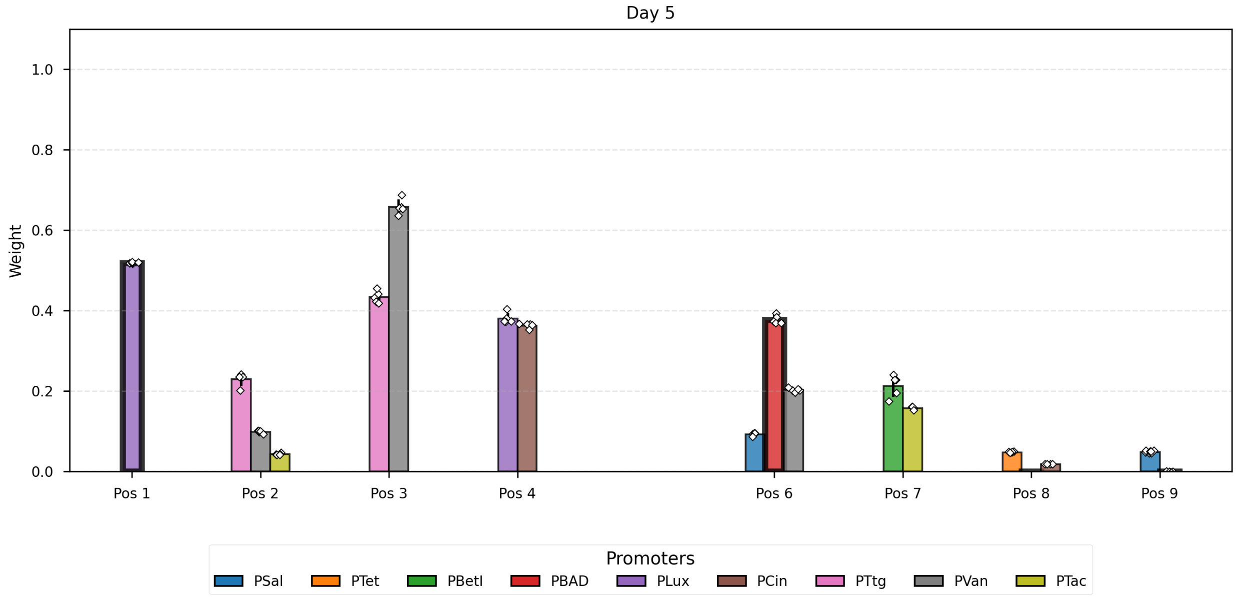

**Supplementary Fig. 77 | Summary of results: Single-culture learning: Pos 1 (PLux, Δ=-0.048, tol=0.0192, pchange=5.4 × 10-11, pstable=1.000); Co-culture stability (no Kan): Pos 2 (PTtg, Δ=0.01.**

#### Day 6

Visual Guide: The bars corresponding to Pos 2 (PTac), Pos 4 (PLux), Pos 6 (PVan), Pos 7 (PBetI) are drawn with thicker contours to indicate they were targets of learning in the previous step.
Analysis of Day 6 vs Day 5:

- Pos 1 PLux: Weight shifted (Δ=0.023, $p_{change}$=3.3 × $10^{-9}$), showing a lack of weight stability in mono-culture.
- Pos 2 PTtg: Weight shifted (Δ=-0.214, $p_{change}$=6.2 × $10^{-8}$), showing a lack of weight stability in co-culture in presence of kanamycin.
- Pos 2 PVan: Weight shifted (Δ=-0.053, $p_{change}$=3.6 × $10^{-8}$), showing a lack of weight stability in co-culture in presence of kanamycin.
- Pos 2 PTac: Weight shifted (Δ=-0.053, $p_{change}$=2.5 × $10^{-9}$), showing a negative weight update due to kanamycin learning.
- Pos 3 PTtg: Weight shifted (Δ=0.045, $p_{change}$=7.5 × $10^{-4}$), showing a lack of weight stability in co-culture.
- Pos 3 PVan: Weight shifted (Δ=-0.052, $p_{change}$=0.009), showing a lack of weight stability in co-culture.
- Pos 4 PLux: Weight shifted (Δ=-0.054, $p_{change}$=1.6 × $10^{-4}$), showing a negative weight update due to kanamycin learning.

- Pos 4 PCin: Weight shifted (Δ=-0.023, $p_{change}$=0.032), showing a lack of weight stability in co-culture in presence of kanamycin.
- Pos 6 PSal: Weight shifted (Δ=-0.026, $p_{change}$=$1.4 \times 10^{-7}$), showing a lack of weight stability in co-culture in presence of kanamycin.
- Pos 6 PBAD: Weight shifted (Δ=-0.114, $p_{change}$=$2.6 \times 10^{-5}$), showing a lack of weight stability in co-culture in presence of kanamycin.
- Pos 6 PVan: Weight shifted (Δ=-0.131, $p_{change}$=$6.2 \times 10^{-18}$), showing a negative weight update due to kanamycin learning.
- Pos 7 PBetI: Weight shifted (Δ=-0.123, $p_{change}$=0.012), showing a negative weight update due to kanamycin learning.
- Pos 7 PTac: Weight shifted (Δ=-0.059, $p_{change}$=$6.6 \times 10^{-11}$), showing a lack of weight stability in co-culture in presence of kanamycin.
- Pos 8 PTet: Weight stabilized (Δ=0.007, tol=0.0757, $p_{stable}$=$2.0 \times 10^{-9}$), showing weight stability in co-culture.
- Pos 8 PBetI: Inconclusive data (Δ=0.090, tol=0.139, $p_{change}$=0.109, $p_{stable}$=0.166), showing a lack of weight stability in co-culture.
- Pos 8 PCin: Weight stabilized (Δ=-0.004, tol=0.0117, $p_{stable}$=$5.5 \times 10^{-5}$), showing weight stability in co-culture.
- Pos 9 PSal: Weight stabilized (Δ=0.006, tol=0.0126, $p_{stable}$=0.002), showing weight stability in co-culture.
- Pos 9 PTet: Weight stabilized (Δ=-0.007, tol=0.0757, $p_{stable}$=$8.3 \times 10^{-11}$), showing weight stability in co-culture.

Summary of results: Co-culture stability (no Kan): Pos 8 (PTet, Δ=0.007, tol=0.0757, $p_{change}$=0.002, $p_{stable}$=$2.0 \times 10^{-9}$), Pos 8 (PCin, Δ=-0.004, tol=0.0117, $p_{change}$=0.009, $p_{stable}$=$5.5 \times 10^{-5}$), Pos 9 (PSal, Δ=0.006, tol=0.0126, $p_{change}$=0.005, $p_{stable}$=0.002), Pos 9 (PTet, Δ=-0.007, tol=0.0757, $p_{change}$=0.002, $p_{stable}$=$8.3 \times 10^{-11}$); Co-culture learning success: Pos 2 (PTac, Δ=-0.053, tol=0.0317, $p_{change}$=$2.5 \times 10^{-9}$, $p_{stable}$=1.000), Pos 4 (PLux, Δ=-0.054, tol=0.0192, $p_{change}$=$1.6 \times 10^{-4}$, $p_{stable}$=0.999), Pos 6 (PVan, Δ=-0.131, tol=0.0284, $p_{change}$=$6.2 \times 10^{-18}$, $p_{stable}$=1.000), Pos 7 (PBetI, Δ=-0.123, tol=0.139, $p_{change}$=0.012, $p_{stable}$=0.348).

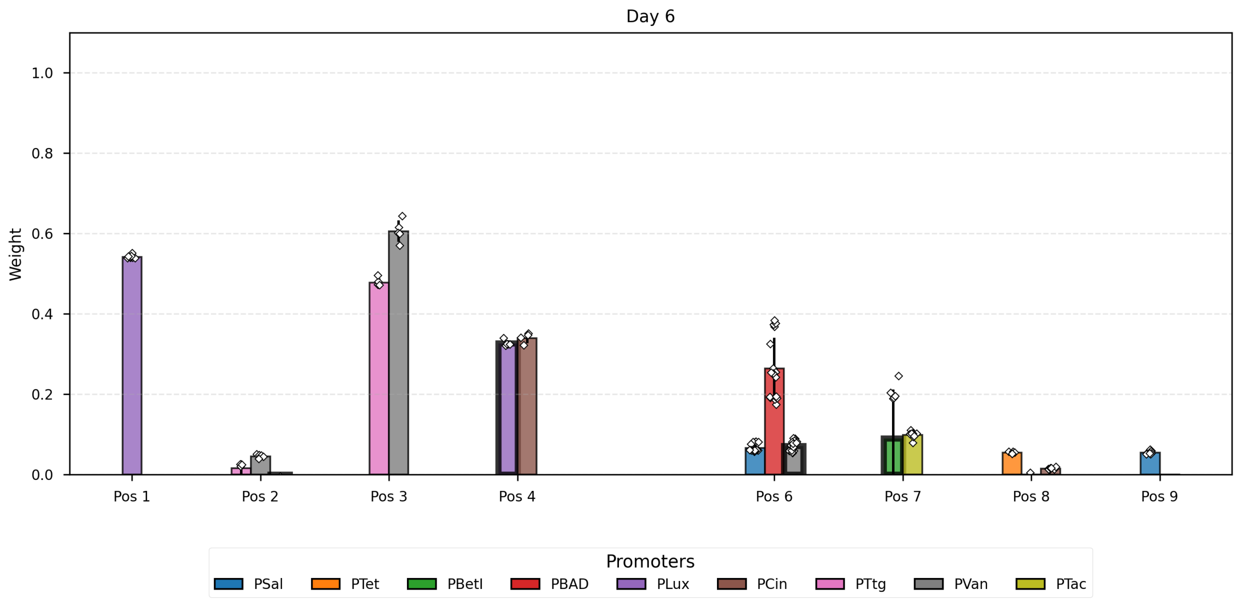

**Supplementary Fig. 78 | Summary of results: Co-culture stability (no Kan): Pos 8 (PTet, Δ=0.007, tol=0.0757, pchange=0.002, pstable=2.0 × 10-9), Pos 8 (PCin, Δ=-0.004, tol=0.0117, pchange=0.0.**

#### Day 7

Visual Guide: The bars corresponding to Pos 1 (PLux), Pos 6 (PBAD), Pos 8 (PBetI), Pos 9 (PTet) are drawn with thicker contours to indicate they were targets of learning in the previous step.

Analysis of Day 7 vs Day 6:

- Pos 1 PLux: Weight shifted (Δ=-0.106, $p_{change}$=2.8 × $10^{-12}$), showing a negative weight update due to kanamycin learning.
- Pos 2 PTtg: Weight shifted (Δ=0.112, $p_{change}$=0.002), showing a lack of weight stability in co-culture.
- Pos 2 PVan: Inconclusive data (Δ=0.026, tol=0.0284, $p_{change}$=0.060, $p_{stable}$=0.413), showing a lack of weight stability in co-culture.
- Pos 2 PTac: Weight stabilized (Δ=0.010, tol=0.0317, $p_{stable}$=2.0 × $10^{-5}$), showing weight stability in co-culture.
- Pos 3 PTtg: Weight stabilized (Δ=-0.027, tol=0.054, $p_{stable}$=0.004), showing weight stability in co-culture.
- Pos 3 PVan: Weight shifted (Δ=0.101, $p_{change}$=2.2 × $10^{-4}$), showing a lack of weight stability in co-culture.
- Pos 4 PLux: Weight shifted (Δ=-0.047, $p_{change}$=1.1 × $10^{-5}$), showing a lack of weight stability in co-culture.
- Pos 4 PCin: Inconclusive data (Δ=0.005, tol=0.0117, $p_{change}$=0.541, $p_{stable}$=0.206), showing a lack of weight stability in co-culture.
- Pos 6 PSal: Weight shifted (Δ=-0.019, $p_{change}$=1.6 × $10^{-5}$), showing a lack of weight stability in co-culture in presence of kanamycin.
- Pos 6 PBAD: Weight shifted (Δ=-0.103, $p_{change}$=8.0 × $10^{-5}$), showing a negative weight update due to kanamycin learning.
- Pos 6 PVan: Weight stabilized (Δ=-0.021, tol=0.0284, $p_{stable}$=0.016), showing **weight stability in co-culture in presence of kanamycin.**

- Pos 7 PBetI: Weight stabilized (Δ=-0.008, tol=0.139, $p_{stable}$=0.004), showing weight stability in co-culture.
- Pos 7 PTac: Weight stabilized (Δ=-0.005, tol=0.0317, $p_{stable}$=0.004), showing weight stability in co-culture.
- Pos 8 PTet: Weight stabilized (Δ=-0.039, tol=0.0757, $p_{stable}$=1.1 × $10^{-8}$), showing **weight stability in co-culture in presence of kanamycin.**
- Pos 8 PBetI: Weight stabilized (Δ=-0.087, tol=0.139, $p_{stable}$=0.007), showing a lack of negative weight update as a result of the kanamycin learning.
- Pos 8 PCin: Weight shifted (Δ=-0.016, $p_{change}$=3.4 × $10^{-5}$), showing a lack of weight stability in co-culture in presence of kanamycin.
- Pos 9 PSal: Weight shifted (Δ=-0.042, $p_{change}$=2.1 × $10^{-8}$), showing a lack of weight stability in co-culture in presence of kanamycin.
- Pos 9 PTet: Weight stabilized (Δ=-0.008, tol=0.0757, $p_{stable}$=1.1 × $10^{-12}$), showing a lack of negative weight update as a result of the kanamycin learning.

Summary of results: Single-culture learning: Pos 1 (PLux, Δ=-0.106, tol=0.0192, $p_{change}$=2.8 × $10^{-12}$, $p_{stable}$=1.000); Co-culture stability (no Kan): Pos 2 (PTac, Δ=0.010, tol=0.0317, $p_{change}$=0.005, $p_{stable}$=2.0 × $10^{-5}$), Pos 3 (PTtg, Δ=-0.027, tol=0.054, $p_{change}$=0.007, $p_{stable}$=0.004), Pos 7 (PBetI, Δ=-0.008, tol=0.139, $p_{change}$=0.842, $p_{stable}$=0.004), Pos 7 (PTac, Δ=-0.005, tol=0.0317, $p_{change}$=0.548, $p_{stable}$=0.004); Co-culture stability (with Kan): Pos 6 (PVan, Δ=-0.021, tol=0.0284, $p_{change}$=4.6 × $10^{-6}$, $p_{stable}$=0.016), Pos 8 (PTet, Δ=-0.039, tol=0.0757, $p_{change}$=1.4 × $10^{-8}$, $p_{stable}$=1.1 × $10^{-8}$); Co-culture learning success: Pos 6 (PBAD, Δ=-0.103, tol=0.0878, $p_{change}$=8.0 × $10^{-5}$, $p_{stable}$=0.775).

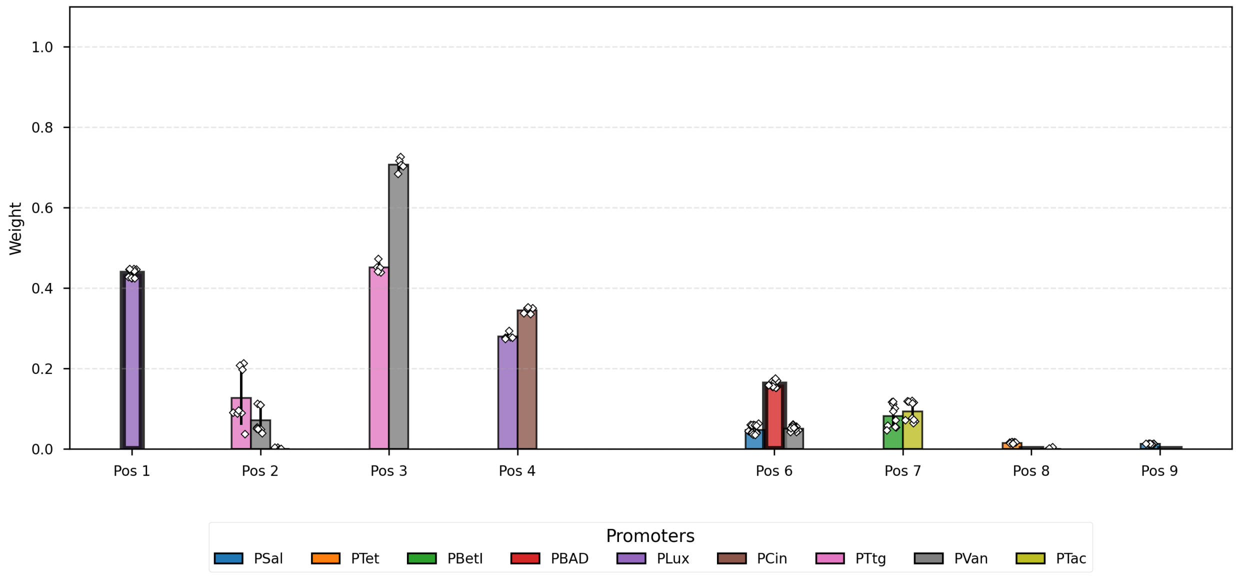

**Supplementary Fig. 79 | Summary of results: Single-culture learning: Pos 1 (PLux, Δ=-0.106, tol=0.0192, pchange=2.8 × 10-12, pstable=1.000); Co-culture stability (no Kan): Pos 2 (PTac, Δ=0.01.**

#### Day 8

Visual Guide: The bars corresponding to Pos 2 (PVan), Pos 3 (PTtg), Pos 4 (PLux), Pos 6 (PSal) are drawn with thicker contours to indicate they were targets of learning in the previous step.
Analysis of Day 8 vs Day 7:

- Pos 1 PLux: Weight shifted ($\Delta$=0.062, $p_{change}$=6.9 × $10^{-7}$), showing a lack of weight stability in mono-culture.
- Pos 2 PTtg: Weight shifted ($\Delta$=-0.086, $p_{change}$=0.010), showing a lack of weight stability in co-culture in presence of kanamycin.
- Pos 2 PVan: Inconclusive data ($\Delta$=-0.020, tol=0.0284, $p_{change}$=0.141, $p_{stable}$=0.271), showing a lack of negative weight update as a result of the kanamycin learning.
- Pos 2 PTac: Weight stabilized ($\Delta$=-0.017, tol=0.0317, $p_{stable}$=3.6 × $10^{-4}$), showing **weight stability in co-culture in presence of kanamycin.**
- Pos 3 PTtg: Weight stabilized ($\Delta$=0.011, tol=0.054, $p_{stable}$=2.8 × $10^{-4}$), showing a lack of negative weight update as a result of the kanamycin learning.
- Pos 3 PVan: Weight stabilized ($\Delta$=0.002, tol=0.0284, $p_{stable}$=0.028), showing **weight stability in co-culture in presence of kanamycin.**
- Pos 4 PLux: Weight shifted ($\Delta$=-0.046, $p_{change}$=1.7 × $10^{-5}$), showing a negative weight update due to kanamycin learning.
- Pos 4 PCin: Weight shifted ($\Delta$=-0.016, $p_{change}$=0.008), showing a lack of weight stability in co-culture in presence of kanamycin.
- Pos 6 PSal: Weight shifted ($\Delta$=-0.031, $p_{change}$=2.7 × $10^{-9}$), showing a negative weight update due to kanamycin learning.
- Pos 6 PBAD: Weight shifted ($\Delta$=0.499, $p_{change}$=8.9 × $10^{-11}$), showing a lack of weight stability in co-culture in presence of kanamycin.
- Pos 6 PVan: Weight shifted ($\Delta$=-0.048, $p_{change}$=3.9 × $10^{-18}$), showing a lack of weight stability in co-culture in presence of kanamycin.
- Pos 7 PBetI: Weight shifted ($\Delta$=0.160, $p_{change}$=0.011), showing a lack of weight stability in co-culture.
- Pos 7 PTac: Weight shifted ($\Delta$=-0.062, $p_{change}$=6.7 × $10^{-4}$), showing a lack of weight stability in co-culture.
- Pos 8 PTet: Weight stabilized ($\Delta$=0.011, tol=0.0757, $p_{stable}$=3.7 × $10^{-12}$), showing weight stability in co-culture.

Summary of results: Co-culture stability (no Kan): Pos 8 (PTet, $\Delta$=0.011, tol=0.0757, $p_{change}$=4.5 × $10^{-5}$, $p_{stable}$=3.7 × $10^{-12}$); Co-culture stability (with Kan): Pos 2 (PTac, $\Delta$=-0.017, tol=0.0317, $p_{change}$=2.5 × $10^{-4}$, $p_{stable}$=3.6 × $10^{-4}$), Pos 3 (PVan, $\Delta$=0.002, tol=0.0284, $p_{change}$=0.848, $p_{stable}$=0.028); Co-culture learning success: Pos 4 (PLux, $\Delta$=-0.046, tol=0.0192, $p_{change}$=1.7 × $10^{-5}$, $p_{stable}$=1.000), Pos 6 (PSal, $\Delta$=-0.031, tol=0.0126, $p_{change}$=2.7 × $10^{-9}$, $p_{stable}$=1.000).

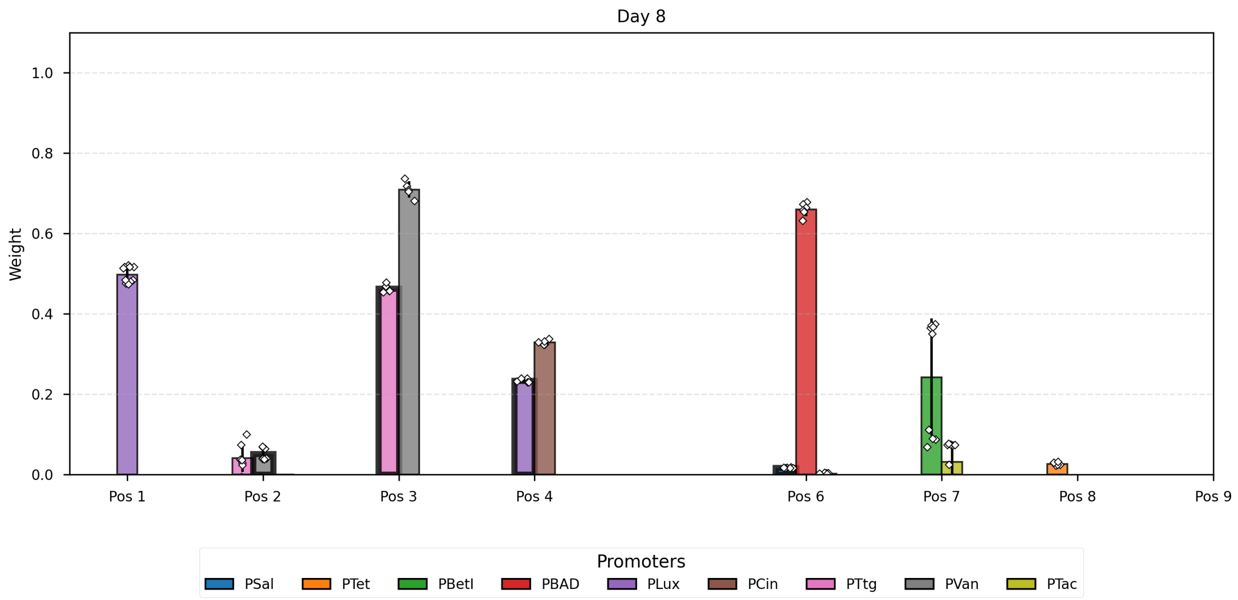

**Supplementary Fig. 80 | Summary of results: Co-culture stability (no Kan): Pos 8 (PTet, Δ=0.011, tol=0.0757, pchange=4.5 × 10-5, pstable=3.7 × 10-12); Co-culture stability (with Kan): Pos 2 (.**

#### Time Course Analysis by Position

Evolution of weights across days. Bars with thicker contours indicate learning events.

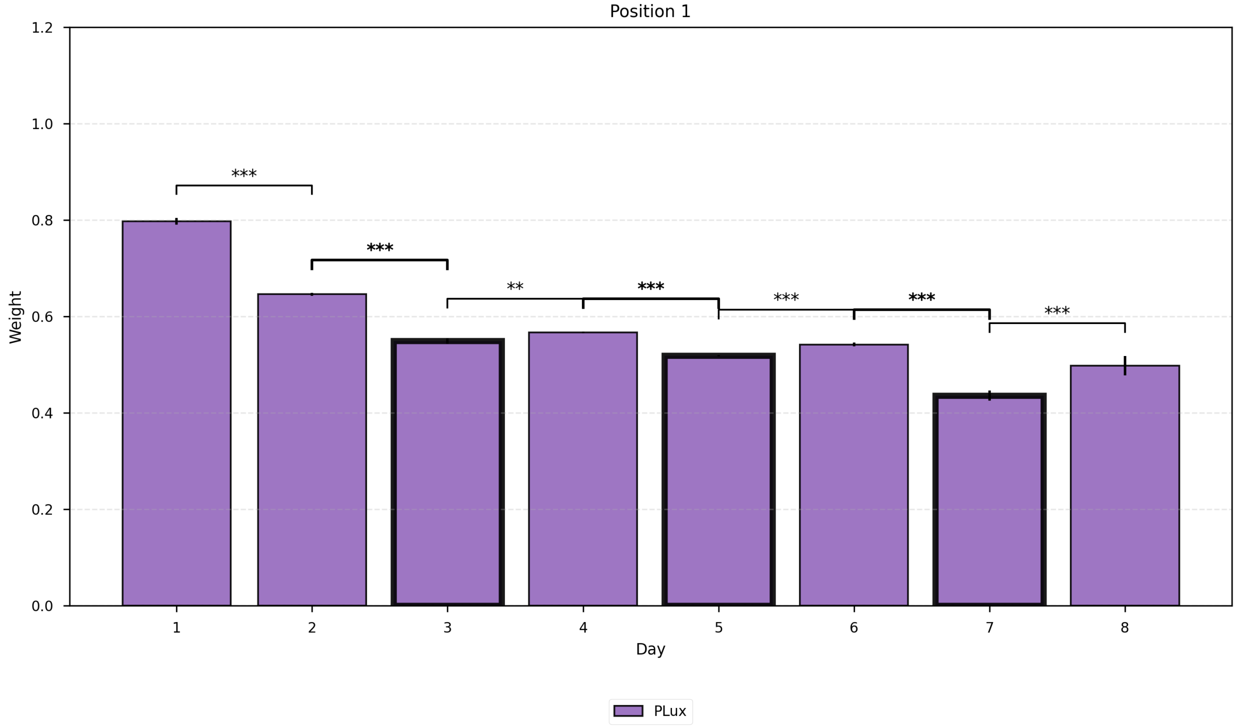

#### Supplementary Fig. 81 | Time Course Analysis by Position.

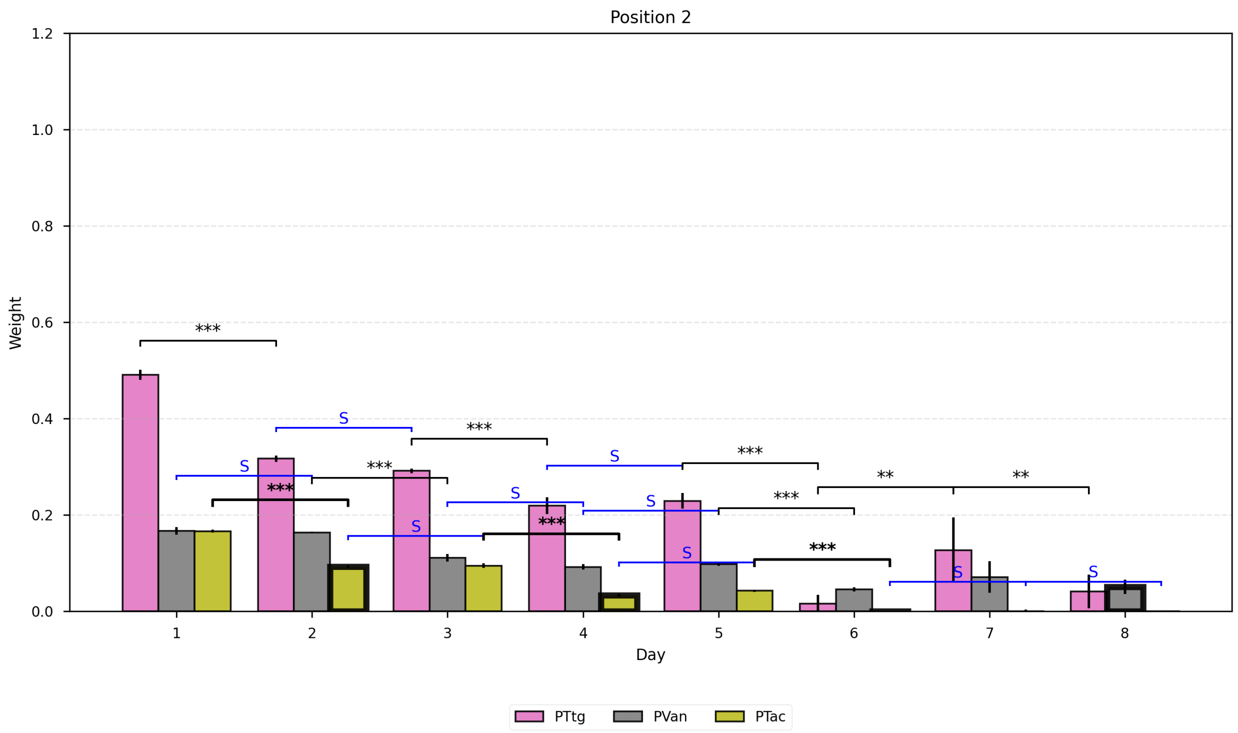

#### Supplementary Fig. 82 | Time Course Analysis by Position.

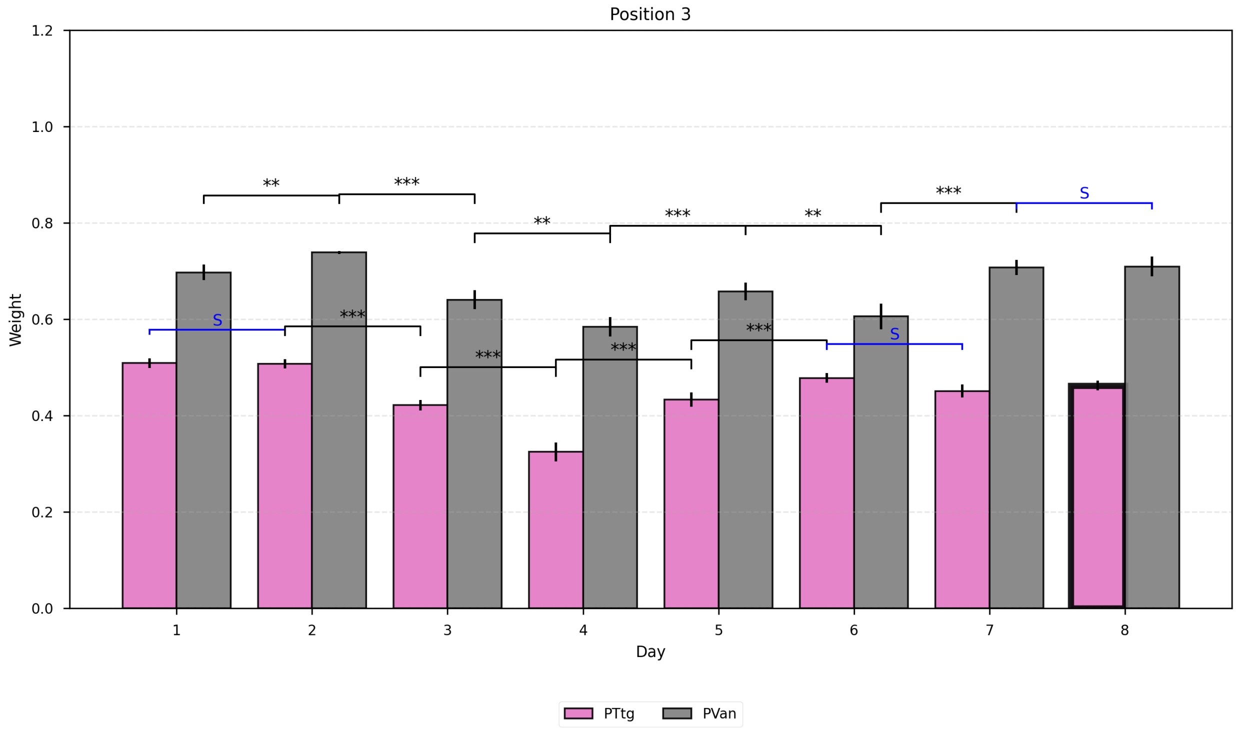

#### Supplementary Fig. 83 | Time Course Analysis by Position.

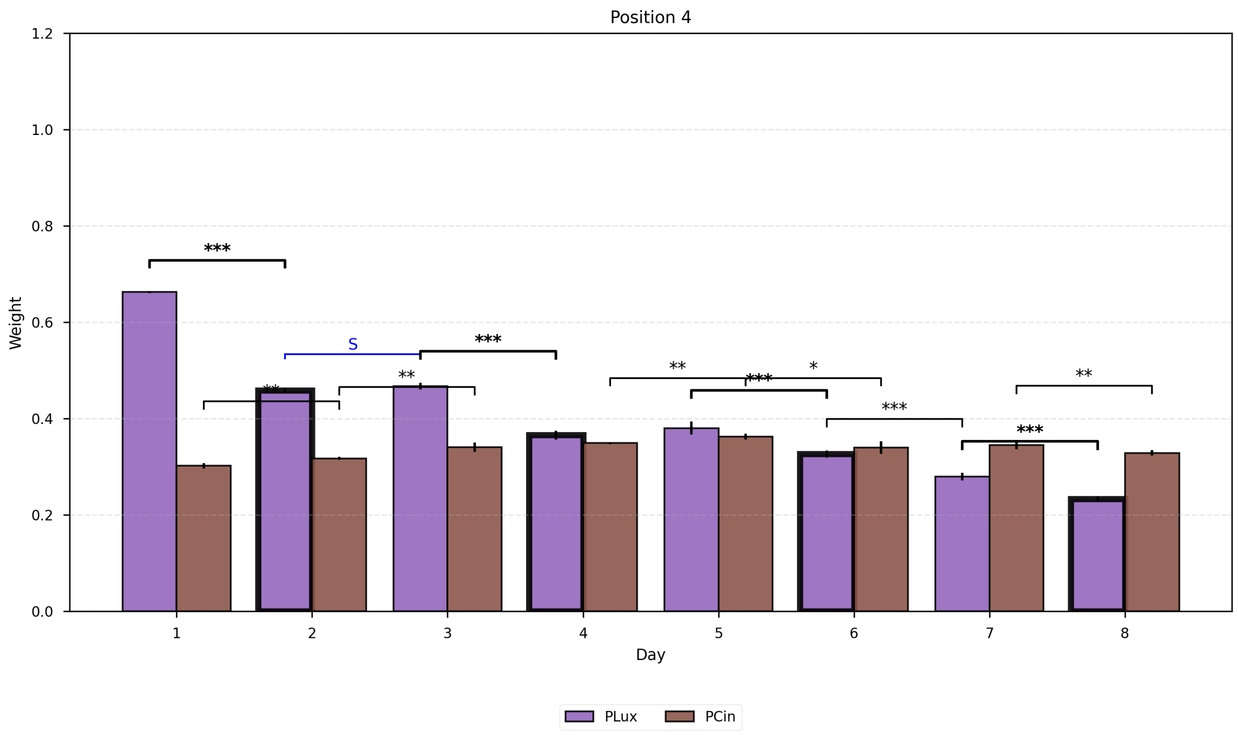

#### Supplementary Fig. 84 | Time Course Analysis by Position.

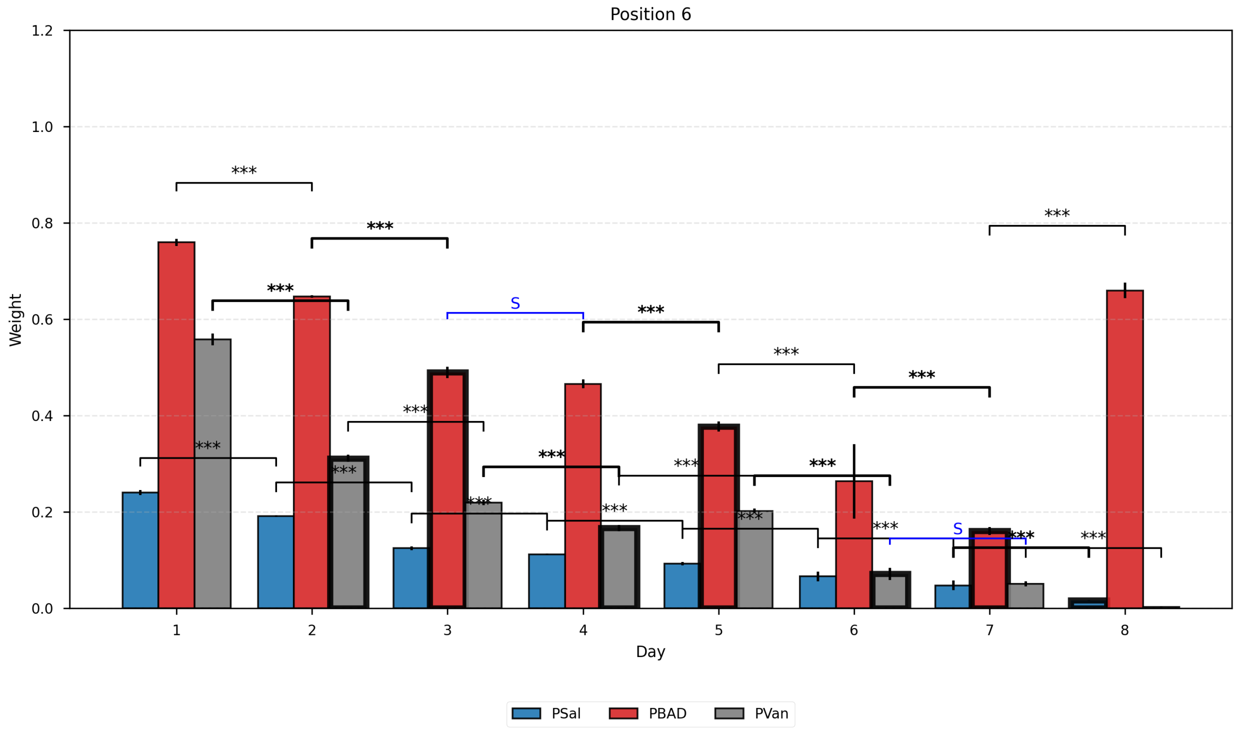

#### Supplementary Fig. 85 | Time Course Analysis by Position.

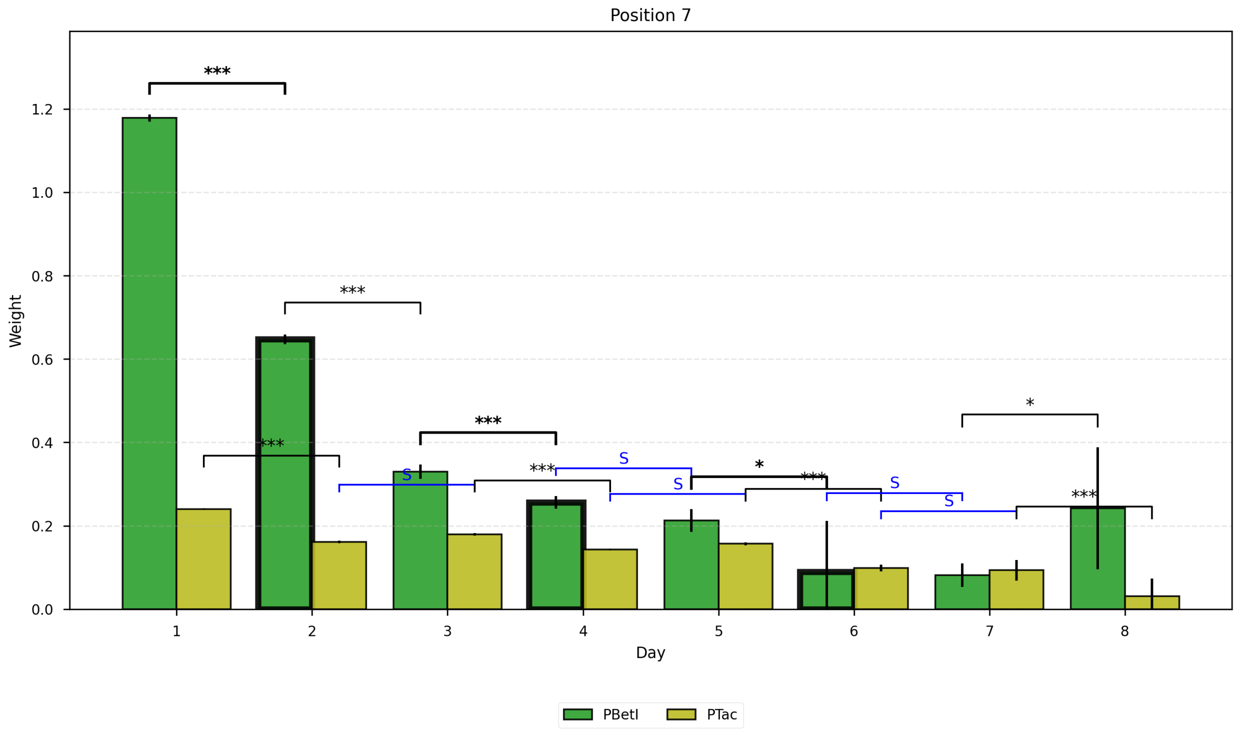

#### Supplementary Fig. 86 | Time Course Analysis by Position.

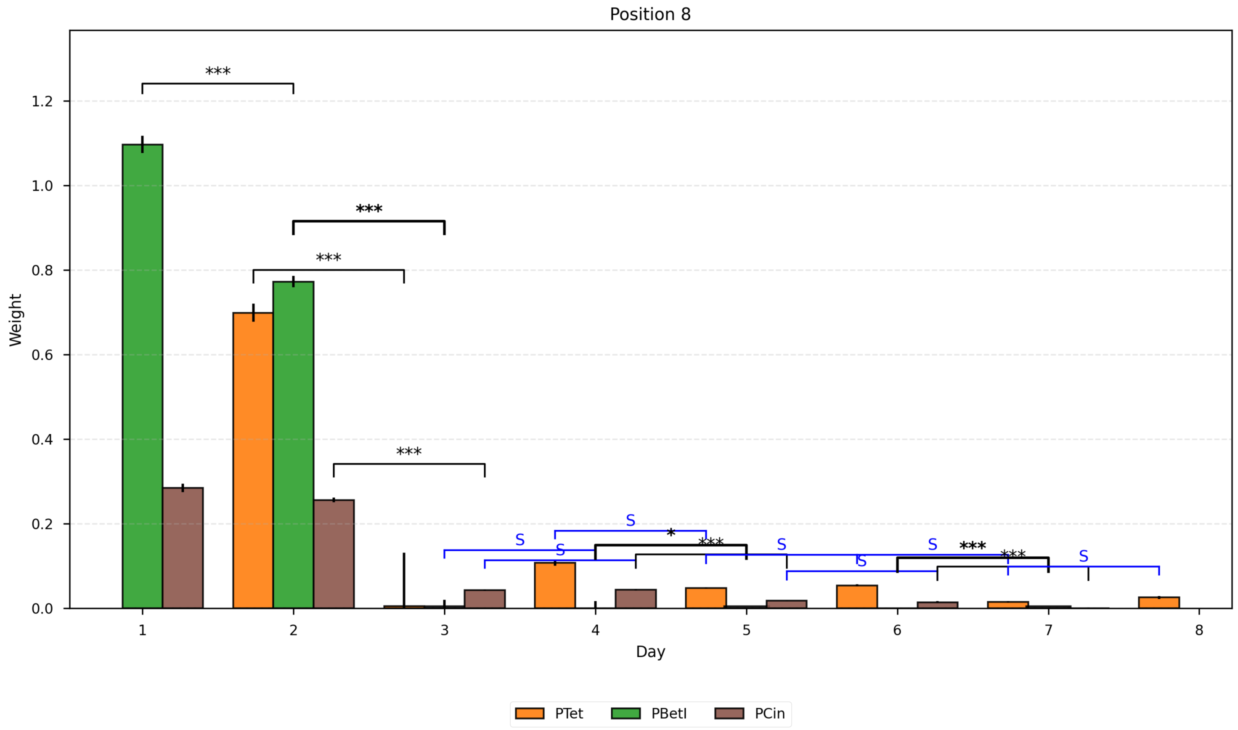

#### Supplementary Fig. 87 | Time Course Analysis by Position.

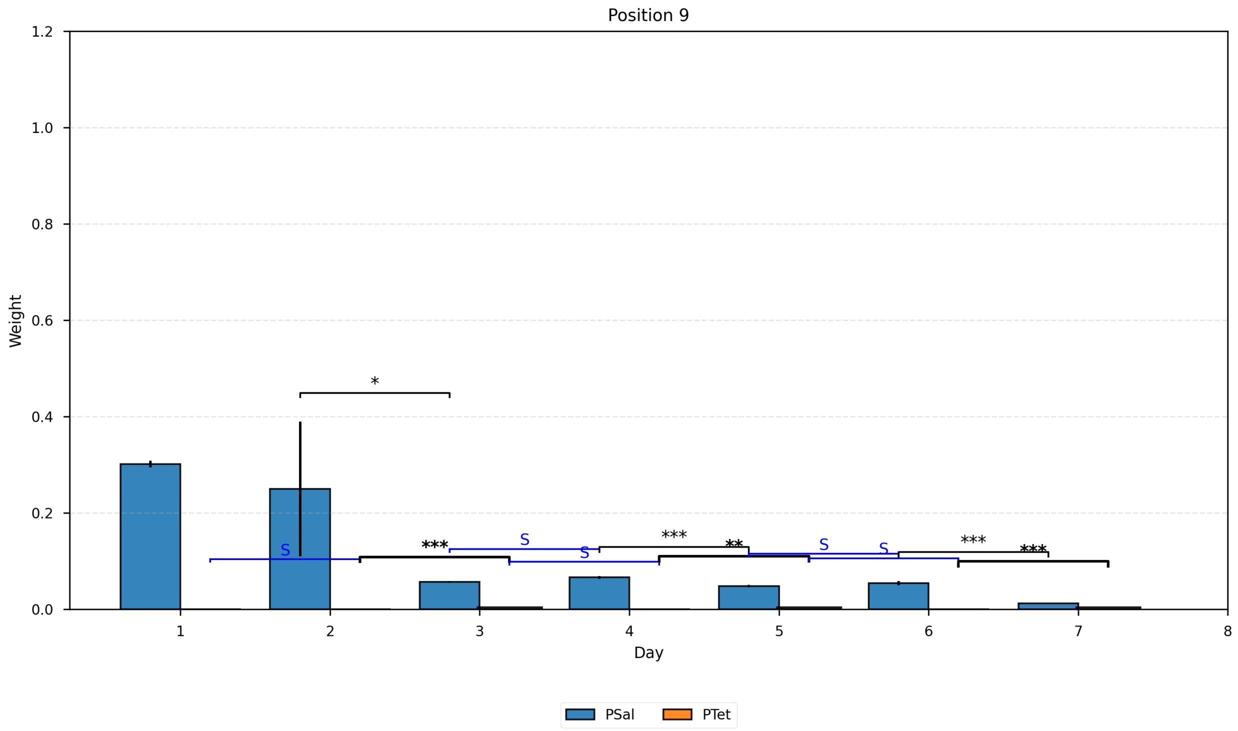

#### Supplementary Fig. 88 | Time Course Analysis by Position.

#### Supplementary Note 12: Simulation framework for one-stage WTA and multilayer XOR with human-in-the-loop signalling and negative local learning

This note describes the simulation framework used to test whether memregulon WTA architectures can implement XOR under biologically meaningful assumptions. The XOR analyses are not presented as reinforcement learning or as a molecular implementation of an RL agent. Instead, they are negative-only pruning and scalability analyses that ask what constrained WTA architectures can do when incorrect outputs trigger local negative updates in active branches. Throughout the framework, synaptic weights are plasmid-fraction variables constrained to the interval [0,1]; inter-layer communication is carried out explicitly by a human operator who reads the state of one co-culture and applies the corresponding inducer to the next layer; learning is implemented as a direct irreversible local negative-update step that only decreases weights; and the uncertainty assigned to each synapse is taken from the experimentally measured promoter-specific weight variability and propagated by Monte Carlo sampling. Within these constraints, the framework yields four concrete outputs. First, a one-stage WTA readout with a physical combinatorial promoter feature can realise XOR. Second, a biologically valid multilayer network with two logical inputs and one bias term can reach XOR by negative-only pruning from a suitable OR-like start. Third, the exact final networks can be drawn directly from the simulated weights and node values. Fourth, propagation of the measured weight uncertainty shows that deterministic XOR correctness does not automatically imply strong statistical separation, particularly in the multilayer case.

In this framework, one co-culture well is modelled as one analogue readout unit for the human-in-the-loop scalability analysis. The neuron does not transmit a digital bit to the next layer. Instead, the operator reads pooled red and pooled green fluorescence, decodes one scalar neuron state, converts that state into an inducer concentration, and then applies that inducer to the next layer. This formulation keeps the biological meaning of the update rule intact while separating it from the current practical limits of inter-layer signalling.

##### Human-in-the-loop signalling

In the human-in-the-loop protocol, the operator performs the inter-layer communication step explicitly. Each co-culture contains a mixture of strains carrying different memregulons, which form the incoming channels of that neuron. During inference, the operator applies inducer concentrations to each co-culture, measures the pooled red and pooled green fluorescence from that co-culture, computes the co-culture output, converts that output into an inducer concentration, and then applies that inducer concentration to the relevant co-cultures in the next layer.

This separation decouples two distinct problems:

1. **Learning**: changes in plasmid fractions inside each co-culture.

2. **Communication**: routing one neuron's output to the correct co-cultures in the next layer.

This abstraction also maps naturally onto future biological implementations. For example, if one upstream layer contains co-cultures 1 and 2, and a downstream layer contains co-cultures A, B, and C, then two orthogonal quorum-sensing channels could replace the human routing step. The OC6 and OHC14 AHL-responsive channels characterised in this work illustrate this type of local activation input, although we did not implement autonomous cell-cell routing during learning. Co-culture 1 could produce signalling molecule 1, and A, B, and C could use that molecule to activate the strains carrying incoming channel 1. Co-culture 2 could do the same for signalling molecule 2. Diffusion, degradation, and dilution would then act as connection-efficiency losses. In the simulator, these losses can be absorbed into the mapping between a neuron's computed output and the signal level experienced by the next layer.

**Physical definition of a weight and of one neuron**

For one incoming channel $k$ in one co-culture, the synaptic weight is constrained by

$$0 \leq w_k \leq 1.$$

Biologically, $w_k$ is the fraction of that incoming plasmid population that is in the red-memory state rather than the green-memory state. Because this variable is a plasmid fraction, the simulator clips every weight to [0,1].

One neuron corresponds to one co-culture. If that co-culture receives several incoming channels $k = 1, \dots, N$, then each channel contributes to pooled red and pooled green fluorescence. The experimentally measured transfer function of channel $k$ is

$$T_k(C) = y_{\min,k} + \left(y_{\max,k} - y_{\min,k}\right)\frac{C^{n_k}}{K_k^{n_k} + C^{n_k}},$$

where $C$ is the inducer concentration in $\mu$M, and $y_{\min,k}$, $y_{\max,k}$, $K_k$, and $n_k$ are the usual Hill parameters.

If channel $k$ is induced at concentration $C_k$, then the mixed co-culture contributes approximately

$$R_k \propto w_k T_k(C_k), \qquad G_k \propto (1 - w_k)T_k(C_k)$$

to the pooled red and green signals. Summing over all incoming channels gives

$$\Sigma R = \sum_k w_k\, T_k(C_k), \qquad \Sigma G = \sum_k (1 - w_k)T_k(C_k).$$

The pooled observable used by the decoder is

$$\rho = \frac{\Sigma G}{\Sigma R}.$$

Using the notation of Supplementary Note 8, the ratiometric decoding uses the same functional form but with effective parameters that account for pooling across multiple memregulons:

$$f(\rho) = \frac{1 - \gamma^{**}\rho}{1 - \lambda^{**}\beta^{**} + (\lambda^{**} - \gamma^{**})\rho}.$$

Here $f(\rho)$ is the decoded co-culture state in [0,1]. With

$$\lambda^{**} = \frac{\sum_{k=1}^{N} r_k}{\sum_{k=1}^{N} r_k\,/\lambda_k}, \qquad \beta^{**} = \frac{\sum_{k=1}^{N} r_k\, \beta_k}{\sum_{k=1}^{N} r_k}, \qquad \gamma^{**} = \frac{\sum_{k=1}^{N} (r_k/\lambda_k)\gamma_k}{\sum_{k=1}^{N} r_k\,/\lambda_k}.$$

We define the effective parameters $(\lambda^{**}, \beta^{**}, \gamma^{**})$ from saturating-inducer measurements, so they are constants for a given $N$-memregulon co-culture design. Here $(\lambda_k, \beta_k, \gamma_k)$ are the single-memregulon parameters defined in Supplementary Note 8, and $r_k$ is the maximal red fluorescence of incoming channel $k \in \{1, \dots, N\}$ at saturating inducer for the P1-only plasmid reference,

$$r_k = T_k(C_{k,\max}).$$

These expressions reduce to the one-edge parameters in the single-channel limit.

The decoder is monotone decreasing because $f(\rho)$ is a linear-fractional transform of $\rho$. It satisfies two anchor identities,

$$f(\beta^{**}) = 1, \qquad f\left(\frac{1}{\gamma^{**}}\right) = 0,$$

and its derivative has constant sign on its domain:

$$\frac{\mathrm{d}f}{\mathrm{d}\rho} = \frac{\lambda^{**}(\gamma^{**}\beta^{**} - 1)}{[1 - \lambda^{**}\beta^{**} + (\lambda^{**} - \gamma^{**})\rho]^2}.$$

Because $\lambda^{**} > 0$ and the denominator is strictly positive whenever $f$ is defined, the sign of $\mathrm{d}f/\mathrm{d}\rho$ is the sign of $(\gamma^{**}\beta^{**} - 1)$. For the parameter regime used here, $\beta^{**} < 1/\gamma^{**}$ and therefore $\gamma^{**}\beta^{**} < 1$, which implies $\mathrm{d}f/\mathrm{d}\rho < 0$. Thus $f(\rho)$ is strictly decreasing. Because $f$ is continuous and strictly decreasing, it follows that

$$0 \le f(\rho) \le 1 \qquad \text{for} \qquad \beta^{**} \le \rho \le \frac{1}{\gamma^{**}}.$$

The linear-mixture model used for pooling constrains each incoming channel to contribute a red/green ratio between its two anchor ratios, and pooling preserves these bounds, so the measured co-culture ratio $\rho = \Sigma G/\Sigma R$ remains within the interval above. Therefore, the decoded state remains in $[0,1]$.

The decoded state is still not a chemical concentration. To communicate with the next layer, the operator must convert that state into the concentration of the outgoing inducer channel $i$. If the normalised target activity is $a \in [0,1]$, then the inverse Hill relation is

$$C_i = K_i\left(\frac{a}{1-a}\right)^{1/n_i},$$

with clipping to the experimentally characterised range $[0, \mathrm{Act}_{i,\max}]$. The activation function therefore outputs an inducer concentration (in $\mu$M) on channel $i$:

$$\mathrm{Act}_i(\Sigma R, \Sigma G) = K_i\left(\frac{1 - \gamma^{**}\rho}{\lambda^{**}(\rho - \beta^{**})}\right)^{1/n_i}, \qquad \rho = \frac{\Sigma G}{\Sigma R}.$$

In practice, the code combines these two steps:

1. read $(\Sigma R, \Sigma G)$ and decode one neuron state $f(\rho)$;
2. convert that state into the output inducer concentration $\mathrm{Act}_i(\Sigma R, \Sigma G)$ for the neuron's assigned channel.

##### Output channels used by the activation function

The output channel of a neuron determines which inverse Hill parameters $(K_i, n_i, \mathrm{Act}_{i,\max})$ are used when that neuron's state is converted into a real inducer concentration. The nine validated output channels used by the simulator are listed in Table 1.

Output-channel parameters used by the activation function

| Id | Inducer | Promoter | $Act_{max}$ ($\mu$M) | $K$ ($\mu$M) | $n$ |
|---|---|---|---|---|---|
| 1 | Sal | PSal | 100 | 43.0 | 1.8 |
| 2 | aTc | PTet | 0.2 | 0.013 | 3.8 |

| | | | | | |
|---|---|---|---|---|---|
| 3 | Cho | PBetI | 10000 | 4100.0 | 2.7 |
| 4 | Ara | PBAD | 4000 | 37.0 | 1.5 |
| 5 | OC6 | PLux | 10 | 0.12 | 1.8 |
| 6 | OHC14 | PCin | 10 | 0.43 | 2.3 |
| 7 | Nar | PTtg | 1000 | 95.0 | 1.9 |
| 8 | Van | PVan | 100 | 26.0 | 2.3 |
| 9 | IPTG | PTac | 1000 | 140.0 | 1.8 |

**Supplementary Table 82 | Output-channel parameters used by the activation function.**

One-stage WTA proof of principle with a combinatorial promoter

The first architecture is a one-stage WTA readout with an explicit physical combinatorial promoter channel. It receives three ordinary incoming channels and one combinatorial incoming channel:

- a constant bias,
- $x_1$,
- $x_2$,
- and one genuine combinatorial promoter channel, denoted $x_{\text{combo}}$, which responds to the joint presence of two inducers.

This arrangement is biologically acceptable because the combinatorial promoter is a real physical regulatory channel. It is not a hidden multiplication performed in software. The simulator searches over 100 random seeds and retains the highest-margin valid XOR solution found during that search.

##### Multilayer network for negative-only learning

The second architecture is the biologically valid multilayer network used for negative-only learning. It contains:

- two logical inputs: $x_1$ and $x_2$,
- one constant bias term,
- four hidden co-cultures,
- and two output co-cultures representing labels 0 and 1.

The wiring is:

- each hidden co-culture receives bias through PLux, $x_1$ through PTtg, and $x_2$ through PVan,
- the hidden co-cultures communicate forward through PSal, PTet, PBetI, and PBAD,
- each output co-culture receives the constant bias plus all four hidden outputs.

##### Inference rule, winner-take-all, accuracy, and cross-entropy

For one input pattern $x = (x_1, x_2)$, the network produces two output values, $o_0$ and $o_1$.

The biological classification rule is winner-take-all (WTA):

$$\hat{y} = \begin{cases} 0 & \text{if } o_0 > o_1, \\ 1 & \text{if } o_1 > o_0. \end{cases}$$

This rule determines whether the network answered correctly.

For reporting and for training diagnostics, the two output values are also converted into a probability-like number by a two-class softmax:

$$\hat{p}(y = 1 \mid x) = \frac{\exp(o_1/\tau)}{\exp(o_0/\tau) + \exp(o_1/\tau)}.$$

where $\tau$ is a small temperature. This softmax does not add a biological layer; it only converts two output values into one number between 0 and 1 so that a standard loss can be computed.

Truth-table accuracy is defined by

$$\text{Accuracy} = \frac{1}{N}\sum_{k=1}^{N} \mathbf{1}\,\{\hat{y}_k = y_k\},$$

where $N = 4$ for the XOR truth table.

Binary cross-entropy is defined by

$$CE(y, p) = -(y \log p + (1 - y)\log(1 - p)).$$

In this note, cross-entropy is the main prediction-error quantity shown in the convergence figure because it is standard in machine learning, directly interpretable, and decreases towards zero as the truth-table prediction improves. An earlier draft used a more specialised quantity, running-best excess cross-entropy, which measured the distance from the best XOR-consistent state encountered so far. That quantity proved useful internally but less transparent for a broad readership. For that reason, Fig. 4d shows ordinary binary cross-entropy.

**Negative Local Learning Rule**

The negative-learning rule is implemented directly as a local negative biological update.

**Step 1: run inference.** The operator presents one truth-table input and applies WTA to the two output co-cultures.

**Step 2: apply the negative update to the wrong winning output co-culture if the winner is incorrect.** Each incoming edge $i \to j$ of that wrong output co-culture is updated by

$$w_{ij} \leftarrow \text{clip}_{[0,1]}\big(w_{ij} - \eta a_i w_{ij}\big),$$

where $a_i$ is the activity measured on that edge during the same inference step and $\eta$ is the learning rate.

This rule has the intended local interpretation:

- if an edge was inactive, then $a_i \approx 0$ and the update is negligible;
- if an edge was strongly active, it receives a stronger negative update;
- if a weight is already small, its absolute change is small because the update is proportional to $w_{ij}$ itself;
- the update is always negative, so the rule can only prune red-memory fraction and cannot create a missing pathway.

**Step 3: deep negative update of the hidden layer. In the same biological update cycle, the same local negative rule is applied to all hidden co-cultures. The final implementation does not use stochastic subsampling of hidden wells.**

**Step 4: no rollback.** The update is applied directly to the current state. The code does not create a trial network and does not accept or reject a candidate update.

**Suitable Start for Multilayer Negative Learning**

Because the rule is purely negative, it cannot create a missing pathway by increasing a weight. The relevant biological question is therefore whether a suitable near-solution can be refined by pruning. Specifically, if the starting network is already an OR-like near-solution, negative local learning can prune the remaining incorrect alternative and reach XOR.
The fixed start used here has accuracy 0.75 before learning: it classifies three truth-table rows correctly and only the (1,1) row incorrectly.

#### One-stage WTA XOR with a combinatorial promoter

The one-stage WTA search found many valid XOR solutions. For Fig. 4a,b, the representative architecture and truth-table readout were chosen as the highest-margin valid XOR solution returned by the 100-seed search. Its deterministic truth table is perfect, its binary cross-entropy is essentially zero, and its mean gate probability is high.

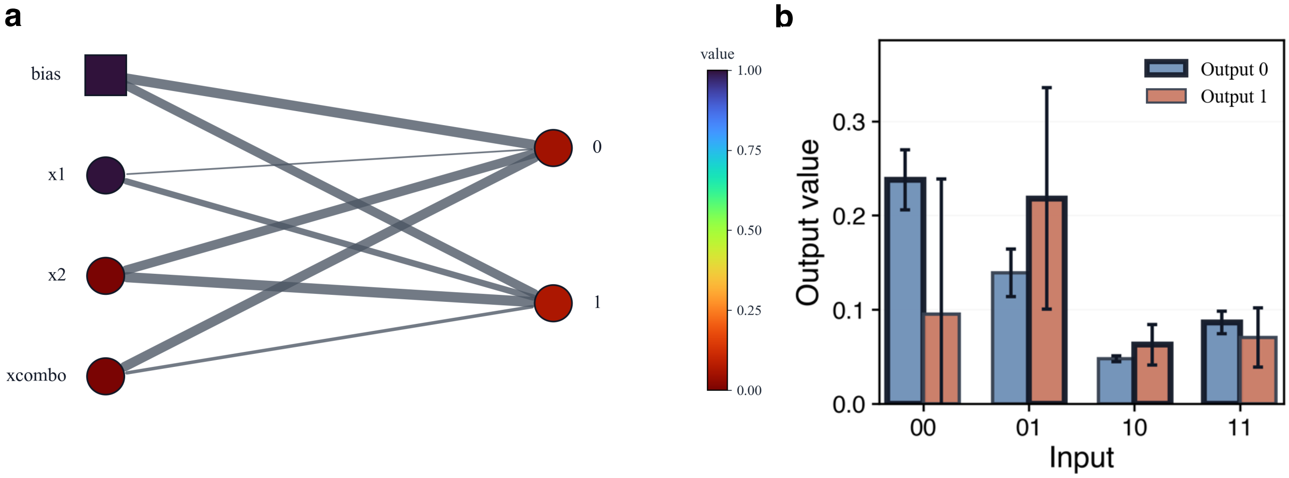

**Supplementary Fig. 89 | One-stage WTA XOR with one combinatorial promoter channel. a**, One-stage WTA combinatorial XOR readout corresponding to main Fig. 4**a**,b. The visible inputs are the constant bias, x1, x2, and the explicit combinatorial channel x1 AND x2. Edge thickness is proportional to the exact simulated weight, and node colour is proportional to the exact node value on the global 0-1 scale for the displayed input condition. **b**, Output values assuming weight-error propagation. Two bars are shown for each input condition, one bar per output neuron. The bar heights are the deterministic output values of the selected representative readout. Error bars are standard deviations obtained by Monte Carlo propagation of the measured promoter-specific weight uncertainty. The thicker bar outline marks the winner-take-all decision.

This architecture matters because it shows that XOR does not require a hidden layer if one physical incoming channel is itself combinatorial. The non-linearity can therefore be implemented in promoter logic rather than in network depth.

Under Monte Carlo propagation of the measured weight uncertainty, the deterministic class ordering remains visible, but the confidence is not equally strong for all four rows. The estimated correct-label fractions are approximately 0.905 for input 00, 0.883 for 01, 0.830 for 10, and 0.694

for 11. Thus the one-stage WTA combinatorial solution is markedly more robust than the multilayer learned solution under the same uncertainty model, although it is not perfectly separated when weight noise is propagated independently across all synapses.

Extended Fig. 4a | Monte Carlo output distributions for the single-layer combinatorial XOR

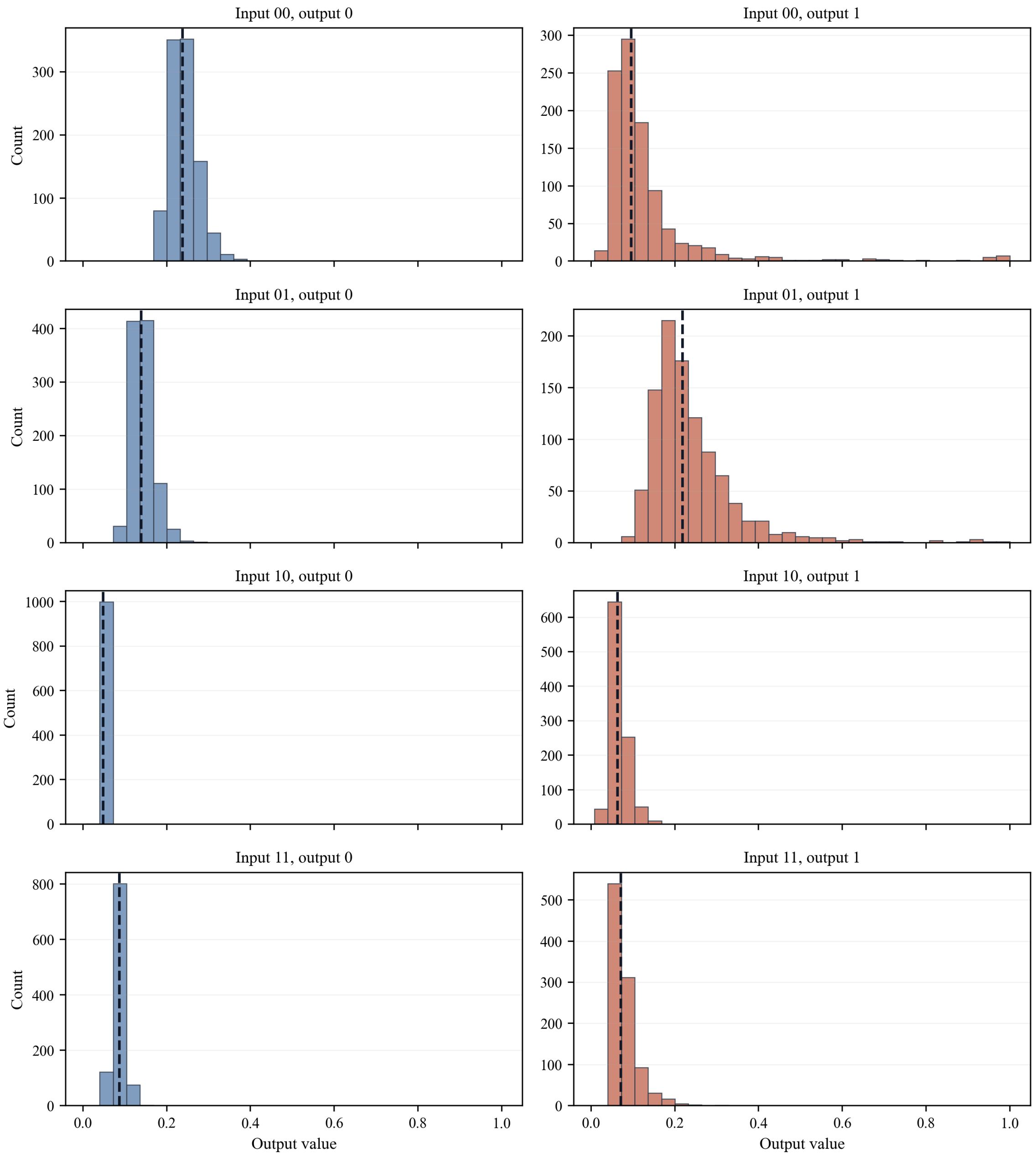

**Supplementary Fig. 90 | Monte Carlo output distributions for the one-stage WTA combinatorial XOR readout.** Eight histogram panels are shown, one for each pair of input condition and output neuron. The first 1000 Monte Carlo draws are shown so the distributions are smooth and easy to inspect. In each panel, the dashed line marks the deterministic output value of the representative XOR architecture/readout shown in Fig. 4**a**,b.

#### Human-in-the-loop multilayer scalability analysis

##### Negative-only Convergence from the OR-like Start

The multilayer negative-learning reference run starts from the fixed OR-like state described above and applies the direct negative-update rule with learning rate $\eta = 0.01$. In the deterministic reference run used for the convergence plot, XOR is reached after 10 epochs and 134 direct weight changes. Final deterministic performance is perfect (Accuracy = 1.0), with mean gate probability 0.9837 and cross-entropy 0.0166.

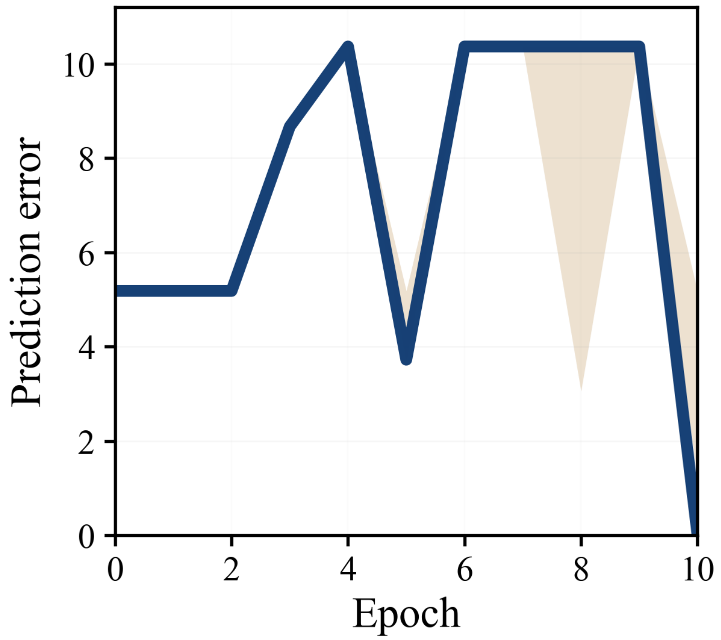

**Supplementary Fig. 91 | Convergence of negative-only pruning in the multilayer network.** The vertical axis is labelled as prediction error for readability. Here prediction error is the binary cross-entropy of the simulated WTA readout, which measures how strongly the network assigns probability to the correct XOR output: smaller values are better, and zero means a perfect prediction. This is a task-level simulation metric, not evidence that the biological implementation can run a general error-minimising optimisation algorithm. The dark line is the single convergent reference run. The light band shows the 10th to 90th percentile range of the non-convergent runs over the same epoch window.

The curve is not monotone. That is expected. Negative-only learning can temporarily worsen one truth-table row while it is pruning another. Nevertheless, the final deterministic XOR is reached without any positive update and without any accept-reject logic.

Figure 4c shows the human/operator/computational workflow for the multilayer scalability analysis. Weight updates are physically implemented in the model and constrained by measured memregulon dynamics, whereas inter-layer communication, inducer routing and WTA handoff are externally supplied by the operator/computational workflow. Edge thickness is proportional to the exact final weight, and node colour is proportional to the exact node value on the global 0 to 1 scale. The bias is shown as a square because it is a constant source term rather than a measured input well.

To make the visible colours more informative, the network is shown for the asymmetric input condition $(x_1, x_2) = (0,1)$ rather than for $(1,1)$.

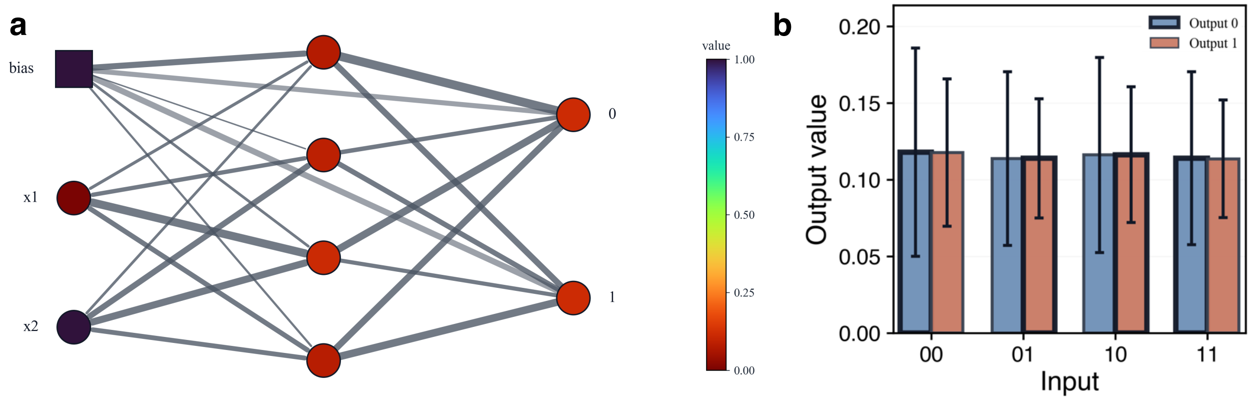

**Supplementary Fig. 92 | Human-in-the-loop multilayer scalability analysis after negative-only learning. a**, The figure shows the simulated architecture for the asymmetric input condition. Hidden nodes are intentionally left unlabeled in the drawing, but all node identities, node values, edge weights and rendered edge thicknesses are listed in the supplementary tables below. **b**, Output/readout values with propagated weight uncertainty. These are deterministic truth-table outputs of the final multilayer scalability-analysis state together with error bars produced by Monte Carlo propagation of experimentally measured weight uncertainty.

This is not a fully autonomous biological multilayer neural network. The architecture is a human-in-the-loop multilayer scalability analysis with non-negative memregulon weights constrained to [0,1]. The nonlinear readout arises from calibrated ratiometric co-culture decoding and the WTA decision rule. Weight updates are physically implemented and constrained by measured memregulon dynamics, whereas inter-layer communication is externally supplied by the human/operator/computational workflow; the analysis does not implement self-contained inter-layer signalling.

Two bars in Supplementary Fig. 92b are shown for each input condition, one per output neuron. The bar heights are the deterministic output values of the learned network. Error bars are standard deviations obtained by Monte Carlo propagation of the measured promoter-specific weight uncertainty. The thicker bar outline marks the winner-take-all decision. This figure requires careful interpretation. Deterministically, the network is correct on all four XOR rows. However, under the deliberately conservative uncertainty model used here, each synaptic weight is sampled independently around its measured promoter-specific uncertainty. Under that assumption, the class-confidence fractions are close to 0.5 for all four rows, approximately 0.521, 0.494, 0.479 and 0.511. The deterministic multilayer solution therefore exists, but its output margins remain small compared with the propagated experimental weight variability. This result does not invalidate the deterministic negative-learning solution. Rather, it shows that a multilayer XOR solution is reachable by negative-only learning from a suitable start, while also indicating that tighter experimental control of weight variability will be needed for statistically reliable biological execution.

The main-text panel reports the deterministic correct-output advantage, defined as the target-output score minus the non-target-output score. This representation shows the sign and size of the deterministic WTA decision margin directly. The Monte Carlo analysis is reported separately as correct-label fractions and output distributions, because propagating the experimentally measured weight uncertainty gives broad output distributions relative to the small deterministic margins.

Extended Fig. 4d | Monte Carlo output distributions for the learned multilayer XOR

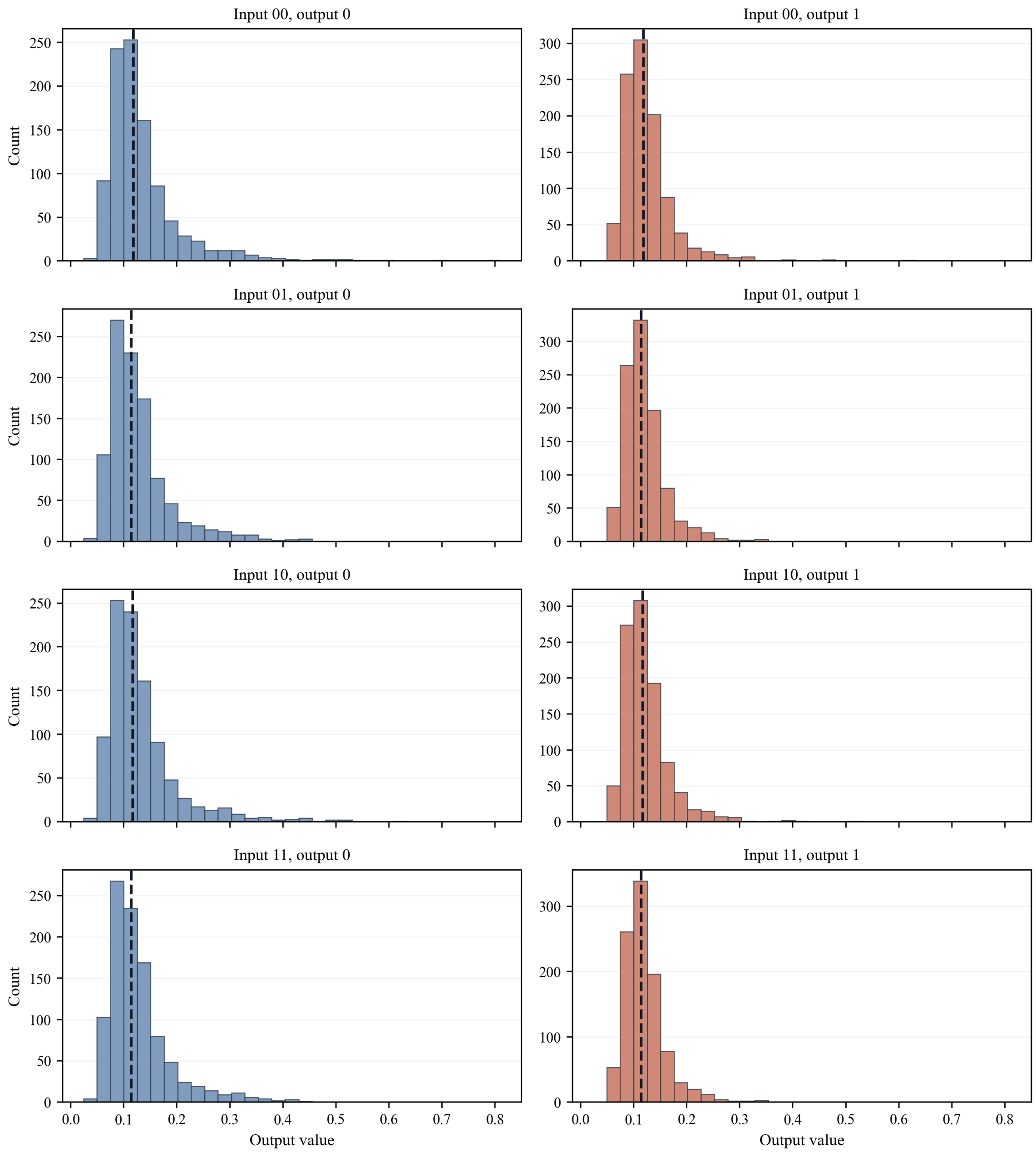

**Supplementary Fig. 93 | Monte Carlo output distributions for the learned multilayer XOR. a**, Eight histogram panels are shown, one for each pair of input condition and output neuron. The first 1000 Monte Carlo draws are shown so the shapes of the sampled output distributions can be inspected directly. In each panel, the dashed line marks the deterministic output value of the learned XOR WTA readout shown in Fig. 4e.

#### Promoters, weights, and standard deviations used in the displayed networks

Because the purpose of these simulations is to mirror what a human experimenter would measure and apply in a real human-in-the-loop workflow, it should be explicit which memregulon promoter was assigned to each edge, which weight value was used, and which standard deviation was propagated for that edge. The two tables below list that information for the displayed one-stage WTA combinatorial XOR readout and for the displayed learned multilayer XOR network. In every case, the reported standard deviation is the value actually used by the Monte Carlo propagation, obtained by interpolation from the corresponding promoter-specific experimental uncertainty estimates.

The companion truth-table tables below provide the corresponding displayed node values, WTA labels, correct-output advantages and Monte Carlo correct-label fractions for the displayed networks.

#### One-stage WTA combinatorial XOR readout

| Source | Target | Edge promoter | Inducer | Deterministic Edge Weight (W) | Propagated SD | Rendered Edge Thickness |
|---|---|---|---|---|---|---|
| *bias* | y0 | PLux | OC6 | 0.94045 | 0.00936 | 6.97 |
| *x1* | y0 | PTtg | Nar | 0.12091 | 0.04035 | 1.20 |
| *x2* | y0 | PVan | Van | 0.86569 | 0.02688 | 6.44 |
| *xcombo* | y0 | PVan-Ttg | Van+Nar | 0.90441 | 0.12050 | 6.71 |
| *bias* | y1 | PLux | OC6 | 0.84105 | 0.07679 | 6.27 |
| *x1* | y1 | PTtg | Nar | 0.58198 | 0.04090 | 4.44 |
| *x2* | y1 | PVan | Van | 0.97351 | 0.02688 | 7.20 |
| *xcombo* | y1 | PVan-Ttg | Van+Nar | 0.29043 | 0.33202 | 2.39 |

**Supplementary Table 83 | One-stage WTA combinatorial XOR readout.** Source is the upstream input or nonlinear feature, target is the output readout, edge promoter identifies the memregulon promoter assigned to that connection, and inducer gives the chemical input used for activation. Deterministic edge weight is the displayed simulated weight, propagated SD is the promoter-specific uncertainty used for Monte Carlo sampling, and rendered edge thickness is the visual line-width scale.

| Input x1 | Input x2 | AND feature value | y0 | y1 | Target Label | WTA Label | Correct-output advantage | Monte Carlo correct-label fraction |
|---|---|---|---|---|---|---|---|---|
| *0* | 0 | 0 | 0.23770 | 0.09545 | 0 | 0 | 0.14225 | 0.9052 |
| *0* | 1 | 0 | 0.13905 | 0.21811 | 1 | 1 | 0.07906 | 0.8830 |
| *1* | 0 | 0 | 0.04798 | 0.06280 | 1 | 1 | 0.01483 | 0.8304 |
| *1* | 1 | 1 | 0.08645 | 0.07059 | 0 | 0 | 0.01586 | 0.6944 |

**Supplementary Table 83b | One-stage WTA combinatorial XOR truth-table values.** Input x1 and input x2 are the Boolean truth-table states. AND feature value is the physical nonlinear feature supplied by the combinatorial promoter. y0 and y1 are deterministic output scores, target label is the XOR class, WTA label is the selected class, correct-output advantage is the target-output score minus the non-target-output score, and Monte Carlo correct-label fraction is the fraction of sampled networks that retained the correct WTA label.

#### Learned multilayer XOR network

| Source | Target | Edge promoter | Inducer | Deterministic Edge Weight (W) | Propagated SD | Rendered Edge Thickness |
|---|---|---|---|---|---|---|
| *bias* | h1 | PLux | OC6 | 0.87613 | 0.05299 | 5.05 |
| *x1* | h1 | PTtg | Nar | 0.77859 | 0.03704 | 2.52 |
| *x2* | h1 | PVan | Van | 0.76631 | 0.01962 | 2.20 |
| *bias* | h2 | PLux | OC6 | 0.72763 | 0.15375 | 1.20 |
| *x1* | h2 | PTtg | Nar | 0.81809 | 0.03704 | 3.54 |
| *x2* | h2 | PVan | Van | 0.86810 | 0.02688 | 4.84 |
| *bias* | h3 | PLux | OC6 | 0.76783 | 0.12647 | 2.24 |
| *x1* | h3 | PTtg | Nar | 0.94025 | 0.03704 | 6.71 |
| *x2* | h3 | PVan | Van | 0.90651 | 0.02688 | 5.84 |
| *bias* | h4 | PLux | OC6 | 0.75102 | 0.13788 | 1.81 |
| *x1* | h4 | PTtg | Nar | 0.83939 | 0.03704 | 4.10 |
| *x2* | h4 | PVan | Van | 0.83378 | 0.02688 | 3.95 |
| *bias* | y0 | PLux | OC6 | 0.84106 | 0.07679 | 4.14 |
| *h1* | y0 | PSal | Sal | 0.95915 | 0.10054 | 7.20 |
| *h2* | y0 | PTet | aTc | 0.82251 | 0.01176 | 3.66 |
| *h3* | y0 | PBetI | Cho | 0.91292 | 0.03277 | 6.00 |
| *h4* | y0 | PBAD | Ara | 0.88920 | 0.00316 | 5.39 |
| *bias* | y1 | PLux | OC6 | 0.87217 | 0.05568 | 4.95 |
| *h1* | y1 | PSal | Sal | 0.87669 | 0.10054 | 5.06 |
| *h2* | y1 | PTet | aTc | 0.84649 | 0.01176 | 4.28 |
| *h3* | y1 | PBetI | Cho | 0.81191 | 0.05901 | 3.38 |
| *h4* | y1 | PBAD | Ara | 0.92223 | 0.00316 | 6.24 |

**Supplementary Table 84 | Learned multilayer XOR network.** Source is the upstream bias, input or hidden node, target is the downstream hidden or output readout, edge promoter identifies the memregulon promoter assigned to that connection, and inducer gives the chemical input used for activation. Deterministic edge weight is the displayed simulated weight after negative-only learning, propagated SD is the promoter-specific uncertainty used for Monte Carlo sampling, and rendered edge thickness is the visual line-width scale.

| *Input x1* | *Input x2* | *h1* | *h2* | *h3* | *h4* | *y0* | *y1* | *Target Label* | *WTA Label* | *Correct-output advantage* | *Monte Carlo correct-label fraction* |
|---|---|---|---|---|---|---|---|---|---|---|---|
| *0* | 0 | 0.12478 | 0.05103 | 0.06258 | 0.05726 | 0.11799 | 0.11773 | 0 | 0 | 0.00026 | 0.5206 |
| *0* | 1 | 0.07475 | 0.08640 | 0.11326 | 0.07813 | 0.11369 | 0.11392 | 1 | 1 | 0.00023 | 0.4938 |
| *1* | 0 | 0.10205 | 0.05764 | 0.08175 | 0.06494 | 0.11616 | 0.11631 | 1 | 1 | 0.00015 | 0.4788 |
| *1* | 1 | 0.07375 | 0.08604 | 0.12061 | 0.07970 | 0.11400 | 0.11355 | 0 | 0 | 0.00045 | 0.5106 |

**Supplementary Table 84b | Learned multilayer XOR network truth-table values.** Input x1 and input x2 are the Boolean truth-table states. h1-h4 are deterministic hidden-node values, y0 and y1 are deterministic output scores, target label is the XOR class, WTA label is the selected class, correct-output advantage is the target-output score minus the non-target-output score, and Monte Carlo correct-label fraction is the fraction of sampled networks that retained the correct WTA label.

Monte Carlo Uncertainty Propagation and Correct-Label Fraction
For each synapse, the Monte Carlo procedure is:

1. take the deterministic simulated weight as the mean,
2. assign that weight a promoter-specific standard deviation by interpolation within the corresponding measured sheet,
3. draw a random weight from a clipped normal distribution on $[0,1]$,
4. run the whole truth table with all sampled synapses,
5. repeat this many times (5000 samples in the figures).

The error bars in the one-stage WTA and multilayer truth-table panels are the standard deviations of the sampled output values. The bar heights themselves are not Monte Carlo means; they are the deterministic output values of the displayed network. This choice keeps the winner-take-all decision visually tied to the actual simulated solution while still reporting how much that solution spreads when the measured weight variability is propagated.

Because standard-deviation error bars can hide the actual shape of a broad or skewed distribution, this note also includes the corresponding histogram figures for the first 1000 Monte Carlo draws of each network. For each input row, the workbook also stores a Monte Carlo correct-label fraction: the fraction of Monte Carlo samples whose WTA label matches the target label. Because this note neglects uncertainty in the enforced input values, that fraction reflects only the propagated uncertainty in the weights.

**Robustness across 100 shuffle seeds**

The negative-only multilayer result is robust in a narrow deterministic sense because the fixed-start reference run converges, but it does not converge for every truth-table presentation order. To quantify that dependence, the same OR-like suitable start was rerun over 100 different shuffle seeds. Only the sample order was changed; the starting weights were identical in all 100 runs. The results were:

| ***Quantity*** | ***Value*** |
|---|---|
| Total shuffle seeds tested | 100 |
| Converged runs | 45 |
| Convergence rate | 45.0% |
| % Wilson interval | [35.6%, 54.8%] |
| One-sided exact binomial $p$-value versus $1/16$ | $1.25 \times 10^{-27}$ |
| One-sided exact binomial $p$-value versus 0.5 | 0.864 |
| Mean gate probability among converged runs | $0.971 \pm 0.034$ |
| Mean cross-entropy among converged runs | $0.031 \pm 0.038$ |

**Supplementary Table 85 | Robustness across 100 shuffle seeds.**

Two conclusions follow directly. First, the success rate is far higher than a naive random-baseline probability, so the method is not behaving like blind chance. Second, the success rate does not exceed 50% under this test, so the present negative-only protocol remains sensitive to sample order even from the same suitable start. The main deterministic claim of the note should therefore remain precise: negative-only pruning can reach XOR from a biologically suitable start, but it is not an order-insensitive optimiser and not an autonomous biological reinforcement-learning system.1/16

.

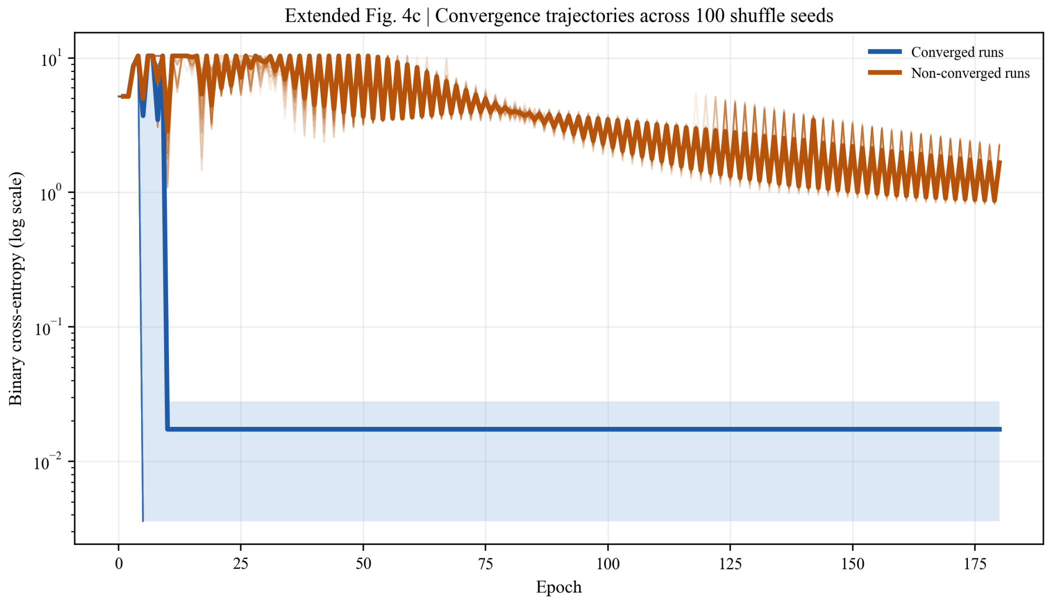

**Supplementary Fig. 94 | Individual convergence trajectories across 100 shuffle seeds.** Each thin line is one run of negative-only learning from the same OR-like suitable start, with only the truth-table presentation order changed. The vertical axis is logarithmic so that both early high-error epochs and late near-zero converged epochs remain visible. Thick lines and shaded bands show the median and interquartile range for converged and non-converged runs separately.
